# Contrasting genomic routes to domestication in Occidental and Oriental pears

**DOI:** 10.1101/2025.11.17.687327

**Authors:** Yuqi Nie, Xilong Chen, Somia Saidi, Yann Bourgeois, Anthony Venon, Johann Confais, Aurélie Mesnil, Shuo Liu, Yuhao Gao, Junbei Ni, Songling Bai, Lijuan Gao, Guofang Li, Monika Hoefer, Nicolas Francillonne, Anamaria Roman, Lehel Lukács, Caroline Denance, Takashi Akagi, Anush Nersesyan, Astghik Papikyan, Ivan George Gabrielyan, Hamid Abdollahi, Karine Alix, Yuanwen Teng, Amandine Cornille

**Affiliations:** Université Paris-Saclay, INRAE, CNRS, AgroParisTech, GQE – Le Moulon, 91190 Gif-sur-Yvette, France; Division of Science, New York University Abu Dhabi, Saadiyat Island, Abu Dhabi, United Arab Emirates; DIADE, University of Montpellier, CIRAD, IRD, 34394, Montpellier, France; Université Paris-Saclay, INRAE, BioinfOmics, URGI, 78026 Versailles, France; Liaoning Institute of Pomology, Yingkou City, Liaoning 115009, China; College of Agriculture and Biotechnology, Zhejiang University, Hangzhou, Zhejiang 310058, China; The Key Laboratory of Horticultural Plant Growth, Development and Quality Improvement, Ministry of Agriculture of China, Hangzhou, 310058, Zhejiang, China; Hainan Institute of Zhejiang University, Sanya, Hainan 572000, China; Changli Institute of Pomology, Hebei Academy of Agriculture and Forestry Sciences, Changli 066600, Hebei, China; College of Horticulture, Hebei Agricultural University, Hebei 071000, China; Institute of Breeding Research on Fruit Crops, Federal Research Centre for Cultivated Plants (JKI), Dresden, Germany; Institute of Biological Research Cluj – National Institute for Research and Development in Biological Sciences, 400015, Cluj-Napoca, Romania; University of Agricultural Sciences and Veterinary Medicine, 400372, Cluj-Napoca, Romania; Univ Angers, Institut Agro, INRAE, IRHS, SFR QUASAV, F-49000 Angers, France; Statistical Genetics Team, RIKEN Center for Advanced Intelligence Project, Nihonbashi 1-chome Mitsui Building 15F, 1-4-1 Nihonbashi, Chuo-ku, Tokyo 103-0027, Japan; Bioorganic Research Institute, Suntory Foundation for Life Sciences, Seika-dai 8-1-1, Seika-cho, Kyoto 619-0284, Japan; A.Takhtajan Institute of Botany NAS RA, Achryan Street 1, 0063 Yerevan, Armenia; Nature Heritage NGO, Luxemburg Street 9-38, 0090 Yerevan, Armenia; Temperate Fruits Research Center, Horticultural Sciences Research Institute, Agricultural Research, Education and Extension Organization (AREEO), Karaj, Iran

**Author notes:** Corresponding authors: Karine ALIX, Xilong CHEN, Yuanwen TENG, and Amandine CORNILLE.

**Keywords:** Pear, domestication, population genetics, demographic history, gene flow, positive selection, genetic load, transposable elements, perennial fruit crops

## Abstract

Despite their global importance, the genomic foundations of domestication in perennial fruit trees remain incompletely integrated across demography, selection, and structural variation. Pear (*Pyrus* spp.), domesticated in Europe and East Asia, provides a powerful system to compare alternative evolutionary trajectories within a balanced genomic framework. Using lineage-specific reference genomes, we analyzed nucleotide and transposable element (TE) polymorphisms across 396 well-documented wild and cultivated individuals spanning both regions. We uncovered weak domestication bottlenecks and extensive wild–crop gene flow, consistent with a protracted evolutionary process. In Europe, dessert and perry *P. communis* diverged independently based on fruit use. In East Asia, *P. pyrifolia* diversified geographically in China and Japan, following distinct domestication trajectories. Selection scans identify lineage- and population-specific targets associated with fruit texture, metabolism, and immunity. We find no support for the cost-of-domestication hypothesis: cultivated pears show a lower realized burden of deleterious mutations than their wild relatives. TE insertion patterns largely reflect demographic history, and are not enriched near the genes under positive selection. Together, our results support a protracted, multi-regional model of pear domestication shaped by recurrent gene flow and lineage-specific genomic architectures.

## Introduction

Plant domestication represents one of the most profound evolutionary transitions in human history, providing a natural experiment to study how selection, gene flow, and demography shape adaptation on short evolutionary timescales^1–3^. Researchers have studied domestication in annual crops extensively, but less is known about its genomic consequences in long-lived perennial fruit tree species.

Perennials differ fundamentally from annuals in the way selection and demography interact during domestication. Unlike annual crops, which were shaped by rapid domestication and intensive, human-driven selection, perennial fruit trees, with their long generation times and outcrossing systems, maintain high heterozygosity, enabling extensive gene flow between their wild and cultivated forms^4–7^. As a result, perennial crops often experience weak or no domestication bottlenecks and frequently exhibit bidirectional introgression between wild and cultivated populations. These features make perennial fruit trees ideal systems for exploring how adaptive evolution proceeds under continuous gene flow and long-term selection. In several perennial models, including apple (*Malus domestica* Borkh.), grapevine (*Vitis vinifera* L.), olive (*Olea europaea* L.), and peach (*Prunus persica* (L.) Batsch), genes under selection during domestication have been linked to fruit development, ripening, and stress tolerance^8–13^. These studies have primarily focused on gene coding regions, whereas the role of non-coding regions, particularly transposable elements (TEs), mobile DNA sequences capable of reshaping gene regulation and chromatin structure^14^, remains largely unexplored. TEs can generate regulatory and structural diversity, influence epigenetic states, and create new cis-regulatory variants, but their population-level and evolutionary dynamics during domestication are poorly understood^15^. Investigating how TE insertion polymorphisms contribute to genome variation and adaptation is therefore essential to fully understand perennial plant domestication.

Among temperate fruit trees, pear (*Pyrus* L., Rosaceae) provides an excellent model for studying perennial evolution and domestication. The genus *Pyrus* comprises roughly 30 intercompatible wild species distributed across Eurasia, forming two major evolutionary lineages: the Occidental and Oriental pears, each comprising both wild and cultivated representatives^16–18^. The European pear (*Pyrus communis* L.) represents the main cultivated form of the Occidental lineage, while the cultivars of the Oriental lineage encompass several cultivated species (also called Asian pears)—*P. × sinkiangensis* T.T. Yu, *P. ussuriensis* Maxim., and *P. pyrifolia* (Burm. f.) Nakai. Among them, *P. pyrifolia* is the most widely cultivated and genetically diverse, comprising three major cultivar groups: the White Pear Group, Sand Pear Group, and Japanese Pear Group^19,20^. European and Asian pears differ markedly in their fruit morphology and use: European pears (*P. communis*) include dessert varieties, which typically ripen post-harvest, developing a soft, melting texture suitable for fresh consumption and specialized varieties used for fermentation (perry, an alcoholic beverage). In contrast, Asian pears (*P. pyrifolia*) bear crisp fruits that are mainly consumed fresh without post-harvest ripening^18,21,22^. Despite their agricultural importance, the domestication history of pears remains unresolved for the Occidental lineage because the wild relatives of European pears, *P. pyraster* (L.) Burgsd. and *P. caucasica* Fed., considered the most probable ancestors of the cultivated *P. communis*^23^, in population genomic studies^24^. In addition, while recent studies have greatly advanced the characterization of diversity among Oriental pears^25–27^, their analyzed samples were heavily biased toward Asian cultivars and lacked field-verified wild populations (e.g., *Pyrus pashia,* Buch.-Ham. ex D. Don, the Himalayan pear)^17,28^, limiting inferences about the independent origins, demographic histories, and comparative selective pressures experienced by these two lineages, and the role of TEs in shaping their genomes.

Here, we investigated the demographic and adaptive processes that shaped the genomes of the two major cultivated pear lineages, aiming to reconstruct the tempo and mode of pear domestication. Specifically, we asked (i) how many distinct gene pools exist among Occidental and Oriental pears, and how they genetically relate to local wild relatives; (ii) whether cultivated populations diverged independently from wild gene pools within each lineage; (iii) whether parallel selection targeted shared functional pathways or distinct genetic routes; (iv) whether TE insertion polymorphism follows the demographic history inferred from SNPs or is associated with genes under positive selection. We addressed these questions by combining new and publicly available whole-genome resequencing data from 396 non-clonal individuals spanning Eurasia. Using nucleotide and transposable element insertion polymorphisms (SNPs and TIPs, respectively) called from lineage-matched reference genomes for *P. communis* and *P. pyrifolia*, we reconstructed the population structure, demographic history, and genomic signatures of selection and genetic load. Our integrative analyses go beyond confirming the dual origins of *Pyrus* domestication to reveal contrasting demographic and adaptive pathways in Occidental and Oriental pears, illustrating independent adaptive trajectories in perennial crops, where long lifespan, self-incompatibility, and recurrent introgression have shaped distinctive genomic outcomes.

## Results

### Occidental and Oriental pears form distinct gene pools

A global analysis of 396 non-duplicate pear genomes revealed a strong genetic split between Occidental (incl. *P. communis*) and Oriental (incl. *P. pyrifolia*) lineages (Fig. 1a,b). Using ∼21.8K synonymous and unlinked SNPs derived from 17.7 M high-quality variants (Supplementary Table 1; Supplementary Figs. 1,2), fastSTRUCTURE, Neighbor-Net, and principal component analysis (PCA) consistently distinguished the two gene pools, with 136 of 396 individuals showing admixed ancestry (Fig. 1a–e; Supplementary Tables 2,3; Supplementary Figs. 3–8; Supplementary Note 1). The strongest genetic differentiation among Oriental pears suggests a longer or more complex divergence history.

**Figure 1.**
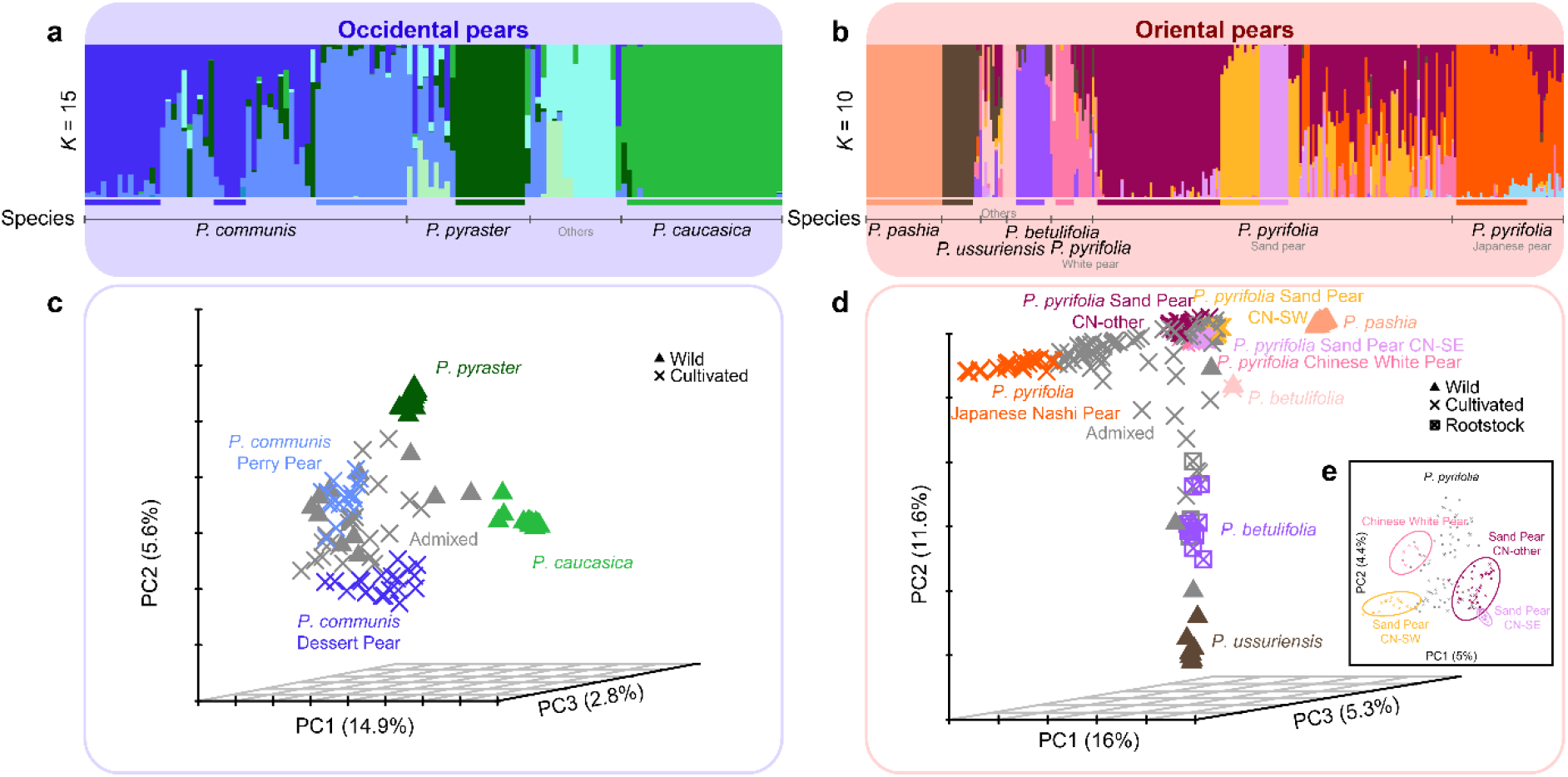
Population genetic structure, differentiation, and variation in Occidental and Oriental pears (*Pyrus* spp.). a,b. Population structure of wild and cultivated pears (*N* = 396 individuals) inferred with fastSTRUCTURE: Occidental pears (*N* = 130; 20,298 synonymous, unlinked SNPs) at *K* = 15 (a) and Oriental pears (*N* = 266; 21,141 synonymous, unlinked SNPs) at *K* = 10 (b). **c,d** Principal component analysis (PCA) of 113 Occidental (20,263 synonymous and unlinked SNPs mapped onto *P. communis*) (c) and 256 Oriental individuals (21,163 synonymous and unlinked SNPs mapped onto *P. pyrifolia*) (d), after excluding 17 and 10 “other” individuals, respectively. **e** PCA plot of Chinese cultivated *P. pyrifolia* sand and white pears.

Because *P. communis* and *P. pyrifolia* were genetically distinct (Supplementary Figs. 3–15), we analyzed each lineage separately using its respective reference genome to minimize mapping bias (Supplementary Figs. 16–21). Within *P. communis*, dessert and perry pears formed distinct clusters (Fisher’s exact test *p* = 1.04 × 10^−7^), more closely related to wild *P. pyraster* than *P. caucasica* (Fig. 1a,c; Supplementary Fig. 23; Supplementary Note 1). Several admixed *P. pyraster* individuals likely reflect localized introgression or historical gene flow from nearby cultivated stands (Supplementary Figs. 9–13; Supplementary Table 3).

Among Oriental pears, we identified nine genetic clusters (*K* = 10; Fig. 1b,d,e; Supplementary Figs. 19–21 and 23), corresponding to three wild species and six cultivated groups. The three wild Oriental pear species in China, namely *P. pashia*, *P. ussuriensis*, and *P. betulifolia* (used as rootstocks), possess distinct gene pools. Among these, *P. pashia* exhibited the highest genetic purity, showing no signs of admixture with any cultivated pear genetic group. *P. ussuriensis*, distributed in northeastern China, showed relatively low levels of admixture with other Chinese cultivar pear gene pools. By contrast, *P. betulifolia* showed extensive admixture with Chinese cultivated pear gene pools, likely because it overlaps geographically with major cultivated pear production regions in China. Among cultivated pears, the white pear and Japanese pear gene pools showed strong regional differentiation. The white pear individuals surveyed here are mainly distributed in North China, the historical heartland of Asian white pear cultivation. The extensive admixture observed across the white and sand pear gene pools (Fig. 1b; Supplementary Fig. 22) indicates that our sampling effectively captures the broad spectrum of the *P. pyrifolia* complex. We found that CN-other sand pears, distributed across nearly all production regions of China, overlapping with the other two sand pear clusters, likely represent the ancestral cluster of all regionally differentiated *P. pyrifolia* sand pears (Fig. 1b). Among Japanese pears, approximately one-third of individuals are admixed, primarily with the Chinese cultivar cluster referred to as CN-other (Fig. 1b). The gene pool of sand pears in Eastern China (e.g., Zhejiang Province) showed substantial admixture with that of Japanese pears (Fig. 1b; Supplementary Fig. 22). The admixture patterns likely result from both the retention of common ancestral alleles and episodic gene flow, rather than solely intensive modern breeding over the last century^29^.

Overall, population-structure analyses revealed deep divergence and limited admixture between Occidental and Oriental pears, consistent with separate evolutionary histories of the two lineages. After excluding admixed and uncertainly assigned individuals (Fig. 1c,d; Supplementary Fig. 23) to minimize the effects of recent breeding-related hybridization and to remove potential escaped cultivars showing introgression into local wild populations, we retained 229 individuals representing 12 Occidental (four) and Oriental (eight) panmictic populations for demographic and selection analyses (Supplementary Table 4; Supplementary Note 1; Supplementary Fig. 23).

### Divergence and demographic histories of Oriental and Occidental pears

Evolutionary relationships among the 12 pear populations were inferred using SVDQuartets based on 21,237 unlinked synonymous SNPs mapped to the *P. pyrifolia* reference genome (Fig. 2a; Supplementary Table 4). Loquat (*Eriobotrya japonica*) was utilized as the outgroup^30,31^. The phylogenetic reconstruction revealed a deep split between Occidental and Oriental pears, confirming that these lineages evolved independently and underwent separate divergence histories (Fig. 2a; Supplementary Fig. 24). Within the Occidental clade, *P. caucasica* formed a distinct branch from *P. pyraster* and the European pear, *P. communis*. Nodes within the *P. pyraster–P. communis* clade exhibited moderate bootstrap support, suggesting incomplete lineage sorting or historical gene flow between wild and cultivated populations. In the Oriental lineage, *P. pashia* and *P. betulifolia* were clearly separated from the cultivated *P. pyrifolia* complex, although relationships among the Asian pears (white, sand, and Japanese pears) remained weakly resolved, likely reflecting recent divergence and recurrent introgression (see below). To further elucidate these relationships, we tested alternative divergence scenarios using demographic modeling. To do this, we first quantified historical gene flow among populations and reconstructed historical effective population size (*N*_e_) to inform model optimization.

**Figure 2.**
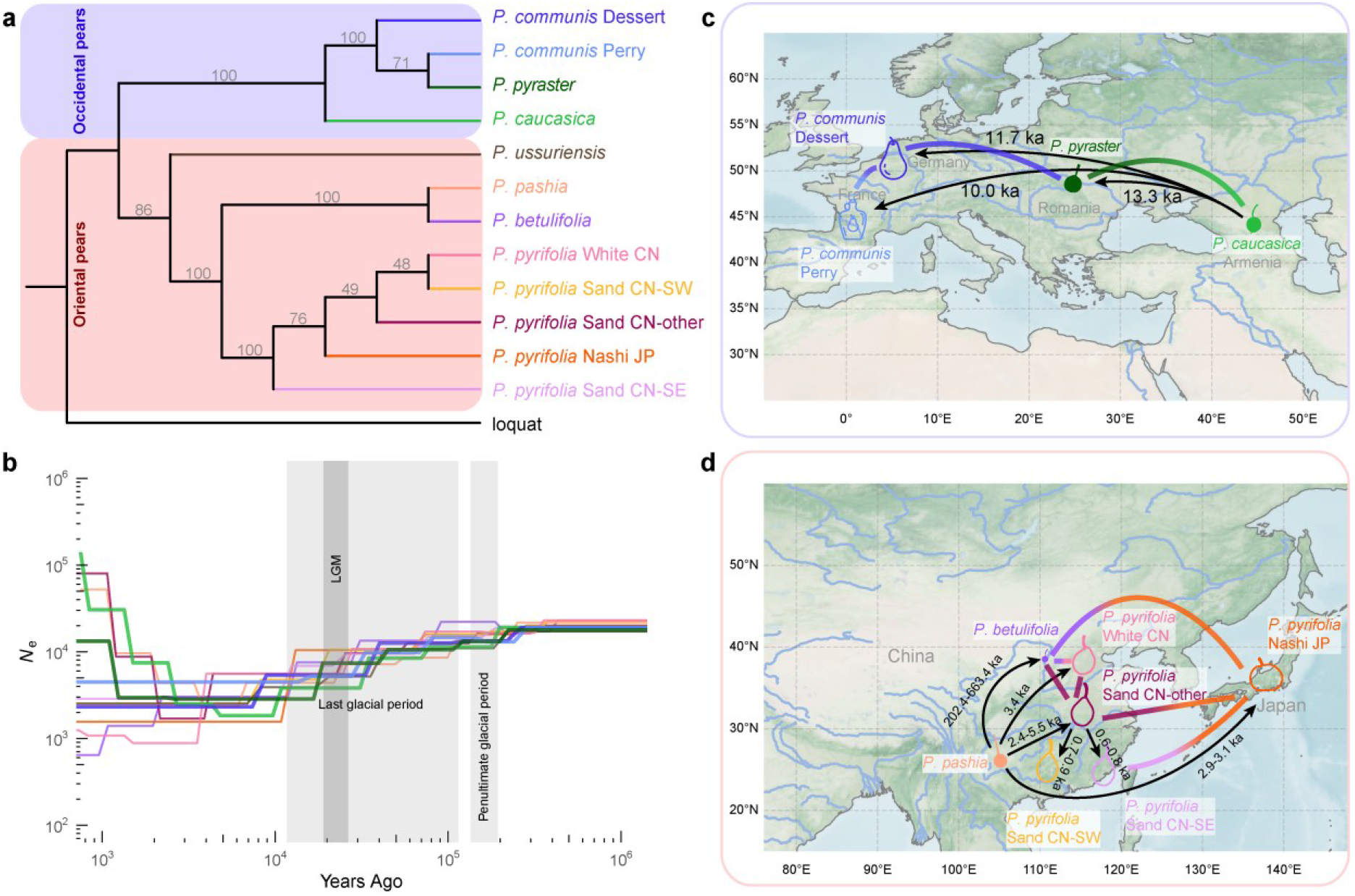
The demographic history of Occidental and Oriental pear populations. **a** Phylogenetic tree of pear populations under the coalescent model generated using SVDquartets. Standard bootstrap values are shown on the branches. **b** Historical changes in effective population size for Occidental and Oriental pear populations inferred using SMC++, assuming a generation time of 7.5 years and a mutation rate of 3.9 × 10^−8^ per site per generation. LGM, Last Glacial Maximum. **c,d** Best-supported scenarios (lowest AIC among the models tested) of domestication and divergence inferred for Occidental (c) and Oriental (d) pear populations. Arrows denote divergence events with associated time estimates; colored lines indicate gene flow, with line color representing the source population. Populations are color-coded based on their fastSTRUCTURE assignments (Supplementary Table 4). Split times were converted assuming a generation time of 7.5 years and were estimated from the best-fitting models based on AIC (Supplementary Table 7). When a split time was inferred in multiple rounds, the minimum–maximum range across the best-fitting model of each round was reported. Country names indicate approximate geographical origins only and do not imply any political position on territorial status. The basemap was obtained from NASA Visible Earth.

We quantified historical gene flow using the *f*_4_-ratio, *D*-statistics (ABBA-BABA), and the *f*-branch test (Supplementary Figs. 25,26; Supplementary Tables 5,6). Under strict isolation (no gene flow), these statistics approached zero, with divergence explained solely by drift^32,33^. Significant non-zero values identified multiple episodes of gene flow. In Occidental pears, we detected gene flow between wild and European pears (*P. pyraster* and dessert *P. communis*; *P. caucasica* and perry *P. communis*) and between wild populations (*P. caucasica* and *P. pyraster*). Notably, in Oriental pears, our analysis revealed previously underappreciated gene flow between *P. pashia* or *P. ussuriensis* and *P. pyrifolia* cultivars. However, we detected no gene flow between the two wild populations, *P. ussuriensis* (Ussurian pear) and *P. pashia* (Himalayan pear), consistent with their geographically distant and isolated distributions. Cultivar–cultivar introgression coincided with low phylogenetic bootstrap support (Fig. 2a). Elevated *f*_4_-ratios between *P. betulifolia* and white pears suggested recurrent gene flow, likely facilitated by their overlapping distributions.

We estimated unbiased nucleotide diversity (*π*) for each population and pairwise *F*_ST_ and *d*_XY_ for population comparisons using pixy with SNP and invariant sites data and estimated other demographic statistics using Stacks with SNPs (Table 1; Supplementary Figs. 27,28). In Occidental pears, cultivar populations (dessert and perry *P. communis*; *π* = 5.335 × 10^−3^ and 5.829 × 10^−3^; Wilcoxon rank-sum test, two-sided, *p* < 0.01) had significantly higher diversity than wild populations (*P. pyraster* and *P. caucasica*; *π* = 5.106 × 10^−3^ and 4.284 × 10^−3^; *p* < 0.01). In Oriental pears, *P. ussuriensis* (*π* = 4.168 × 10^−3^; *p* < 0.01) had the lowest diversity, indicating a relatively isolated population, consistent with relatively high *F*_ST_ values with other Oriental populations (Supplementary Fig. 28), and an isolated branch on the population tree (Fig. 2a). However, *f*_4_-ratio statistics revealed gene flow signals between *P. ussuriensis* and multiple Oriental populations (Supplementary Fig. 25), which is consistent with ancient introgression. *P. pashia* (*π* = 5.29 × 10^−3^, *p* < 0.01) was more diverse than white and Japanese pears (*π* = 4.905 × 10^−3^ and 4.833 × 10^−3^; *p* < 0.01; between white and Japanese pears, *p >* 0.01), but less diverse than sand pears (*π* = 5.879 × 10^−3^, 5.857 × 10^−3^ and 5.982 × 10^−3^; *p* < 0.01; within sand pears, *p* > 0.01). *P. betulifolia* (represented in our dataset by individuals commonly used as rootstocks) showed relatively high diversity (*π* = 5.575 × 10^−3^; observed heterozygosity *H*_o_ = 0.28), consistent with its wide distribution. The *F*_ST_ and *d*_XY_ results revealed clear genetic differentiation between the Occidental and Oriental pear populations (Supplementary Fig. 28). Overall, differences in *π*, *H*_o_, *H*_e_, and the negative inbreeding coefficient (*F*_IS_) suggest substantial gene flow and only moderate bottleneck effects in most domesticated pears.

**Table 1.**
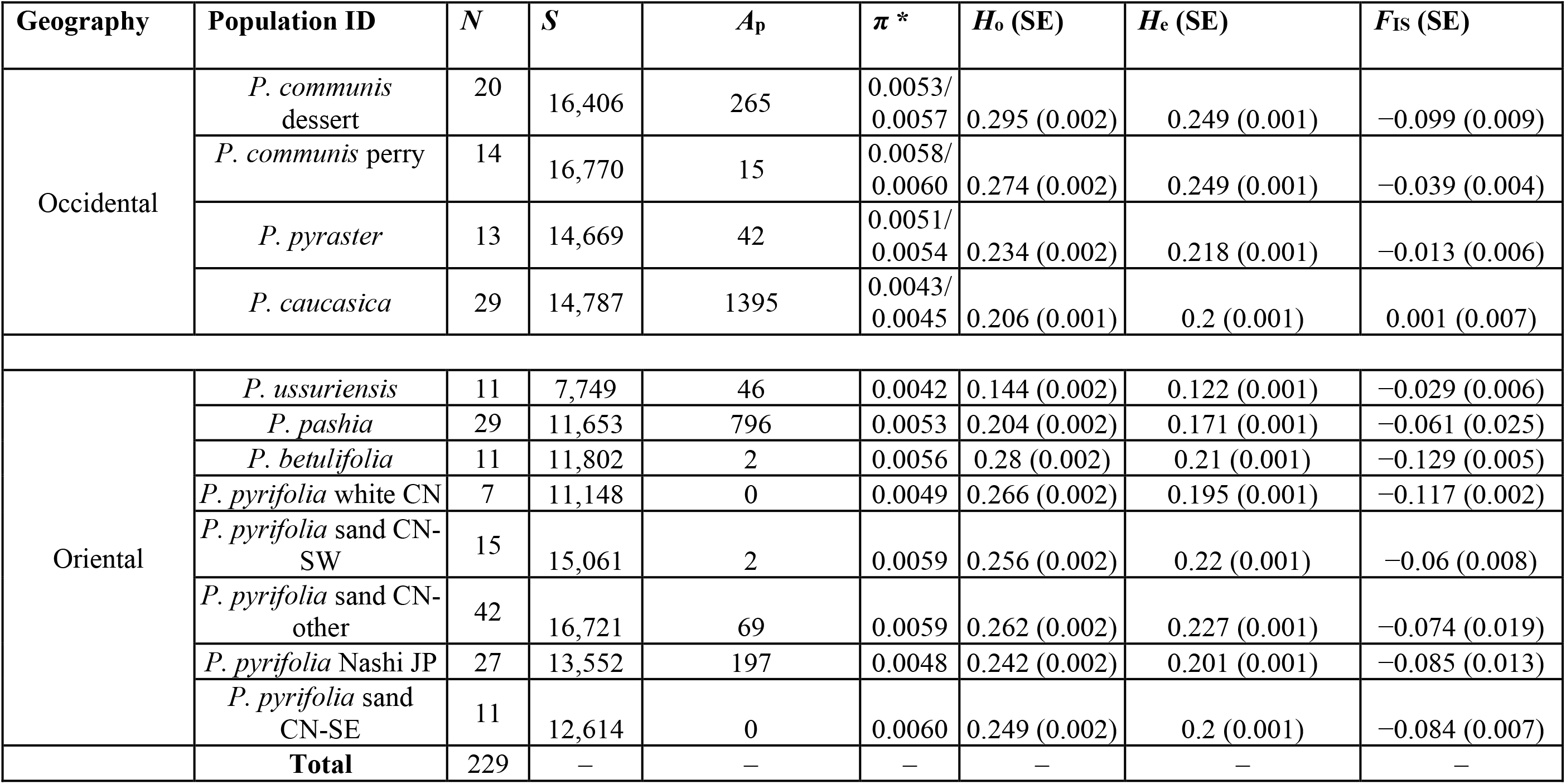
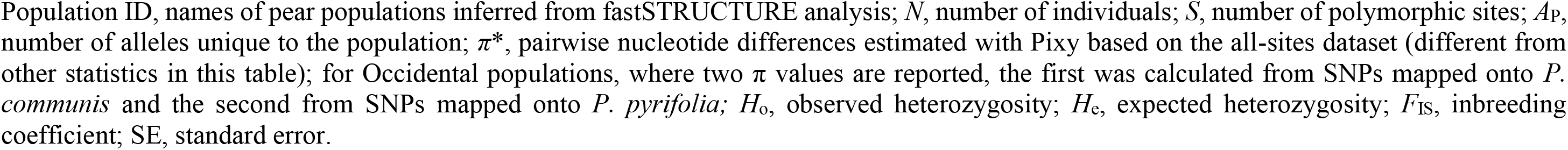
Genetic diversity estimates for each population of Occidental and Oriental pears. (*N* = 76; 20,178 Occidental and *N =* 153; 21,141 Oriental SNPs mapped onto *P. communis* and *P. pyrifolia,* respectively, detected with fastSTRUCTURE, excluding 167 admixed individuals that could not be assigned to any cluster with a membership coefficient ≥ 0.8, or non-consensus individuals whose genetic assignment did not match their taxonomic classification).

| Geography | Population ID | $N$ | $S$ | $A_p$ | $\pi^*$ | $H_o$ (SE) | $H_e$ (SE) | $F_{is}$ (SE) |
| --- | --- | --- | --- | --- | --- | --- | --- | --- |
| Occidental | *P. communis* dessert | 20 | 16,406 | 265 | 0.0053/<br>0.0057 | 0.295 (0.002) | 0.249 (0.001) | -0.099 (0.009) |
|  | *P. communis* perry | 14 | 16,770 | 15 | 0.0058/<br>0.0060 | 0.274 (0.002) | 0.249 (0.001) | -0.039 (0.004) |
|  | *P. pyraster* | 13 | 14,669 | 42 | 0.0051/<br>0.0054 | 0.234 (0.002) | 0.218 (0.001) | -0.013 (0.006) |
|  | *P. caucasica* | 29 | 14,787 | 1395 | 0.0043/<br>0.0045 | 0.206 (0.001) | 0.2 (0.001) | 0.001 (0.007) |
| Oriental | *P. ussuriensis* | 11 | 7,749 | 46 | 0.0042 | 0.144 (0.002) | 0.122 (0.001) | -0.029 (0.006) |
|  | *P. pashia* | 29 | 11,653 | 796 | 0.0053 | 0.204 (0.002) | 0.171 (0.001) | -0.061 (0.025) |
|  | *P. betulifolia* | 11 | 11,802 | 2 | 0.0056 | 0.28 (0.002) | 0.21 (0.001) | -0.129 (0.005) |
|  | *P. pyrifolia* white CN | 7 | 11,148 | 0 | 0.0049 | 0.266 (0.002) | 0.195 (0.001) | -0.117 (0.002) |
|  | *P. pyrifolia* sand CN-SW | 15 | 15,061 | 2 | 0.0059 | 0.256 (0.002) | 0.22 (0.001) | -0.06 (0.008) |
|  | *P. pyrifolia* sand CN-other | 42 | 16,721 | 69 | 0.0059 | 0.262 (0.002) | 0.227 (0.001) | -0.074 (0.019) |
|  | *P. pyrifolia* Nashi JP | 27 | 13,552 | 197 | 0.0048 | 0.242 (0.002) | 0.201 (0.001) | -0.085 (0.013) |
|  | *P. pyrifolia* sand CN-SE | 11 | 12,614 | 0 | 0.0060 | 0.249 (0.002) | 0.2 (0.001) | -0.084 (0.007) |
| **Total** |  | 229 | — | — | — | — | — | — |

Population ID, names of pear populations inferred from fastSTRUCTURE analysis; $N$ , number of individuals; $S$ , number of polymorphic sites; $A_P$ , number of alleles unique to the population; $\pi^*$ , pairwise nucleotide differences estimated with Pixy based on the all-sites dataset (different from other statistics in this table); for Occidental populations, where two $\pi$ values are reported, the first was calculated from SNPs mapped onto *P.* *communis* and the second from SNPs mapped onto *P. pyrifolia*; $H_o$ , observed heterozygosity; $H_e$ , expected heterozygosity; $F_{IS}$ , inbreeding coefficient; SE, standard error.

Demographic reconstruction revealed long-term declines predating domestication, broadly coinciding with the Last Glacial Period (115–11.7 ka)^34^, followed by stabilization or recovery (Fig. 2b). Crucially, these demographic histories explain the observed variation in linkage disequilibrium (LD) decay rates measured by the *r*² statistic. The *N*_e_ recovery observed in *P. caucasica*, sand CN-other, and *P. pashia* pear populations aligns with reduced *r*^2^ in these populations (Supplementary Fig. 29). In contrast, the increased *r*^2^ observed in white pear and *P. betulifolia* is consistent with the admixture events detected by the *f*_4_-ratio statistics (Supplementary Fig. 25). The parallel *N*_e_ trajectories in wild and cultivated populations indicate that domestication imposed at most weak bottlenecks.

We fitted demographic models with gene flow using fastsimcoal2^35^ (Table 2; Supplementary Figs. 30–37; Supplementary Tables 7,29). For Occidental pears, independent divergence of both dessert and perry *P. communis* was the best-supported model (the lowest AIC), favored over post-domestication diversification from a shared cultivated ancestor. Perry *P. communis* diverged later, consistent with its later adoption for fermentation (Fig. 2c; Supplementary Note 1; Supplementary Figs. 32–34). The population trees (SVDquartets, CASTER) instead place *P. pyraster* closest to the cultivars, as sister to perry, whereas the best-supported model derives both cultivated populations from the ancestral pool represented by *P. caucasica* (Supplementary Table 29); we therefore infer separate divergence of dessert and perry but not their wild source. The model also supported continuous bidirectional gene flow between wild *P. pyraster* and *P. caucasica*, along with introgression from cultivated dessert pears back into wild *P. pyraster* (Supplementary Table 7).

**Table 2.** Demographic model selection with fastsimcoal2. Each row is one analysis. For each analysis, combinations of divergence topology and gene-flow configuration were fitted, and the best-supported model was selected by AIC (60 models for the Occidental lineage; 109 for the Oriental lineage, run in five rounds; all models in Supplementary Table 29; parameter estimates of the best-supported models in Supplementary Table 7). ΔAIC values are comparable only within a row. ΔAIC to closest alternative divergence: AIC of the best model with a different divergence topology minus that of the best-supported model. ΔAIC to no gene flow involving cultivars: AIC of the model with the same topology and without gene flow involving cultivated populations minus that of the best-supported model. Divergence and gene flow are given forward in time: A → B, gene flow modeled from A to B only; A ⇄ B, gene flow modeled in both directions, with the higher estimated rate from A to B.

| Analysis (populations) | Alternatives compared | *N* models tested (topologies × gene-flow configurations) | Best-supported divergence (model ID) | Gene flow in the best model | ΔAIC to closest alternative divergence | ΔAIC to no gene flow involving cultivars |
| --- | --- | --- | --- | --- | --- | --- |
| Occidental lineage: *P. pyraster*, *P. caucasica*, *P. communis* dessert, and *P. communis* perry | Dessert and perry diverged separately from wild populations (3 topologies) vs perry diverged from dessert (2) | 60 ($5 \times 12$) | Dessert, then perry, diverged from *P. caucasica* (westG10D3) | *P. pyraster* ⇌ *P. caucasica*; perry ⇌ dessert; dessert → *P. pyraster* | 8,808 (perry from dessert); 43,866 (perry from dessert, with wild-to-perry gene flow) | 84,223 |
| Oriental lineage, round 1: *P. betulifolia*, *P. pashia*, *P. pyrifolia* sand CN-other, and *P. pyrifolia* sand CN-SW | Sand CN-SW diverged from sand CN-other or from the *P. pashia* population, or sand CN-other diverged from sand CN-SW | 8 ($4 \times 2$) | Sand CN-SW diverged from sand CN-other (eastS2G1) | sand CN-other ⇌ *P. betulifolia*; *P. betulifolia* ⇌ sand CN-SW | 13,043 (sand CN-other from sand CN-SW) | 140,918 |
| Oriental lineage, round B1: as round 1, with sand CN-SE replacing sand CN-SW | As round 1, for sand CN-SE | 8 ($4 \times 2$) | Sand CN-SE diverged from sand CN-other (eastS2G1) | sand CN-other ⇌ *P. betulifolia* | 77,037 (sand CN-other from sand CN-SE) | 129,200 |
| Oriental lineage, round 2: round 1 + *P. pyrifolia* Nashi JP | Nashi JP diverged from the *P. pashia* population (4 topologies) vs from sand CN-other or sand CN-SW (6) | 35 ($10 \times 4 - 5$) | Nashi JP diverged from the *P. pashia* population after sand CN-other; sand CN-SW diverged from sand CN-other (eastJ31G3) | Nashi JP ⇌ *P. betulifolia*; Nashi JP ⇌ sand CN-other | 120,506 (sand CN-other from sand CN-SW); 123,090 (Nashi JP from sand CN-SW) | 342,392 |
| Oriental lineage, round B2: round B1 + *P. pyrifolia* Nashi JP | As round 2, with sand CN-SE | 40 ($10 \times 4$) | Nashi JP diverged from the *P. pashia* population after sand CN-other; sand CN-SE diverged from sand CN-other (eastJ31G3) | Nashi JP ⇌ *P. betulifolia*; Nashi JP ⇌ sand CN-other; Nashi JP ⇌ sand CN-SE | 3,852 (sand CN-other from the sand CN-SE); 125,961 (Nashi JP from sand CN-SE) | 344,862 |
| Oriental lineage, round C1: *P. betulifolia*, *P. pashia*, *P. pyrifolia* sand CN-other, and *P. pyrifolia* white CN | White CN diverged from the *P. pashia* population (2 topologies), sand CN-other (1) or *P. betulifolia* (2), or sand CN-other diverged from white CN (1) | 18 ($6 \times 3$) | White CN diverged from the *P. pashia* population after sand CN-other (eastW5G2) | white CN ⇌ *P. betulifolia*; white CN → sand CN-other; *P. betulifolia* → sand CN-other | 2,440 (white CN from sand CN-other) | 216,424 |

For Oriental pears, we used a staged strategy testing overlapping populations subsets across five rounds, given that joint modeling of all eight populations was computationally intractable; the best-supported topologies were consistent across rounds (Fig. 2d; Supplementary Note 1; Supplementary Table 7; Supplementary Figs. 31, 35–37). We used *P. pashia* (Himalayan pear) as an ancestral proxy^17,28,36^, the most divergent extant lineage closely related to *P. pyrifolia* in our dataset. This does not imply that *P. pashia* is the direct ancestor of cultivated sand pears; rather, it provides a reference point in the absence of a clearly identified wild progenitor, and downstream inferences carry that limitation. Under this framework, sand CN-SW and sand CN-SE populations diversified regionally after domestication of sand CN-other, the basal population among the three, despite the geographic proximity of sand CN-SW to the native range of *P. pashia* (Fig. 2d; Supplementary Figs. 31,37). White pears formed a distinct domesticated population that branched alongside the sand pear populations (Fig. 2d; Supplementary Figs. 31,37). For Japanese *P. pyrifolia*, AIC-based comparisons favored divergence of Japanese pears from the wild proxy over divergence from Chinese *P. pyrifolia* cultivars; we present this as the best-supported scenario among those tested, not a settled conclusion, given its dependence on the staged models and the *P. pashia proxy* above. Our results are consistent with separate divergence of Chinese and Japanese pears from a common ancestral gene pool, rather than linear expansion from a single Chinese source. The true wild progenitor may be unsampled or extinct, and we did not model an unsampled ancestral population.

Cultivars diverged from approximately 1,554 (dessert) and 1,332 (perry) generations ago (95% CI: 1545 to 1673, and 1145 to 1476) for Occidental pears, and 324 to 737 generations ago for Oriental populations (range across modeling rounds; Supplementary Table 7). Assuming a 7.5-year generation time, these estimates correspond to approximately 10.0–11.7 ka and 2.4–5.5 ka, respectively.

### Signatures of positive selection during domestication in the pear genomes

Because population structure followed fruit use in Europe and geography in Asia. We anticipated that selective sweeps would be predominantly population-specific with minimal sharing across lineages. As expected, our analysis using two complementary statistics, including *ω* (OmegaPlus^37^) and unbiased Tajima’s *D* (pixy^38^), identified 358 to 652 candidate genes per population, the vast majority of which were indeed specific to a single population (Fig. 3; Supplementary Figs. 39–49; Supplementary Tables 8–12).

**Figure 3.**
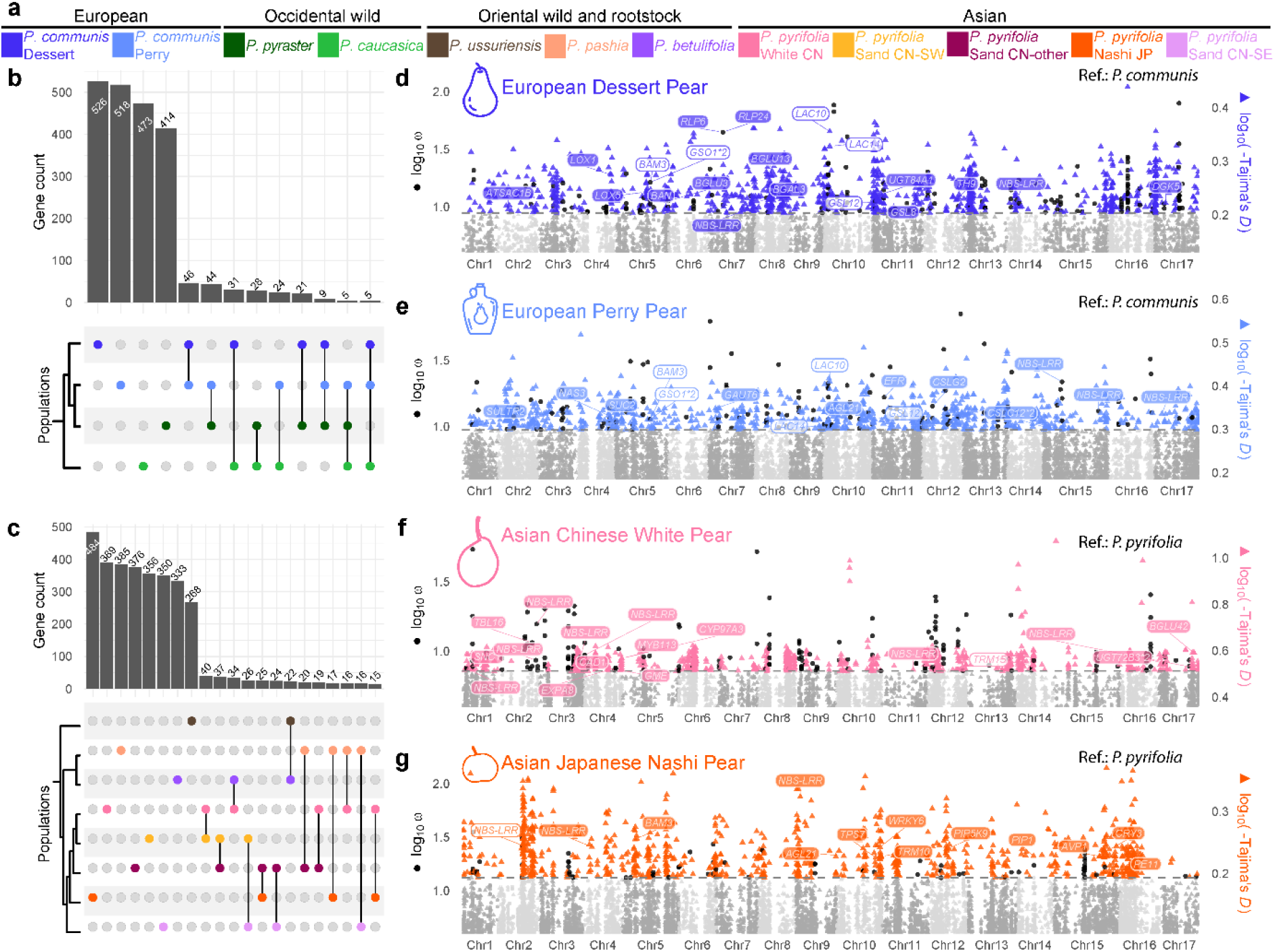
Signatures of positive selection in Occidental and Oriental pears and domestication footprints. **a** Population color key used across all panels. **b, c** Intersection counts of genes under positive selection within and among Occidental (b) and Oriental (c) pear populations; only intersections containing >3 genes are shown. **d–g** Genome-wide scans for selective sweeps in four representative populations: *P. communis* ‘dessert’ (d), *P. communis* ‘perry’ (e), *P. pyrifolia* ‘White CN’ (f), and *P. pyrifolia* ‘Nashi JP’ (g). Positive selection was inferred using Tajima’s *D* (pixy; triangles colored by population) and *ω* (OmegaPlus; black circles) statistics from SNPs mapped to the reference genome for each species. Centromeric and low-mappability regions were excluded to reduce false positives. Each point represents a statistic in a sliding window; the dashed horizontal line indicates the optimized cutoffs (99.45th percentile for *ω*, 0.40th percentile for Tajima’s *D*) determined by 1,000 permutations in regioneR. Values of log_10_(-Tajima’s *D*) were rescaled so that both cutoffs coincide. Points below cutoffs are shown in grey. Points with weak signals (*ω* < 1 or Tajima’s *D* > −0.5) are not plotted. For *P. pyrifolia* panels, chromosome numbers were reassigned to the *P. communis* reference using MCScanX and minimap2 synteny (Supplementary Note 1). Gene labels show official NCBI symbols when available; a white background indicates a gene shared with ≥1 other cultivar population, and an asterisk denotes a duplicated gene or gene family.

Among the few shared candidate genes, *P. pashia* (Himalayan pear) showed significantly greater overlap with *P. pyrifolia* cultivars than *P. betulifolia* or *P. ussuriensis* (Fig. 3c; Supplementary Fig. 47). This elevated overlap suggests that some adaptive haplotypes were already segregating in wild gene pools. Shared sweeps overlapped windows with excess allele sharing more often than expected in the Occidental lineage (2.8- to 6.8-fold, FDR ≤ 0.033), but the association weakened after adjusting for local diversity, window length, and gene density (1.3- to 4.4-fold, FDR ≥ 0.082) and was absent in the Oriental lineage (Supplementary Tables 13–15). We therefore found no robust evidence that sweeps are enriched in regions of gene flow.

In cultivated pears, selective-sweep regions encompassed genes involved in metabolism, fruit quality, use traits, defense, and adaptation to biotic/abiotic stresses (Fig. 3d–g). European and Asian pears differ mainly in postharvest ripening traits and fruit shape, suggesting distinct domestication footprints. Gene Ontology enrichment of lineage-specific genes highlighted contrasting functional profiles between lineages: Occidental pears were primarily enriched in growth, lipid metabolism, and fruit development processes (Supplementary Table 10), while Oriental pears showed a predominance of abiotic and oxidative stress–related terms (Supplementary Table 11).

Despite their different end uses, dessert and perry *P. communis* populations shared 46 candidate genes within selective-sweep regions (Fig. 3d,e). These genes include *BARELY ANY MERISTEM 3* (*BAM3*; pycom05g17790) and *GASSHO1* (*GSO1*; pycom05g17800, pycom05g17770), encoding leucine-rich repeat receptor-like kinases that may affect fruit peel structure and contribute to disease resistance^39^. Population-specific selective-sweep regions in the dessert *P. communis* contained genes implicated in flavor/texture pathways, including *BETA GLUCOSIDASE* (*BGLU3*; pycom05g32120), *BGLU13* (pycom08g12030), and *LIPOXYGENASE 1* (*LOX1*; pycom04g18070); Selective-sweep regions in the perry *P. communis* population comprised genes involved in cell-wall biosynthesis and modification, including the pectin biosynthetic gene *GALACTURONOSYLTRANSFERASE 6* (*GAUT6*; pycom07g00090), the cellulose synthase genes *CSLC12* (pycom15g06110, pycom15g06120) (Fig. 3e).

For Asian pears, we identified genes under selection associated with fruit quality and adaptation to Japan’s abiotic environment (Fig. 3f,g). White pear sweep regions contained genes consistent with their crisp texture, including two *EXPANSIN 8* homologs (*EXPA8*; GWHGBAOS027806, GWHGBAOS026111). These regions also contained *CYTOCHROME P450 97A3* (*CYP97A3*; GWHGBAOS028374), involved in lutein biosynthesis, and a *MYB* transcription factor gene (*Arabidopsis* homolog *MYB113*; GWHGBAOS028351) (Fig. 3f). In Japanese pear, selective-sweep regions included genes involved in plant adaptation to humid, mild monsoon climates while maintaining fruit quality, such as genes related to hydraulics and osmotic regulation (*PLASMA MEMBRANE INTRINSIC PROTEIN 1* [*PIP1*], GWHGBAOS012726; *PHOSPHATIDYL INOSITOL MONOPHOSPHATE 5 KINASE 9* [*PIP5K9*], GWHGBAOS008238) (Fig. 3g).

### Lower realized but higher potential deleterious burden in cultivated pears

To assess whether domestication altered the burden of putatively deleterious variants, we annotated nonsynonymous SNPs in both lineage-matched references (*P. communis* and *P. pyrifolia*) using the SIFT 4G algorithm^40^.

Cultivated populations carried fewer deleterious alleles per synonymous site than their wild relatives. Within the Oriental lineage, wild populations exhibited the highest deleterious burden, whereas *P. pyrifolia* Japanese cultivars showed lower levels of deleterious variants than Chinese cultivars (Fig. 4a; Supplementary Table 16). Cultivated pears carried fewer homozygous (realized) but more heterozygous (potential) deleterious alleles than wild pears in both lineages (Supplementary Fig. 52). A linear mixed model accounting for population and sequencing depth gave effects of similar size in both lineages, significant in Oriental pears but not in Occidental pears, which comprise only four populations (Supplementary Table 18). Loss-of-function (LoF) variants, annotated with SnpEff and VEP independently of SIFT 4G, showed the same pattern (Supplementary Fig. 54; Supplementary Tables 19,20), although the total LoF burden of White pear and sand CN-SW did not differ significantly from that of *P. ussuriensis*. When we compared selective sweeps with matched controls, selective-sweep regions carried a higher deleterious burden than controls (two-sided Wilcoxon tests; Fig. 4b; Supplementary Table 16), consistent with linked selection (genetic hitchhiking), whereby positively selected haplotypes carry linked deleterious variants to higher frequency and locally reduce the efficacy of purifying selection. Across populations, the potential (heterozygous) load exceeded the realized (homozygous) load (Fig. 4c; Supplementary Table 16). This local excess in sweep regions was notably evident for realized deleterious variants (Fig. 4d; Supplementary Table 16; heterozygous burden in Supplementary Fig. 53). Together, these results indicate that, at swept loci, positive selection locally elevated the relative deleterious load.

**Figure 4.**
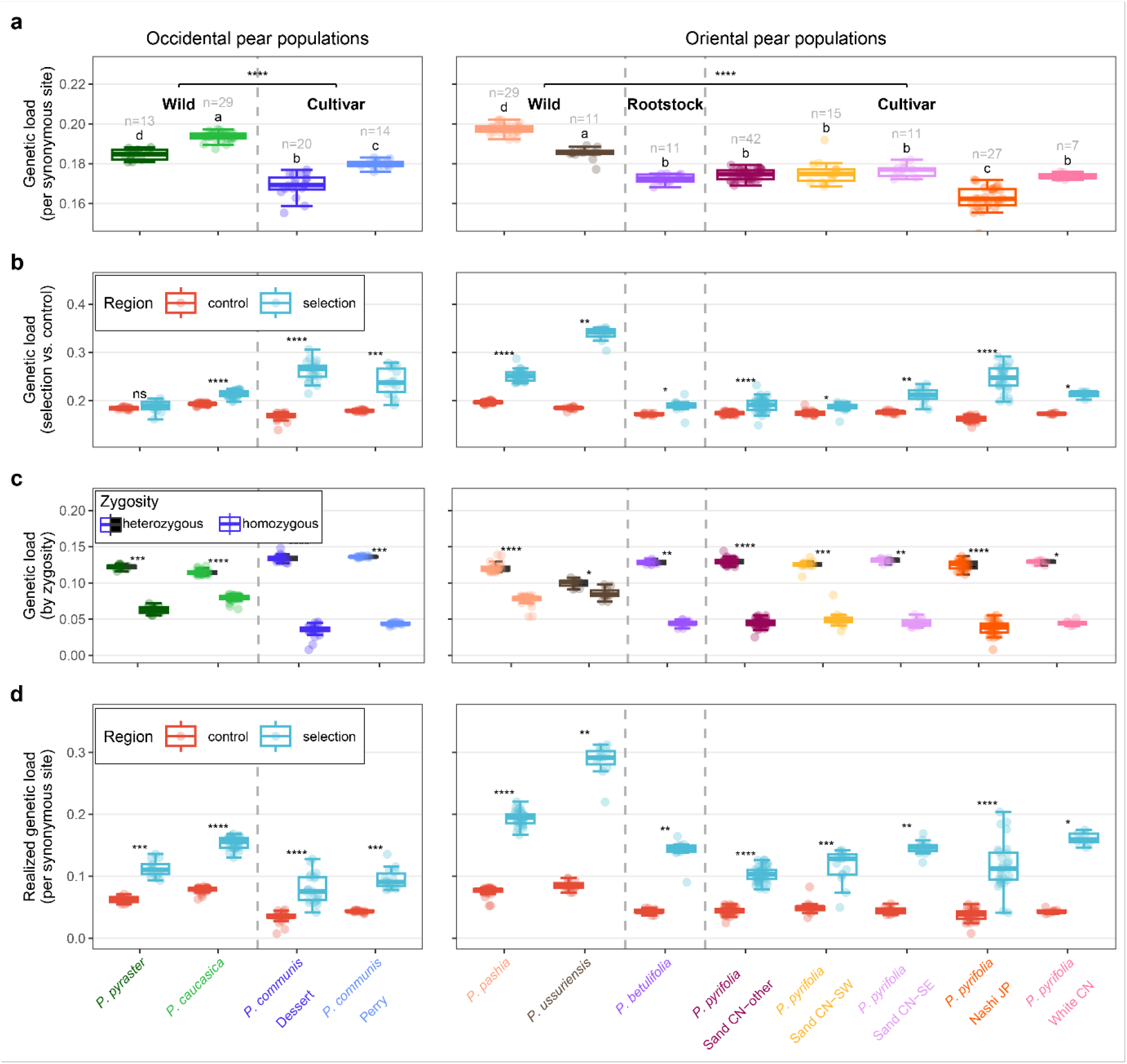
The genetic burden during pear domestication. **a** Number of deleterious mutations (SIFT score < 0.05; SNPs or fixed sites) per carried synonymous site, using *P. communis* and *P. pyrifolia* as reference genomes for the Occidental and Oriental pear populations, respectively. Stars above boxplots indicate significant differences between cultivar and wild populations (two-sided Wilcoxon rank-sum test of pooled cultivated versus pooled wild individuals, performed separately within each lineage; P. betulifolia rootstocks are excluded from the wild group; one pre-planned comparison per lineage, so no multiplicity correction). Comparisons were between individuals and did not account for population structure. A population-level analysis, in which wild and cultivated groups were compared with population as a random effect and sequencing depth as a covariate, is given in Supplementary Table 18. Different letters above boxplots indicate significant differences (Wilcoxon test, Holm-adjusted, *p* < 0.01) in the number of deleterious alleles among populations. Comparisons were performed separately for the Occidental and Oriental populations. **b** Comparison between selection sweeps and control regions (the rest of the genome after excluding low-mappability and centromeric regions). **c** Comparison of deleterious mutations between homozygous and heterozygous (half-black boxes represent heterozygous mutations) states. **d** Comparison of homozygous deleterious mutations between selection sweep regions and control regions. Boxplot elements: center line, median; box limits, 25th and 75th percentiles; whiskers, extending to the most extreme data point within 1.5 × the interquartile range from the box; points, individual data. Source data are provided in Supplementary Table 16. *P* values are provided in Supplementary Table 17. Significance levels (two-sided paired Wilcoxon test, Holm-adjusted within each lineage): ns, *p* > 0.05 (not significant); * *p* ≤ 0.05; *\*\* p ≤* 0.01; *\*\*\* p ≤* 0.001; *\*\*\*\* p ≤* 0.0001.

Cultivated populations showed higher *F*_ROH_ (the fraction of the genome in runs of homozygosity) than wild populations (two-sided Wilcoxon rank-sum test), consistent with recent inbreeding (Supplementary Fig. 55). Grouping individuals by lineage revealed that the realized genetic load correlated negatively with *F*_ROH_, whereas the correlation was positive within populations (11 of 12 populations, significant in four; Supplementary Fig. 56). Within individuals, the realized burden was higher inside than outside ROH (Supplementary Fig. 57). Recent inbreeding therefore increased the realized burden within cultivated populations, but it does not explain their lower burden relative to wild populations.

Thus, cultivated pears do not show the higher deleterious burden predicted by the cost-of-domestication hypothesis^41^.

### TE variation tracks demographic structure

We detected 13,574 reference and 169,796 non-reference TIPs in the four Occidental populations, and 12,064 and 203,296, respectively, in the eight Oriental populations (TIP analyses are based on the 67 Occidental and 82 Oriental non-admixed individuals with TE calls; Methods; Supplementary Figs. 58,59). Neutral unlinked and synonymous SNPs revealed population differentiation in Oriental pears (PC1 = 46.4%) and in Occidental pears (18.1%), whereas TE insertions consistently explained∼10% of the variation across all populations. Thus, we find no evidence that heterogeneous TE activity obscures genome-wide demographic patterns. Wild populations (*P. caucasica* in the Occidental lineage; *P. ussuriensis* in the Oriental lineage) formed well-defined clusters with both SNPs and TIPs. In Occidental pears, admixed individuals clustered closer to *P. pyraster* in the TE-based PCA, suggesting relatively recent introgression between wild and cultivated gene pools. In contrast, Oriental admixed individuals grouped consistently with cultivated populations across markers (Figs. 5a,b).

**Figure 5.**
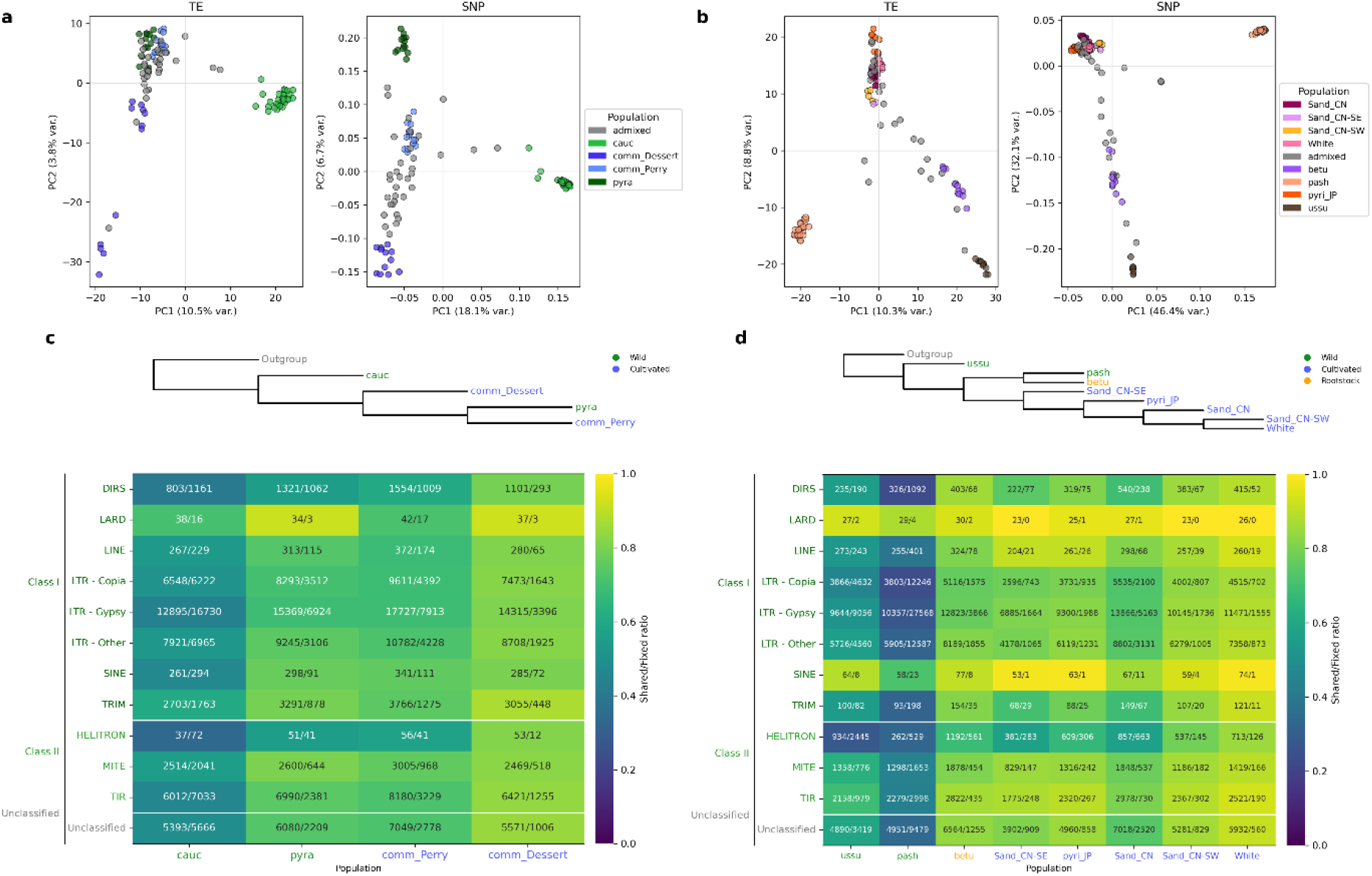
Shared and fixed transposable element insertions in Occidental and Oriental pear populations. Throughout the figure, the left panels (a, c) correspond to Oriental populations, while the right panels (b, d) correspond to Occidental populations. **a, b** PCA plot based on TE insertions (left) and SNP variation computed with PLINK2 (right). **c, d** Heatmaps showing the ratio of shared versus fixed insertions for each TE order, grouped into Class I (retrotransposons) and Class II (DNA transposons). Each cell indicates the number of shared and fixed insertions within a population, with color gradients reflecting the ratio of shared to fixed insertions (0–1). Above each heatmap, phylogenetic trees depict relationships among populations from SVDquartets. Population colors represent wild (green), cultivated (blue), and rootstock (orange) individuals.

We then tested whether TIPs were associated with genes under positive selection, comparing selected and non-selected genes within each population with cluster-stratified Cochran–Mantel–Haenszel (CMH) tests, covariate-adjusted logistic models and 1,000 matched control gene sets (Supplementary Note 4). No population showed TIP enrichment near selected genes (Supplementary Table 25). Four populations showed a depletion in both the adjusted models and the matched controls, upstream of selected genes in *P. caucasica* and downstream in *P. pashia*, *P. ussuriensis,* and White pear (adjusted odds ratio = 0.60–0.70, FDR ≤ 0.026; matched-control FDR = 0.032), but none did in the CMH test (FDR ≥ 0.23). Three further depletions were significant only in the adjusted models and not supported by the matched controls (FDR ≥ 0.07): upstream in sand CN-other, in introns in Japanese pear, and in the 3′ UTR of sand CN-other, the last being a degenerate estimate resulting from zero insertions among selected genes. An insertion within *PE11* (pectinesterase 11), a gene under selection in Japanese pear, was carried by 18 cultivated *P. pyrifolia* individuals and was not detected in wild or rootstock individuals (Supplementary Table 22). In selection regions, Gypsy insertions outnumbered Copia insertions per individual in every population (paired Wilcoxon test, Holm-adjusted *P* < 0.05 in 8 of 12; Supplementary Fig. 61), whereas Copia consensus sequences had more of their insertions within 2 kb of genes. No single consensus sequence was significantly enriched or depleted in selection regions (Supplementary Fig. 62).

## Discussion

We reconstructed the mode and tempo of pear domestication by integrating SNP- and TIP-based analyses across 396 individuals and lineage-matched reference genomes. Our study refines the domestication narrative of this woody perennial crop, revealing independent divergence of cultivated populations within each lineage, sustained gene flow with wild relatives, and candidate selection signals in genes annotated to immunity, cell wall, carbohydrate metabolism, and ripening pathways.

### A protracted, multi-regional domestication shaped by recurrent gene flow

The dataset of 130 Occidental (42 wild) and 266 Oriental (40 wild and 11 rootstock) unique individuals provides high-resolution insight into domestication processes in Europe and East Asia. Coalescent-based modeling indicates that cultivated Occidental and Oriental pears diverged independently from their wild gene pools, consistent with long-standing hypotheses^17,26^, but further reveals that pear domestication occurred under weak demographic bottlenecks and sustained wild–crop gene flow (Figure 6). *N*_e_ estimates show no evidence of severe reductions, consistent with the view that perennial domestication avoids the strong genetic erosion typical of annual crops, likely owing to long generation times, and continued introgression^4,5,42^. The inferred divergence between cultivated and wild gene pools aligns broadly with archaeological and literature evidence^18,43,44^. However, these dates likely reflect initial population separation rather than the precise beginning of domestication.

**Figure 6.**
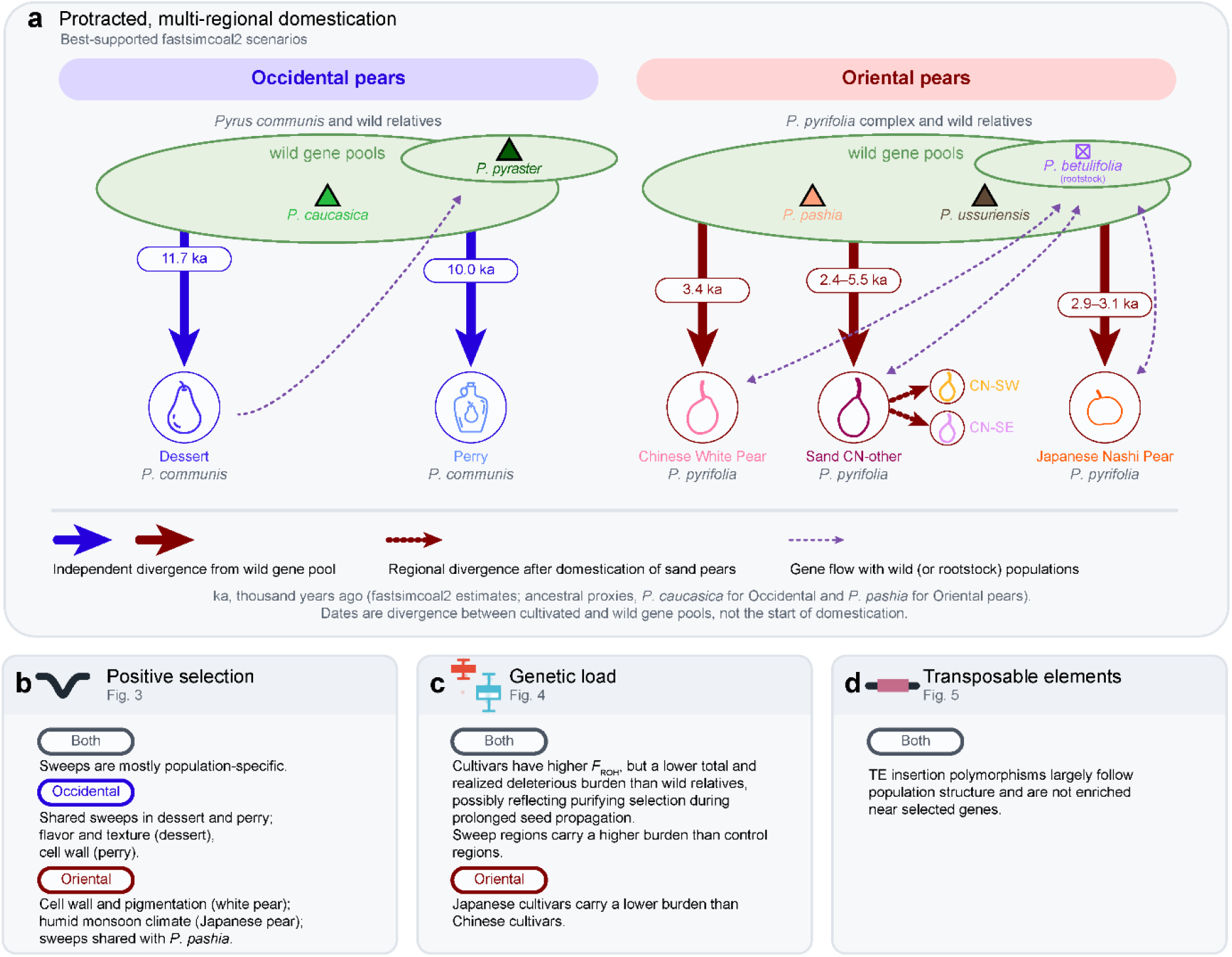
Schematic graph of Occidental and Oriental pear domestication histories. Ancestral gene pools are proxies; in the best-supported model both cultivated populations diverge from the *P. caucasica* proxy.

For Japanese pears, genomic signals consistent with a *P. pashia*-related wild component are detectable, despite *P. pashia* currently being absent from Japan. This pattern is more parsimoniously explained by dispersal of a mainland wild gene pool, followed by separate divergence of cultivated pears in Japan, than by direct domestication from extant *P. pashia*. Originating in southwestern China, *Pyrus* likely dispersed to Japan through natural range shifts or early human-mediated movement before or during the early stages of cultivation^17,45,46^. Subsequent geographic isolation of the Japanese archipelago likely facilitated the differentiation of Japanese pears into a distinct cultivated lineage.

Our analyses also help clarify several taxonomic inconsistencies, framed here as germplasm-classification insights rather than formal taxonomic conclusions. Our analyses show that the “*P. bretschneideri*” reported by Li et al.^47^ has high genetic admixture with white pear cultivars, suggesting a feral rather than wild origin. Similarly, “*P. bretschneideri*” individuals described by Wu et al.^26^ show clonality with our *P. pyrifolia* samples. Together, these results are consistent with the Chinese white pear being a differentiated subgroup within *P. pyrifolia*^19,48,49^. Because our study includes no type material or morphological data, we make no claim about the delimitation of *P. bretschneideri*.

Overall, pear domestication followed a gradual, spatially diffuse trajectory characterized by parallel selection and recurrent gene flow rather than an abrupt or centrally driven origin^50^.

### Contrasting selection histories shaped by divergent use and environmental factors

In European pears, domestication was primarily driven by use, paralleling patterns reported in apples^51^. Dessert and perry *P. communis* underwent distinct selective pressures: dessert pears showed selection on *BGAL3*, *BGLU*, and *LOX*, linked to fruit softening and aroma^52–54^, whereas perry pears displayed selection in *GAUT6* and *CSLC*, linked to cell-wall biosynthesis and firmness^55,56^. Selection for *BGAL3* in dessert pears is consistent with their characteristic melting texture^53^. In Asian pears, domestication was more geographically structured. Japanese *P. pyrifolia* shows selective sweeps in *PIP1*, *AVP1*, and *CRY3,* consistent with adaptation to a humid monsoon climate^57–60^, whereas Chinese white pears show selection on a cell-wall-loosening gene, *EXPA8*, and a carotenoid biosynthetic gene, *CYP97A3*, together with *MYB113*^61–64^. Sweep regions in Asian *P. pyrifolia* contained *TRM* homologs, key determinants of fruit shape, aligning with the selection for round, apple-like fruit^65^.

Across both lineages, selection converged on immunity and fruit quality. Sweep intervals harbor multiple *NBS-LRR* and receptor-like kinase genes, consistent with recurrent selection for disease resistance under orchard conditions^66,67^. In Asian pears, overlap between selective-sweep regions in cultivated *P. pyrifolia* and *P. pashia* suggests some adaptive haplotypes were already segregating in ancestral wild gene pools, potentially reflecting retained standing variation predating domestication or historical introgression between wild and cultivated populations. Shared sweeps were not robustly enriched for local excess allele sharing (Supplementary Table 15); distinguishing standing variation from introgression requires denser sampling of wild populations. However, this remains particularly challenging in pears (and in general), as putative wild progenitors are rare, taxonomically ambiguous, or potentially extinct, complicating efforts to distinguish shared ancestry from secondary gene flow. Although we did not explicitly test locus-specific introgression, the observed overlap is consistent with pre-existing variants from *P. pashia*–related gene pools contributing to Asian pear domestication, without implying direct ancestry or human-driven selection in wild populations.

Collectively, European domestication prioritized fruit utility and texture, whereas Asian domestication integrated fruit quality with local ecological adaptation. Because callable fractions differ between assemblies (59.4% versus 76.7%), absolute numbers of candidate regions are not comparable across lineages. Further functional validation of the genes found in regions under selection is needed to demonstrate a causal role in the corresponding pear phenotype, for example through expression analyses, association mapping in segregating populations, or genome editing.

### Genetic load: no support for a cost of domestication in pears

Our genetic load analysis illustrates how perennial domestication can diverge from annual models^68^. Cultivated pears harbored fewer deleterious variants than their wild relatives, for both SIFT 4G-predicted and LoF variants, arguing against extending the cost-of-domestication hypothesis^41^ in this system rather than refuting it generally. The total burden differed only modestly, whereas cultivars carried markedly fewer homozygous (realized) and more heterozygous (potential) deleterious alleles. Heterozygosity maintained by outcrossing, admixture, and clonal propagation can shelter recessive variants, as proposed for grapevine^69^. The lower total burden, which contrasts with grapevine, may instead reflect purifying selection during the extended period of seed-based propagation in pears before grafting became standard practice, as in apple (“chance seedling” methods)^4,9^: pears root poorly from cuttings and therefore relied longer on sexual reproduction (obligate outcrossing)^70^. Our data do not demonstrate this explanation: both reference genomes derive from cultivars, which may reduce the number of deleterious variants called in cultivated individuals, and the sampled wild populations may also not represent the direct progenitors of cultivated pears. Cultivated pears showed higher genome-wide *F*_ROH_ than wild relatives, reflecting recent inbreeding, but realized deleterious load did not offset their lower genome-wide burden. Sweep regions likewise carried a higher deleterious burden than matched controls, most pronounced for realized load, consistent with genetic hitchhiking. However, both effects remained locally restricted and did not offset the lower genome-wide burden of cultivated pears. These findings echo patterns in apples^8^. Whether other species with similarly extended sexual reproduction due to rooting difficulties, such as plums, show the same pattern remains to be tested.

### TE variation largely follows demography

TE variation in *Pyrus* largely mirrors genome-wide demographic structure rather than lineage-specific bursts of activity. TE-based clustering recapitulates SNP-defined population structure, consistent with drift and gene flow being the primary forces shaping TE landscapes, although we did not test other forces directly.

TIPs were not enriched near genes under positive selection, so insertions within selected genes, such as that in *PE11*, remain candidates for regulatory variation. Fewer TIPs than expected were found near selected genes in four populations, which we do not interpret further. Because selection status is binary, non-selected genes include undetected targets of selection; such misclassification biases associations toward the null, so our tests do not exclude weak effects. Testing them requires expression data and functional assays, as performed for an LTR retrotransposon insertion upstream of *MdMYB1* in apple^71^. Future pan-genome frameworks^72^ will be essential to minimize calling bias, refine the genomic landscape of TE dynamics, and clarify the balance between deleterious accumulation and adaptive cooption.

### Outcomes and perspectives

Modern pears represent a genomic mosaic shaped by parallel, asynchronous domestication and gene flow between wild and cultivated populations. The asymmetric pace of this process, with a geographically structured origin in Oriental pears and a use-driven trajectory in Occidental lineages, underscores that perennial domestication is a gradual and regionally diffuse process driven by both ecological and cultural selection. Our findings position *Pyrus* as a compelling model for studying adaptive evolution in perennials, where long lifespans, self-incompatibility, and recurrent introgression generate unique genomic outcomes^1,73,74^. Recent and emerging pangenome-scale resources^25,75^ will further clarify the impact of structural variants and introgression on this architecture. Ultimately, these results illustrate how life history, demography, and selection collectively govern the evolution of long-lived fruit trees.

## Methods

### DNA extraction and short-read sequencing of new samples

Genomic DNA was extracted from the leaf tissue of 235 new *Pyrus* samples from 10 (sub-) species, representing wild and cultivated pears (Supplementary Table 1), using commercial kits followed by sequencing (Supplementary Note 1). In addition, raw short-read data for 439 *Pyrus* individuals were downloaded from the ENA (European Nucleotide Archive) database (Project Accession numbers: PRJNA381668, PRJNA724820, PRJNA563813, and PRJDB5584). In total, data from 674 pear individuals sequenced as short paired-end reads were used for SNP calling.

### Quality control of whole-genome sequencing data

The bioinformatic analysis workflow is shown in Supplementary Fig. 1. To remove insufficient sequences or reads during downstream analysis from all 674 raw paired-end sequencing datasets, fastp version 0.21.0^76^ was employed. Based on the fastp default filtering level, our additional sequence-trimming targets included several bases at the start of sequences, depending on the GC ratio and specific FastQ files. The quality of clean reads was validated using FastQC v0.11.8^77^ and MultiQC v1.18^78^.

### Dual-reference mapping and variant calling

To account for the pronounced genomic divergence between Occidental and Oriental lineages, we employed a dual-reference mapping strategy using lineage-matched assemblies. Clean reads from all 674 individuals were aligned independently to both the *Pyrus pyrifolia* Cuiguan v1.0 and *P. communis* Bartlett DH v2 reference genomes^79,80^ using bwa-mem2 v2.2^81^ with default parameters. The resulting alignments were sorted with SAMtools v1.1^82^, and duplicates were marked with GATK v4.1.9.0^83^. All preprocessing steps were automated in a custom wrapper script. Variant calling was performed using the GATK HaplotypeCaller to generate both SNP data and invariant (monomorphic) sites, facilitating accurate calculations of *π* and *d_XY_*. For downstream analyses, we used 396 non-clonal individuals mapped to their lineage-matched references, unless otherwise stated. This approach ensured optimal sequencing depth, genome coverage, and heterozygosity (*H*ₒ) estimates, minimizing mapping bias across the two lineages (Supplementary Figs. 9, 10; Supplementary Note 1). To confirm that this choice did not bias cross-lineage inference, we additionally cross-mapped each lineage onto the reciprocal reference and re-estimated population structure from the same ∼20,000 synonymous, unlinked SNPs; assignments were highly concordant between references, with only a single Occidental individual differing. The Cuiguan assembly served as the reference for analyses requiring a common coordinate system across lineages, yielding 29,826,531 SNPs before filtering.

### Horizontal SNP filtering: removing low-quality markers

The variant-filtering strategy operated along two complementary dimensions: horizontal (filtering specific sites) and vertical (excluding individuals that failed quality thresholds). Horizontal filters were applied to the raw variant files using GATK (version 4.1.9.0) VariantFiltration, GATK SelectVariants tools, and bcftools (version 1.14) filter command^82^ as described previously^8^. Variants were excluded if they met any of the following criteria: (i) SNPs with a ReadPosRankSum score <-8.0 or InDels/mixed variants with a score <-20.0 were removed, as extreme values indicate a strong positional bias in the supporting reads. (ii) SNPs with a Fisher Strand (FS) value > 60.0 or Strand Odds Ratio (SOR) > 3.0, and InDels/mixed variants with FS > 200.0 or SOR > 10.0, were excluded, as these indicate potential sequencing or alignment artifacts. (iii) SNPs with a Mapping Quality (MQ) score < 40.0 or an MQRankSum < −12.5 were removed, as they reflect poor or biased read mapping. iv) SNPs, InDels, or mixed variants with an overall quality (QUAL) score < 30.0 were excluded. Only biallelic SNPs were retained, assuming the pear individuals are diploid, and additional genotype-level filters were applied to ensure reliable calls. Specifically, for each genotype, sequencing depth (DP) had to be between 3- and 100-fold, genotype quality (GQ) had to be ≥ 20, and SNPs within 10 bp of any InDel were excluded. Equivalent filtering for invariant sites replaced GQ with the reference genotype quality score (RGQ ≥ 20).

### Vertical SNP filtering: removing individuals

For vertical filtering, potential issues stemming from clones or closely related samples in our dataset were mitigated. Colonies, mutants, and synonyms are common phenomena in *Pyrus* germplasm and collections from herbaria, botanical gardens, and gene banks. However, previous studies have not adequately addressed these issues^26,44^. Therefore, the pairwise KING kinship coefficients were computed, using Plink2 v2.0_alpha2^84^ for all sample pairs, and a pruning list was generated for samples based on a coefficient cutoff of 0.354. This threshold represents the geometric mean of 0.5 (corresponding to clones) and 0.25 (corresponding to first-degree relatives). A custom script was used to improve this pruning list, prioritizing the removal of samples with a higher rate of missing variants between each pair of closely related samples. The objective was to ensure that all KING-robust coefficients for each sample pair in our dataset remained below the cutoff. Samples with a missing rate >40% (indicating lower sequencing coverage) were also excluded (Supplementary Figs. 1,2; Supplementary Table 1). Moreover, individuals with ambiguous passport information were excluded from the dataset. After applying the vertical filtering procedures, an SNP dataset of 17,769,455 variants was obtained from 396 non-clonal individuals, all with missing rates < 20%.

### Further filtering of SNPs: lowering LD and selection effects

The full SNP dataset was further filtered to obtain a non-selected, unlinked set for population structure and demographic analyses. Variants under potential selection, including nonsynonymous and linked sites, were excluded. After applying a 5% minor allele frequency filter, removing nonsynonymous variants, and thinning to one SNP per 8 kb using PLINK v1.9^85^, 21,756 unlinked SNPs from 396 individuals mapped to the *P. pyrifolia* reference genome were retained (Supplementary Table 26). Variant annotation was performed with SnpEff v5.1^86^.

### Inference of population genetic structure with passport information

The (sub-)species names, regions, and type information (wild, cultivar, or rootstock) of pear samples were obtained from the samplers or from the original data sources. Wild (and rootstock) pears were classified as Occidental or Oriental based on their native distributions, with the Tianshan-Hindu Kush Mountains serving as the biogeographical boundary. Wild populations from Central Asia and regions further west, including Asia Minor, Europe, and the Mediterranean, were designated as Occidental, whereas those confined to East Asia were designated as Oriental. A more conservative approach was used for cultivated pears, assigning them to Occidental or Oriental lineages primarily based on genetic affinities (derived from gene pool structure, phylogenetic, and PCA) rather than current cultivation regions, which may reflect human-mediated dispersal (Supplementary Table 1).

Population structure and admixture were inferred using the variational Bayesian framework, fastSTRUCTURE^87^. The number of clusters (*K*) ranged from 2 to 16, with 30 replicates per *K*. Results were consolidated using CLUMPAK^88^ and visualized using PopHelper^89^ (Fig. 1a,b; Supplementary Figs. 4, 11, 16; Supplementary Tables 2,3). The optimal *K* was determined by inspecting cross-validation errors around the plateau (“elbow”) and by visually assessing cluster consistency across *K* values (Supplementary Figs. 4–6, 11–13, 16–18; Supplementary Note 1) rather than relying solely on Δ*K*^90^.

1–IBS distance matrices were computed with PLINK v1.9, and Neighbor-Net trees were visualized using SplitsTree4 v4.18.3^91^ (Supplementary Figs. 7, 14 and 23). Principal component analysis (PCA) was performed in PLINK, and the results were plotted using ggplot2^92^ (Fig. 1c–e; Supplementary Figs. 8, 15). Consistent color schemes were used across analyses, with admixed individuals (membership < 0.8) shown in gray. Color palettes were carefully refined for accessibility using Adobe Color (Supplementary Fig. 3).

Using fastSTRUCTURE, 14 genetic clusters were identified across Occidental and Oriental pears (Fig. 1a,b). Clusters with < 7 conspecific individuals were excluded, and discordant outliers between fastSTRUCTURE and Neighbor-Net were removed (Fig. 1a,b; Supplementary Fig. 23), yielding 12 well-supported populations. Occidental populations were cross-validated using SNPs mapped to both the *P. communis* and *P. pyrifolia* references (Supplementary Figs. 11–18; Supplementary Tables 2–4). After excluding hybrids and poorly documented individuals, 229 individuals with ≥ 80% ancestry assignment were retained for downstream population-level analyses (Table 1; Supplementary Table 4). This threshold retains individuals with up to 20% secondary ancestry and lies in a low-density shoulder of the distribution of maximum membership coefficients, where few individuals fall (Supplementary Figs. 6,13,18 and 21). With thresholds of 0.7 and 0.9, the wild–cultivated difference in deleterious burden kept its direction, and re-estimated demographic models (Occidental lineage and Oriental Round 1) gave unchanged gene-flow directions, whereas individual candidate genes were more sensitive (Supplementary Figs. 63–65; Supplementary Tables 4,27).

### Estimating genomic diversity and differentiation

Low-mappability and centromeric regions were masked prior to demographic inference. Mappability profiles for both reference genomes were generated with GenMap^93^ using 140-bp k-mers (*k* = 140, *e* = 0). Mean mappability was calculated in 100-kb windows (50-kb step) with a custom script; windows with mean > 0.9 were retained as high-mappability. Centromeric regions inferred with CentIER^94^ were then subtracted. The resulting callable set comprised 302.3 Mb (59.4%) of the 17 *P. pyrifolia* chromosomes and 342.3 Mb (76.7%) of the 17 *P. communis* chromosomes; unplaced scaffolds were excluded from all analyses (Supplementary Table 28).

Pairwise nucleotide diversity (π)^95^, genetic distance (*d*_XY_)^95,96^, and differentiation (*F*_ST_)^97^ were estimated using pixy v1.2.7.beta1^98^ on “all-sites” VCFs restricted to high-mappability windows (Supplementary Figs. 27,28). Normality was tested using Anderson–Darling and Shapiro–Wilk tests (*p* < 0.05), and π differences among populations were tested using the Wilcoxon test (*p* < 0.01). Genome-wide values were computed from total pairwise differences rather than window averages. Observed (*H*ₒ) and expected (*H*ₑ) heterozygosity^99^, along with the inbreeding coefficient (*F*_IS_), were calculated using Stacks Populations v2.65^100^.

### Phylogenetic species tree construction and historical demography inference

The population-wide phylogenetic tree was reconstructed using SVDquartets implemented in PAUP version 4.0a^101,102^ with standard bootstrap tests (Fig. 2a). Additional population-tree analyses used CASTER^103^ (site- and pair-based approaches) under two alternative rooting schemes (Supplementary Fig. 24). The historical *N*_e_ was analyzed using SMC++^104^, assuming a mutation rate of 3.9 × 10^−8^ mutations per site per generation and 7.5 years per generation, and multiple EM iterations were performed 50 times (Fig. 2b). *D*-statistics (Patterson’s *D*, ABBA-BABA) and the *f_4_*-ratio test via DtriosParallel of Dsuite^105^ were used to estimate the genome-wide introgression between pear populations. Local admixture statistics (*f*_d_ and *f*_dM_) were computed with Dinvestigate (Dsuite) in sliding windows of 50 SNPs advanced in 25-SNP steps.

### Demographic modeling based on site frequency spectrum (SFS)

To investigate the demographic history of pear populations, fastsimcoal2 v2.8^35^ was employed. Informed by the phylogenetic topology inferred from SVDquartets and Neighbour-Net, along with *F*_ST,_ *d*_XY_, and gene flow signals identified using *D*-statistics and *f*_4_-ratio (Supplementary Note 1), the demographic history of pear populations was investigated.

For Occidental pears, alternative divergence and gene flow scenarios were explored between wild and cultivated populations. Five divergence scenarios (two in which perry derives from dessert and three in which dessert and perry diverge separately from wild populations) were designed covering the two cultivar populations (dessert and perry *P. communis*). Under each divergence scenario, a set of 12 demographic models was constructed combining possible gene flow between wild *P. pyraster* and *P. caucasica*, with or without gene flow between wild and cultivated or between cultivated populations, yielding 60 alternative models (Table 2; Supplementary Fig. 30; Supplementary Table 29). For Oriental pears, to keep the simulations computationally tractable, five rounds of analysis were performed, with each round containing four or five populations, to test candidate phylogenetic relationships and gene flow. In Rounds 1 and B1, sand CN-SW or sand CN-SE was modeled together with sand CN-other, *P. pashia*, and *P. betulifolia* to test the most ancestral sand pear (*P. pyrifolia*) population. In Rounds 2 and B2, based on the best-fitting model from the first two rounds, the Japanese cultivar population was added to test its origin and possible gene flow (Supplementary Fig. 31). In Round C1, various scenarios were tested, including white pear together with sand CN-other, *P. pashia,* and *P. betulifolia.* Each Occidental and Oriental round included models with and without gene flow, yielding a total of 169 alternative models (Supplementary Figs. 30,31; Supplementary Table 29). All the folded two-dimensional site frequency spectra (SFS) were generated from SNPs using easySFS (option: -a)^106,107^. Each model was optimized through 50 independent runs with parameters including 100,000 coalescent simulations per likelihood estimation (-n 100,000), 40 cycles of the conditional maximization algorithm (-L 40), and a minimum of 10 observed SFS entries for likelihood calculation (-C 10). The best-fitting model from these runs was identified based on the Akaike Information Criterion (AIC)^108^. Because AIC values are computed from composite likelihoods that ignore linkage among SNPs, we used them as relative support only. Parameter uncertainty was assessed by calculating 95% confidence intervals (CIs) using a nonparametric block bootstrap analysis (Supplementary Note 1). Based on Rounds 2, B2, and C1, we additionally tested models in which *P. ussuriensis* replaced *P. pashia* as the ancestral proxy (Rounds 2 ussu, B2 ussu, and C1 ussu; Supplementary Table 30).

### Detection of positive selection signatures

Positive selection during domestication leaves distinct genomic footprints in LD, the SFS, and diversity patterns. Because selection can generate both peaks and troughs in diversity^109^, complementary statistics were combined rather than relying solely on diversity-based measures (e.g., *F*_ST_, *π*). Selective sweeps were detected using *ω* from OmegaPlus v3.0.3^37^ and unbiased Tajima’s *D* from pixy v2.0.0beta^38^. SNP datasets were mapped to lineage-matched references (*P. communis* for Occidental, *P. pyrifolia* for Oriental), masking low-mappability and centromeric regions.

For OmegaPlus, the grid number was set to 1,000th of the chromosome length (∼1 kb), with 10–200-kb window limits. Candidate *ω* thresholds were scanned across the 95th–99.9th percentiles, and candidate Tajima’s *D* thresholds across the 0.1–5% left tail, after excluding regions with >20% overlap with low-mappability or centromeric regions (Supplementary Figs. 39–46). Tajima’s *D* was estimated with pixy from “all-sites” VCFs in 1-kb windows, excluding sites with >20% missing data (Supplementary Figs. 43,44). Cross-lineage chromosome numbering was harmonized using an MCScanX/minimap2 synteny map (Supplementary Note 1). Enrichment of *f*_d_/*f*_dM_ introgression outliers within shared positive-selection regions was assessed by circular-permutation and Fisher’s exact tests (Supplementary Note 2).

Candidate genes were defined as those whose coding or 1-kb upstream regions overlapped with outlier regions. Thresholds (*ω* > 99.45th percentile; Tajima’s *D* < 0.40th percentile) were chosen based on 1,000 permutations with the R package regioneR^110^, yielding *p* < 0.001 across all populations (Supplementary Figs. 41,42,45,46). No functional or phenotypic information was used to define candidate regions. Overlap between these candidate genes and *f*_d_/*f*_dM_ outlier windows was then examined to identify selection candidates coinciding with local introgression signals (Supplementary Note 2). Selection candidates were additionally tested for enrichment within protein clusters derived from unsupervised embedding-based clustering, both within and across populations (Supplementary Note 3).

### Gene annotation of the two reference genomes

Gene structure predictions were obtained from previous reports^79,80^. Functional annotation was performed with eggNOG-mapper v2.1.12 ^111^ using a custom Viridiplantae sub-database to assign Gene Ontology (GO) terms and Kyoto Encyclopedia of Genes and Genomes (KEGG) pathways. To refine these annotations, *Pyrus* protein sequences were retrieved from UniProtKB and *Arabidopsis thaliana* models (version 20110103) from the Arabidopsis Information Resource (TAIR), and BLASTp v2.15.0 searches (*E*-value < 1e−5) were conducted using both datasets. GO and KEGG identifiers were interpreted and enriched using *Arabidopsis thaliana* GO annotations (release 2024-11-03; Gene Ontology Consortium) and *Pyrus* KEGG pathways (https://www.kegg.jp) (Supplementary Tables 9–12). Enrichment analyses were conducted with the R packages AnnotationForge, clusterProfiler, and enrichplot^112^ using *p*-value and *q*-value thresholds of 0.05 (Supplementary Tables 10–12).

### Mutation load analysis

To assess the deleterious genetic burden in the pear populations, SIFT scores for SNP sites were calculated using the Sorting Intolerant From Tolerant (SIFT 4G) algorithm^40^. To facilitate the installation of SIFT 4G and its dependencies without root access, we developed a Singularity container recipe (SIFT4G_Create_Genomic_DB.def) available on GitHub (see Data Availability). The SIFT genomic databases for both reference genome assemblies were precomputed against UniRef90 (2025 release)^113^ reference sequences. The resulting databases covered 98.8% and 99.9% of possible amino acid substitutions with SIFT scores, with 79.6% and 87.7%, respectively, having high-confidence predictions. Using these databases, variants remaining after masking low-mappability and centromeric regions were annotated as synonymous or nonsynonymous using the SIFT 4G annotator. Nonsynonymous variants were further classified as deleterious (SIFT < 0.05) or tolerated (SIFT ≥ 0.05), excluding low-confidence deleterious entries (SIFT median sequence information > 3.5). Synonymous variants were used as the reference category. A custom script was used to count alleles for each annotated variant per genotype and population and to distinguish homozygous from heterozygous states. The robustness of the burden estimates to sequencing depth, genotype missingness, and SIFT annotation coverage was assessed in Supplementary Table 18. Loss-of-function (LoF) variants were annotated separately with SnpEff and VEP^114^, and *F*_ROH_ was estimated with ROHan^115^ (Supplementary Note 2).

### TE library building, TE annotation, and TE polymorphism detection

Two TE libraries were built *de novo* from each reference whole-genome assembly using the TEdenovo pipeline^116^ from REPET v3.0 (https://urgi.versailles.inra.fr/Tools/REPET), identifying 53.28% and 52.93% repeat content in *P. pyrifolia* (PPY) and *P. communis* (PCOM), respectively. The TE libraries were automatically curated using “the second TEannot process”^117^. Each genome reference assembly was then annotated using TEannot from the REPET package v3.0^118^. TE classification was performed using PASTEC v2.0^119^. MEGAnE^120^ was used to detect mobile element (ME) variants. Using the -rep option, we ensured that MEGAnE specifically targeted TEs from our TE library. The TE GFF annotations from REPET were converted to RepeatMasker format to be compatible with the MEGAnE tool. For the Occidental populations, the *P. communis* genome assembly was used as the reference, against which short reads from 102 samples (67 pure) were mapped with sequencing depths ranging from 18× to 50× (Supplementary Tables 1,4). For the Oriental populations, the *P. pyrifolia* whole-genome assembly served as the reference, and short reads from 114 samples (82 pure) were mapped with sequencing depths^121^ also ranging from 18× to 50× (Supplementary Tables 1,4). In MEGAnE, the non-reference TE insertions (named mobile element insertions or MEIs in this tool) are encoded such that the alternative allele indicates the presence of a TE. Therefore, the genotype values can be used directly. By contrast, the reference TE insertions (named mobile element absences or MEAs) are encoded in the opposite manner: the reference allele indicates presence. To make the two datasets comparable, the MEA genotypes were inverted before counting, ensuring that “presence” consistently represented a TE allele in both cases. Diploid genotypes with one missing allele were treated as non-missing; e.g., 1/. was imputed as 1/0 and 0/. as 0/0. Associations between TIPs and positive selection were tested as described in Supplementary Note 4 (cluster-stratified Cochran–Mantel–Haenszel tests, covariate-adjusted logistic GEE models, and 1,000 matched control gene sets). Detailed information on the visualization of TE genotypes is provided in the Supplementary Note 1 (“Analysis of TE polymorphisms”).

## Data availability

Raw whole-genome short-read data generated in this study have been deposited in the European Nucleotide Archive (ENA) at EMBL-EBI under accession number PRJEB104237 (https://www.ebi.ac.uk/ena/browser/view/PRJEB104237). The SNP data have been deposited on Zenodo (<u>DOI: 10.5281/zenodo.17673915</u>). The TE consensus sequences and annotations are available on RepetDB^122^ (https://urgi.versailles.inrae.fr/repetdb). Source Data are provided with this paper.

## Code availability

All custom scripts developed and used in this study are freely available. All scripts, pipelines, and analysis workflows are hosted and maintained in the GitHub repository (https://github.com/CornilleEclecticLab/pear-SNP).

## Acknowledgments

We thank Pierre Baduel, Timothée Flutre, and Aurélie Hua-Van for their valuable discussions and helpful suggestions. We thank the Plant Bioinformatics Facility, BioinfOmics, INRAE, Université Paris-Saclay, the GenoToul bioinformatics facility, the France National Bioinformatics Infrastructure (IFB), and the New York University Abu Dhabi (NYUAD) High Performance Computing (HPC) Center for generously providing computing resources. We thank Y. Hurand and T. Griaud for help sampling *P. pyraster,* which turned out to be introgressed with *P. communis*. We thank Gayle Volk, Serena Steffensmeier, and Nahla Victor Bassil from the USDA for providing *P. caucasica* samples. The authors thank the Biological Resource Center “RosePom - Pome Fruits and Roses” (https://www6.angers-nantes.inrae.fr/irhs/Ressources-mutualisees/Ressources-genetiques/CRB-Fruits-a-pepins-et-rosier) and associated staff for maintaining the plant material and associated datasets used in the present study.

## Author contributions

A.C. and Y.T. obtained funding, and A.C. and Y.T. designed the experiments; Y.N., X.C., S.L., Y.T., and A.C. prepared the materials for genome resequencing; Y.N., X.C., J.C., S.S., and A.C. analyzed the data; all co-authors discussed the results; Y.N., X.C., S.S., and A.C. wrote the manuscript with critical input from other co-authors.

## References

1. Miller, A. J. & Gross, B. L. From forest to field: Perennial fruit crop domestication. American Journal of Botany 98, 1389–1414 (2011).

2. Purugganan, M. D. What is domestication? Trends in Ecology & Evolution 37, 663–671 (2022).

3. Purugganan, M. D. & Fuller, D. Q. The nature of selection during plant domestication. Nature 457, 843–848 (2009).

4. Cornille, A., Giraud, T., Smulders, M. J. M., Roldán-Ruiz, I. & Gladieux, P. The domestication and evolutionary ecology of apples. Trends in Genetics 30, 57–65 (2014).

5. Gaut, B. S., Díez, C. M. & Morrell, P. L. Genomics and the Contrasting Dynamics of Annual and Perennial Domestication. Trends in Genetics 31, 709–719 (2015).

6. Gross, B. L., Henk, A. D., Richards, C. M., Fazio, G. & Volk, G. M. Genetic diversity in Malus ×domestica (Rosaceae) through time in response to domestication. American Journal of Botany 101, 1770–1779 (2014).

7. Mesnil, A. et al. Genomic architecture of the self-incompatibility locus in apple provides insights into the evolution of collaborative non-self recognition. 2025.10.28.683340 Preprint at 10.1101/2025.10.28.683340 (2025).

8. Chen, X. et al. Gene flow from the European wild apple and selection shaped the domesticated apple genome. Current Biology 36, 2104–2118.e7 (2026).

9. Cornille, A. et al. A Multifaceted Overview of Apple Tree Domestication. Trends in Plant Science 24, 770–782 (2019).

10. Julca, I. et al. Genomic evidence for recurrent genetic admixture during the domestication of Mediterranean olive trees (Olea europaea L.). BMC Biol 18, 148 (2020).

11. Duan, N. et al. Genome re-sequencing reveals the history of apple and supports a two-stage model for fruit enlargement. Nat Commun 8, 249 (2017).

12. Yu, Y. et al. Genome re-sequencing reveals the evolutionary history of peach fruit edibility. Nat Commun 9, 5404 (2018).

13. Dong, Y. et al. Dual domestications and origin of traits in grapevine evolution. Science 379, 892–901 (2023).

14. Feschotte, C., Jiang, N. & Wessler, S. R. Plant transposable elements: where genetics meets genomics. Nat Rev Genet 3, 329–341 (2002).

15. Gaut, B. S. & Ross-Ibarra, J. Selection on Major Components of Angiosperm Genomes. Science 320, 484–486 (2008).

16. Challice, J. S. & Westwood, M. N. Numerical taxonomic studies of the genus Pyrus using both chemical and botanical characters. Botanical Journal of the Linnean Society 67, 121–148 (1973).

17. Zheng, X. et al. Phylogeny and evolutionary histories of Pyrus L. revealed by phylogenetic trees and networks based on data from multiple DNA sequences. Molecular Phylogenetics and Evolution 80, 54–65 (2014).

18. Silva, G. J., Souza, T. M., Barbieri, R. L. & Costa de Oliveira, A. Origin, Domestication, and Dispersing of Pear (Pyrus spp.). Advances in Agriculture 2014, e541097 (2014).

19. Bao, L., Chen, K., Zhang, D., Li, X. & Teng, Y. An assessment of genetic variability and relationships within Asian pears based on AFLP (amplified fragment length polymorphism) markers. Scientia Horticulturae 116, 374–380 (2008).

20. Teng, Y., Yu, P., Bai, S. & Jiang, S. The origin of Asian pear cultivars inferred from DNA markers. Acta Hortic. 1–6 (2021) doi:10.17660/ActaHortic.2021.1308.1.

21. Hiwasa, K. et al. European, Chinese and Japanese pear fruits exhibit differential softening characteristics during ripening. J Exp Bot 55, 2281–2290 (2004).

22. Mou, Y. et al. Identification and Evaluation of Flesh Texture of Crisp Pear Fruit Based on Penetration Test Using Texture Analyzer. Horticulturae 11, 359 (2025).

23. Asanidze, Z., Akhalkatsi, M., Henk, A. D., Richards, C. M. & Volk, G. M. Genetic relationships between wild progenitor pear (*Pyrus* L.) species and local cultivars native to Georgia, South Caucasus. *Flora - Morphology, Distribution*, Functional Ecology of Plants 209, 504–512 (2014).

24. Wagner, I. & Wagner, S. Identity of Pyrus pyraster (L.) Burgsdorf, 1787 and implications for conservation and silviculture. Forestry: An International Journal of Forest Research cpae059 (2025) doi:10.1093/forestry/cpae059.

25. Sun, M. et al. Haplotype-resolved, gap-free genome assemblies provide insights into the divergence between Asian and European pears. Nat Genet 57, 2040–2051 (2025).

26. Wu, J. et al. Diversification and independent domestication of Asian and European pears. Genome Biol 19, 77 (2018).

27. Zhang, X. et al. Genomic insights into deleterious mutations and their impact on agronomic traits during pear domestication. Horticulture Research uhaf140 (2025) doi:10.1093/hr/uhaf140.

28. Teng, Y., Liu, J. & Hu, C. Genetic diversity of *Pyrus pashia* (*Rosaceae*) revealed by microsatellite loci. Acta Hortic. 21–26 (2018) doi:10.17660/ActaHortic.2018.1190.4.

29. Teng, Y. THE PEAR INDUSTRY AND RESEARCH IN CHINA. Acta Hortic. 161–170 (2011) doi:10.17660/ActaHortic.2011.909.16.

30. Wang, Y. & Paterson, A. H. Loquat (Eriobotrya japonica (Thunb.) Lindl) population genomics suggests a two-staged domestication and identifies genes showing convergence/parallel selective sweeps with apple or peach. The Plant Journal 106, 942–952 (2021).

31. Su, W. et al. Polyploidy underlies co-option and diversification of biosynthetic triterpene pathways in the apple tribe. Proceedings of the National Academy of Sciences 118, e2101767118 (2021).

32. Lipson, M. Applying f4-statistics and admixture graphs: Theory and examples. Molecular Ecology Resources 20, 1658–1667 (2020).

33. Reich, D., Thangaraj, K., Patterson, N., Price, A. L. & Singh, L. Reconstructing Indian population history. Nature 461, 489–494 (2009).

34. Corrick, E. C. et al. Synchronous timing of abrupt climate changes during the last glacial period. Science 369, 963–969 (2020).

35. Excoffier, L. et al. fastsimcoal2: demographic inference under complex evolutionary scenarios. Bioinformatics 37, 4882–4885 (2021).

36. Liu, J. et al. Genetic structure and phylogeography of Pyrus pashia L. (Rosaceae) in Yunnan Province, China, revealed by chloroplast DNA analyses. Tree Genetics & Genomes 9, 433–441 (2013).

37. Alachiotis, N., Stamatakis, A. & Pavlidis, P. OmegaPlus: a scalable tool for rapid detection of selective sweeps in whole-genome datasets. Bioinformatics 28, 2274–2275 (2012).

38. Bailey, N., Stevison, L. & Samuk, K. Correcting for Bias in Estimates of θw and Tajima’s D From Missing Data in Next-Generation Sequencing. Molecular Ecology Resources 25, e14104 (2025).

39. Tsuwamoto, R., Fukuoka, H. & Takahata, Y. GASSHO1 and GASSHO2 encoding a putative leucine-rich repeat transmembrane-type receptor kinase are essential for the normal development of the epidermal surface in Arabidopsis embryos. The Plant Journal 54, 30–42 (2008).

40. Vaser, R., Adusumalli, S., Leng, S. N., Sikic, M. & Ng, P. C. SIFT missense predictions for genomes. Nat Protoc 11, 1–9 (2016).

41. Lu, J. et al. The accumulation of deleterious mutations in rice genomes: a hypothesis on the cost of domestication. Trends in Genetics 22, 126–131 (2006).

42. Purugganan, M. D. Evolutionary Insights into the Nature of Plant Domestication. Current Biology 29, R705–R714 (2019).

43. Teng, Y., Tanabe, K., Tamura, F. & Itai, A. Genetic Relationships of Pyrus Species and Cultivars Native to East Asia Revealed by Randomly Amplified Polymorphic DNA Markers. jashs 127, 262–270 (2002).

44. Zhang, M.-Y. et al. Genome-wide association studies provide insights into the genetic determination of fruit traits of pear. Nat Commun 12, 1144 (2021).

45. Qi, X.-S., Yuan, N., Comes, H. P., Sakaguchi, S. & Qiu, Y.-X. A strong ‘filter’ effect of the East China Sea land bridge for East Asia’s temperate plant species: inferences from molecular phylogeography and ecological niche modelling of Platycrater arguta (Hydrangeaceae). BMC Evol Biol 14, 41 (2014).

46. Iketani, H. Native fruit tree genetic resources in Japan. Breed Sci 66, 82–89 (2016).

47. Li, Y., Zhang, J., Wang, S., Zhang, Y. & Yang, M. The Distribution and Origins of Pyrus hopeiensis-“Wild Plant With Tiny Population” Using Whole Genome Resequencing. Front Plant Sci 12, 668796 (2021).

48. Wu, J. et al. The genome of the pear (Pyrus bretschneideri Rehd.). Genome Res. 23, 396–408 (2013).

49. Li, Q. et al. Unbiased subgenome evolution following a recent whole-genome duplication in pear (Pyrus bretschneideri Rehd.). Horticulture Research 6, 34 (2019).

50. Allaby, R. Integrating the processes in the evolutionary system of domestication. J Exp Bot 61, 935–944 (2010).

51. Cornille, A. et al. New Insight into the History of Domesticated Apple: Secondary Contribution of the European Wild Apple to the Genome of Cultivated Varieties. PLOS Genetics 8, e1002703 (2012).

52. Li, M. et al. Characterization of the lipoxygenase (LOX) gene family in the Chinese white pear (Pyrus bretschneideri) and comparison with other members of the Rosaceae. BMC Genomics 15, 444 (2014).

53. Mwaniki, M. W. et al. Expression characteristics of seven members of the β-galactosidase gene family in ‘La France’ pear (Pyrus communis L.) fruit during growth and their regulation by 1-methylcyclopropene during postharvest ripening. Postharvest Biology and Technology 36, 253–263 (2005).

54. Wang, Z. et al. Genome-wide identification and characterization of active ingredients related β-Glucosidases in Dendrobium catenatum. BMC Genomics 23, 612 (2022).

55. Kim, S.-J. et al. The synthesis of xyloglucan, an abundant plant cell wall polysaccharide, requires CSLC function. Proceedings of the National Academy of Sciences 117, 20316–20324 (2020).

56. Lund, C. H. et al. Pectin Synthesis and Pollen Tube Growth in Arabidopsis Involves Three GAUT1 Golgi-Anchoring Proteins: GAUT5, GAUT6, and GAUT7. Front. Plant Sci. 11, (2020).

57. Brini, F., Hanin, M., Mezghani, I., Berkowitz, G. A. & Masmoudi, K. Overexpression of wheat Na+/H+ antiporter TNHX1 and H+-pyrophosphatase TVP1 improve salt- and drought-stress tolerance in Arabidopsis thaliana plants. J Exp Bot 58, 301–308 (2007).

58. Chaumont, F. & Tyerman, S. D. Aquaporins: Highly Regulated Channels Controlling Plant Water Relations. Plant Physiol 164, 1600–1618 (2014).

59. Kleine, T., Lockhart, P. & Batschauer, A. An Arabidopsis protein closely related to Synechocystis cryptochrome is targeted to organelles. The Plant Journal 35, 93–103 (2003).

60. Uchiyama, K. et al. Climate-Associated Genetic Variation and Projected Genetic Offsets for Cryptomeria japonica D. Don Under Future Climate Scenarios. Evolutionary Applications 18, e70077 (2025).

61. Gonzalez, A., Zhao, M., Leavitt, J. M. & Lloyd, A. M. Regulation of the anthocyanin biosynthetic pathway by the TTG1/bHLH/Myb transcriptional complex in Arabidopsis seedlings. Plant J 53, 814–827 (2008).

62. Huang, B. et al. The MdMYB44-MdTPR1 repressive complex inhibits MdCCD4 and MdCYP97A3 expression through histone deacetylation to regulate carotenoid biosynthesis in apple. The Plant Journal 119, 540–556 (2024).

63. Ireland, H. S. et al. Ethylene Regulates Apple (Malus × domestica) Fruit Softening Through a Dose × Time-Dependent Mechanism and Through Differential Sensitivities and Dependencies of Cell Wall-Modifying Genes. Plant Cell Physiol 55, 1005–1016 (2014).

64. Yao, G. et al. Map-based cloning of the pear gene MYB114 identifies an interaction with other transcription factors to coordinately regulate fruit anthocyanin biosynthesis. Plant J 92, 437–451 (2017).

65. Wu, S. et al. A common genetic mechanism underlies morphological diversity in fruits and other plant organs. Nat Commun 9, 4734 (2018).

66. Jia, Y., Yuan, Y., Zhang, Y., Yang, S. & Zhang, X. Extreme expansion of NBS-encoding genes in Rosaceae. BMC Genetics 16, 48 (2015).

67. Lv, L. et al. A TIR-NBS-LRR Gene MdTNL1 Regulates Resistance to Glomerella Leaf Spot in Apple. International Journal of Molecular Sciences 23, 6323 (2022).

68. Gaut, B. S., Seymour, D. K., Liu, Q. & Zhou, Y. Demography and its effects on genomic variation in crop domestication. Nature Plants 4, 512–520 (2018).

69. Zhou, Y., Massonnet, M., Sanjak, J. S., Cantu, D. & Gaut, B. S. Evolutionary genomics of grape (Vitis vinifera ssp. vinifera) domestication. Proceedings of the National Academy of Sciences 114, 11715–11720 (2017).

70. Mudge, K., Janick, J., Scofield, S. & Goldschmidt, E. E. A History of Grafting. in *Horticultural Reviews* 437–493 (John Wiley & Sons, Ltd, 2009). doi:10.1002/9780470593776.ch9.

71. Zhang, L. et al. A high-quality apple genome assembly reveals the association of a retrotransposon and red fruit colour. Nat Commun 10, 1494 (2019).

72. Saidi, S., Blaison, M., del Pilar Rodríguez-Ordóñez, M., Confais, J. & Quesneville, H. A reference-free pipeline for detecting shared transposable elements from pan-genomes to retrace their dynamics in a species. Genome Biol (2026) doi:10.1186/s13059-026-03984-5.

73. Alam, O. & Purugganan, M. D. Domestication and the evolution of crops: variable syndromes, complex genetic architectures, and ecological entanglements. The Plant Cell 36, 1227–1241 (2024).

74. Meyer, R. S., DuVal, A. E. & Jensen, H. R. Patterns and processes in crop domestication: an historical review and quantitative analysis of 203 global food crops. New Phytol 196, 29–48 (2012).

75. Labbancz, J. & Dhingra, A. Pangenome Analysis of Pyrus Reveals Potential Role of Transcription Factors in Recent Adaptation to Arid Environments. 2026.01.20.700718 Preprint at 10.64898/2026.01.20.700718 (2026).

76. Chen, S., Zhou, Y., Chen, Y. & Gu, J. fastp: an ultra-fast all-in-one FASTQ preprocessor. Bioinformatics 34, i884–i890 (2018).

77. Andrews, S. FastQC: A Quality Control Tool for High Throughput Sequence Data. (2010).

78. Ewels, P., Magnusson, M., Lundin, S. & Käller, M. MultiQC: summarize analysis results for multiple tools and samples in a single report. Bioinformatics 32, 3047–3048 (2016).

79. Gao, Y. et al. High-quality genome assembly of ‘Cuiguan’ pear (*Pyrus pyrifolia*) as a reference genome for identifying regulatory genes and epigenetic modifications responsible for bud dormancy. Hortic Res 8, 1–16 (2021).

80. Linsmith, G. et al. Pseudo-chromosome-length genome assembly of a double haploid ‘Bartlett’ pear (Pyrus communis L.). Gigascience 8, giz138 (2019).

81. Vasimuddin, Md., Misra, S., Li, H. & Aluru, S. Efficient Architecture-Aware Acceleration of BWA-MEM for Multicore Systems. in 2019 *IEEE International Parallel and Distributed Processing Symposium (IPDPS)* 314–324 (IEEE, Rio de Janeiro, Brazil, 2019). doi:10.1109/IPDPS.2019.00041.

82. Danecek, P. et al. Twelve years of SAMtools and BCFtools. Gigascience 10, giab008 (2021).

83. McKenna, A. et al. The genome analysis toolkit: a mapreduce framework for analyzing next-generation DNA sequencing data. Genome Res. 20, 1297–1303 (2010).

84. Manichaikul, A. et al. Robust relationship inference in genome-wide association studies. Bioinformatics 26, 2867–2873 (2010).

85. Purcell, S. et al. PLINK: A Tool Set for Whole-Genome Association and Population-Based Linkage Analyses. The American Journal of Human Genetics 81, 559–575 (2007).

86. Cingolani, P. et al. A program for annotating and predicting the effects of single nucleotide polymorphisms, SnpEff: SNPs in the genome of Drosophila melanogaster strain w1118; iso-2; iso-3. Fly (Austin) 6, 80–92 (2012).

87. Raj, A., Stephens, M. & Pritchard, J. K. fastSTRUCTURE: Variational Inference of Population Structure in Large SNP Data Sets. Genetics 197, 573–589 (2014).

88. Kopelman, N. M., Mayzel, J., Jakobsson, M., Rosenberg, N. A. & Mayrose, I. Clumpak: a program for identifying clustering modes and packaging population structure inferences across K. Mol Ecol Resour 15, 1179–1191 (2015).

89. Francis, R. M. pophelper: an R package and web app to analyse and visualize population structure. Mol Ecol Resour 17, 27–32 (2017).

90. Puechmaille, S. J. The program structure does not reliably recover the correct population structure when sampling is uneven: subsampling and new estimators alleviate the problem. Mol Ecol Resour 16, 608–627 (2016).

91. Huson, D. H. & Bryant, D. Application of Phylogenetic Networks in Evolutionary Studies. Molecular Biology and Evolution 23, 254–267 (2006).

92. Wickham, H. Ggplot2: Elegant Graphics for Data Analysis. (Springer, New York, NY, 2009). doi:10.1007/978-0-387-98141-3.

93. Pockrandt, C., Alzamel, M., Iliopoulos, C. S. & Reinert, K. GenMap: ultra-fast computation of genome mappability. Bioinformatics 36, 3687–3692 (2020).

94. Xu, D. et al. CentIER: Accurate centromere identification for plant genomes. Plant Communications 5, 101046 (2024).

95. Nei, M. & Li, W. H. Mathematical model for studying genetic variation in terms of restriction endonucleases. Proc Natl Acad Sci U S A 76, 5269–5273 (1979).

96. Wakeley, J. H. Coalescent Theory: An Introduction. (Roberts & Co, Greenwood Village, 2009).

97. Weir, B. S. & Cockerham, C. C. Estimating F-Statistics for the Analysis of Population Structure. Evolution 38, 1358–1370 (1984).

98. Korunes, K. L. & Samuk, K. pixy: Unbiased estimation of nucleotide diversity and divergence in the presence of missing data. Molecular Ecology Resources 21, 1359–1368 (2021).

99. Nei, M. Analysis of Gene Diversity in Subdivided Populations. Proc Natl Acad Sci U S A 70, 3321–3323 (1973).

100. Catchen, J., Hohenlohe, P. A., Bassham, S., Amores, A. & Cresko, W. A. Stacks: an analysis tool set for population genomics. Molecular Ecology 22, 3124–3140 (2013).

101. Chifman, J. & Kubatko, L. Quartet Inference from SNP Data Under the Coalescent Model. Bioinformatics 30, 3317–3324 (2014).

102. Chifman, J. & Kubatko, L. Identifiability of the unrooted species tree topology under the coalescent model with time-reversible substitution processes, site-specific rate variation, and invariable sites. Journal of Theoretical Biology 374, 35–47 (2015).

103. Zhang, C., Nielsen, R. & Mirarab, S. CASTER: Direct species tree inference from whole-genome alignments. Science 387, eadk9688 (2025).

104. Terhorst, J., Kamm, J. A. & Song, Y. S. Robust and scalable inference of population history from hundreds of unphased whole genomes. Nat Genet 49, 303–309 (2017).

105. Malinsky, M., Matschiner, M. & Svardal, H. Dsuite - Fast D-statistics and related admixture evidence from VCF files. Molecular Ecology Resources 21, 584–595 (2021).

106. Coffman, A. J., Hsieh, P. H., Gravel, S. & Gutenkunst, R. N. Computationally Efficient Composite Likelihood Statistics for Demographic Inference. Mol Biol Evol 33, 591–593 (2016).

107. Gutenkunst, R. N., Hernandez, R. D., Williamson, S. H. & Bustamante, C. D. Inferring the Joint Demographic History of Multiple Populations from Multidimensional SNP Frequency Data. PLOS Genetics 5, e1000695 (2009).

108. Akaike, H. A new look at the statistical model identification. IEEE Transactions on Automatic Control 19, 716–723 (1974).

109. Jasper, R. J. & Yeaman, S. Local adaptation can cause both peaks and troughs in nucleotide diversity within populations. G3 Genes|Genomes|Genetics 14, jkae225 (2024).

110. Gel, B. et al. regioneR: an R/Bioconductor package for the association analysis of genomic regions based on permutation tests. Bioinformatics 32, 289–291 (2016).

111. Huerta-Cepas, J. et al. eggNOG 5.0: a hierarchical, functionally and phylogenetically annotated orthology resource based on 5090 organisms and 2502 viruses. Nucleic Acids Research 47, D309–D314 (2019).

112. Wu, T. et al. clusterProfiler 4.0: A universal enrichment tool for interpreting omics data. Innovation (Camb) 2, 100141 (2021).

113. Suzek, B. E., Huang, H., McGarvey, P., Mazumder, R. & Wu, C. H. UniRef: comprehensive and non-redundant UniProt reference clusters. Bioinformatics 23, 1282–1288 (2007).

114. McLaren, W. et al. The Ensembl Variant Effect Predictor. Genome Biol. 17, 122 (2016).

115. Renaud, G., Hanghøj, K., Korneliussen, T. S., Willerslev, E. & Orlando, L. Joint estimates of heterozygosity and runs of homozygosity for modern and ancient samples. Genetics 212, 587–614 (2019).

116. Flutre, T., Duprat, E., Feuillet, C. & Quesneville, H. Considering Transposable Element Diversification in De Novo Annotation Approaches. PLOS ONE 6, e16526 (2011).

117. Jamilloux, V., Daron, J., Choulet, F. & Quesneville, H. De Novo Annotation of Transposable Elements: Tackling the Fat Genome Issue. Proceedings of the IEEE 105, 474–481 (2017).

118. Quesneville, H. et al. Combined Evidence Annotation of Transposable Elements in Genome Sequences. PLOS Computational Biology 1, e22 (2005).

119. Hoede, C. et al. PASTEC: An Automatic Transposable Element Classification Tool. PLOS ONE 9, e91929 (2014).

120. Kojima, S. et al. Mobile element variation contributes to population-specific genome diversification, gene regulation and disease risk. Nat Genet 55, 939–951 (2023).

121. Quadrana, L. et al. The *Arabidopsis thaliana* mobilome and its impact at the species level. eLife 5, e15716 (2016).

122. Amselem, J. et al. RepetDB: a unified resource for transposable element references. Mobile DNA 10, 6 (2019).

