## Supplementary Note 1 for "Contrasting genomic routes to domestication in Occidental and Oriental pears"

#### Occidental and oriental pears form distinct gene pools

Out of the 674 pear individuals initially collected, 396 non-duplicate individuals (207 newly sequenced from field and orchard collections and 189 retrieved from public databases) were retained for downstream analysis after quality control, with an average mapping depth of 19.3×. Reads were first mapped onto the high-quality *Pyrus pyrifolia* 'Cuiguan' reference genome[1], generating 17,769,455 single-nucleotide polymorphisms (SNPs) after stringent filtering (Supplementary Table 1; Supplementary Figs. 1,2). To minimize the confounding effects of selection and linkage, 21,756 synonymous and unlinked SNPs were retained for population structure analysis.

Using these markers, fastSTRUCTURE[2] revealed a clear genetic split between Occidental and Oriental pears, with only 136 admixed individuals at $K$ = 10 (Supplementary Figs. 3–6; Supplementary Table 2). This bipartition was further supported by Neighbor-Net analysis (Supplementary Fig. 7) and principal component analysis (PCA) (Supplementary Fig. 8), both of which indicated greater genetic differentiation among Oriental pears than among Occidental pears. This pattern suggests that the Oriental lineage experienced a longer or more complex domestication history.

Because *P. communis* (Occidental) and *P. pyrifolia* (Oriental) formed distinct gene pools, and higher $K$ values revealed substructure within the Oriental lineage, the dataset was partitioned into two lineages, Occidental and Oriental, using the Tianshan–Hindu Kush mountain range as a natural geographical boundary based on the origin of wild accessions (Fig. 1a,b; Supplementary Figs. 3–8). To minimize reference bias, a second round of SNP calling was performed using identical filtering criteria but with reads mapped to the *P. communis* reference genome. Unless otherwise specified, all subsequent analyses were conducted on lineage-specific datasets, with SNPs mapped and called on *P. communis* for Occidental pears (Supplementary Figs. 16–18) and *P. pyrifolia* for Oriental pears (Supplementary Figs. 19–21).

Occidental cultivated gene pools corresponded to both use classes (dessert and perry). At $K = 15$, fastSTRUCTURE resolved five well-supported clusters: two cultivated *P. communis* clusters, wild *P. pyraster*, wild *P. caucasica*, and one minor wild-species group. Within *P. communis*, dessert and perry pears formed distinct clusters. The dessert cluster comprised 20 individuals (14 with prior reported "dessert" usage, 6 unassigned). After excluding one 'other species' (likely *P. nivalis*) and three samples that deviated from the Perry core in the Neighbor-Net (Supplementary Fig. 23), the Perry cluster contained 14 individuals (12 with prior known "Perry" usage, 2 unassigned). A Fisher's exact test confirmed a strong association between cluster membership and reported use (two-sided $p = 1.04 \times 10^{-7}$). Admixed dessert pears lie between wild *P. pyraster* and cultivated *P. communis* (Fig. 1a; Supplementary Figs. 16–18). The embedding of *P. communis* perry pears within wild *P. pyraster* in the Neighbor-Net (Supplementary Fig. 23) and their proximity in the PCA plot (Fig. 1c) further support the close relationship between perry pears and *P. pyraster*. Wild *P. caucasica* formed a clearly separate cluster (Fig. 1a).

Overall, *P. communis* cultivars exhibited complex admixture patterns, reflecting contributions from dessert, perry, and wild individuals. Several admixed *P. pyraster* individuals collected in France were likely orchard escapees (Fig. 1a; Supplementary Fig. 23). Considering differences in sequencing depth, coverage, and observed heterozygosity ($H_o$) between the genome sequencing data of Occidental and Oriental pears when mapped to the two genome assemblies (Supplementary Figs. 9,10), Occidental pears were analyzed using SNP markers referenced to both

*P. pyrifolia* and *P. communis* to ensure robust inference of population structure (Supplementary Figs. 11–18). Comparative analyses at $K = 9$ (reference as *P. pyrifolia*) and $K = 15$ (reference as *P. communis*) produced consistent results, with only one individual differing in assignment (admixed in the former, pure in the latter), which was ultimately treated as admixed (Supplementary Figs. 11,16; Supplementary Tables 2,3). Therefore, analyses using both reference genomes supported consistent population structures for Occidental pears.

In Oriental pears, fastSTRUCTURE and CLUMPAK identified nine major genetic clusters at $K = 10$ (Fig. 1b; Supplementary Figs. 19–21). These included three wild populations (*P. pashia*, *P. ussuriensis*, and *P. betulifolia*) and six cultivated groups corresponding to *P. pyrifolia*, white pear (also known as *P.* × *bretschneideri*), and four sand pear clusters (CN-SW, CN-SE, CN-other, and the Japanese Nashi pear). Among cultivated pears, the sand pear clusters displayed clear geographical structuring: CN-SW and CN-SE were isolated by major mountain barriers such as the Hengduan, Himalayan, and Nanling ranges, whereas samples assigned as CN–other were widely distributed across China (Supplementary Fig. 22). The gene pools of white pear and Japanese pear exhibit strong regional differentiation. The White pear gene pool shows high admixture with other Oriental pear groups (Supplementary Fig. 22). In particular, the CN-other population occurs across nearly all production regions of China, suggesting it may represent the ancestral cluster of all regionally differentiated *P. pyrifolia* populations. The gene pool of sand pears in Eastern China (e.g., Zhejiang Province) shows substantial admixture with that of Japanese pears (Supplementary Fig. 22). Chinese sand, white, and Japanese *P. pyrifolia* cultivars formed five coherent clusters that grouped together in the Neighbor-Net and were distinct in the PCA (Fig. 1d,e; Supplementary Fig.23). The wild species *P. ussuriensis* and *P. betulifolia* (rootstock) formed genetically distinct clusters (Fig. 1d). Notably, previously published *P. betulifolia* individuals grouped inconsistently with other species and were distinct from our newly sequenced rootstock individuals, whose species identity was verified. This discrepancy suggests that some *P. betulifolia* samples reported in earlier studies were misidentified; we therefore excluded these admixed or inconsistent clusters from further analysis.

**Divergence and demographic history of Oriental and Occidental pears**

The evolutionary relationships among the 12 defined pear populations were reconstructed using SVDQuartet[3,4] with 21,237 SNPs mapped to the *Pyrus pyrifolia* reference genome (Fig. 2a). Population assignments followed those identified with fastSTRUCTURE (Fig. 1; Supplementary Table 4). Loquat (*Eriobotrya japonica*) SNPs were used as an outgroup[5,6]. The resulting phylogeny revealed a deep split between Occidental and Oriental pears, supporting separate evolutionary histories for the two cultivar groups. Within the Occidental lineage, *P. communis* dessert and perry cultivars diverged from distinct ancestral nodes, suggesting separate divergence histories. In Oriental *P. pyrifolia*, shallow divergences and low bootstrap support among several branches likely reflect historical gene flow among populations.

Historical admixture was quantified using $f_4$-ratio and $D$-statistics (ABBA–BABA tests) implemented in Dsuite[7], using *P. ussuriensis* and *P. caucasica* as outgroups for Occidental and Oriental pears, respectively (Supplementary Fig. 25; Supplementary Tables 5,6). Under strict isolation, these statistics approached zero, with divergence explained solely by drift[8,9]. Significant non-zero values indicated multiple episodes of gene flow. In Occidental pears, introgression occurred between wild and cultivated populations (*P. pyraster* and dessert *P. communis*; *P. caucasica* and perry *P. communis*) and between wild populations (*P. caucasica* and *P. pyraster*). In Oriental pears, we detected extensive gene flow between *P. pashia* and *P. pyrifolia* cultivars, but not between *P. pashia* and *P. ussuriensis*. Introgression among cultivars (e.g., between white and sand pear populations) coincided with low phylogenetic bootstrap support.

**Demographic simulations using fastsimcoal2**

We reconstructed the demographic history of pear domestication using the coalescent-based simulator fastsimcoal2 v2.8[10]. The objectives were to infer (i) the tempo of domestication events—successive versus independent divergence from a shared ancestral pool, (ii) whether gene flow occurred among specific wild and cultivated populations during domestication. Scenarios were established based on population clusters identified by fastSTRUCTURE, complemented by genetic differentiation ($F_{ST}$, $d_{XY}$), introgression (*D*-statistics and $f_4$-ratio; Fig. 2a), and phylogenetic relationships inferred by SVDquartets (Fig. 2a). All scripts are available at:

- https://github.com/CornilleEclecticLab/pear-SNP/blob/QC/run_fastsimcoal2_refer_on_communis_k_IFB/
- https://github.com/CornilleEclecticLab/pear-SNP/tree/QC/run_fastsimcoal2_refer_on_pyrifolia
- https://github.com/CornilleEclecticLab/pear-SNP/tree/QC/run_fastsimcoal2_refer_on_pyrifolia_round1_c_4pop_IFB

In addition, we provide example *.obs files, the inputs for fastsimcoal, along with step-by-step instructions for running the analyses on a computing cluster:

- https://github.com/CornilleEclecticLab/pear-SNP/tree/QC/run_fastsimcoal2_refer_on_pyrifolia_round1_c_4pop_IFB

### Demographic simulations using fastsimcoal2: Occidental pears

Occidental pears were divided into four main populations. To maintain computational feasibility, *P. communis* was modeled jointly with its putative wild progenitors, *P. pyraster* and *P. caucasica*. The phylogeny did not fully resolve relationships between *P. pyraster* and the cultivated *P. communis* lineages (dessert and perry types, Fig. 2a). However, genetic distance (Supplementary Fig. 28) and network analyses (Fig. 1) indicated that perry pears were genetically closer to *P. pyraster* than to dessert pears. Furthermore, no signal of recent gene flow was detected between perry pears and *P. pyraster* (Supplementary Fig. 25), suggesting shared ancestry rather than secondary introgression.

We therefore tested alternative divergence scenarios: (i) independent divergence of the dessert and perry pears from *P. pyraster* or *P. caucasica* or (ii) secondary domestication of one cultivar from the other (Supplementary Fig. 30). Five divergence topologies were designed to represent the possible origins of the two cultivar populations, each combined with models incorporating gene flow between *P. pyraster* and *P. caucasica*, and between wild and cultivated populations. Five divergence topologies (two in which perry derives from dessert, three in which dessert and perry diverge separately from wild populations) were each combined with 12 gene-flow configurations, giving 60 alternative models (Supplementary Fig. 30). The model parameters used for these models are provided at:

- https://github.com/CornilleEclecticLab/pear-SNP/tree/QC/run_fastsimcoal2_refer_on_communis_k_IFB/run_fastsimcoal2/analysis/DemoInfer/demo_files/west

### Demographic simulations using fastsimcoal2: Oriental pears

The Oriental dataset initially comprised eight populations; *P. ussuriensis* was excluded from the main models. Previous studies reported introgression from this wild species into *P. pyrifolia* mainly in northern China[11,12]. Including this species would have markedly increased the number of parameters and migration edges to estimate, complicating model fitting and convergence. Consistent with this exclusion, we observed low nucleotide diversity ($\pi$) in *P. ussuriensis*, together with relatively high $F_{ST}$ and $d_{XY}$ values compared with cultivated populations —patterns that also support its distinct position in the species tree (Fig. 2a; Supplementary Figs. 27,28). To test whether this choice affected the inferred topologies, we repeated Rounds 2, B2 and C1 with *P. ussuriensis* replacing *P. pashia* as the wild root proxy (Supplementary Table 30).

The final Oriental dataset for demographic simulations comprised seven populations: two non-cultivars (*P. betulifolia* and *P. pashia*), and five cultivars (*P. pyrifolia* sand CN, sand CN-SE, sand CN-SW, white, and Japanese). As testing all populations simultaneously was computationally prohibitive, we performed five rounds of analyses using subsets of four to five populations, each representing alternative topologies and migration schemes (Supplementary Fig. 31).

Neighbor-Net (Supplementary Fig. 23) and pairwise differentiation ($F_{ST}$, $d_{XY}$; Supplementary Fig. 28) showed that Japanese pears were genetically closer to Sand pears, consistent with either gene flow or shared divergence. Assuming *P. pashia* as the ancestral population[13–15], we tested models of (i) independent divergence of Japanese pears from the wild proxy and (ii) divergence of Japanese pears from Chinese Sand pears, both with and without post-divergence gene flow. Rounds 1 and B1 modeled Sand CN-SW or CN-SE with Sand CN, *P. pashia*, and *P. betulifolia* to identify the most ancestral Sand pear population. Rounds 2 and B2 incorporated Japanese pears to test their origin and migration routes. Round C1 evaluated scenarios including White pears alongside Sand CN, *P. pashia*, and *P. betulifolia*. Each round included models with and without gene flow, resulting in a total of 109 alternative models (Supplementary Fig. 31; Supplementary Table 29). The parameter files used for these models are provided at:

- https://github.com/CornilleEclecticLab/pear-SNP/tree/QC/run_fastsimcoal2_refer_on_pyrifolia_round1_c_4pop_IFB/input/demography_models_config
- https://github.com/CornilleEclecticLab/pear-SNP/tree/QC/run_fastsimcoal2_refer_on_pyrifolia_roundB1_d_4pop_IFB/input/demography_models_config
- https://github.com/CornilleEclecticLab/pear-SNP/tree/QC/run_fastsimcoal2_refer_on_pyrifolia_round2_d_5pop_IFB/input/demography_models_config
- https://github.com/CornilleEclecticLab/pear-SNP/tree/QC/run_fastsimcoal2_refer_on_pyrifolia_roundB2_c_5pop_IFB/input/demography_models_config
- https://github.com/CornilleEclecticLab/pear-SNP/tree/QC/run_fastsimcoal2_refer_on_pyrifolia_roundC1_e_4pop_IFB/input/demography_models_config

**Demographic simulations using fastsimcoal2: Model optimization and evaluation**

Folded two-dimensional site frequency spectra (SFS) were generated from SNP data using easySFS[16,17]. Each model was optimized through 50 independent runs with parameters set to: 100,000 coalescent simulations per likelihood estimation (-n 100000), 40 conditional maximization cycles (-L 40), and a minimum of 10 observed SFS entries for likelihood calculation

(-C 10). The best-fitting model for each comparison was selected using the Akaike Information Criterion (AIC)[18]. To assess the uncertainty in the estimated parameters, we performed a nonparametric block bootstrap analysis. The SNP dataset for each round was divided into 100 contiguous blocks, and 50 bootstrapped datasets were generated by resampling blocks with replacement. Parameter estimation was repeated for the best-fitting model on each bootstrapped SFS, and 95% confidence intervals (CIs) were derived from the 2.5th and 97.5th percentiles of the bootstrap distribution.

**Identification of colinear genes**

To identify syntenic genes between the two *P. communis*[19] and *P. pyrifolia*[1] reference genomes used in this study, interspecies BLASTp (v2.15.0)[20] searches were performed with an E-value threshold of 1e−10, retaining the top five hits per query, following the MCScanX guidelines (https://github.com/wyp1125/MCScanX/blob/master/README.rst), using the putative protein sequences of each species as a reference database in separate runs. MCScanX[21] was then applied to the combined set of 75,840 genes using the reciprocal BLAST results. When using *P. pyrifolia* as the reference, 53,761 (70.9%) colinear genes were detected, whereas when using *P. communis* as the reference, 53,107 (70.0%) colinear genes were identified. The datasets were derived from 50,683 and 50,755 intercolinear gene pairs, respectively. To define syntenic genes, the intersection of the two MCScanX runs was used to identify 45,289 interspecies colinear gene pairs shared between the two genomes, corresponding to 49,725 (65.6%) genes (Supplementary Fig. 38).

**DNA extraction and short-read sequencing of new individuals**

Genomic DNA was extracted from the leaf tissues of 235 newly sequenced *Pyrus* samples of 10 (sub-) species, representing wild and cultivated pears (Supplementary Table 1), using standard commercial extraction kits, such as the NucleoSpin Plant II kit (Macherey-Nagel), following the manufacturer's instructions. The DNA was eluted in Elution Buffer PE (5 mM Tris-HCl, pH 8.5). DNA concentrations were quantified using a Nanodrop spectrophotometer. Sequencing libraries were prepared using 1.0 µg of DNA per sample. Libraries were constructed using standard DNA library prep kits and sequenced on high-throughput short-read platforms (producing 150 bp paired-end reads).

**Mapping of SNP markers to the Occidental pear genome**

Substantial genomic differences were detected between Occidental and Oriental pears, e.g., the genomic annotations revealed 37,445 "mRNA" items in the 497-Mb Occidental pear (*P. communis*) genome assembly[19] with an average length of 2,326 bp, compared to 42,622 "mRNA" items in the 541-Mb Oriental pear (*P. pyrifolia*) assembly[1] with an average length of 3,150 bp. Consequently, the above SNP-calling and filtering steps were applied to the genome sequences of the 396 non-clone individuals, using the *P. communis* genome assembly as the reference. The sequencing depth, genome coverage, and $H_o$ values were compared for both Occidental and Oriental pears when reads were mapped to either reference genome (Supplementary Figs. 9 and 10). Unless stated otherwise, analyses were based on lineage-matched reference assemblies.

**Inference of population genetic structure using passport information**

The (sub-)species names, regions, and type information (wild, cultivar, or rootstock) of pear samples were obtained from the samplers or from the original data sources. Wild (and rootstock) pears were classified into Occidental and Oriental groups according to their native distributions, with the Tianshan–Hindu Kush Mountains serving as the biogeographical boundary. Wild populations from Central Asia and regions further west, including Asia Minor, Europe, and the Mediterranean, were designated as Occidental, whereas those confined to East Asia were designated as Oriental. For cultivated pears, a more conservative approach was adopted, assigning them to Occidental or Oriental groups primarily based on genetic closeness inferred from gene pool population structure, phylogenetic analyses, and PCA, rather than their current cultivation regions, which may reflect human-mediated dispersal (Supplementary Table 1).

A Bayesian framework was adopted for posterior inference of population structure and admixture from SNP genotypes using fastSTRUCTURE[2]. $K$ values (assumed population number) were set from 2 to 16; 30 iterations were performed with random seeds for each $K$. Subsequently, the results were consolidated into consensus solutions using CLUMPAK[22] (Supplementary Tables 2,3). The R package Pophelper[23] was used to visualize the CLUMPAK results (Fig. 1a,b; Supplementary Figs. 4, 11, 16). To evaluate the optimal number of clusters, cross-validation errors were examined around and after the "elbow" of the curve, where model likelihoods plateaued (Supplementary Figs. 5, 12, 17). However, $\Delta K$ alone may not fully capture the complexity of population structure observed in fastSTRUCTURE[24]. Therefore, this statistical criterion was complemented by visually comparing bar plots across $K$ values (Supplementary Figs. 4, 6, 11, 13, 16, 18), selecting the solution in which clusters were most clearly defined, and individuals were consistently well assigned. This combined approach provided the finest resolution of genetic structure, rather than relying solely on the strongest $\Delta K$ signal.

To infer the Neighbor-Net algorithm-based tree, the 1-IBS (i.e., one minus the identity-by-state value) distance matrices were calculated using Plink v1.9 and visualized with SplitsTree4 (version 4.18.3)[25] (; Supplementary Figs. 7,14 and 23). PCA was conducted using Plink1.9[26] , and the results were visualized with the R package ggplot2[27] (Fig. 1c–e; Supplementary Figs. 8, 15). The colors assigned to each cluster in the population structure bar plots were also used for both PCA and Neighbor-Net tree plots, except for admixed individuals whose membership coefficients were all $< 0.8$ (colored in gray). Additionally, to enhance accessibility, the color schemes were carefully improved using the Adobe Color tool (https://color.adobe.com/) (Supplementary Fig. 3).

Fourteen genetic clusters were inferred across Occidental and Oriental pears using fastSTRUCTURE (Fig. 1a,b). For downstream analyses, two clusters were excluded that contained fewer than seven conspecific individuals. Within each retained population, individuals whose placements were discordant outliers in the Neighbor-Net tree relative to their fastSTRUCTURE assignments were removed (Fig. 1a,b; Supplementary Fig.23), yielding a population set with consistent structure across Neighbor-Net and fastSTRUCTURE. For Occidental pears, population assignments were resolved using SNPs independently mapped to both the *P. communis* and *P. pyrifolia* reference genomes (Supplementary Figs. 11–18; Supplementary Tables 2–4). Summary statistics were then calculated for the 12 final populations (Table 1; Supplementary Table 4). After excluding putative hybrids and accessions with unclear passport information, 229 genetically "pure" individuals (admixture proportion $\geq 80\%$) were retained for population-level analyses (Supplementary Table 4).

To mask low-mappability regions within genotypes for demographic inference, mappability profiles were first computed for both reference genomes using GenMap[28] with simulated 140-bp reads ($k$-mers, $k = 140$) and no mismatches allowed ($e = 0$). A custom post-processing script was then implemented to smooth the discrete GenMap scores by scanning the genome in 100-kb windows with 50-kb steps and calculating the mean mappability score per window. Windows with a mean mappability score > 0.9 were classified as high-mappability, with the remainder classified as low-mappability. As a result, 59.4% of the 17 *P. pyrifolia* chromosomes and 76.7% of the 17 *P. communis* chromosomes were identified as high-mappability, with the remainder considered low-mappability. The centromeric regions were also inferred using CentIER[29] for both reference genomes to obtain low-confidence regions. The predicted centromeric regions were consistently found to be subsets of low-mappability regions in both reference genomes.

Pairwise nucleotide diversity ($\pi$)[30] within each population, genetic distance ($d_{XY}$)[30,31], and genetic differentiation ($F_{ST}$)[32] were calculated between each population pair with Pixy v1.2.7.beta1[33], with windows defined by high-mappability regions, using "all sites" VCFs that contained both variant and invariant sites as input (Supplementary Figs. 27,28). Anderson-Darling and Shapiro-Wilk tests were used to assess normality (*p*-values at the 0.05 level). The Wilcoxon test was used to assess differences in π among populations (*p*-values at the 0.01 level). Additionally, a script was written to compute the genome-wide π, $d_{XY}$, and $F_{ST}$ values from the Pixy output. The genome-wide $\pi$ values were calculated by dividing the total pairwise differences by the total number of comparisons rather than by averaging window-based $\pi$ values. Using Stacks Populations v2.65[34], the observed ($H_O$) and expected ($H_E$) heterozygosity[35] values were calculated, as well as the inbreeding coefficient ($F_{IS}$), for each variant and population.

**Analysis of TE polymorphisms**

For each TE insertion identified by MEGAnE[36], presence or absence was assessed across all samples relative to a single reference genome. A quantitative 0-1-2 encoding was used to capture fine-scale genotypic variation among individuals, whereas a simplified binary (0–1) coding was applied in TE dynamics analyses across TE classes and TE orders. PCA was performed on the TE genotypes using a diploid 0-1-2 coding scheme that accounts for heterozygosity at each insertion site. Specifically, for each individual and locus, heterozygotes (0/1) were assigned a value of one, homozygous carriers (1/1) a value of two, and homozygous non-carriers (0/0) a value of zero. In the diploid genotyping scheme, the states 0/0, 0/1, and 1/1 correspond to the absence, heterozygous presence, and homozygous presence of a TE at a given locus, respectively. Both 0/1 and 1/1 were interpreted as indicative of TE presence, without distinguishing allelic dosage. Individuals were subsequently grouped by population, and a TE was considered present in a population if at least one individual in that population carried the insertion. For the heatmaps, a TE insertion was considered fixed when it occurred in only one population and was absent from all others, indicating a population-specific insertion. Conversely, a TE insertion was classified as shared when it was detected in two or more distinct populations. The "shared/fixed" ratio measures the balance between shared and population-specific TE insertions across populations. For the UpSet plots, each population was counted once per TE locus if at least one individual carried a TE insertion. To identify genes involved in immunity-related processes, we applied a custom filtering procedure using the annotation fields "Atha_description" and "gene_Symbol" extracted from Supplementary Tables 8 and 9, together with Level-2 Gene Ontology terms. GO annotations were parsed using the goatools Python package (v1.5.2) and the go-basic.obo ontology file (format 1.2; release 2025-10-10; 42,666 terms). Genes were flagged as immunity-related when at least one of the following substrings was detected: (i) "disease" or "pathogen" within Atha_description; (ii) the motif "NBS" within gene_Symbol; or (iii) any of the immunity-associated keywords "immune", "pathogen", "infection", or "disease" within Level-2 Gene Ontology annotations. The resulting binary annotation is reported in the column “Immunity”, which is provided in Supplementary Tables 21 and 22.

### References


1. Gao, Y. *et al.* High-quality genome assembly of ‘Cuiguan’ pear (*Pyrus pyrifolia*) as a reference genome for identifying regulatory genes and epigenetic modifications responsible for bud dormancy. *Hortic Res* **8**, 1–16 (2021).
2. Raj, A., Stephens, M. & Pritchard, J. K. fastSTRUCTURE: Variational Inference of Population Structure in Large SNP Data Sets. *Genetics* **197**, 573–589 (2014).
3. Chifman, J. & Kubatko, L. Quartet Inference from SNP Data Under the Coalescent Model. *Bioinformatics* **30**, 3317–3324 (2014).
4. Chifman, J. & Kubatko, L. Identifiability of the unrooted species tree topology under the coalescent model with time-reversible substitution processes, site-specific rate variation, and invariable sites. *Journal of Theoretical Biology* **374**, 35–47 (2015).
5. Wang, Y. & Paterson, A. H. Loquat (Eriobotrya japonica (Thunb.) Lindl) population genomics suggests a two-staged domestication and identifies genes showing convergence/parallel selective sweeps with apple or peach. *The Plant Journal* **106**, 942–952 (2021).
6. Su, W. *et al.* Polyploidy underlies co-option and diversification of biosynthetic triterpene pathways in the apple tribe. *Proceedings of the National Academy of Sciences* **118**, e2101767118 (2021).

7. Malinsky, M., Matschiner, M. & Svardal, H. Dsuite - Fast D-statistics and related admixture evidence from VCF files. *Molecular Ecology Resources* **21**, 584–595 (2021).
8. Reich, D., Thangaraj, K., Patterson, N., Price, A. L. & Singh, L. Reconstructing Indian population history. *Nature* **461**, 489–494 (2009).
9. Lipson, M. Applying f4-statistics and admixture graphs: Theory and examples. *Molecular Ecology Resources* **20**, 1658–1667 (2020).
10. Excoffier, L. *et al.* fastsimcoal2: demographic inference under complex evolutionary scenarios. *Bioinformatics* **37**, 4882–4885 (2021).
11. Teng, Y., Yu, P., Bai, S. & Jiang, S. The origin of Asian pear cultivars inferred from DNA markers. *Acta Hortic.* 1–6 (2021) doi:10.17660/ActaHortic.2021.1308.1.
12. Wuyun, T. *et al.* Population Structure of and Conservation Strategies for Wild Pyrus ussuriensis Maxim. in China. *PLOS ONE* **10**, e0133686 (2015).
13. Liu, J. *et al.* Genetic structure and phylogeography of Pyrus pashia L. (Rosaceae) in Yunnan Province, China, revealed by chloroplast DNA analyses. *Tree Genetics & Genomes* **9**, 433–441 (2013).
14. Teng, Y., Liu, J. & Hu, C. Genetic diversity of *Pyrus pashia* ( *Rosaceae* ) revealed by microsatellite loci. *Acta Hortic.* 21–26 (2018) doi:10.17660/ActaHortic.2018.1190.4.
15. Zheng, X. *et al.* Phylogeny and evolutionary histories of Pyrus L. revealed by phylogenetic trees and networks based on data from multiple DNA sequences. *Molecular Phylogenetics and Evolution* **80**, 54–65 (2014).
16. Coffman, A. J., Hsieh, P. H., Gravel, S. & Gutenkunst, R. N. Computationally Efficient Composite Likelihood Statistics for Demographic Inference. *Mol Biol Evol* **33**, 591–593 (2016).

17. Gutenkunst, R. N., Hernandez, R. D., Williamson, S. H. & Bustamante, C. D. Inferring the Joint Demographic History of Multiple Populations from Multidimensional SNP Frequency Data. *PLOS Genetics* **5**, e1000695 (2009).

18. Akaike, H. A new look at the statistical model identification. *IEEE Transactions on Automatic Control* **19**, 716–723 (1974).

19. Linsmith, G. *et al.* Pseudo-chromosome-length genome assembly of a double haploid 'Bartlett' pear (Pyrus communis L.). *Gigascience* **8**, giz138 (2019).

20. Camacho, C. *et al.* BLAST+: architecture and applications. *BMC Bioinformatics* **10**, 421 (2009).

21. Wang, Y. *et al.* MCScanX: a toolkit for detection and evolutionary analysis of gene synteny and collinearity. *Nucleic Acids Research* **40**, e49 (2012).

22. Kopelman, N. M., Mayzel, J., Jakobsson, M., Rosenberg, N. A. & Mayrose, I. Clumpak: a program for identifying clustering modes and packaging population structure inferences across K. *Mol Ecol Resour* **15**, 1179–1191 (2015).

23. Francis, R. M. pophelper: an R package and web app to analyse and visualize population structure. *Mol Ecol Resour* **17**, 27–32 (2017).

24. Puechmaille, S. J. The program structure does not reliably recover the correct population structure when sampling is uneven: subsampling and new estimators alleviate the problem. *Mol Ecol Resour* **16**, 608–627 (2016).

25. Huson, D. H. & Bryant, D. Application of Phylogenetic Networks in Evolutionary Studies. *Molecular Biology and Evolution* **23**, 254–267 (2006).

26. Purcell, S. *et al.* PLINK: A Tool Set for Whole-Genome Association and Population-Based Linkage Analyses. *The American Journal of Human Genetics* **81**, 559–575 (2007).

27. Wickham, H. *Ggplot2: Elegant Graphics for Data Analysis*. (Springer, New York, NY, 2009). doi:10.1007/978-0-387-98141-3.

28. Pockrandt, C., Alzamel, M., Iliopoulos, C. S. & Reinert, K. GenMap: ultra-fast computation of genome mappability. *Bioinformatics* **36**, 3687–3692 (2020).

29. Xu, D. *et al.* CentIER: Accurate centromere identification for plant genomes. *Plant Communications* **5**, 101046 (2024).

30. Nei, M. & Li, W. H. Mathematical model for studying genetic variation in terms of restriction endonucleases. *Proc Natl Acad Sci U S A* **76**, 5269–5273 (1979).

31. Wakeley, J. H. *Coalescent Theory: An Introduction*. (Roberts & Co, Greenwood Village, 2009).

32. Weir, B. S. & Cockerham, C. C. Estimating F-Statistics for the Analysis of Population Structure. *Evolution* **38**, 1358–1370 (1984).

33. Korunes, K. L. & Samuk, K. pixy: Unbiased estimation of nucleotide diversity and divergence in the presence of missing data. *Molecular Ecology Resources* **21**, 1359–1368 (2021).

34. Catchen, J., Hohenlohe, P. A., Bassham, S., Amores, A. & Cresko, W. A. Stacks: an analysis tool set for population genomics. *Molecular Ecology* **22**, 3124–3140 (2013).

35. Nei, M. Analysis of Gene Diversity in Subdivided Populations. *Proc Natl Acad Sci U S A* **70**, 3321–3323 (1973).

36. Kojima, S. *et al.* Mobile element variation contributes to population-specific genome diversification, gene regulation and disease risk. *Nat Genet* **55**, 939–951 (2023).
