## Supplementary Note 2 for "Contrasting genomic routes to domestication in Occidental and Oriental pears"

### Methods

#### Enrichment of $f_d$/$f_{dM}$ introgression outliers within shared positive-selection regions

*Trios.* Within each lineage, we retained trios that included both wild and cultivated populations and showed a significant genome-wide excess of allele sharing (*D*-statistics, $P < 0.01$; Supplementary Tables 5,6). We arranged each trio as the topology ((P1, P2), P3), with P1 and P2 as sister populations in the best-supported fastsimcoal2 scenarios. This gave one Occidental trio (P1 = *P. caucasica*, P2 = dessert *P. communis*, P3 = *P. pyraster*) and three Oriental trios (P1 = *P. pashia*; P2 = Japanese pear, sand pear CN-other or sand pear CN-SW; P3 = *P. betulifolia*).

*Introgression outliers.* We computed $f_d$ and $f_{dM}$ in sliding genomic windows (Dsuite; 50 SNPs, step 25 SNPs) for each population trio. Outlier windows were the top 5% for $f_d$ (one-tailed, as negative values are not interpretable as introgression) and the top and bottom 2.5% for $f_{dM}$. We analyzed the two $f_{dM}$ tails separately because their signs identify excess allele sharing between population pairs within the trio (positive: P2 and P3; negative: P1 and P3). Neither statistic gives the direction of gene flow.

*Enrichment tests.* We tested whether outlier windows were overrepresented within sweep regions shared by the corresponding population pair (the intersection of their sweeps, a deliberately conservative definition), restricted to windows with ≥80% of their length in high-mappability, non-centromeric regions (Supplementary Table 28); sweeps were clipped to the same space. We used: (i) a count-based test and (ii) a test of mean $f_d$/$f_{dM}$ inside versus outside sweeps, both evaluated with 10,000 chromosome-wise circular permutations that preserve chromosomal distribution and spatial clustering of windows; and (iii) a one-sided Fisher's exact test on non-overlapping windows. To control for diversity, window length, and gene density, we repeated the count-based test relative to the number of outliers expected from covariates alone (logistic regression on log $\pi$ of P1, P2, and P3 calculated with pixy in the same windows, log window length, and gene density). We corrected *P* values with the Benjamini-Hochberg procedure within each lineage across trios, metrics, and tails (Occidental, 3 tests; Oriental, 9 tests).

**Loss-of-function (LoF) variant annotation and burden**

LoF variants were annotated independently of SIFT4G with two tools, snpEff v5.1d[1] and Ensembl VEP v105[2]. The input was the same filtered SNP set used for the SIFT4G analysis. Two parallel pipelines were run against different reference assemblies: the *Pyrus communis* Bartlett DHv2.0 genome for the Occidental lineage and the *P. pyrifolia* Cuiguan reference genome for the Oriental lineage. Both tools used the same gene annotation: a local snpEff database built from the reference GFF, and VEP run in GFF/FASTA mode without a cache. CDS phases in the *P. pyrifolia* GFF were recomputed before the snpEff database was built. A SNP was considered LoF by a tool if it was assigned a stop_gained, start_lost, stop_lost, splice_donor_variant, or splice_acceptor_variant consequence on a protein-coding transcript. We retained a high-confidence (HC) LoF set: stop-gained and canonical splice-donor/acceptor SNPs called as LoF by both VEP and snpEff and additionally flagged by snpEff's LOF tag. This excluded start-lost, stop-lost, and NMD-escaping stop-gained variants. Synonymous sites were defined as sites called synonymous_variant by both tools.

Following the per-site normalization used for the SIFT4G burden, we computed for each individual the number of HC LoF alleles (two per homozygous and one per heterozygous genotype) divided by the number of synonymous sites at which the individual carried the alternative allele. This was calculated separately for all genotypes (total burden), homozygous genotypes (realized burden), and heterozygous genotypes (potential burden), within the same mappability-masked regions. Differences between wild and cultivated populations, and pairwise differences among populations within each lineage, were tested with two-sided Wilcoxon rank-sum tests. Pairwise *P* values were Holm-adjusted and summarized as compact letter displays (adjusted $P < 0.01$).

**Runs of homozygosity (ROH) analysis and realized genetic load**

Genome-wide autozygosity was characterized with ROHan v1.0.1[3], which estimates local heterozygosity and ROH state jointly from aligned reads under a hidden Markov model. Reads were filtered to remove duplicate and secondary/supplementary alignments (samtools view -F 0x600) prior to analysis. For each individual, ROHan was run on autosomes only, restricted to a mappability-masked region set, using a population-specific transition/transversion ratio pre-estimated from that population's autosomal, mask-restricted variant calls (bcftools stats) and the recommended ROH mutation-rate prior (--rohmu 1e-3). Two parallel pipelines were run against different reference assemblies: the *Pyrus communis* 'Bartlett' DHv2.0 genome for the Occidental lineage, and the *P. pyrifolia* 'Cuiguan' reference genome for the Oriental lineage. The proportion of the genome in ROH ($F_{\mathrm{ROH}}$) and its bootstrap confidence interval were taken directly from ROHan's per-sample summary output.

Following Wang et al. (2025)[4], we related $F_{\mathrm{ROH}}$ to the realized deleterious burden identified by SIFT4G in two complementary ways. First, for each individual, we computed the ratio of homozygous deleterious alleles to homozygous synonymous SNPs and regressed this ratio on $F_{\mathrm{ROH}}$ using a Gaussian linear model, assessing the Pearson correlation at two levels: within each lineage as a whole (pooling across populations), and separately within each population. Second, we tested directly whether deleterious burden is enriched within ROH. Every genotyped, SIFT4G-annotated site was intersected against each individual's own merged ROH interval set and assigned to the ROH or non-ROH compartment. Deleterious-allele burden was then compared between compartments within individuals using a paired Wilcoxon signed-rank test, separately for each reference genome and for three burden metrics (allele count, homozygous-genotype count, heterozygous-genotype count); individuals with no called ROH were excluded to keep the pairing balanced.

**Results**

**TE variation tracks demographic structure**

Wild populations exhibited lower shared/fixed ratios across most TE orders (Supplementary Fig. 60a,b), consistent with lineage-specific insertions accumulating after divergence. Cultivated populations showed elevated shared/fixed ratios, reflecting shared ancestral insertions and/or gene flow among cultivars (Supplementary Fig. 60a,b). LARDs and SINEs displayed consistently high shared/fixed ratios in both lineages, consistent with their non-autonomous nature, while Helitrons were more abundant and often population-specific in Oriental pears (Supplementary Fig. 60), suggesting lineage-specific accumulation. Differences among TE orders further reflect their biology: LARDs largely represent ancient relics, SINEs include a mixture of old and more recent insertions[5,6], and Helitrons appear more abundant and population-specific in Oriental pears, suggesting lineage-specific retention or recent activity.

We identified 1,170 TIPs within 2 kb upstream of genes under positive selection in Occidental pears and 1,233in Oriental pears (Supplementary Fig. 60c,d; Supplementary Tables 21,22). Across populations, 0.25–0.48% of total TIPs were located within gene bodies or upstream regions (Supplementary Table 23), which describes where TIPs lie relative to genes; because TIPs were not enriched near selected genes (Supplementary Note 4), it is not evidence for a contribution of TEs to regulatory variation at domestication-related loci.

Focusing on immunity genes under positive selection (including *NBS-LRR*[7,8] and *EF-TU RECEPTOR* loci[9]), we observed variation in the proportion of shared upstream TE insertions among cultivated populations. In European pears, perry *P. communis* showed a lower proportion of immunity genes associated with shared upstream insertions compared to dessert cultivars; however, these differences were not statistically significant (Chi-square and pairwise Fisher's exact tests, $p > 0.05$) (Supplementary Fig. 60c; Supplementary Table 21). Although this trend may suggest subtle population-level differences, the lack of statistical support prevents firm conclusions about population-specific purging or divergent selective regimes at these loci.

**Reference**

1. Cingolani, P. *et al.* A program for annotating and predicting the effects of single nucleotide polymorphisms, SnpEff: SNPs in the genome of Drosophila melanogaster strain w1118; iso-2; iso-3. *Fly* **6**, 80–92 (2012).
2. McLaren, W. *et al.* The Ensembl Variant Effect Predictor. *Genome Biol* **17**, 122 (2016).
3. Renaud, G., Hanghøj, K., Korneliussen, T. S., Willerslev, E. & Orlando, L. Joint Estimates of Heterozygosity and Runs of Homozygosity for Modern and Ancient Samples. *Genetics* **212**, 587–614 (2019).
4. Wang, Z. *et al.* Effective Purging and Conservation of Heterozygous Regions During Independent Evolution to High-Level Inbreeding in Wild Paradise Fishes. *Molecular Ecology* **34**, e70045 (2025).
5. Kalendar, R. *et al.* Large retrotransposon derivatives: abundant, conserved but nonautonomous retroelements of barley and related genomes. *Genetics* **166**, 1437–1450 (2004).
6. Weiner, A. M. SINEs and LINEs: the art of biting the hand that feeds you. *Current Opinion in Cell Biology* **14**, 343–350 (2002).
7. Jia, Y., Yuan, Y., Zhang, Y., Yang, S. & Zhang, X. Extreme expansion of NBS-encoding genes in Rosaceae. *BMC Genetics* **16**, 48 (2015).
8. Lv, L. *et al.* A TIR-NBS-LRR Gene MdTNL1 Regulates Resistance to Glomerella Leaf Spot in Apple. *International Journal of Molecular Sciences* **23**, 6323 (2022).
9. Vuong, U. T. *et al.* Engineering plant immune circuit: walking to the bright future with a novel toolbox. *Plant Biotechnology Journal* **21**, 17–45 (2023).
