## Supplementary Note 3 for "Contrasting genomic routes to domestication in Occidental and Oriental pears"

### Material and methods

#### Protein embedding generation and clustering

Protein sequences were taken from the two reference genomes used for read mapping (main text Methods): *P. communis* 'Bartlett' DH v2.0 (hereafter PCOM) and *P. pyrifolia* 'Cuiguan' v1.0 (hereafter PPY). The FASTA headers of both proteomes were normalized to the format 'species|identifier', and then the two files were concatenated into a single multi-species FASTA (80,004 sequences) used for the subsequent analysis.

Vector representations of each protein sequence were calculated using the protein language model ESM2[1] ('esm2_t33_650M_UR50D', 650M parameters, 33 layers), via PyTorch 2.5.1[2]. For each protein, the final embedding corresponds to the average of the per-residue representations of the last layer (layer 33). Sequences longer than 3,000 amino acids were truncated to this length.

The embeddings (80,004 × 1,280) were z-score standardized, then reduced by PCA (scikit-learn 1.8.0[3]) to 50 components. Unsupervised clustering was performed by HDBSCAN[4] on the PCA space, providing for each protein a cluster identifier (or the "noise" label) and a probability of belonging to the assigned cluster.

The distribution of cluster sizes and the proportion of proteins classified as "noise" were examined for each population, without filtering on the probability of membership.

#### Gene cluster enrichment test under selection

For each population, the enrichment of each HDBSCAN cluster in genes from the corresponding selection list was tested using a one-sided hypergeometric test (overrepresentation). The statistical universe for each test is the set of clustered proteins (excluding noise), i.e., 57,248 proteins.

For a given cluster and selection list, the test is defined by four quantities: *N*: the number of proteins in the universe; *K*: the number of genes from the selection list that are present in the universe (the total number of "successes"); *n*: the number of proteins in the cluster under consideration (cluster size); *k*: the number of genes from the selection list that belong to that cluster.

Under the null hypothesis, a cluster's members are a random sample of *n* proteins drawn without replacement from the universe. The number *X* of selected genes in the cluster then follows a hypergeometric distribution, $X \sim$ Hypergeometric(*N, K, n*). The one-sided *p*-value is the probability of observing at least *k* selected genes in the cluster:

$$P(X \geq k) = \sum_{i=k}^{min(n.K)} \frac{C(K,i)\, C(N-K, n-i)}{C(N,n)}$$

where C(a, b) denotes the binomial coefficient. Clusters with no selected gene ($k = 0$) were given $p = 1$.

For each cluster, we also report the fold enrichment, FE = $(k/n)/(K/N)$. FE is the ratio of the proportion of selected genes in the cluster to their proportion in the universe.

*P*-values were corrected for multiple testing with the Benjamini–Hochberg procedure (statsmodels 0.14.6) across all clusters tested within each population. A cluster was considered significantly enriched when FDR < 0.05. Because *K* is specific to each selection list, the test accounts for differences in list size between populations. As a consequence, enrichment of a given cluster cannot be compared between lists of very different sizes, including the pooled lists described below, using the number of selected genes alone.

To assess whether certain clusters are under selection recurrently across several populations rather than in a population-specific manner, the 12 p-values obtained per cluster (one per population) were combined using Fisher's method (scipy.stats.combine_pvalues, method="fisher"), followed by an FDR (Benjamini-Hochberg) correction on all the tested clusters.

To determine whether enrichment signals detected in single populations reflect a broader, shared response, cluster-level p-values were combined pairwise across all population pairs (Fisher's method, one pair at a time, each combination independently BH-FDR-corrected across the 3,294 tested clusters), rather than combining all populations at once as in the global convergence test. This was performed on both the complete and the TIP-filtered selection lists (66 pairs, all selection-scan genes, and lists of genes with a nearby TIP within 2 kb).

For clusters identified as significant in at least one population (enrichment test, see Results), the eggNOG functional tags (preferred name, description, COG category, Pfam domains) of member proteins were aggregated by cluster (cluster_function_network.py) to identify the dominant function(s) of each cluster.

### Results

#### Clustering quality check

HDBSCAN clustering of the ESM2 embedding space of the 80,004 proteins from the two reference proteomes produced 3,294 clusters. 57,248 proteins (71.6%) were assigned to a cluster, with the remaining 22,756 (28.4%) classified as noise. The vast majority of clusters (3,188/3,294, or 96.8%) group proteins from both PCOM and PPY, suggesting that the clustering preferentially captures groups of homologs shared between PCOM and PPY rather than artifacts specific to one assembly.

#### Enrichment tests

Of the 12 enrichment tests performed (one per population, 3,294 clusters tested per population), 9 clusters showed significant gene enrichment under positive selection on the unfiltered gene lists, and a tenth (cluster 428) only on the TIP-filtered lists (Table N1, below). The observed enrichment factors are high, 19 to 57 times the expected value, corresponding to a limited number of genes under selection (4 to 8 proteins) within clusters of 16 to 44 proteins.

**Table N1. Clusters significantly enriched in genes under positive selection (FDR < 0.05).** Unless stated otherwise, tests used all selected genes (unfiltered); cluster 428 was significant only for genes with a TIP within 2 kb. Population abbreviations: ussu, *P. ussuriensis*; betu, *P. betulifolia*; cauc, *P. caucasica*; White, *P. pyrifolia* white CN; Sand_CN-SE and Sand_CN-SW, *P. pyrifolia* sand CN-SE and CN-SW

| **Cluster** | **Cluster size** | **Population** | **Proteins under selection** | **Fold enrichment** | **FDR** |
|---|---|---|---|---|---|
| 1008 | 44 | Sand_CN-SE | 8 | 28.0x | 1.40e-06 |
| 762 | 22 | Sand_CN-SE | 4 | 28.0x | 1.28e-02 |
| 968 | 21 | Sand_CN-SE | 4 | 29.3x | 1.28e-02 |
| 1289 | 16 | ussu | 4 | 56.6x | 2.10e-03 |
| 976 | 32 | ussu | 4 | 28.3x | 2.00e-02 |
| 344 | 30 | betu | 5 | 27.6x | 3.20e-03 |
| 311 | 39 | cauc | 5 | 18.9x | 2.21e-02 |
| 439 | 35 | Sand_CN-SW | 5 | 19.8x | 1.73e-02 |
| 900 | 19 | White | 4 | 26.9x | 4.27e-02 |
| 428 | 39 | betu | 4 | 25.4x | 3.13e-02 |

The convergence test (combination of *p*-values using Fisher's method across the 12 populations, FDR correction on all clusters) identified no significant clusters (FDR < 0.05) after correction for multiple tests. This result is consistent with population-specific selection, although the test has limited power to detect weaker shared signals.

In the pairwise tests, 41 (cluster, population-pair) combinations reached significance across 32/66 pairs with the complete selection lists; under the TIP-filtered set, 40 combinations reached significance across 23/66 pairs.

The majority of these hits were attributable to a single population with a strong individual signal recurring across every pair it was tested in: cluster 1008 co-occurred exclusively with Sand_CN-SE (11/11 of its significant pairs), cluster 1289 exclusively with ussu (11/11), and cluster 344 exclusively with betu

(11/11), consistent with these three clusters' already-significant single-population enrichment (Table N1) rather than evidence of independent convergence between two populations.

In contrast, two clusters reached pairwise significance without being individually significant in either constituent population: cluster 107 (Sand_CN-SE: $p = 1.23 \times 10^{-2}$, FDR = 1; ussu: $p = 2.06 \times 10^{-4}$, FDR = 0.226; combined FDR < 0.05), and cluster 474 (Sand_CN-SW: $p = 4.53 \times 10^{-4}$, FDR = 0.747; ussu: $p = 3.86 \times 10^{-3}$, FDR = 1; combined FDR < 0.05). These may represent selection signals too weak to survive genome-wide correction within any single population, but that reinforce into significance when evidence from two populations is combined without dilution from unrelated, uninformative populations.

Functional eggNOG annotation was recovered for all 12 significant clusters identified across the enrichment tests: 10 significant in at least one single-population test (Table N1) and 2 additional clusters (107, 474) significant only in the pairwise tests(Table N2).

Annotations were assigned to 308 of the 344 constituent proteins (89.5%), and every cluster contained at least one annotated member. Several clusters correspond to gene families with well-established roles in plant defense and abiotic stress tolerance: disease-resistance genes of the NB-LRR family (cluster 1008; 32/44 members carry paired NB-ARC and LRR_8 domains, and all 10 selection-scan members across five populations are NB-LRR), a TIR-NBS-LRR disease-resistance family (cluster 474; 19/21 members carry TIR together with NB-ARC, and all six selection-scan members are TIR-type, four with explicit TIR/NB-ARC Pfam assignments and two annotated as Toll/interleukin-1 resistance proteins without Pfam support), LEA proteins (Late Embryogenesis Abundant, stress/desiccation tolerance; cluster 900, 17/19 members LEA_2), HSP70 heat shock proteins (cluster 762, 22/22 members), potassium transporters of the HAK/KUP family (cluster 439, 32/35 members carrying K_trans, with HAK11, HAK5, HAK15 and HAK17 assignments), monothiol glutaredoxins (oxidative-stress regulation; cluster 1289, 15/16 members), and 2-alkenal reductases (detoxification; cluster 311, 35/39 members).

For the clusters whose eggNOG label alone was uninformative, cross-referencing the selection-scan members against their domain architecture resolved a more specific function than the generic family assignment would suggest. Cluster 107, one of the two clusters recovered only by the pairwise tests, is composed almost entirely of receptor-like proteins (23/26 members sharing an identical LRRNT_2/LRR_1/LRR_6/LRR_8 architecture); all five selection-scan members, spanning six populations and both reference proteomes, including the Occidental population *P. pyraster*, carry this same architecture and lack a kinase domain, identifying them as cell-surface LRR receptor-like proteins rather than the intracellular NLR receptors of clusters 1008 and 474. Cluster 428, which reached significance only when the analysis was restricted to genes carrying a transposable-element insertion polymorphism within 2 kb (betu, fold enrichment 25.4, FDR = 0.031; FDR = 0.15 on the unfiltered gene set), is dominated by SABATH-type S-adenosylmethionine-dependent carboxyl methyltransferases (29/39 members carrying Methyltransf_7, including three IAMT1 assignments), with two terpene synthases also present; three of its five selection-scan members are SABATH methyltransferases and one is a (−)-α-pinene synthase, suggesting a role in volatile biosynthesis rather than general methylation. Three clusters remained functionally unresolved: cluster 344 (28/30 members annotated as MATH domain and coiled-coil proteins), cluster 976 (15/32 members carrying DUF674, the remainder unannotated) and cluster 968 (9/21 members carrying an X8 carbohydrate-binding domain, with four of its five selection-scan members unannotated).

**Table N2. Dominant function of significant clusters.** COG category is the eggNOG COG category aggregated across cluster members and may not match the dominant function.

| Cluster | Size | PCOM | PPY | COG category | Dominant function |
|---|---|---|---|---|---|
| 1008 | 44 | 23 | 21 | T — Signal transduction | Disease resistance, NB-LRR family (39) |
| 311 | 39 | 17 | 22 | Q — Secondary metabolites | 2-alkenal reductase NADP-dependent (35) |
| 428 | 39 | 23 | 16 | L — Replication, recombination and repair | Methyltransferases SAM-dependent (21) ; IAMT1 (3); SAM carboxyl methyltransferase (3) |

| | | | | | |
|---|---|---|---|---|---|
| 439 | 35 | 15 | 20 | P — Inorganic ion transport | Potassium transporter HAK (32) |
| 976 | 32 | 15 | 17 | K — Transcription | DUF674 (8); not characterized (7) |
| 344 | 30 | 12 | 18 | O — Chaperones, proteolysis | MATH domain and coiled-coil (28) |
| 107 | 26 | 9 | 17 | T — Signal transduction | Receptor-like protein 2 (23); LRR receptor-like kinases (3) |
| 762 | 22 | 9 | 13 | O — Chaperones, proteolysis | Heat shock protein HSP70 (22) |
| 474 | 21 | 2 | 19 | T — Signal transduction | Toll/interleukin-1 resistance, TIR (9); TMV resistance N-like (8); resistance protein (4) |
| 968 | 21 | 4 | 17 | S — Function unknown | Glucan endo-1,3-β-glucosidase (5) ; carbohydrate binding (4) |
| 900 | 19 | 2 | 17 | S — Function unknown | Late embryogenesis abundant, LEA (17) |
| 1289 | 16 | 6 | 10 | J — Translation | Glutaredoxins monothiol (15) |

To test whether the selection signal detected within individual populations is shared at a broader biogeographic or domestication scale, gene lists were pooled within two alternative groupings and re-tested for cluster enrichment using the same one-sided hypergeometric procedure (BH-FDR < 0.05 within each grouping). First, populations were pooled by geographic lineage: Occidental (*P. caucasica, P. communis* Dessert, *P. communis* Perry, *P. pyraster*; 2,148 unique genes, 1,486/2,148 [69.2%] resolved in the clustering universe) versus Oriental (*P. betulifolia*, *P. pashia, P. pyrifolia* Japanese, *P. pyrifolia* Sand_CN-other, *P. pyrifolia* Sand_CN-SE, *P. pyrifolia* Sand_CN-SW, *P. ussuriensis, P. pyrifolia* White; 3,492 unique genes,

2,539/3,492 [72.7%] resolved). Second, populations were pooled by domestication status, excluding *P. betulifolia* (rootstock, neither clearly wild nor cultivated): cultivated (*P. communis* Dessert, *P. communis* Perry, *P. pyrifolia* Japanese, *P. pyrifolia* Sand_CN-ohter, *P. pyrifolia* Sand_CN-SE, *P. pyrifolia* Sand_CN-SW, *P. pyrifolia* White; 3,693 unique genes, 2,653/3,693 [71.8%] resolved) versus wild (*P. caucasica, P. pyraster, P. pashia, P. ussuriensis*; 1,921 unique genes, 1,350/1,921 [70.3%] resolved). Both groupings were applied twice: once to the complete per-population selection gene lists, and once to the TIP-filtered lists (Occidental 1,205 genes, 836 [69.4%] resolved; Oriental 2,194, 1,611 [73.4%]; cultivated 2,221, 1,604 [72.2%]; wild 1,136, 810 [71.3%]).

On the complete selection gene lists, neither grouping yielded any significantly enriched cluster (BH-FDR < 0.05) in either group, nor did a Fisher's method convergence test between the two groups within each scheme. The strongest, sub-threshold signals corresponded largely to clusters already identified in single-population tests: cluster 439 (HAK/KUP potassium transporters; Oriental: 8/35 selected, 5.2-fold, FDR = 0.38), cluster 474 (TIR-NBS-LRR; Oriental: 6/21, 6.4-fold, FDR = 0.38), cluster 1008 (NB-LRR; Oriental: 8/44, 4.1-fold, FDR = 0.54; cultivated: 10/44, 4.9-fold, FDR = 0.086), cluster 900 (LEA; Oriental: 5/19, 5.9-fold, FDR = 0.65), and cluster 1289 (monothiol glutaredoxins; wild: 4/16, 10.6-fold, FDR = 0.37), but none was significant after pooling, because pooling increased the number of selected genes ($K$) more than the overlap within each cluster. The Occidental lineage was the exception: none of the clusters significant in its four constituent populations ranked among its strongest pooled signals, cluster 311 (2-alkenal reductases) retaining its five overlapping genes but only 4.9-fold enrichment at FDR = 1.00.

Restricting the same analysis to genes carrying a TIP within 2 kb changed this picture for the domestication grouping alone. The Occidental/Oriental scheme again yielded no significant cluster and no convergence, its strongest signal being cluster 474 in the Oriental group (6/21, 10.2-fold, FDR = 0.061). In the cultivated/wild scheme, however, **cluster 1008 reached significance in the cultivated group (9/44 selected, 7.3-fold, FDR = 0.010)** and was also the single cluster recovered by the Fisher's-method test between the two groups (combined FDR = 0.020). Together with the absence of a convergent signal across all 12 populations individually, these results are consistent with the clusters enriched for positively selected genes in this dataset largely reflecting population-specific selection pressures rather than signals shared at the level of geographic lineage or domestication status, with the NB-LRR cluster 1008 as the one exception, whose signal survives pooling by domestication status once the gene set is restricted to selected genes with a nearby TIP.

### References


1. Lin, Z. *et al.* Evolutionary-scale prediction of atomic-level protein structure with a language model. *Science* **379**, 1123–1130 (2023).
2. Paszke, A. *et al.* PyTorch: An Imperative Style, High-Performance Deep Learning Library. Preprint at https://doi.org/10.48550/arXiv.1912.01703 (2019).
3. Pedregosa, F. *et al.* Scikit-learn: Machine Learning in Python. *J. Mach. Learn. Res.* **12**, 2825–2830 (2011).
4. McInnes, L., Healy, J. & Astels, S. hdbscan: Hierarchical density based clustering. *J. Open Source Softw.* **2**, 205 (2017).
