## Supplementary Note 4 for "Contrasting genomic routes to domestication in Occidental and Oriental pears"

### Methods

#### Association between TIP occurrence, population differences, and positive selection

We investigated TIP occurrence across Occidental (PCOM) and Oriental (PPY) pear populations using two complementary approaches that address distinct biological questions.

First, we investigated whether TIP occurrence upstream of positively selected genes differed between populations, focusing on insertions potentially affecting cis-regulatory elements and gene expression. We restricted this analysis to strand-specific 2-kb upstream regions, considering all positively selected genes and a subset associated with immunity based on *Arabidopsis thaliana* functional annotations, gene symbols, and Gene Ontology terms. For each population, only genes identified as being under positive selection in that population were considered. TIP occurrence was scored as the presence or absence of at least one insertion, including both population-specific and shared TIPs. Population differences were assessed using generalized estimating equations (GEE)[1] with a binomial distribution, logit link, and exchangeable correlation structure, accounting for repeated observations of the same gene across populations. Models were adjusted for local recombination rate, TE density, gene density, and log-transformed gene length, with covariates standardized to Z-scores. Pairwise comparisons were expressed as adjusted odds ratios (ORs) with 95% confidence intervals. Statistical significance was assessed using two-sided Wald tests with robust standard errors, followed by Benjamini–Hochberg false discovery rate (FDR) correction applied separately for each lineage and gene category.

Second, we tested whether TIP occurrence was associated with positive selection within each population by comparing positively selected and non-selected genes. Unlike the first approach, this analysis assessed TIP enrichment or depletion among positively selected genes relative to the genomic background. We assessed TIP presence in coding sequences (CDS), introns, and strand-specific 2-kb upstream and downstream regions, as well as 5′ and 3′ untranslated regions (UTRs) in PPY only, because PCOM lacks UTR annotations. Associations were tested using Cochran–Mantel–Haenszel (CMH) tests[2] stratified by protein language model (PLM) gene cluster[3,4] and complementary binomial generalized estimating equations (GEE) with a logit link, exchangeable correlation structure, and PLM gene cluster as the clustering variable. GEE models were adjusted for log-transformed gene length, local TE density, gene density, and recombination rate, with covariates standardized to Z-scores. TE and gene densities were calculated as the proportions of bases covered by annotated TEs and other genes, respectively, within 50-kb flanking regions excluding the focal gene body. Local recombination rate was calculated as the mean Rho across these regions, weighted by the number of base pairs covered by each recombination interval. Robustness was assessed using 1,000 randomly sampled sets of non-selected genes matched on quartiles of the four covariates. *P* values from CMH, GEE, and matched-control analyses were independently adjusted using the Benjamini–Hochberg procedure across population-by-region comparisons, separately for PCOM and PPY.

Definition of selected genes and limitations. Positively selected genes were those whose coding or 1-kb upstream regions overlapped $\omega$ or Tajima's *D* outlier regions in the population (main-text Methods; Supplementary Tables 8,9); all other genes were treated as non-selected. Because selection status is binary and non-selected genes include undetected targets of selection, misclassification is expected to bias associations toward the null, so these tests have limited power to detect weak effects and non-significant results should not be read as evidence of absence. Adjusted estimates from tests with no insertion among selected genes (e.g., 3′ UTR) are unstable (quasi-complete separation) and are reported in Supplementary Table 25 for completeness only.

### Results

#### TIP occurrence in relation to population history and positive selection

TIP occurrence upstream of selected genes differed between several populations of the same lineage (Supplementary Table 24). In Occidental pears, cultivated Perry differed from dessert (OR = 1.69, FDR < 0.001) but not significantly from wild *P. pyraster*. In Oriental pears, *P. pashia* exhibited higher upstream TIP occurrence than *P. betulifolia* (OR = 2.32, FDR < 0.001), suggesting distinct evolutionary trajectories despite their close phylogenetic relationship and the complex history of introgression within Oriental pears. Comparisons with non-selected genes revealed no widespread TIP enrichment, but rather significant depletion predominantly in wild populations (Supplementary Table 25): upstream in *P. caucasica* (adjusted OR = 0.67, FDR $< 0.001$) and downstream in *P. pashia* and *P. ussuriensis*, as well as cultivated White (adjusted OR = 0.60–0.70, FDR $\leq 0.026$). These associations were independently supported by matched-control analyses (FDR = 0.032) but not by the cluster-stratified CMH tests (FDR $\geq 0.23$). Three further depletions were significant only in the adjusted models and not supported by the matched controls: upstream in sand CN-other (adjusted OR = 0.71, FDR = 0.028; matched-control FDR = 0.07), in introns in Nashi JP (OR = 0.54, FDR = 0.033; matched-control FDR = 0.20) and in the 3′ UTR of sand CN-other (matched-control FDR = 0.44). The 3′ UTR estimate is degenerate (none of the 567 selected genes carried a TIP; adjusted OR $\approx 7 \times 10^{-8}$) and was not interpreted. Nominal enrichments (CDS in perry, introns in *P. betulifolia*; CMH $P < 0.02$) did not survive correction (FDR $\geq 0.10$), so no compartment showed enrichment in any population. These patterns are descriptive: they are compatible with, but do not demonstrate, an effect of introgression on TIP occurrence between populations or of purifying selection against insertions near selected genes.


1. LIANG, K.-Y. & ZEGER, S. L. Longitudinal data analysis using generalized linear models. *Biometrika* **73**, 13–22 (1986).
2. Mantel, N. & Haenszel, W. Statistical Aspects of the Analysis of Data From Retrospective Studies of Disease. *J Natl Cancer Inst* **22**, 719–748 (1959).
3. Lin, Z. *et al.* Evolutionary-scale prediction of atomic-level protein structure with a language model. *Science* **379**, 1123–1130 (2023).
4. McInnes, L., Healy, J. & Astels, S. hdbscan: Hierarchical density based clustering. *Journal of Open Source Software* **2**, 205 (2017).
