## Supplementary Fig for "Contrasting genomic routes to domestication in Occidental and Oriental pears"

### Supplementary Information

This file contains all supplementary figures with their corresponding legends and the legends for all supplementary tables.

All supplementary tables are provided separately in the file “Supplementary_Tables.xlsx”.

### Supplementary Figures

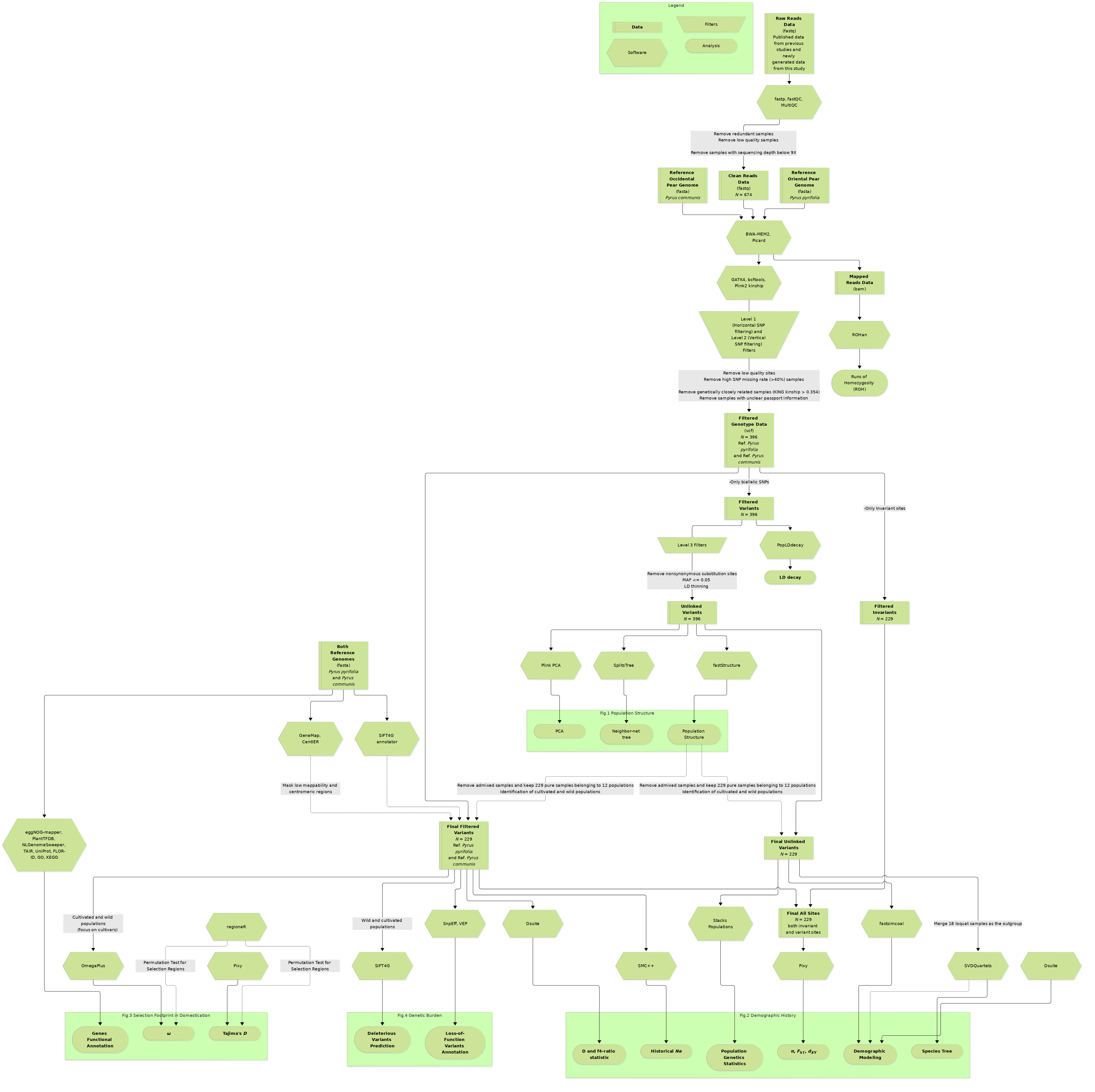

**Supplementary Figure 1 | The workflow for bioinformatic analysis used in this study (SNPs part).**

The original code for rendering this plot in Markdown syntax can be accessed in the Git repository of this study hosted on GitHub (<https://github.com/CornilleEclecticLab/pear-SNP>).

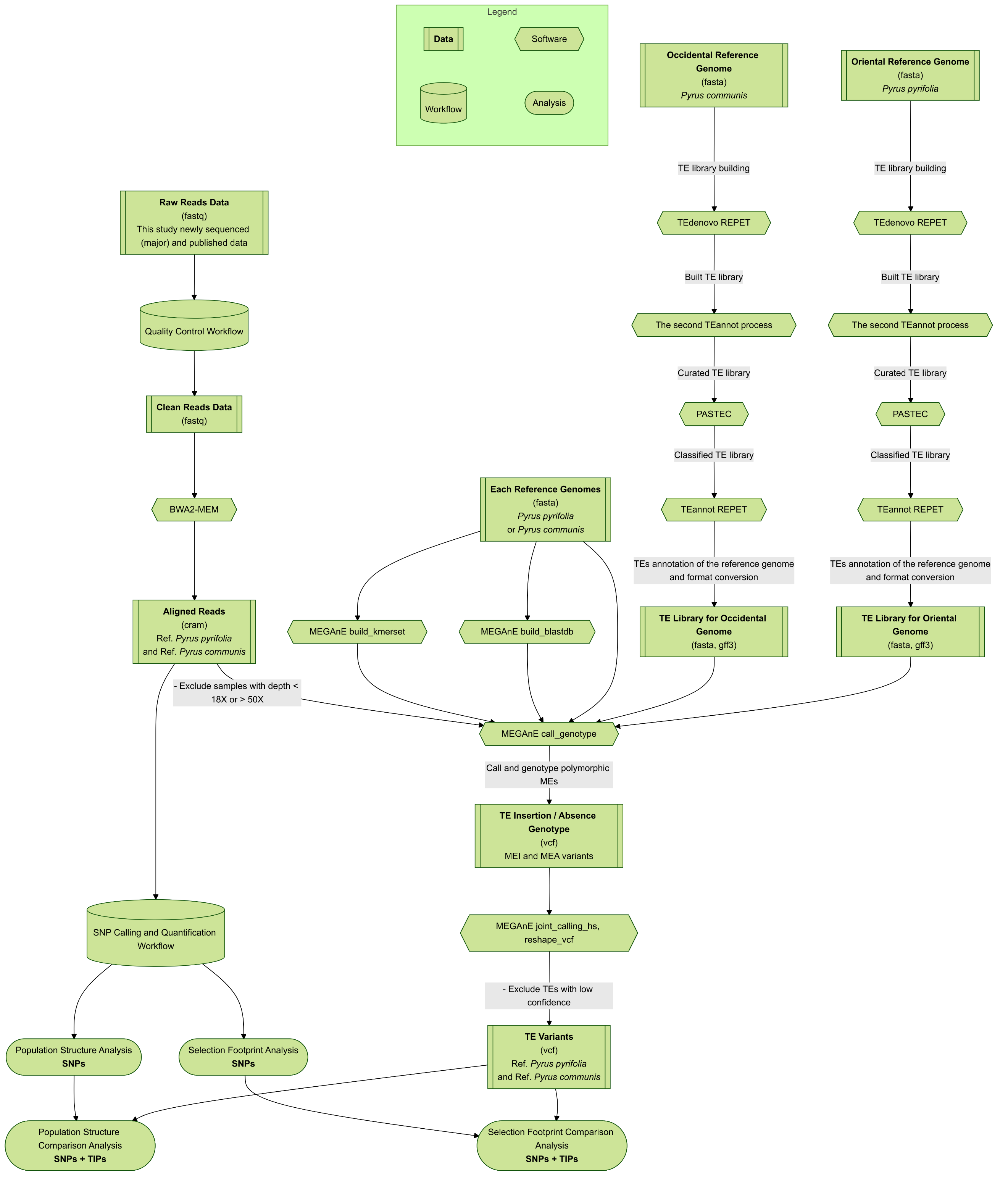

**Supplementary Figure 1 (continued) | The workflow for bioinformatic analysis used in this study (part integrating SNPs and TIPs).**

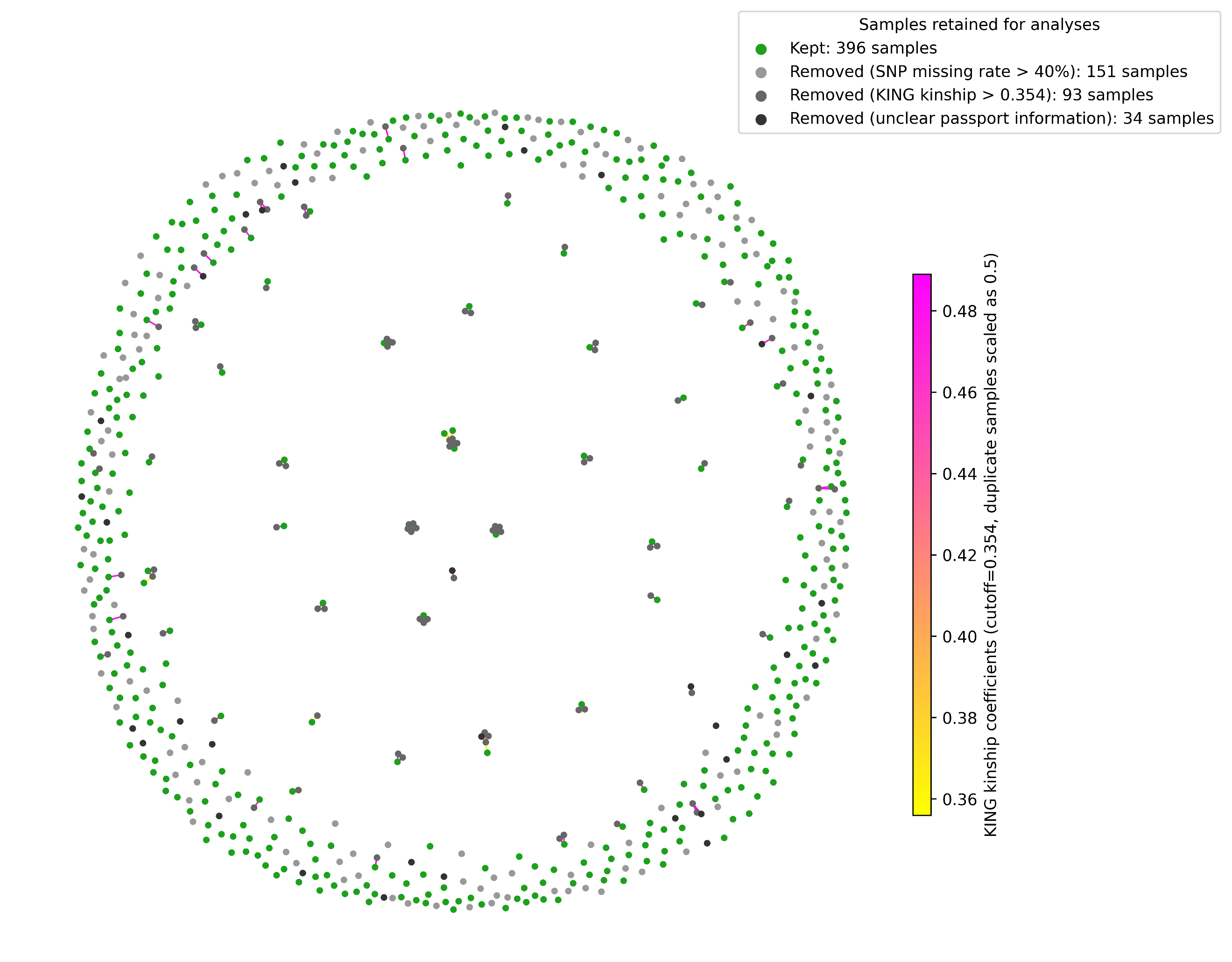

**Supplementary Figure 2 | Diagram of the 396 samples retained for analyses.**

This plot illustrates the samples that were retained and removed, along with their corresponding kinship relationships, before population genetic analysis. Each point represents the genome of one individual called from the whole-genome resequencing data, corresponding to a sample column in a VCF file. Edges and their colors connecting two points indicate a close relationship (KING kinship coefficients ≥ 0.354) between the genotypes, as estimated using the KING-robust kinship estimator implemented in PLINK2. After removing samples with a high SNP missing rate (>40%), one sample from each pair (or several samples from each network) of individuals showing a close kinship relationship (retaining a subset of relatively unrelated individuals to minimize redundancy), and those with inconsistent passport information, a dataset of 396 individuals was obtained. Details are provided in Supplementary Table 1.

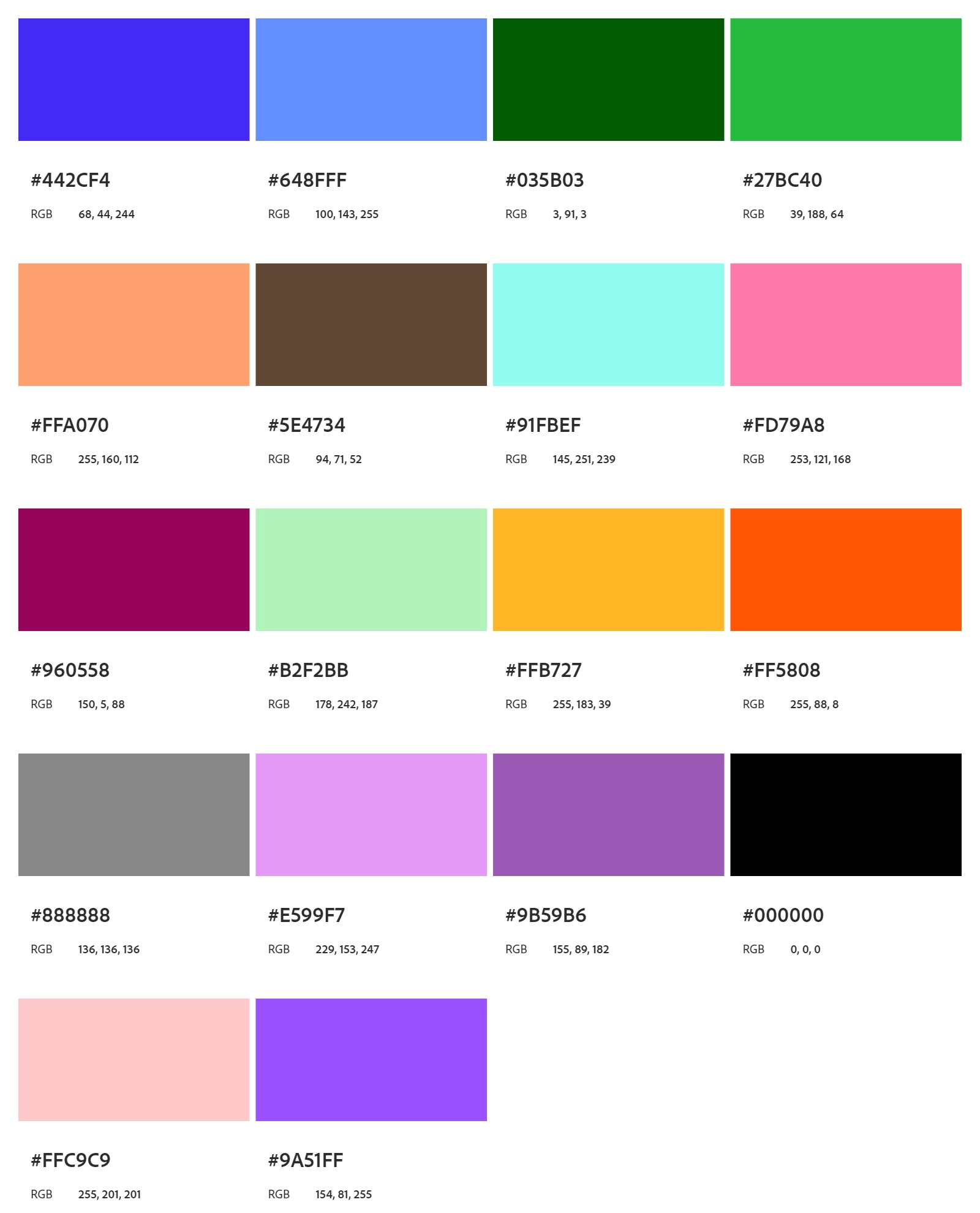

**Supplementary Figure 3 | Color codes for the color-blind-friendly palette used in this study.**

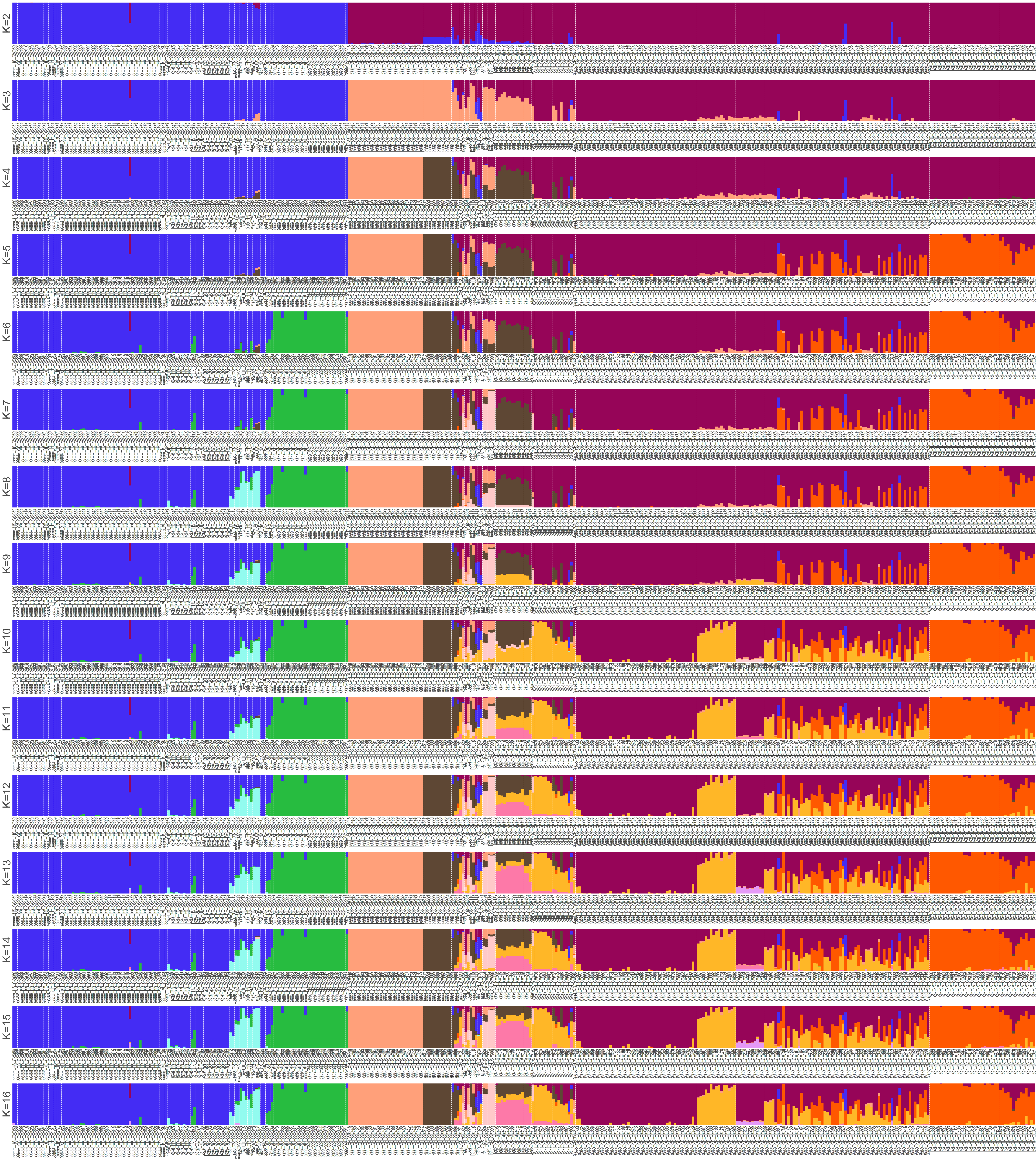

**Supplementary Figure 4 | The population structure landscape (*K*=2 to *K*=16) for the full dataset.**

This plot, based on 21,756 SNPs from the genomes of 396 individuals encompassing both Occidental and Oriental pears mapped onto *P. pyrifolia*, was generated using fastSTRUCTURE and CLUMPAK and visualized with PopHelper. The uniform IDs of the samples under each bar of this plot were coded in the format “AAAA_BB_CC000” for easy recognition during analysis. In this format, the first four letters represent the species name. The next two letters indicate the sampling country code (ISO 3166-1 alpha-2 standard), where "xx" denotes unknown locations, and the last two letters with a unique three-digit number denote the sample type: CW for cultivated Occidental, CE for cultivated Oriental, RE for rootstock Oriental, WW for wild Occidental, and WE for wild Oriental. Details of the *Q* value are shown in Supplementary Table 2.

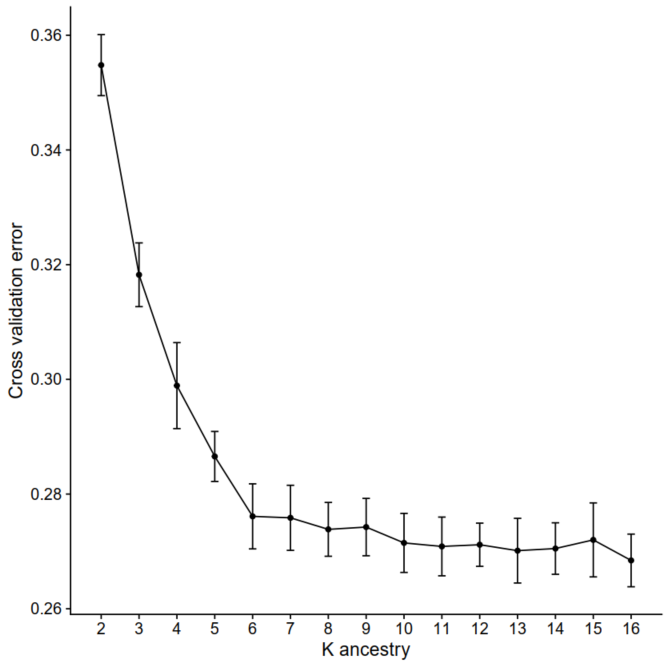

**Supplementary Figure 5 | The cross-validation error for *K* ancestries of the full dataset.**

These values were obtained from the genome data of both Occidental and Oriental pear individuals mapped onto the *P. pyrifolia* reference genome.

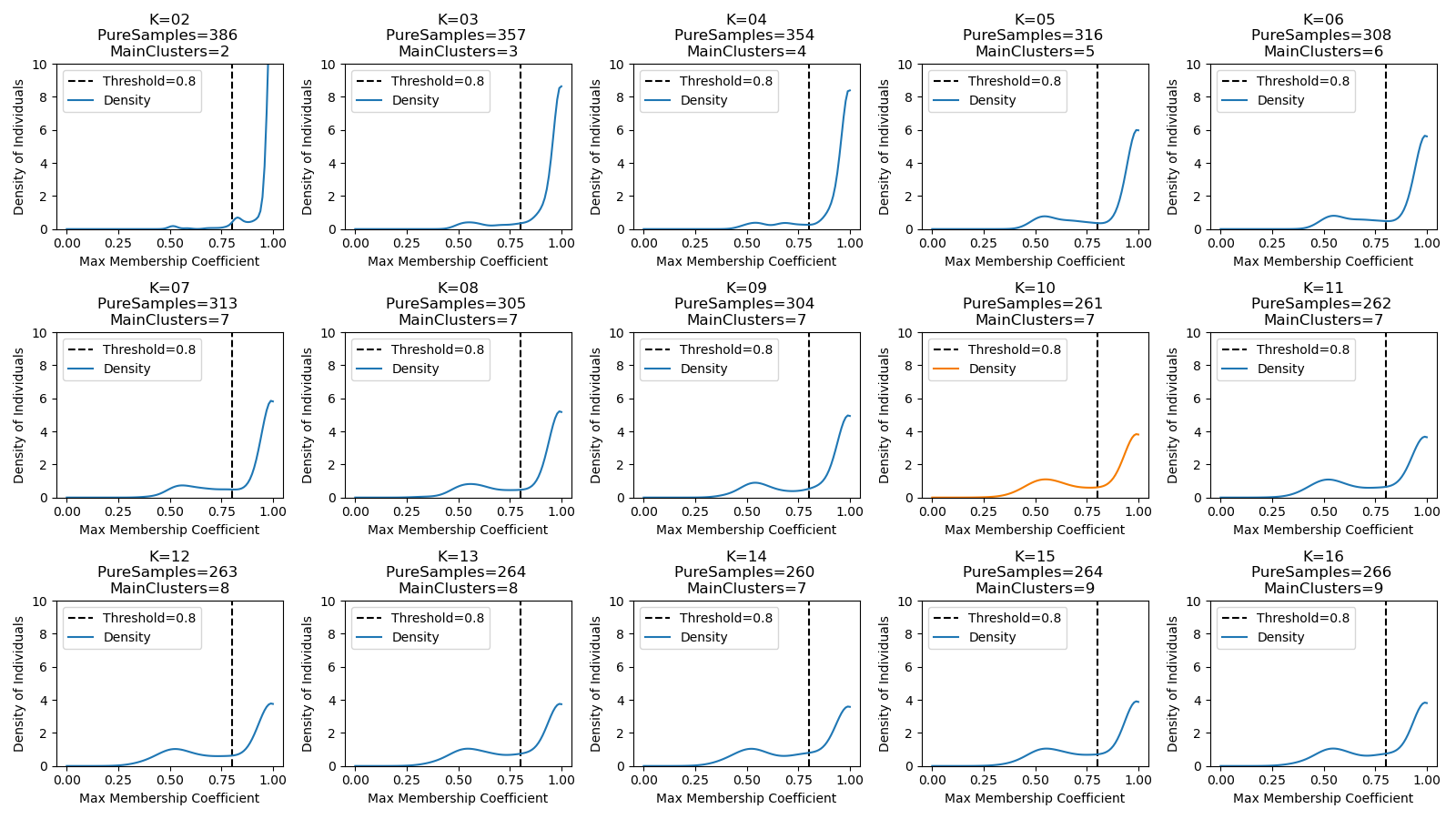

**Supplementary Figure 6 | The density distribution of genetic membership coefficient for the full dataset.**

The analysis was performed using 21,756 SNPs from the genomes of 396 pear individuals representing both Occidental and Oriental groups, mapped onto the *P. pyrifolia* reference genome. The cutoff for defining pure samples was set to 80% of the maximum membership coefficient within each genetic cluster. The “main clusters” correspond to clusters containing at least one pure sample. The optimal number of clusters was determined to be *K* = 10.

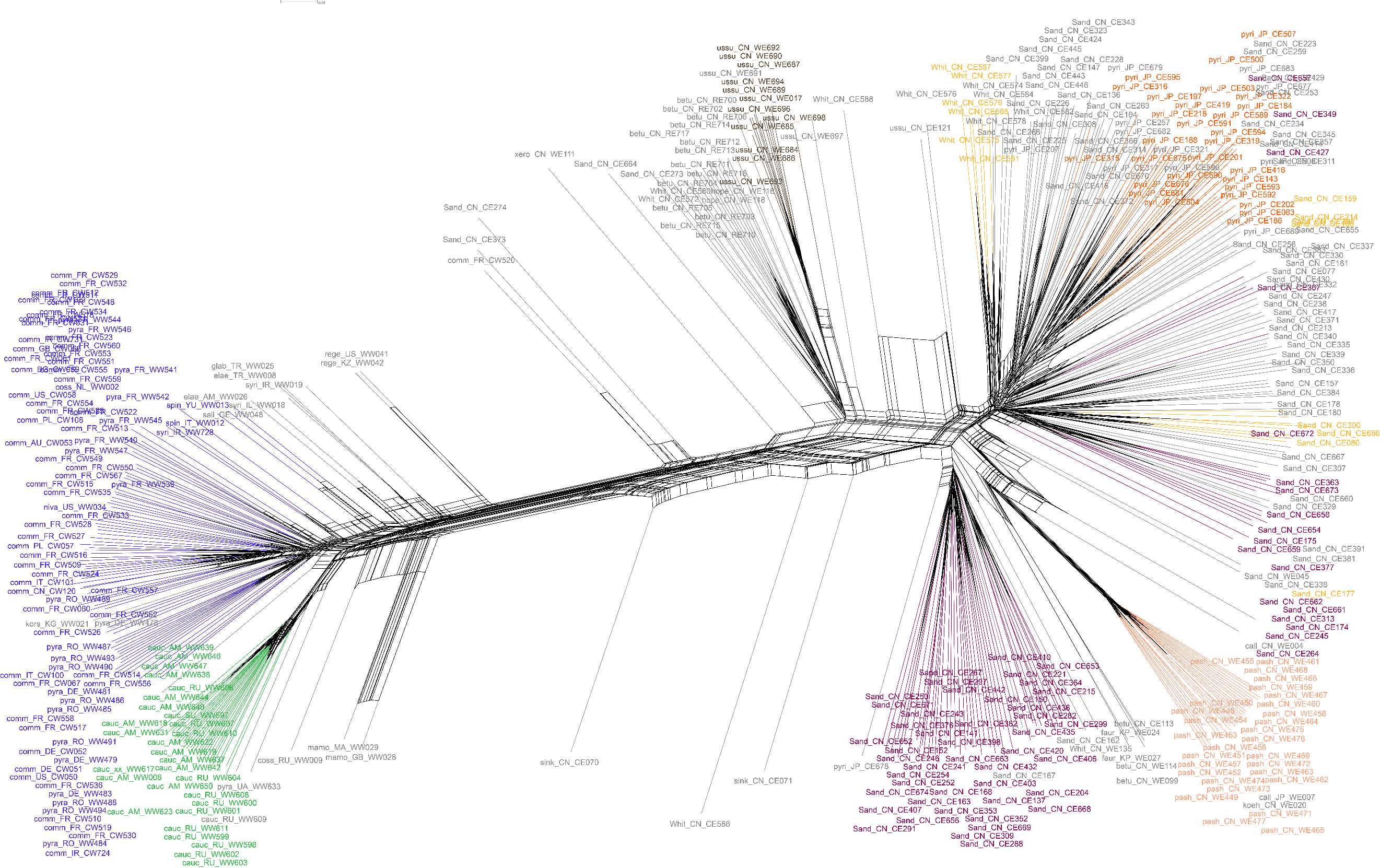

**Supplementary Figure 7 | Neighbor-net tree of the full dataset.**

The tree was reconstructed using 21,756 SNPs from the genomes of 396 individuals representing both Occidental and Oriental groups, mapped onto the *P. pyrifolia* reference genome. The distinct Occidental and Oriental pear branches are primarily concentrated on the left and right sides, respectively, as analyzed by PLINK and SplitsTree. The branches and sample ID colors correspond to their primary genetic pool colors, as assigned in the population structure plot (*K*=10), while admixed samples (with a cutoff of 80% membership) are shown in gray.

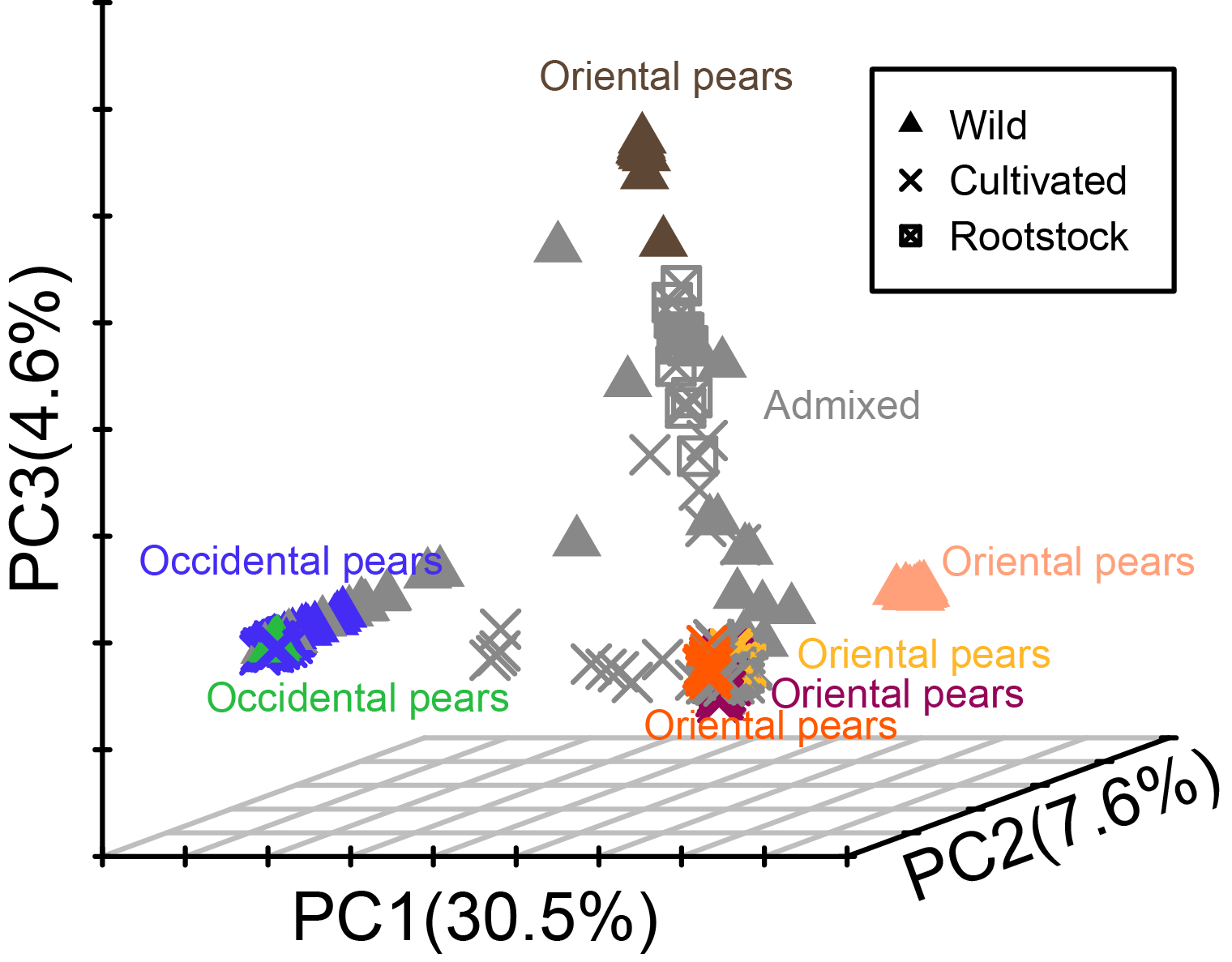

**Supplementary Figure 8 | Principal component analysis (PCA) of the full dataset.**

The analysis was performed using 21,756 SNPs from the genomes of 396 pear individuals mapped onto the *P. pyrifolia* reference genome, including both Occidental and Oriental pears. Each point represents one individual, with colors indicating the primary genetic clusters defined in the population structure analysis (*K* = 10). Admixed samples (membership < 80%) are shown in gray.

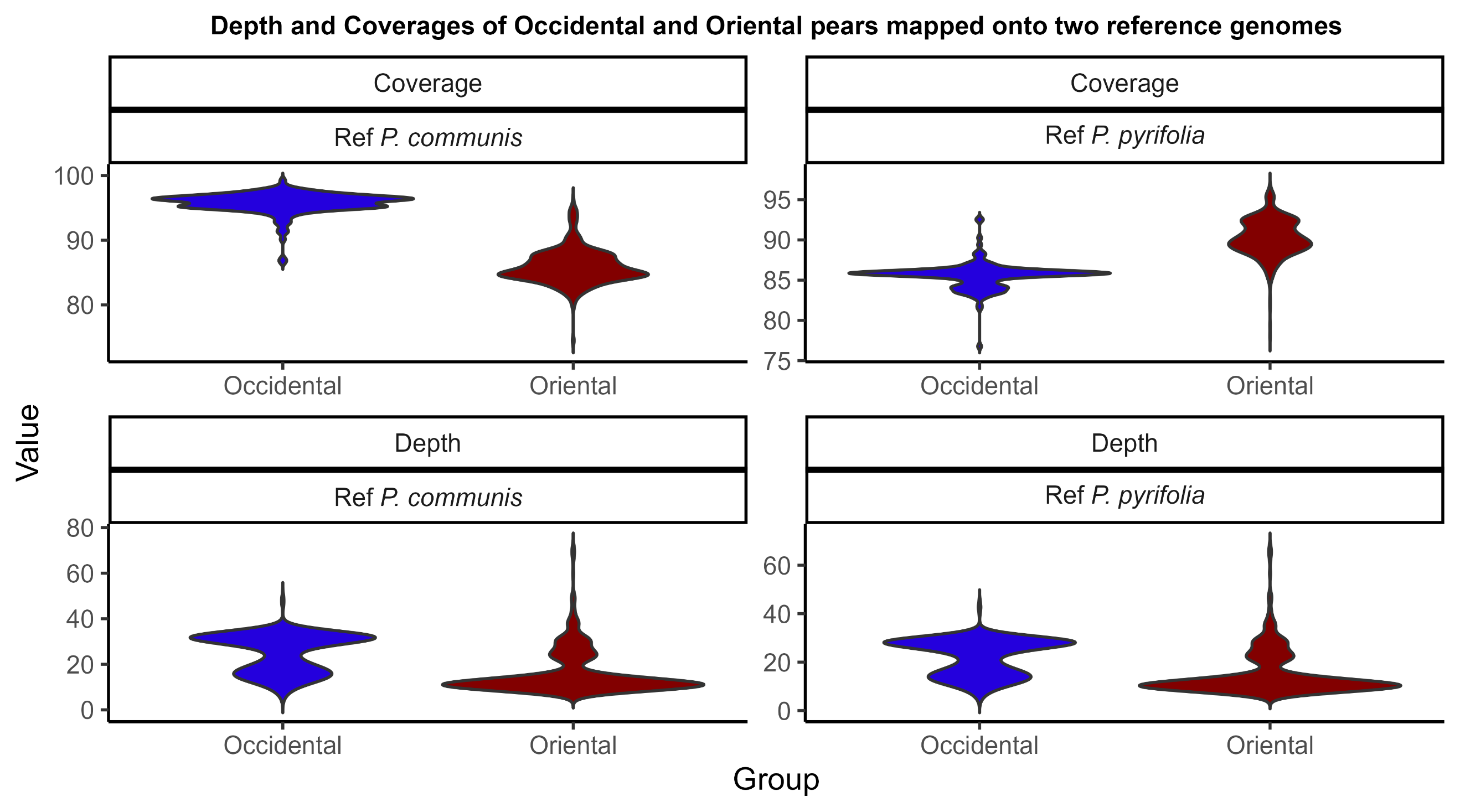

**Supplementary Figure 9 | Sequencing depth and coverage of Occidental and Oriental pears mapped onto the respective *Pyrus communis* or *Pyrus pyrifolia* reference genome.**

Coverage represents the percentage of the reference genome covered by sequencing reads, while depth indicates the average number of reads mapped per site in the reference genome. Source data for this plot is provided in Supplementary Table 1.

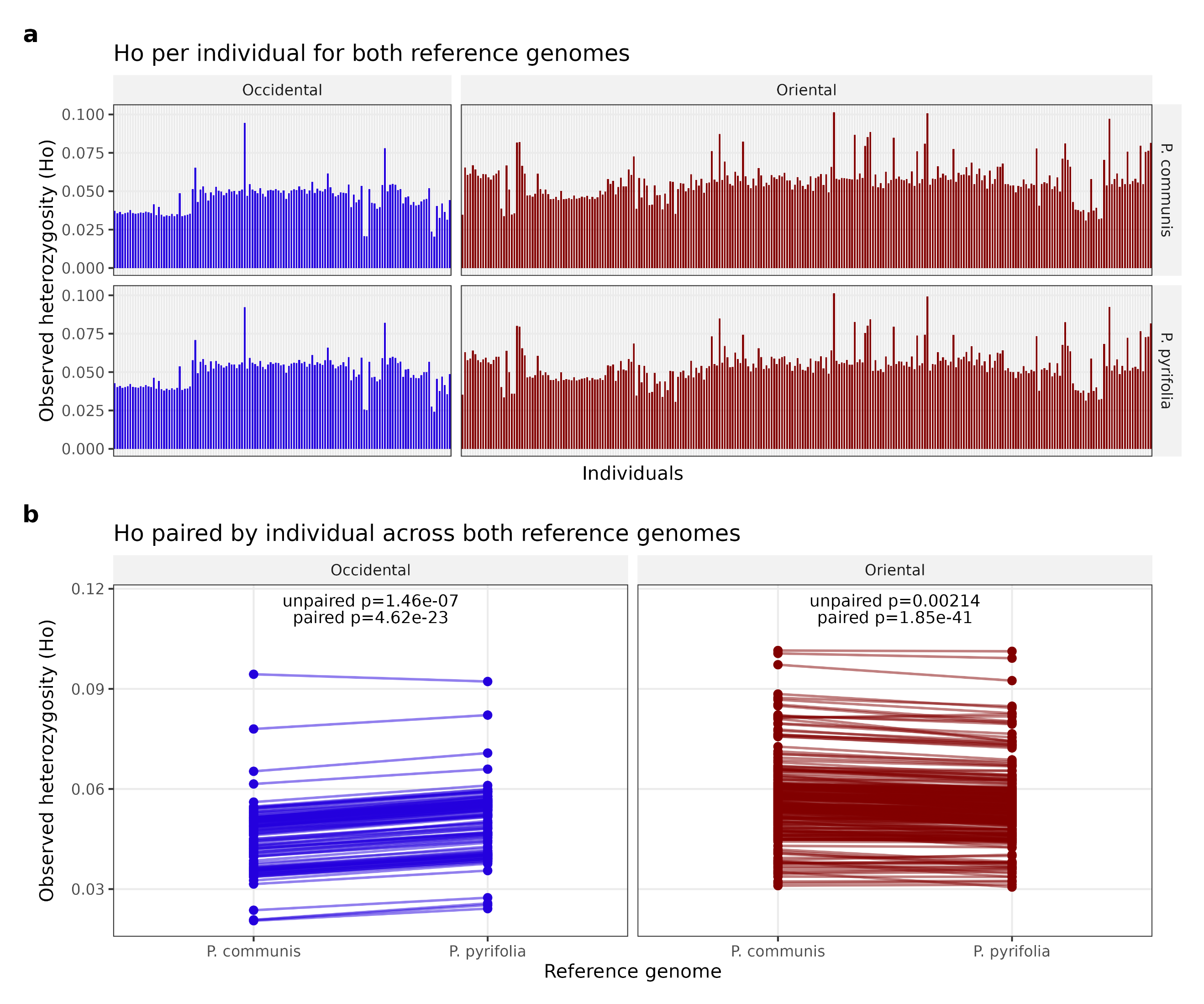

**Supplementary Figure 10 | Observed heterozygosity (*H*_o_) in the genomes of Occidental and Oriental pear individuals mapped onto each reference genome.**

(a) *H*_o_ per individual. (b) Paired *H*_o_ by individual*. P-*values were calculated using the Wilcoxon test (paired and unpaired).

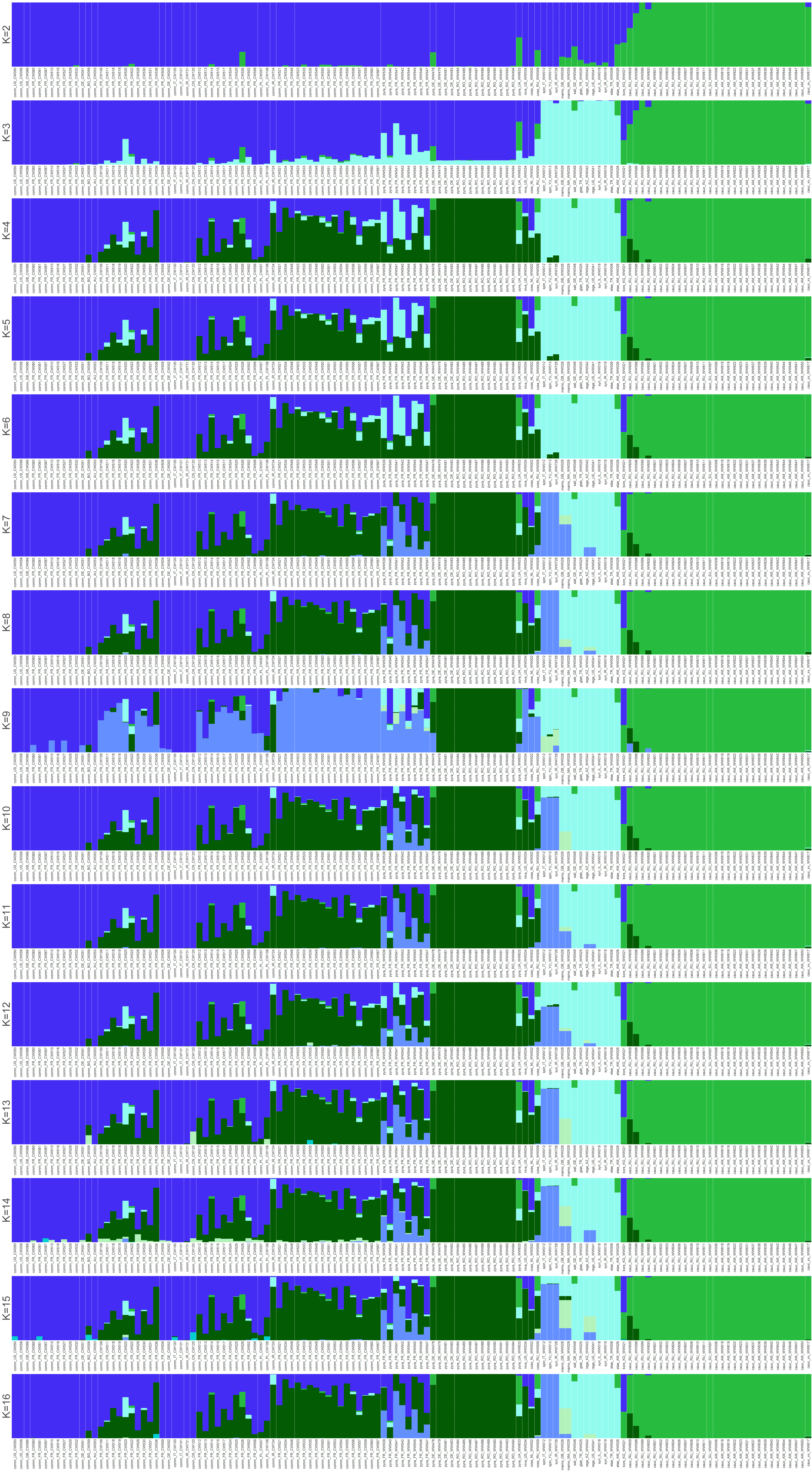

**Supplementary Figure 11 | Population structure landscape (*K* = 2 to 16) of Occidental pears mapped onto the *Pyrus pyrifolia* reference genome.**

The figure was generated using custom pipelines combining fastSTRUCTURE, CLUMPAK, and PopHelper. The Q values for each cluster at the optimal *K* = 9 are provided in Supplementary Table 2.

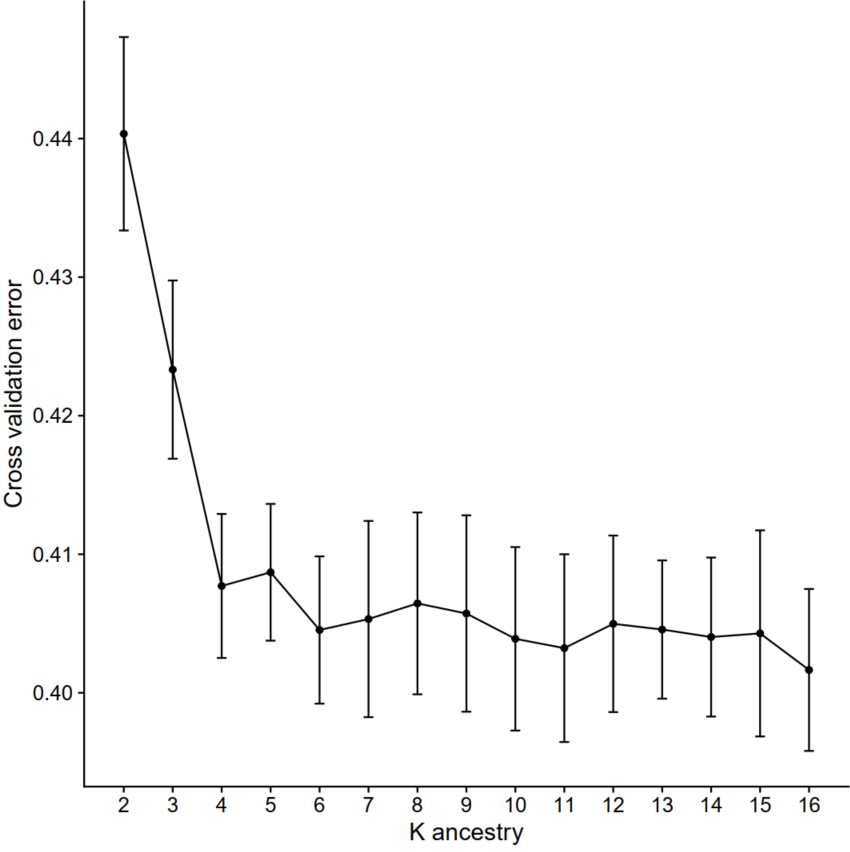

**Supplementary Figure 12 | Cross-validation error for *K* ancestries in Occidental pears mapped onto the *Pyrus pyrifolia* reference genome*.***

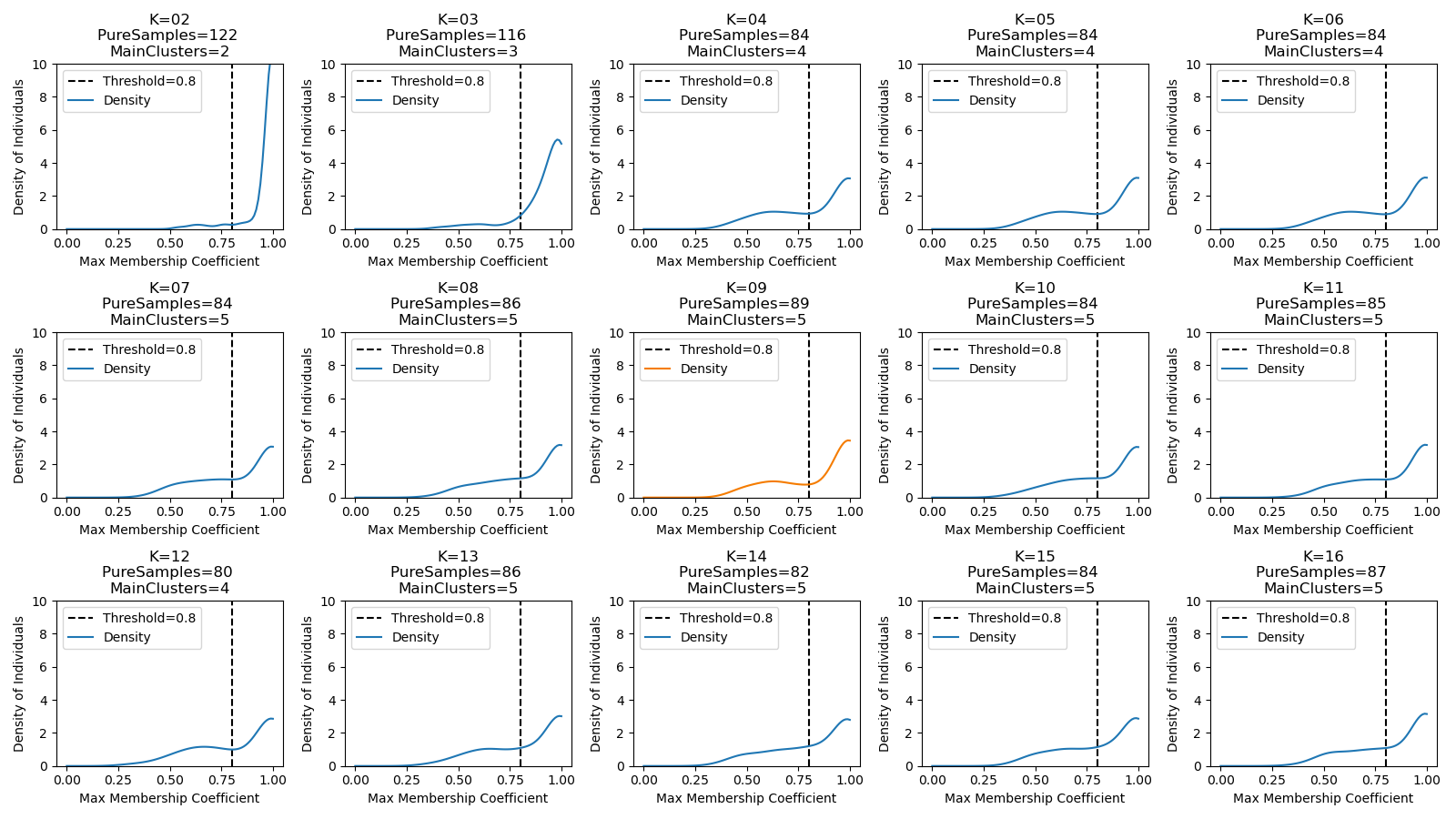

**Supplementary Figure 13 | Density distribution of genetic membership coefficients in Occidental pears mapped onto the *Pyrus pyrifolia* reference genome.**

A cutoff of 80% of the maximum membership coefficient within each genetic cluster was used to define pure samples. “Main clusters” refers to clusters containing at least one pure sample. The optimal number of clusters was *K* = 9.

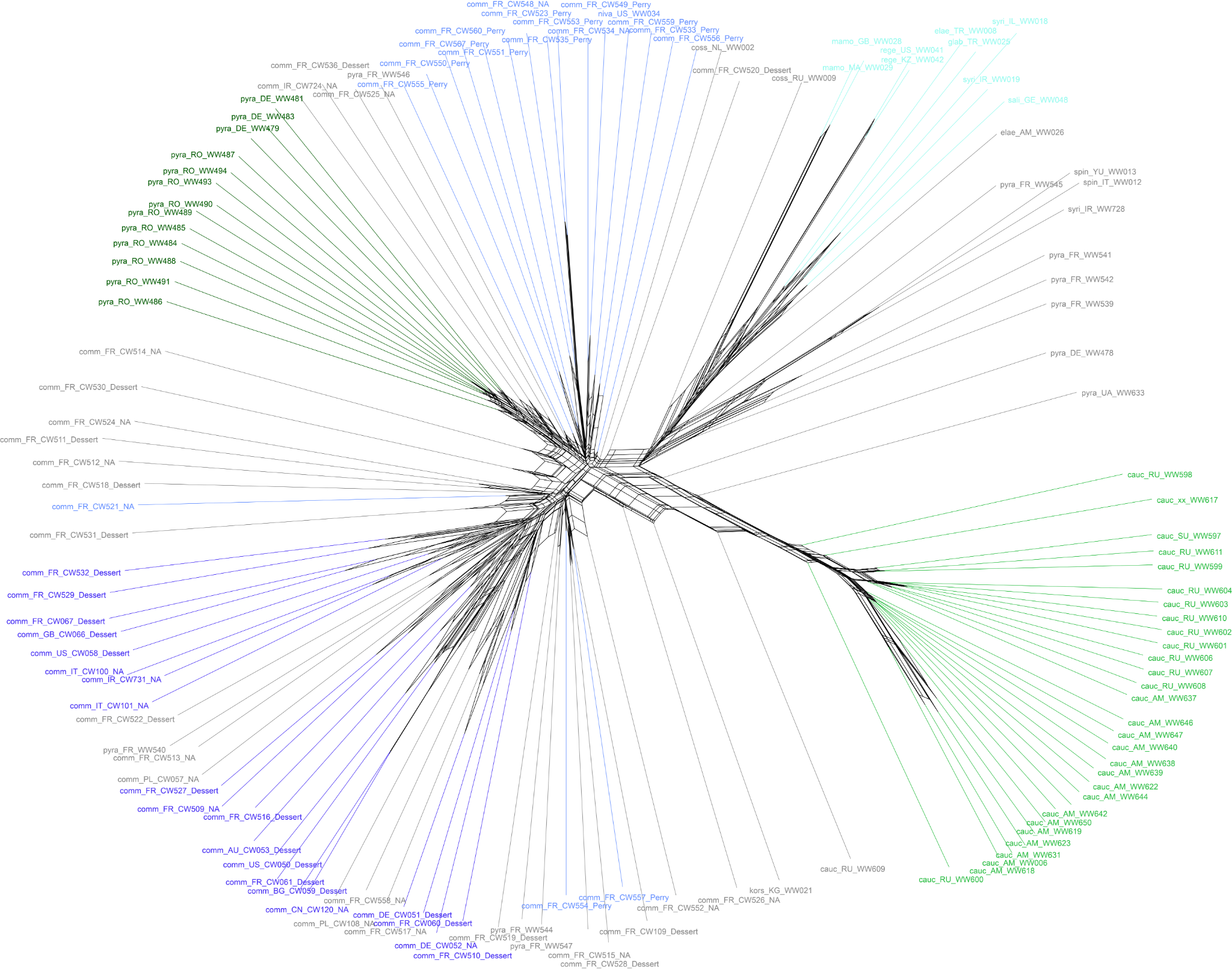

**Supplementary Figure 14 | Neighbor-net tree of Occidental pears.**

The tree was reconstructed using 22,358 synonymous unlinked SNPs from the genomes of 130 Occidental individuals mapped onto the *P. pyrifolia* reference genome*.* The colors of the branches and sample IDs match the primary genetic pool colors assigned in the population structure plot (*K*=9), while admixed samples (with a cutoff of 80% membership) are shown in gray.

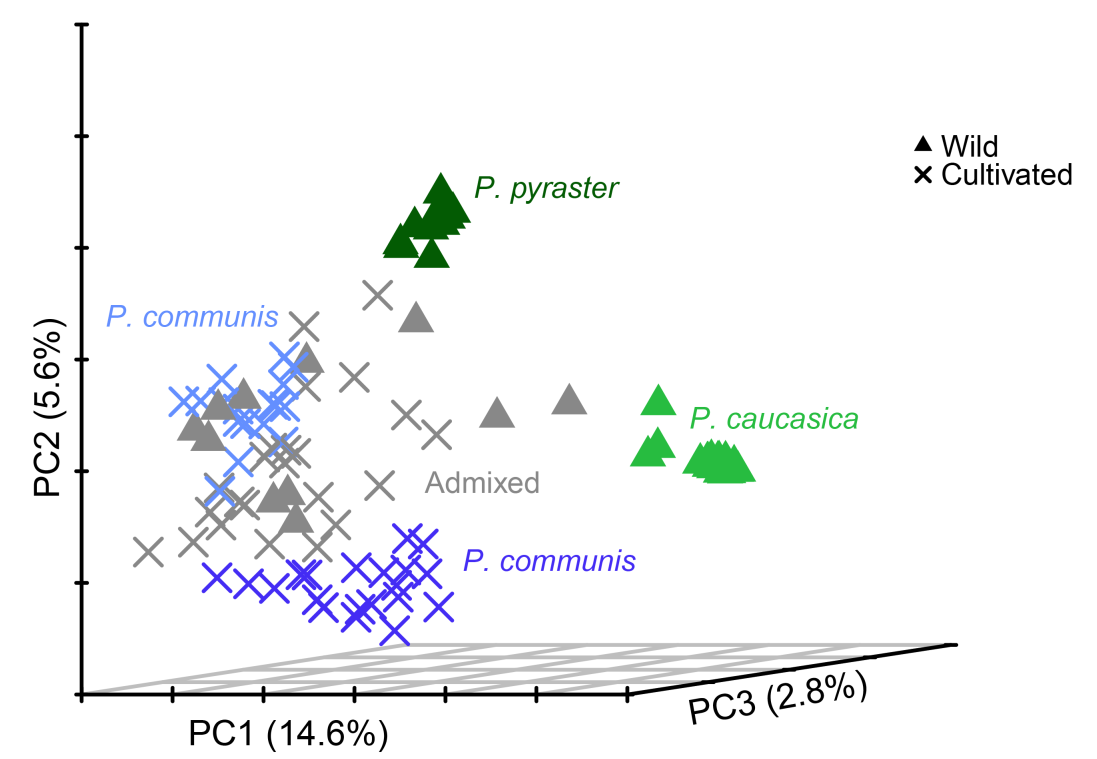

**Supplementary Figure 15 | PCA of Occidental pears mapped onto the *Pyrus pyrifolia* reference genome.**

Each point represents one individual. Colors indicate the primary genetic clusters inferred from the population-structure analysis (*K* = 9), with admixed individuals (membership < 80%) shown in gray.

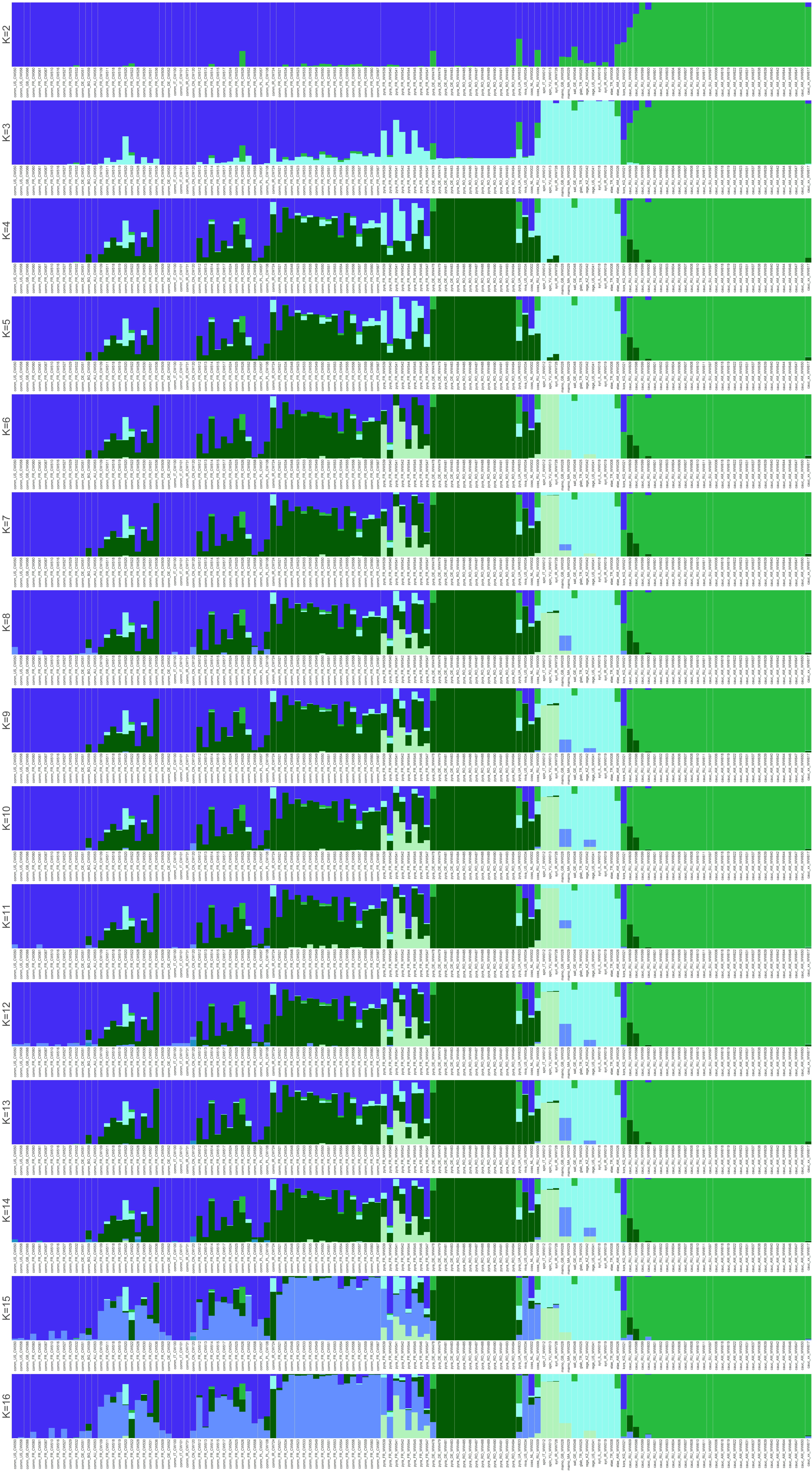

**Supplementary Figure 16 | Population structure landscape (*K*=2 to *K*=16) for Occidental pears mapped onto the *P. communis* reference genome.**

The analysis was performed using 20,298 synonymous unlinked SNPs from the genomes of 130 Occidental pear individuals mapped onto the *P. communis* reference genome. The figure was generated using fastSTRUCTURE, CLUMPAK, and Pophelper. The Q values for optimal *K*=15 are provided in Supplementary Table 3.

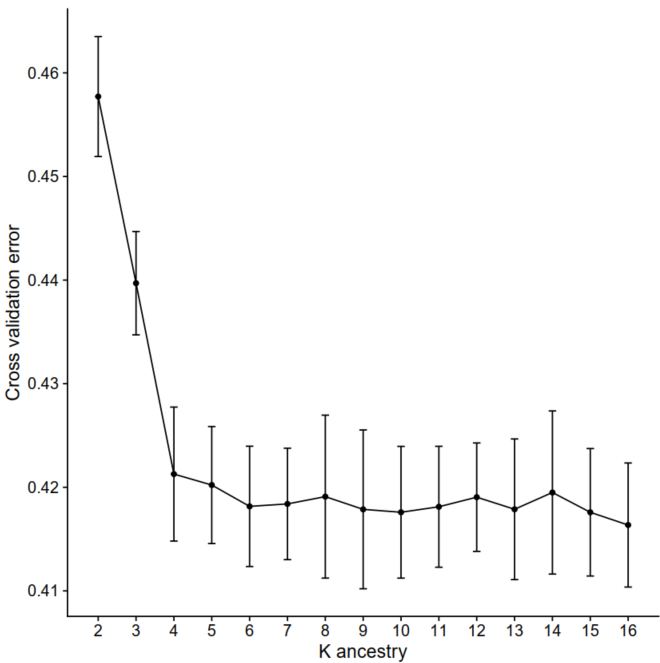

**Supplementary Figure 17 | Cross-validation error for different *K* ancestries in Occidental pears mapped onto the *P. communis* reference genome.**

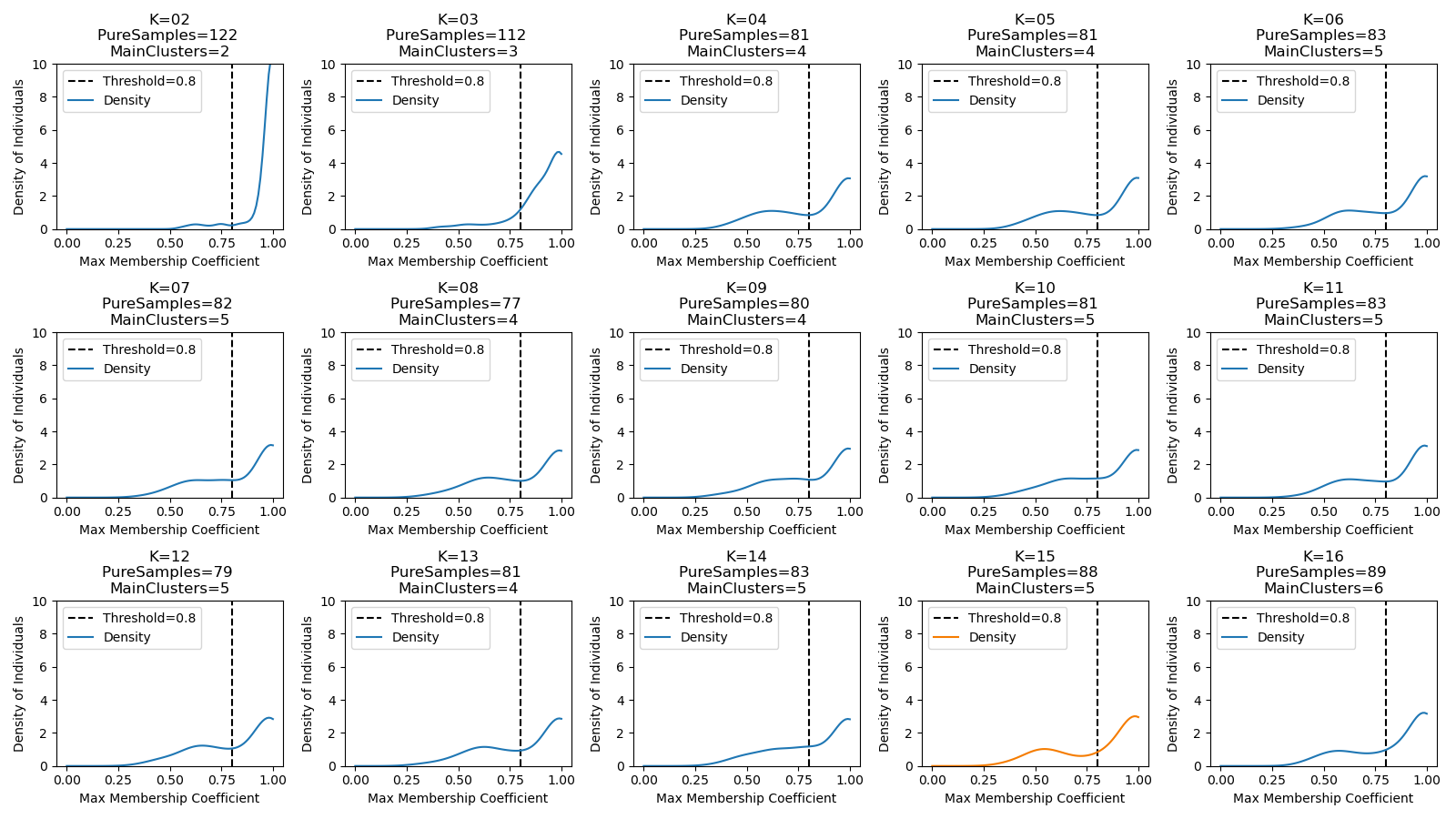

**Supplementary Figure 18 | Density distribution of genetic membership coefficients in Occidental pears mapped onto the *Pyrus communis* reference genome.**

A cutoff of 80% of the maximum membership coefficient within each genetic cluster was used to define pure samples. The optimal number of clusters was *K* = 15.

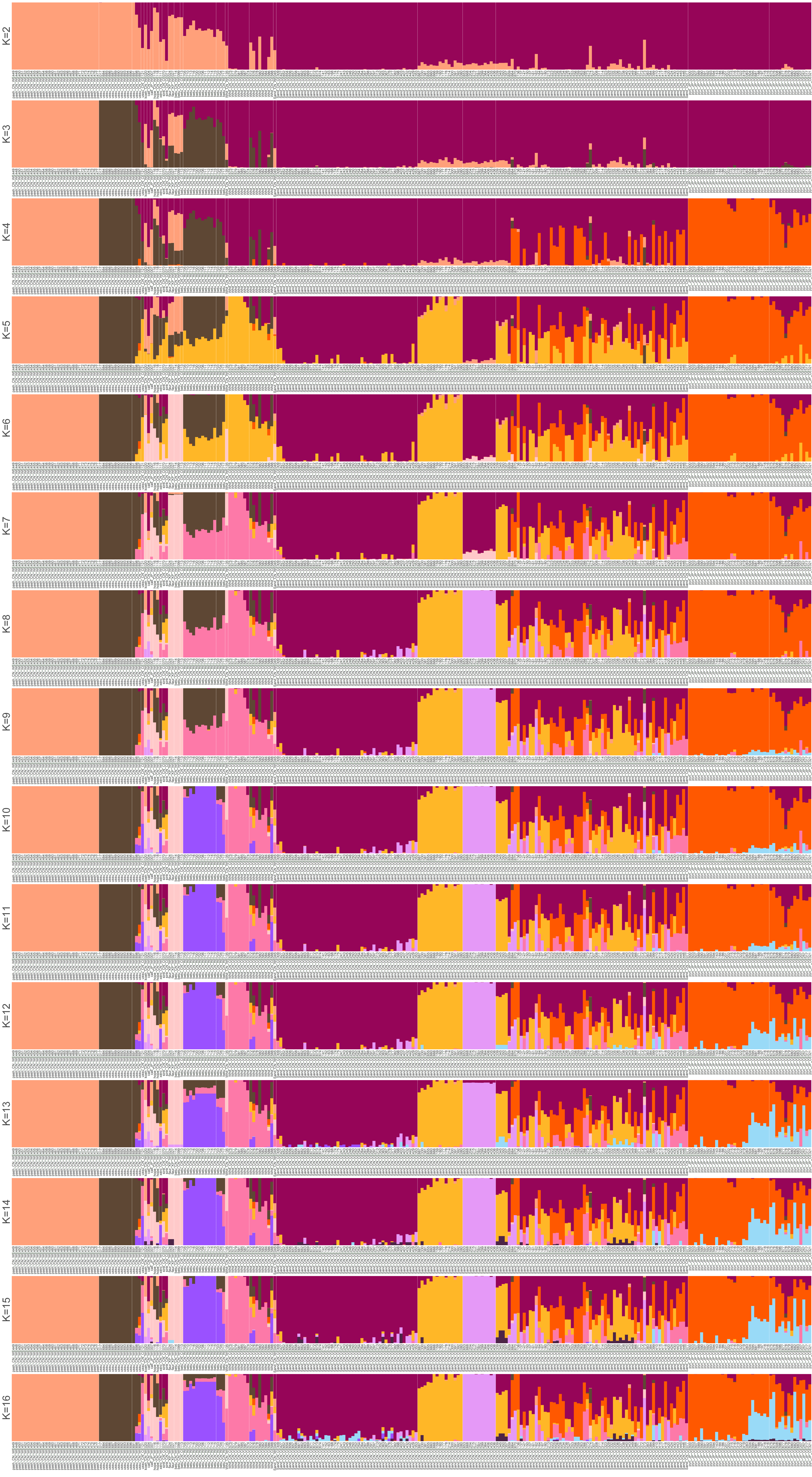

**Supplementary Figure 19 | Population structure landscape (*K*=2 to *K*=16) for Oriental pears mapped onto the *P. pyrifolia* reference genome.**

The figure was generated using fastSTRUCTURE, CLUMPAK, and PopHelper. The Q values for each cluster at the optimal *K* = 10 are provided in Supplementary Table 2.

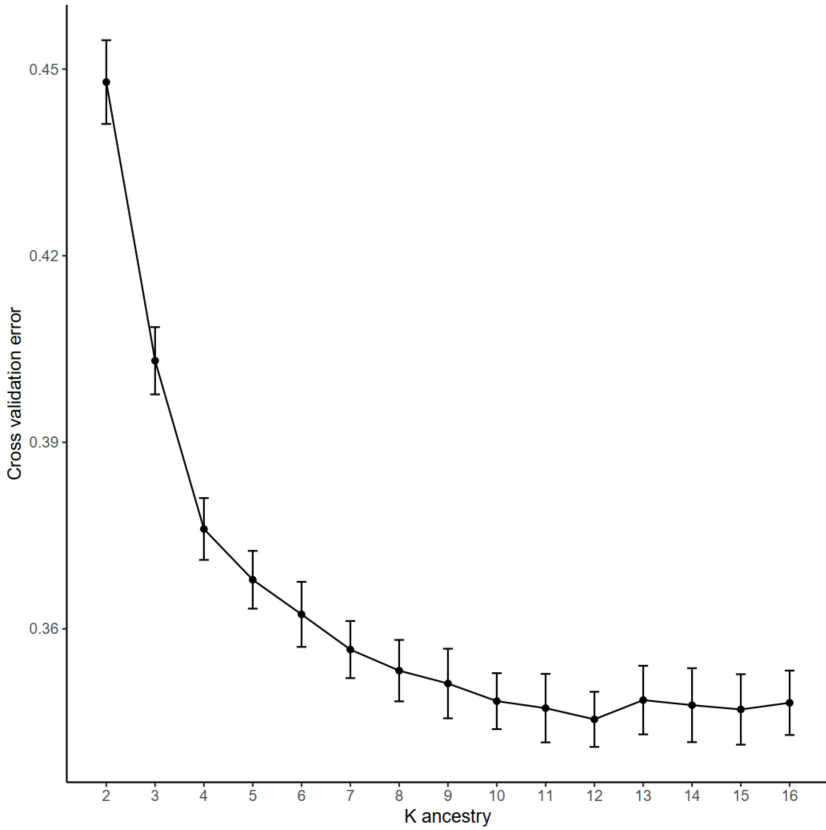

**Supplementary Figure 20 | Cross-validation error for *K* ancestries of Oriental pears mapped onto the *P. pyrifolia* reference genome*.***

The optimal *K* value was 10.

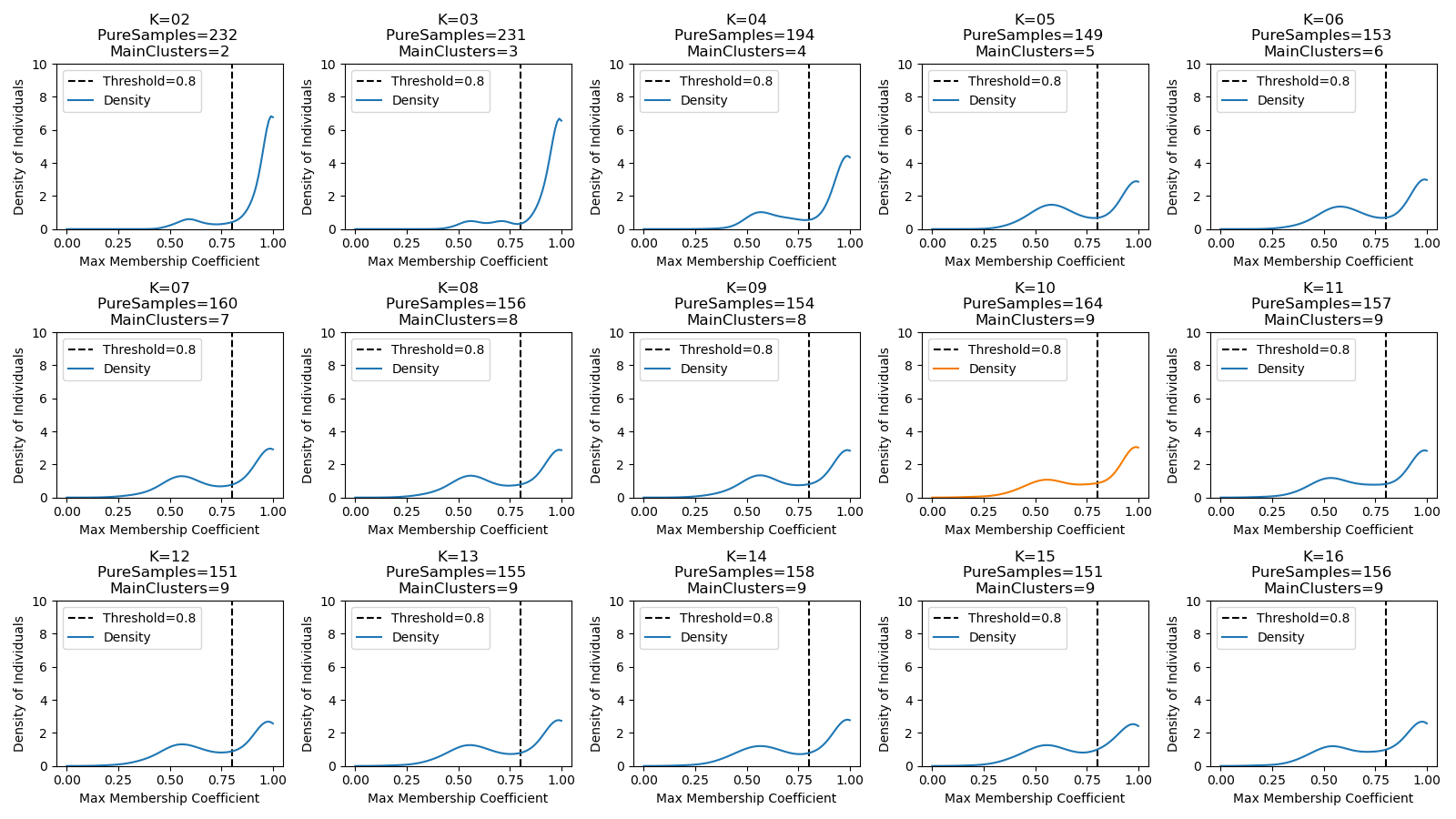

**Supplementary Figure 21 | Density distribution of genetic membership coefficients in Oriental pears mapped onto the *Pyrus pyrifolia* reference genome.**

A cutoff of 80% of the maximum membership coefficient within each genetic cluster was used to define pure samples. “Main clusters” refers to clusters containing at least one pure sample. The optimal number of clusters was *K* = 10.

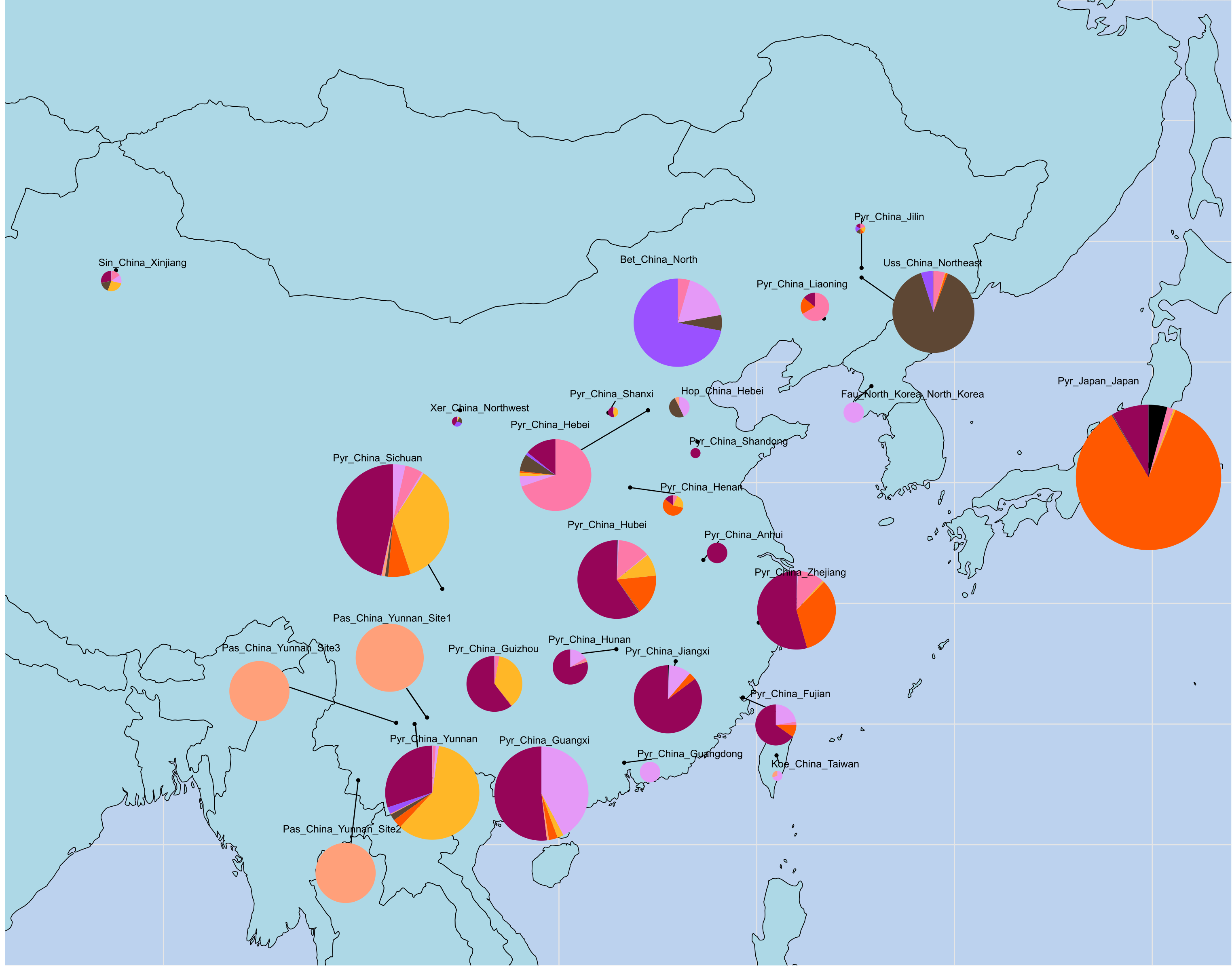

**Supplementary Figure 22 | Geographical distribution and genetic structure of Oriental pears.**

Pie charts represent the membership coefficients of genetic clusters inferred from fastSTRUCTURE, with colors corresponding to those used in the population-structure plots (*K =* 10). Each pie chart is positioned at the approximate sampling location of the corresponding individuals. The first three letters of each label indicate the species abbreviation (for example, Pyr for *Pyrus pyrifolia*), followed by the region or site name.

Species abbreviations: Pyr, *Pyrus pyrifolia*; Bet, *P. betulifolia*; Uss, *P. ussuriensis*; Fau, *P. fauriei*; Sin, *P. sinkiangensis*; Pas, *P. pashia*; Koe, *P. koehnei*; Xer, *P. xerophila*; Hop, *P. hopeiensis*.

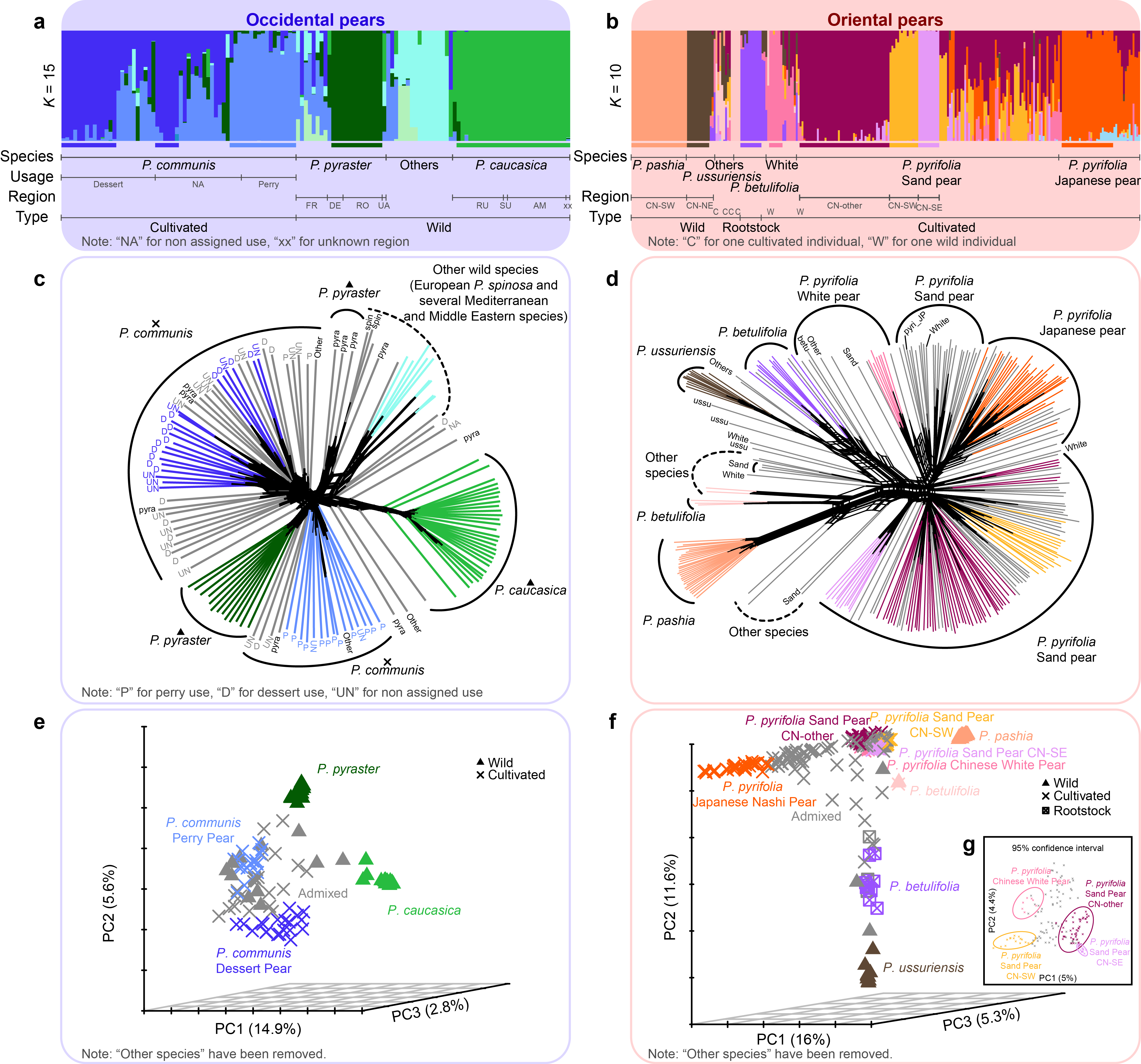

**Supplementary Figure 23 | Population genetic structure, differentiation, and variation in Occidental and Oriental pears (*Pyrus* spp.) (Full annotated Figure 1).**

**a,b** Population structure of wild and cultivated pears (*N* = 396 individuals) inferred with fastSTRUCTURE: Occidental pears (*N* = 130; 20,298 synonymous, unlinked SNPs) at *K* = 15 (a) and Oriental pears (*N* = 266; 21,141 synonymous, unlinked SNPs) at *K* = 10 (b). (Sub)species name, sampling region, and usage type were obtained from sampling records. Wild Occidental and Oriental lineages were defined based on geographical distribution, using the Tianshan-Hindu Kush Mountains as the boundary. Cultivar and rootstock groups were defined by their gene pools and planting locations. Region codes follow the ISO 3166-1 alpha-2 standard, with “xx” indicating an unknown region. “CN-SE”, “CN-SW”, and “CN-NE” refer to China Southeast, China Southwest, and China Northeast, respectively. Regions are ordered by geographic longitude or by cultivar usage category. **c,d** Neighbor-Net tree for the 130 Occidental pears, based on 20,298 unlinked and synonymous SNPs mapped onto *P. communis* (c), and the 266 Oriental pears based on 21,141 SNPs mapped onto *P. pyrifolia* (d). Branch colors correspond to the major gene pool colors in the population structure plots. Admixed or uncertainly assigned individuals (maximal membership coefficient ≤ 0.8), also including one pure Occidental individual that appeared admixed when mapped onto *P. pyrifolia*, are shown in gray. “D” and “P” indicate the two main uses of Occidental *P. communis* cultivars (also called the European pear): dessert and perry, respectively. “UN”, unassigned, denotes individuals that either serve a dual purpose (dessert and perry) or lack prior use metadata. Leaf labels include the abbreviated (sub)species name, with the final two underlined letters indicating the ISO country code of the sampling location. **e,f** Principal component analysis (PCA) of 113 Occidental (20,263 synonymous and unlinked SNPs mapped onto *P. communis*) (e) and 256 Oriental individuals (21,163 synonymous and unlinked SNPs mapped onto *P. pyrifolia*) (f), after excluding 17 and 10 “other” individuals, respectively. **g** PCA plot of Chinese cultivated P. pyrifolia sand and white pears.

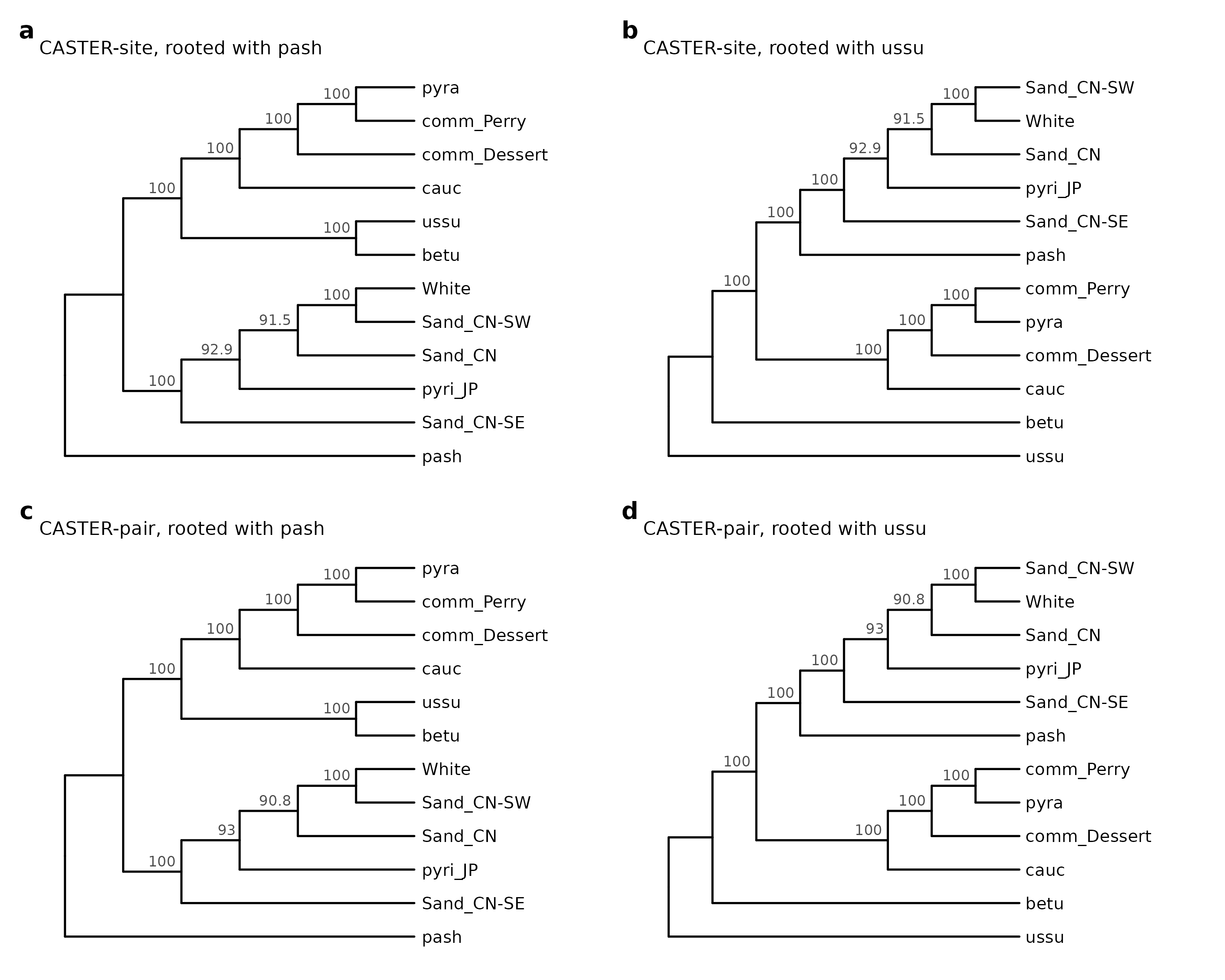

**Supplementary Figure 24 | Population trees of pear inferred with CASTER.**

Population trees were inferred from phased SNPs using CASTER-site (a, b) and CASTER-pair (c, d), rooted with *P. pashia* (a, c) or *P. ussuriensis* (b, d). Branch support values are shown on internal branches; branch lengths are not drawn to scale.

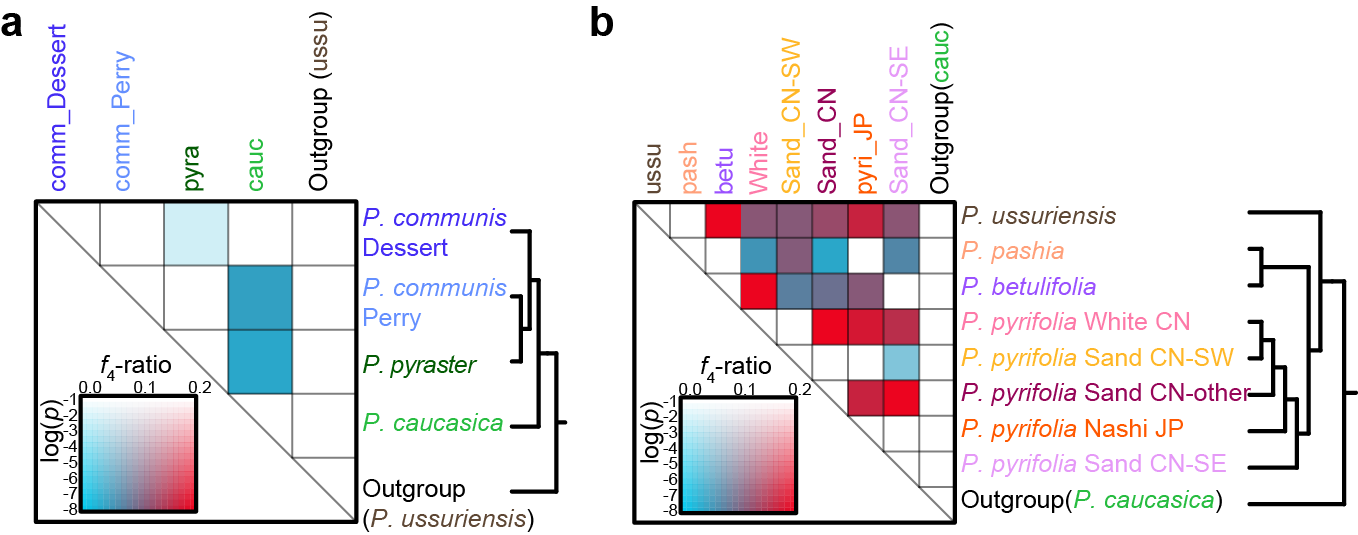

**Supplementary Figure 25 | Heatmaps of *f*_4_-ratio statistics computed with Dsuite under the tree model.**

**a,b.** The heatmaps are accompanied by the corresponding *P* values (*P* < 0.01) for the Occidental (a) and Oriental (b) pear populations. The reference topology trees from Fig. 2a shown beside the heatmaps were used as input for Dsuite to arrange the trios. Source data for this plot are provided in Supplementary Tables 5 and 6.

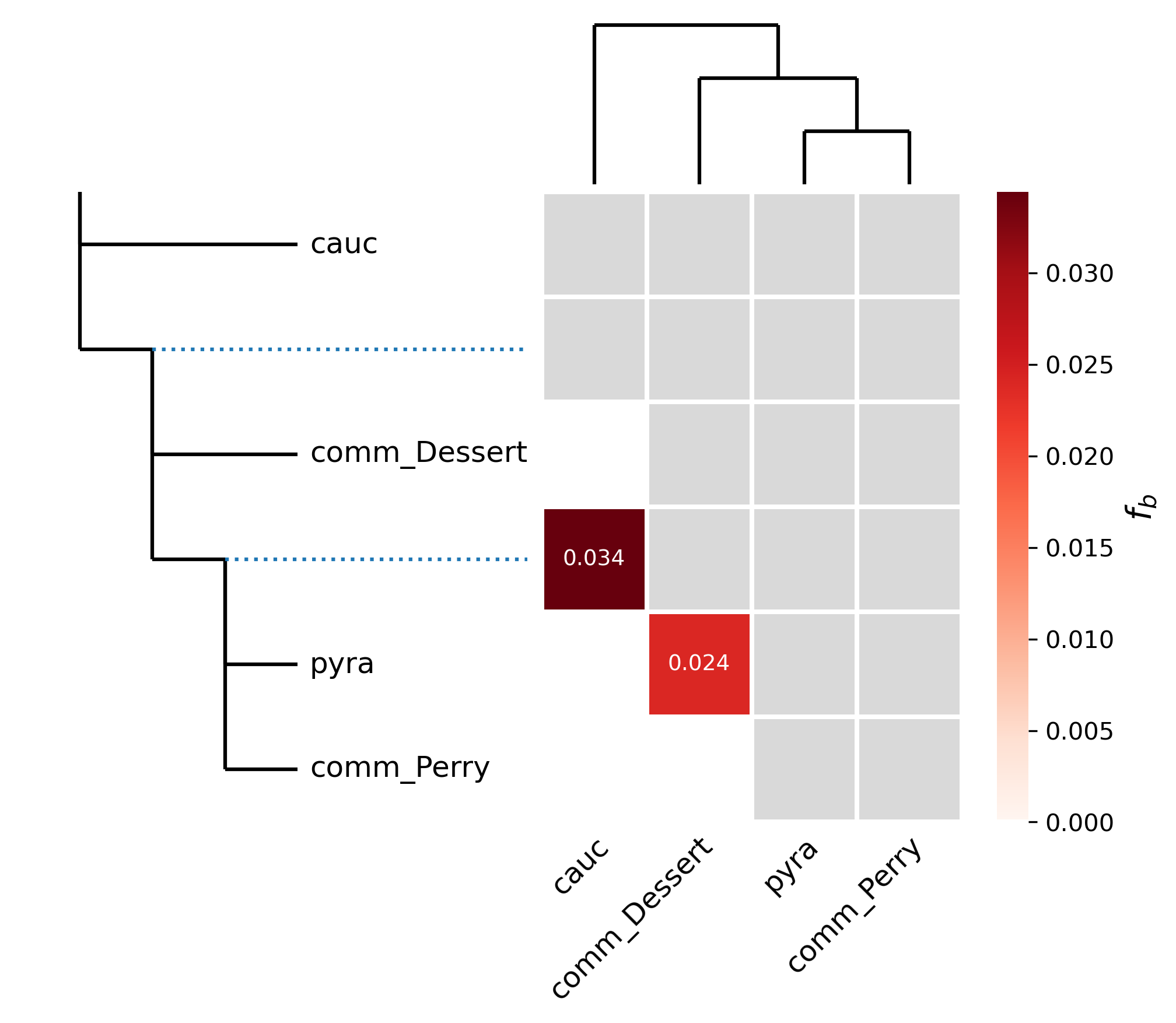

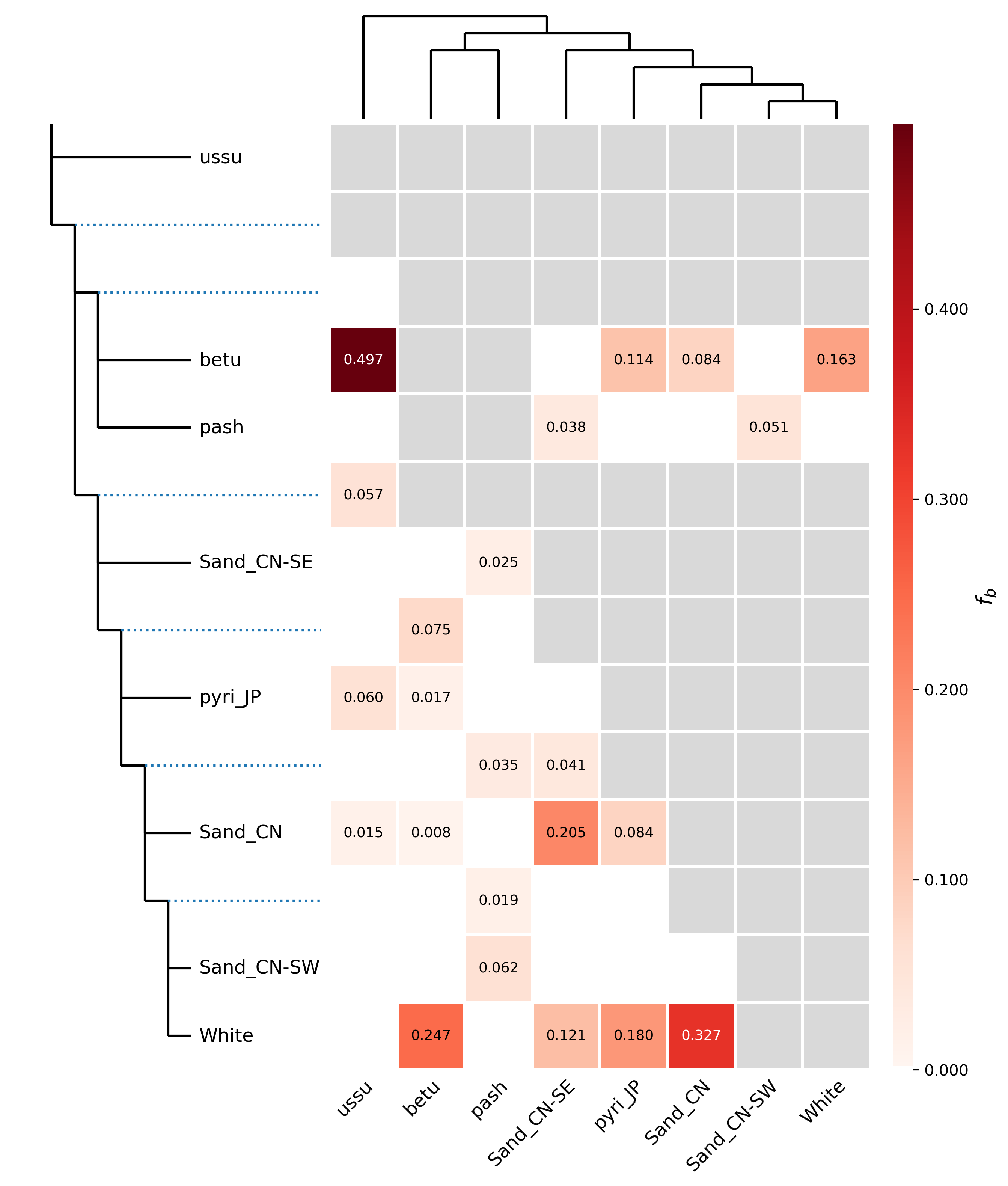

**Supplementary Figure 26 | *F-*branch statistics of introgression among Occidental and Oriental pear lineages.**

*F*-branch (*f*_b_) matrix computed with Dsuite Fbranch from *f*_4_-ratio and *D*-statistic trio results (Dsuite Dtrios). Left pannel: Occidental lineage; right pannel: Occidental lineage. The species tree topology is shown on the left (rooted with *P. caucasica* as outgroup for Oriental pears, and *P. ussuriensis* as outgroup for Occidental pears). Rows correspond to branches of the tree, solid black lines to the terminal (extant) populations and blue dashed lines to internal (ancestral) branches; columns correspond to terminal populations used as P3. Cell color indicates the *f*-branch value, representing excess allele sharing between the lineage descending from branch b and taxon P3 that cannot be explained by incomplete lineage sorting.

Population abbreviations:

betu, *P. betulifolia*;

cauc, *P. caucasica*;

comm Dessert, *P. communis* dessert;

comm Perry, *P. communis* perry;

pyra, *P. pyraster*;

pash, *P. pashia*;

pyri JP, *P. pyrifolia* Nashi Japan;

Sand CN, *P. pyrifolia* sand China-other;

Sand CN-SE, *P. pyrifolia* sand China-Southeast;

Sand CN-SW, *P. pyrifolia* sand China-Southwest;

ussu, *P. ussuriensis*;

White, *P. pyrifolia* white China.

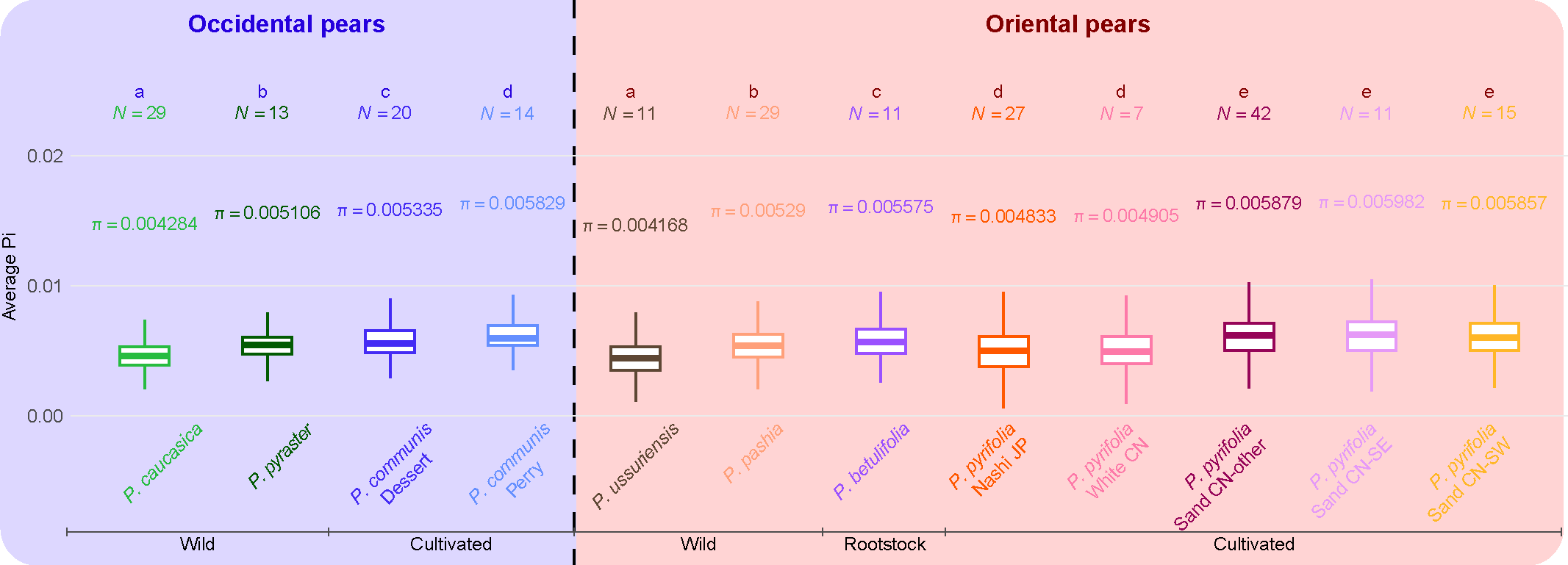

**Supplementary Figure 27 | Pairwise nucleotide diversity (π) across the 12 pear populations.**

π values were calculated using pixy in windows restricted to high-mappability regions, defined using GenMap, and refined by excluding centromeric regions identified with CentIER. Both Anderson-Darling and Shapiro-Wilk tests indicated that π distributions did not follow a normal distribution (*p* < 0.05). Pairwise Wilcoxon tests were therefore used to assess differences among populations; different letters denote populations that differ at Holm-adjusted *P* < 0.01. π was calculated from lineage-specific “all-sites” (variant and invariant) VCFs mapped to each lineage’s own reference genome (149,175,165 sites for *P. pyrifolia*; 127,678,724 sites for *P. communis*), so π values are not directly comparable between the two lineages; different letters were therefore assigned separately within each major lineage (Occidental or Oriental pears). *N* values indicate the number of individuals per population. Genome-wide π values, shown above each box, were obtained by dividing the total pairwise nucleotide differences by the total number of pairwise comparisons, rather than by averaging window-based π estimates.

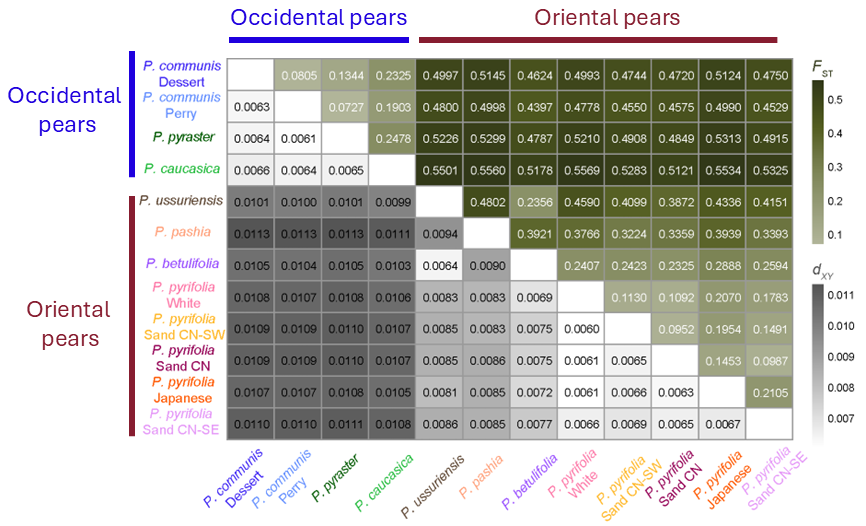

**Supplementary Figure 28 | Matrix of genomic differentiation (upper: *F*_ST_) and absolute divergence (lower: *d_XY_*) of pairwise pear populations.**

The values were calculated with pixy using SNPs mapped onto the *P. pyrifolia* reference genome.

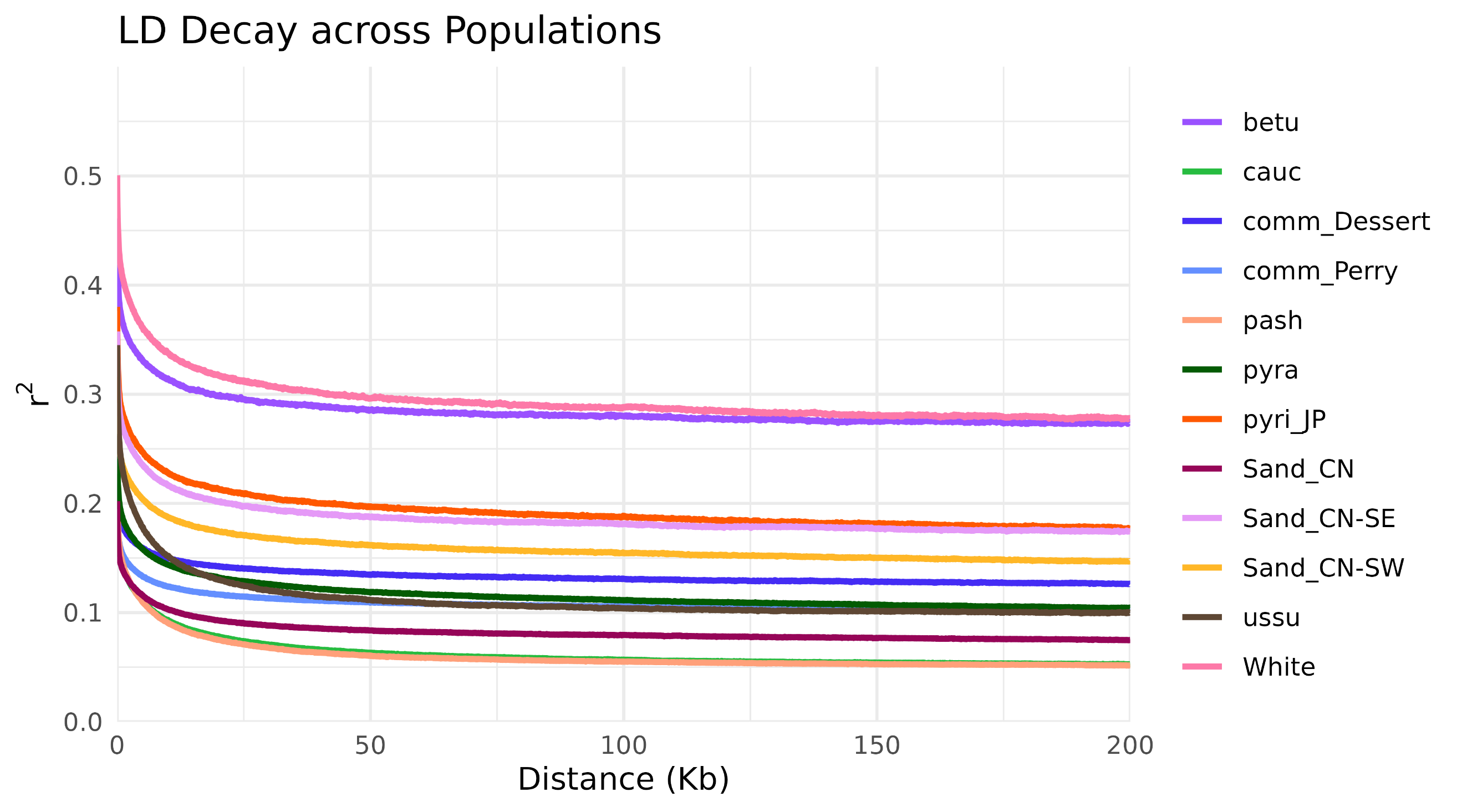

**Supplementary Figure 29 | Linkage disequilibrium (LD) decay patterns across the 12 pear populations.**

LD decay was calculated using PopLDdecay based on SNP markers mapped onto the *P. pyrifolia* reference genome, using the r² statistic.

Population abbreviations in this plot are as follows:

betu, *P. betulifolia*;

cauc, *P. caucasica*;

comm Dessert, *P. communis* dessert;

comm Perry, *P. communis* perry;

pyra, *P. pyraster*;

pash, *P. pashia*;

pyri JP, *P. pyrifolia* Nashi Japan;

Sand CN, *P. pyrifolia* sand China-other;

Sand CN-SE, *P. pyrifolia* sand China-Southeast;

Sand CN-SW, *P. pyrifolia* sand China-Southwest;

ussu, *P. ussuriensis*;

White, *P. pyrifolia* white China.

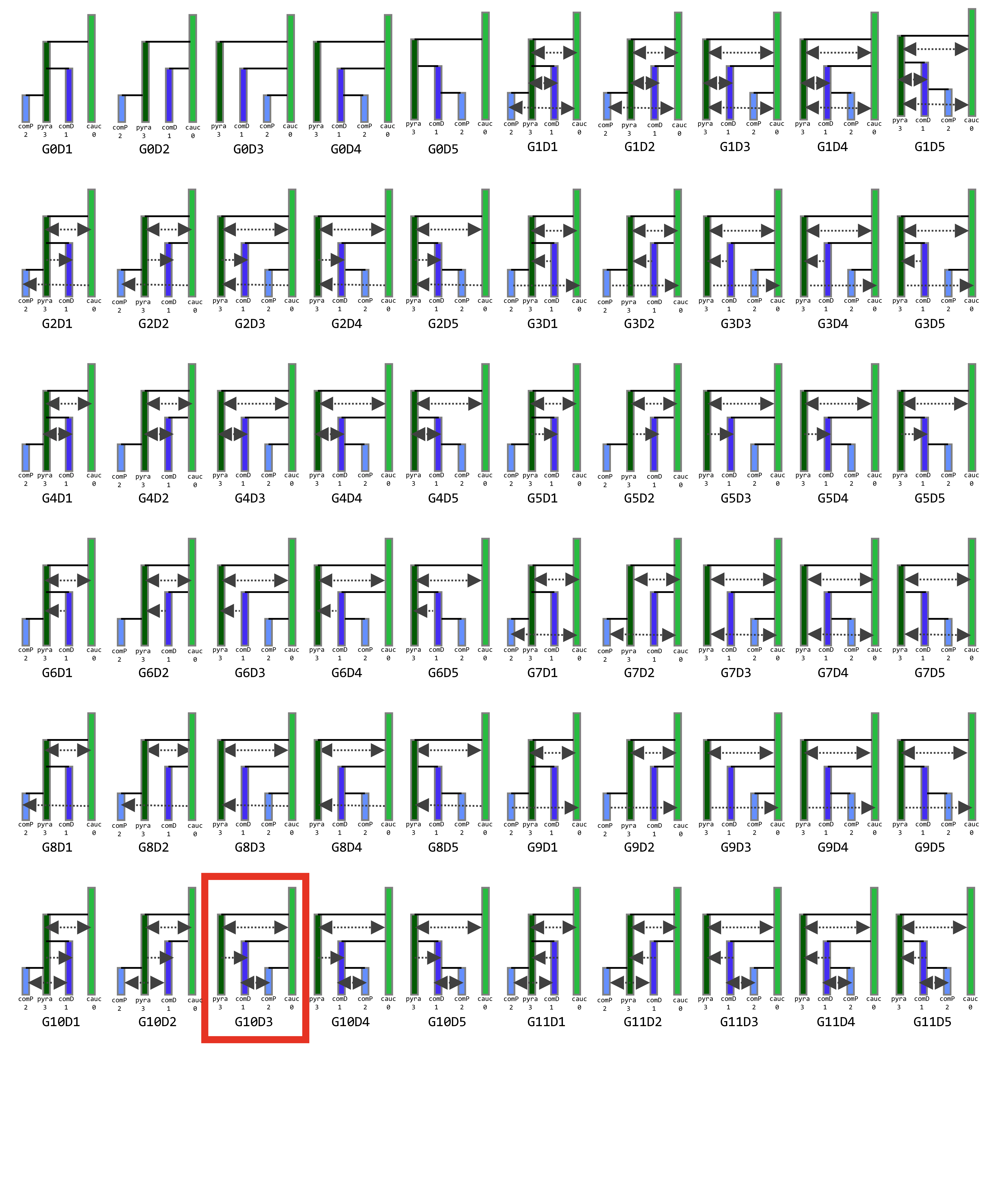

**Supplementary Figure 30 | Diagrams representing the demographic models used for fastsimcoal2 simulations of Occidental pears.**

Each colored bar represents a population, with colors corresponding to those used in the population structure analysis. Numbers indicate the population identifiers, as defined in the model configuration files. Horizontal solid lines denote population divergence events, while dashed lines indicate gene flow (migration), with the arrowheads indicating direction as modeled backward in time, between populations. The model highlighted with a red box represents the best-fitting model.

For compatibility with the simulation framework and scripting conventions, abbreviated population names were used in the model configuration files.

Population abbreviations:

cauc, *P. caucasica*; comD, *P. communis* dessert; comP, *P. communis* perry; pyra, *P. pyraster*.

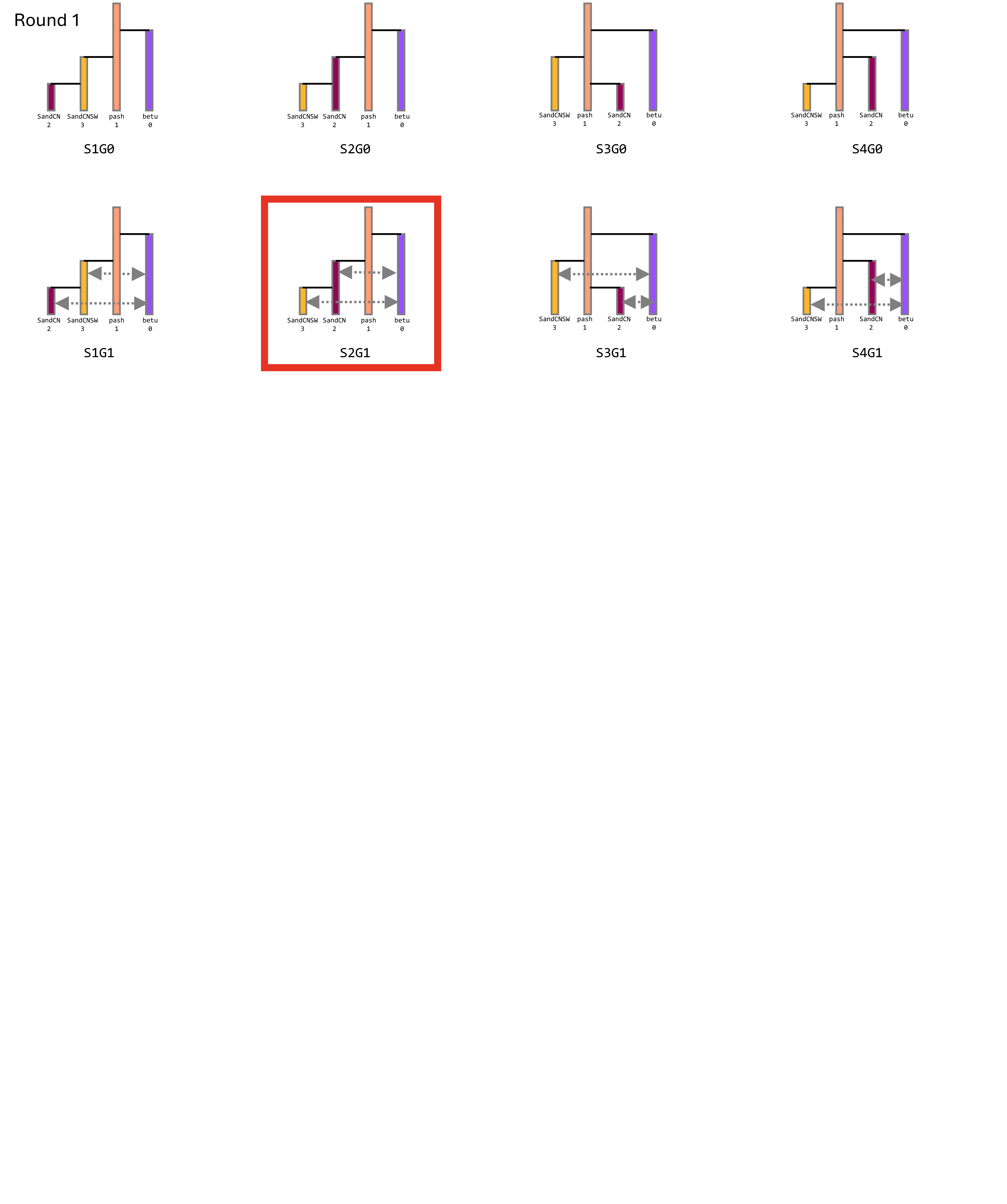

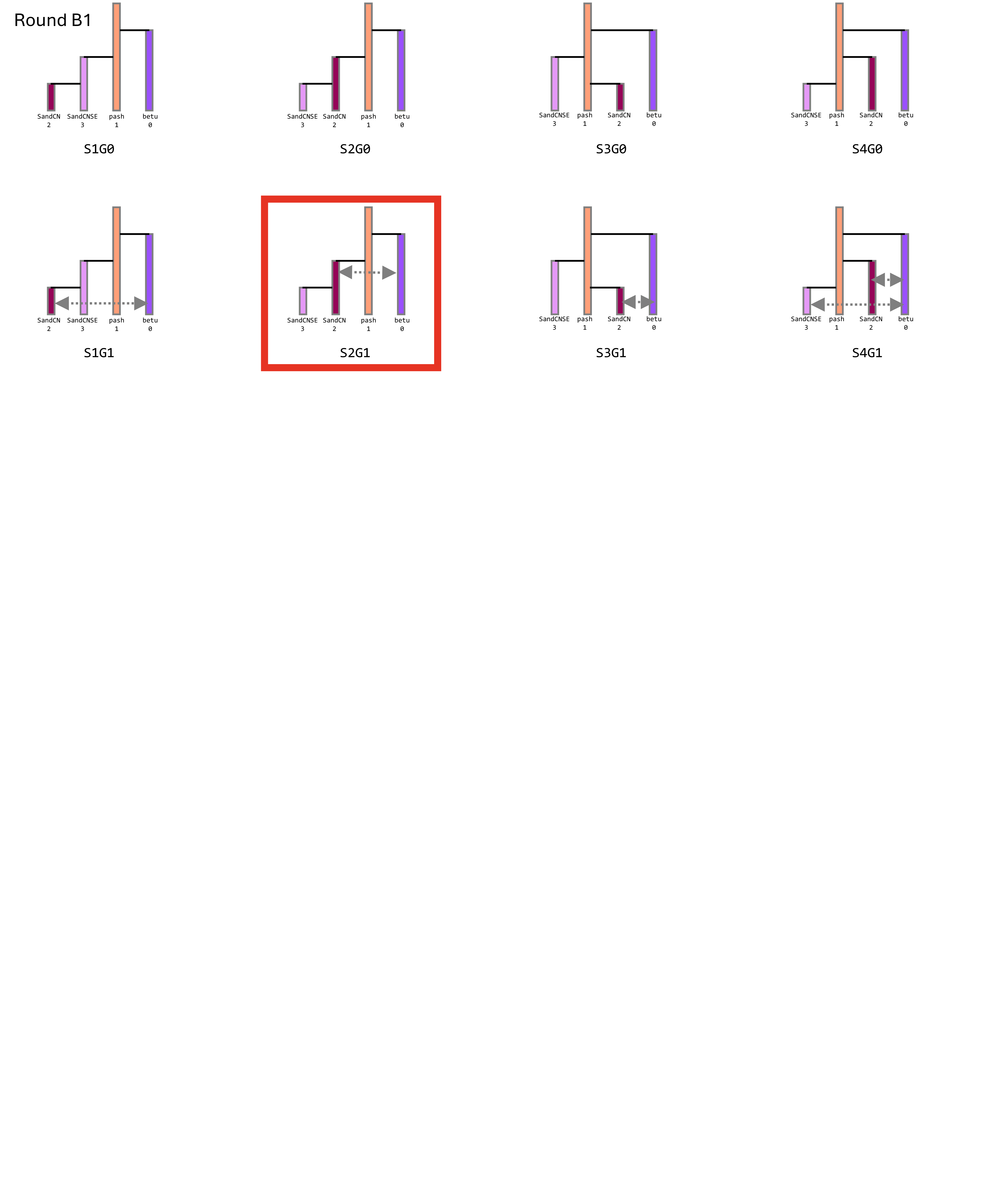

**Supplementary Figure 31 | Diagrams representing the demographic models used for fastsimcoal2 simulations of Oriental pears (Round 1 and Round B1).**

Each colored bar represents a population, with colors corresponding to those used in the population structure analysis. Numbers indicate the population identifiers, as defined in the model configuration files. Horizontal solid lines denote population divergence events, while dashed lines indicate gene flow (migration), with the arrowheads indicating direction as modeled backward in time, between populations. The model highlighted with a red box represents the best-fitting model in this round of analysis.

For compatibility with the simulation framework and scripting conventions, abbreviated population names were used in the model configuration files.

Population abbreviations:

betu, *P. betulifolia*; pash, *P. pashia;* pyriJP, *P. pyrifolia* Nashi Japan; SandCN, *P. pyrifolia* sand China-other; SandCNSE, *P. pyrifolia* sand China-southeast; SandCNSW, *P. pyrifolia* sand China-southwest.

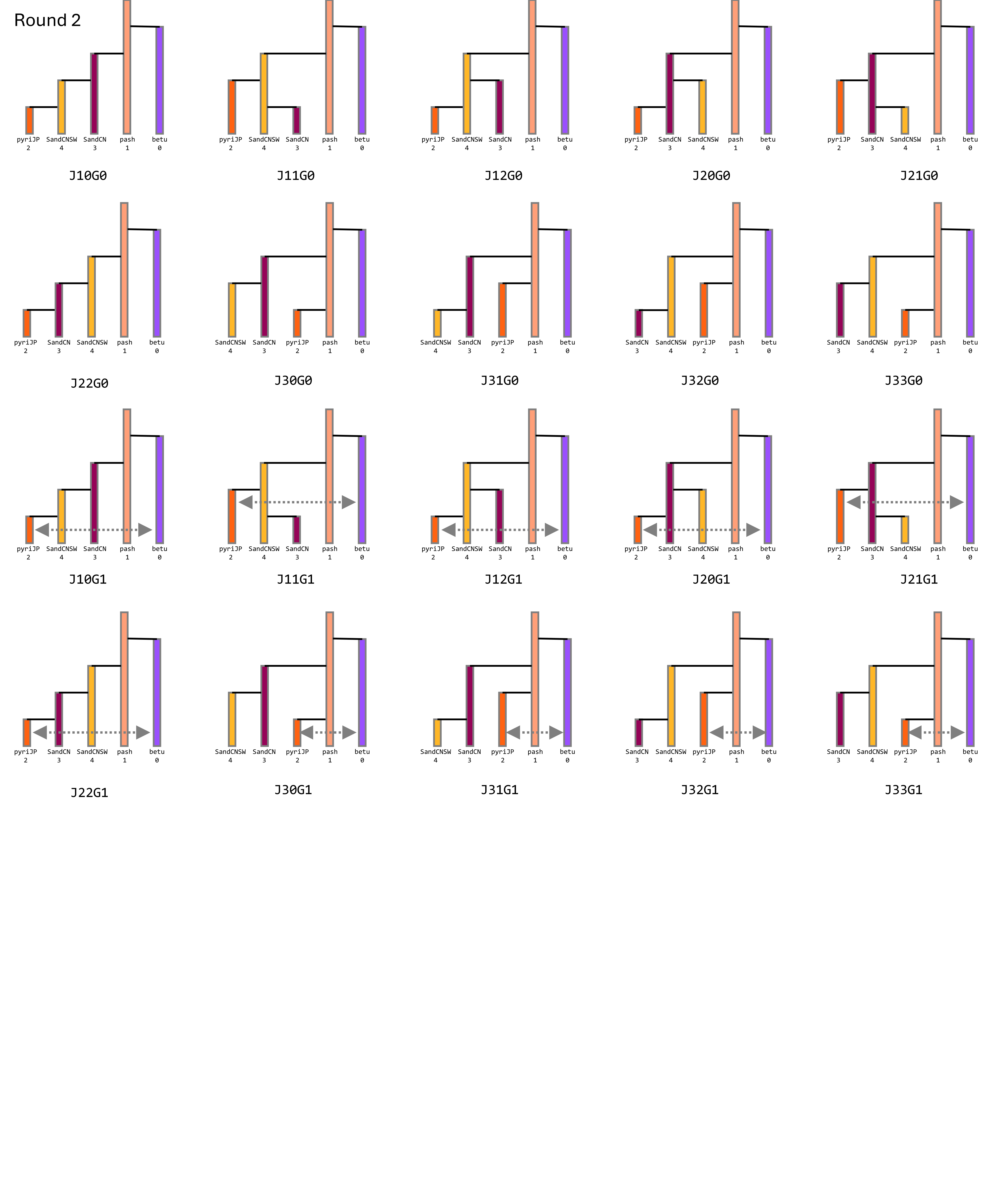

**Supplementary Figure 31 (continued) | Diagrams representing the demographic models used in fastsimcoal2 simulations of Oriental pears (Round 2).**

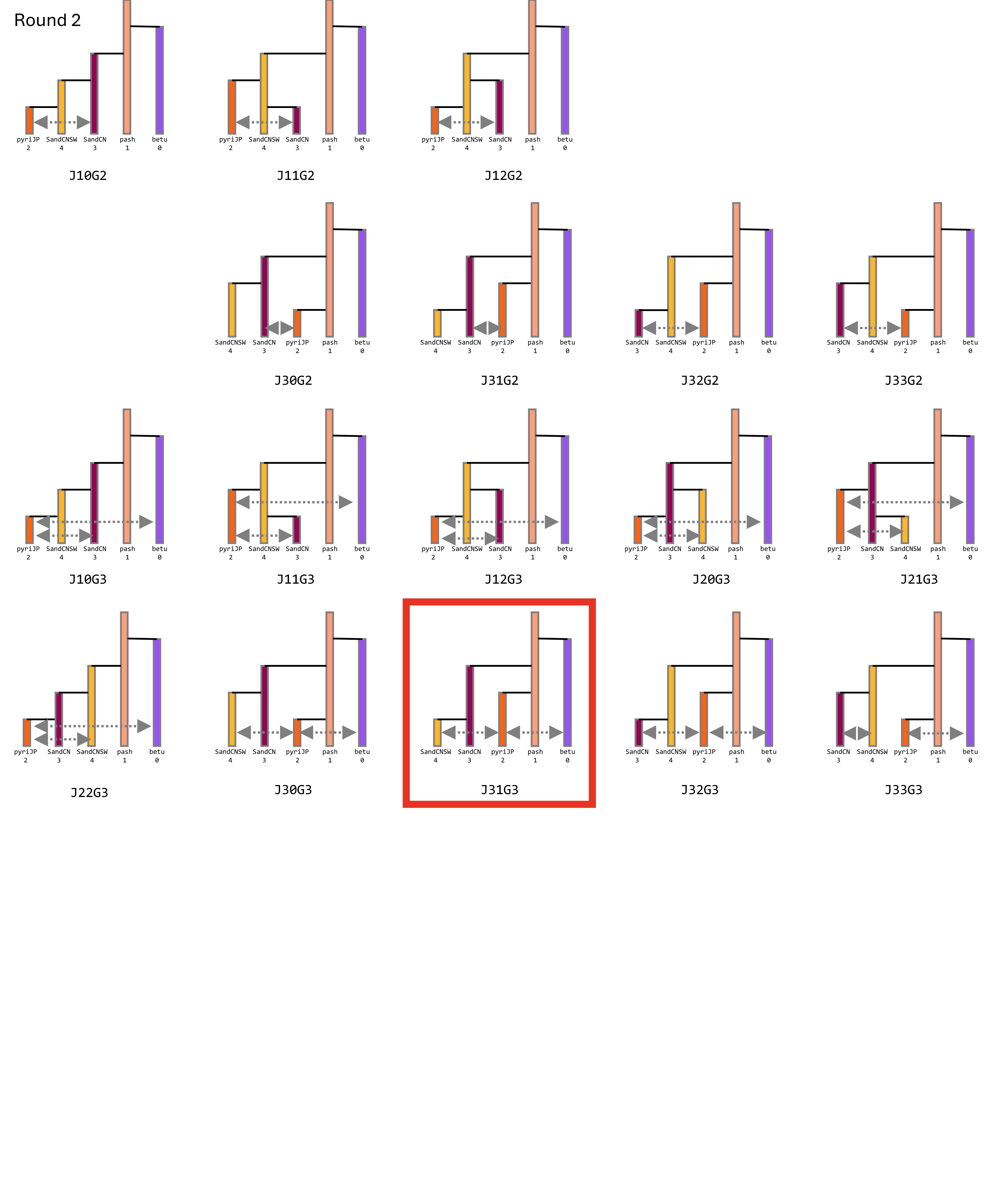

**Supplementary Figure 31 (continued) | Diagrams representing the demographic models used in fastsimcoal2 simulations of Oriental pears (Round 2).**

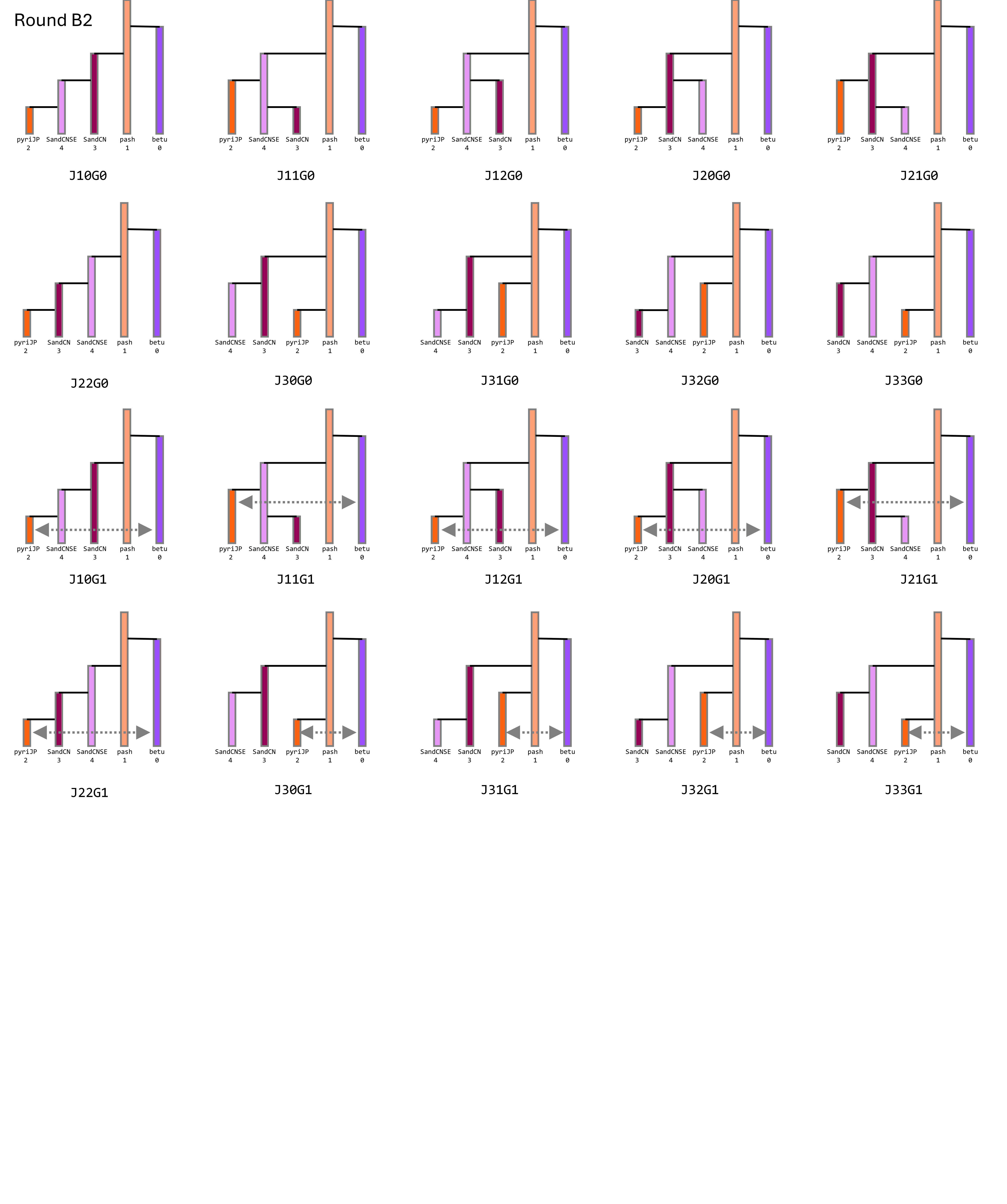

**Supplementary Figure 31 (continued) | Diagrams representing the demographic models used in fastsimcoal2 simulations of Oriental pears (Round B2).**

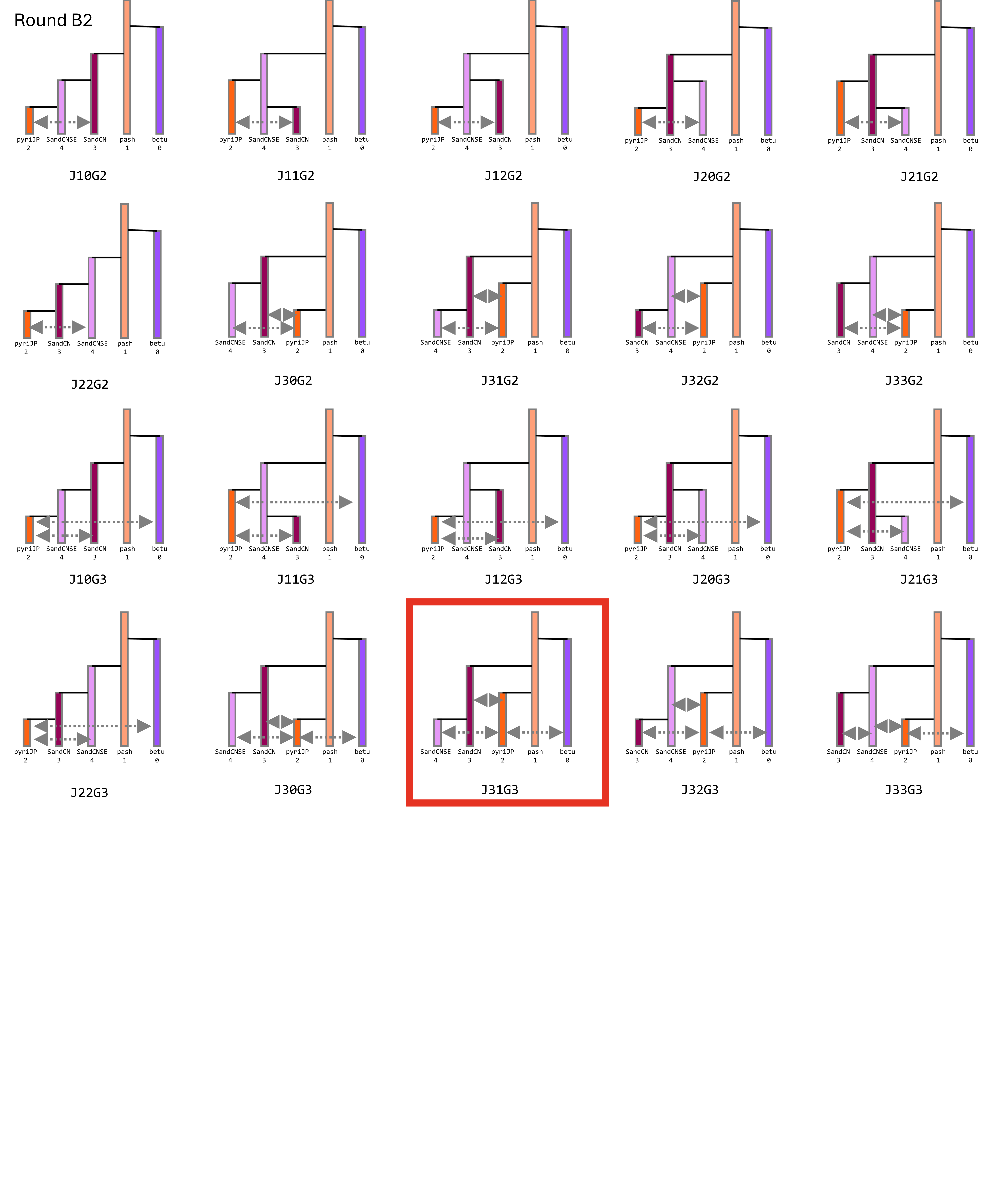

**Supplementary Figure 31 (continued) | Diagrams representing the demographic models used in fastsimcoal2 simulations of Oriental pears (Round B2).**

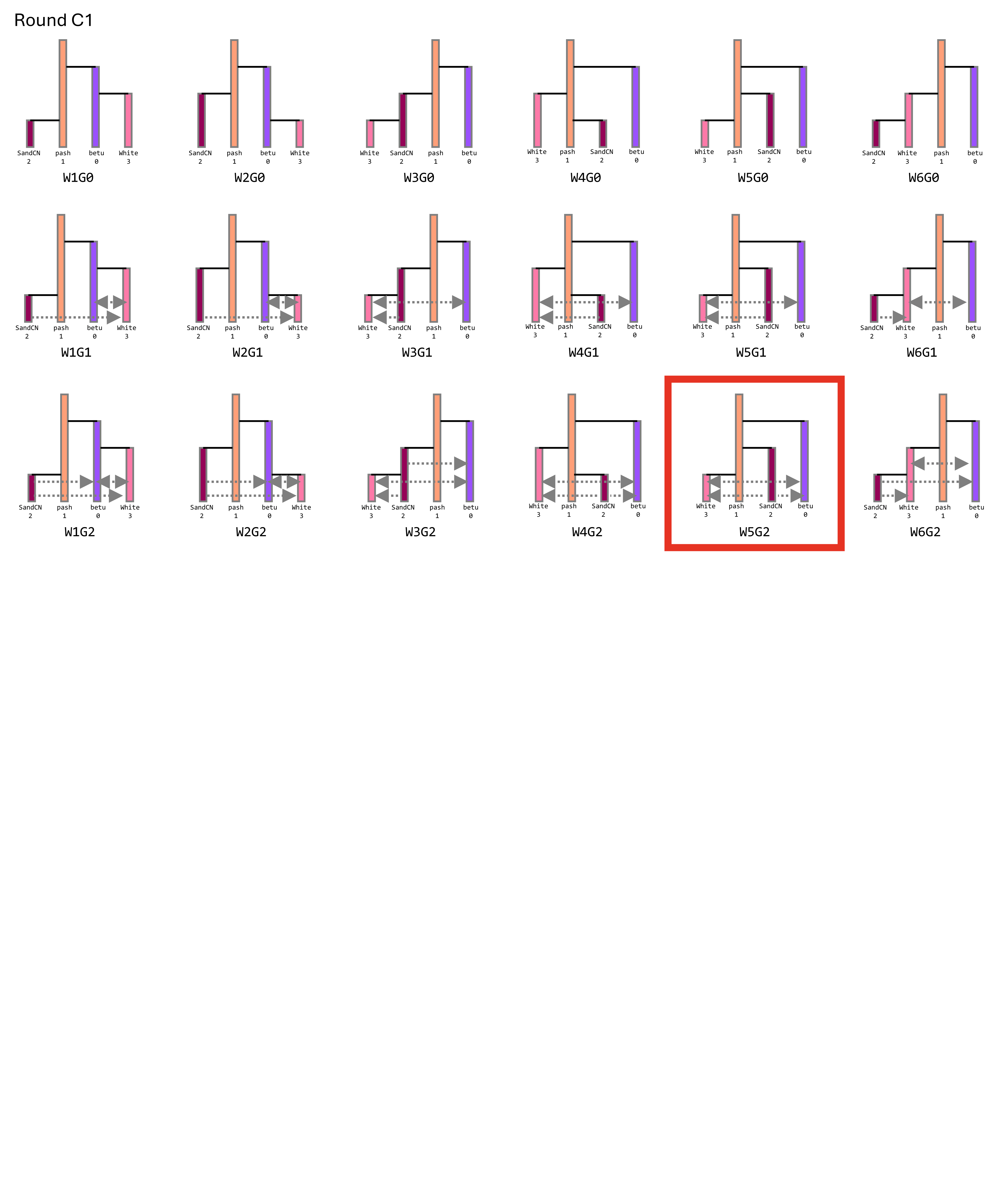

**Supplementary Figure 31 (continued) | Diagrams representing the demographic models used in fastsimcoal2 simulations of Oriental pears (Round C1).**

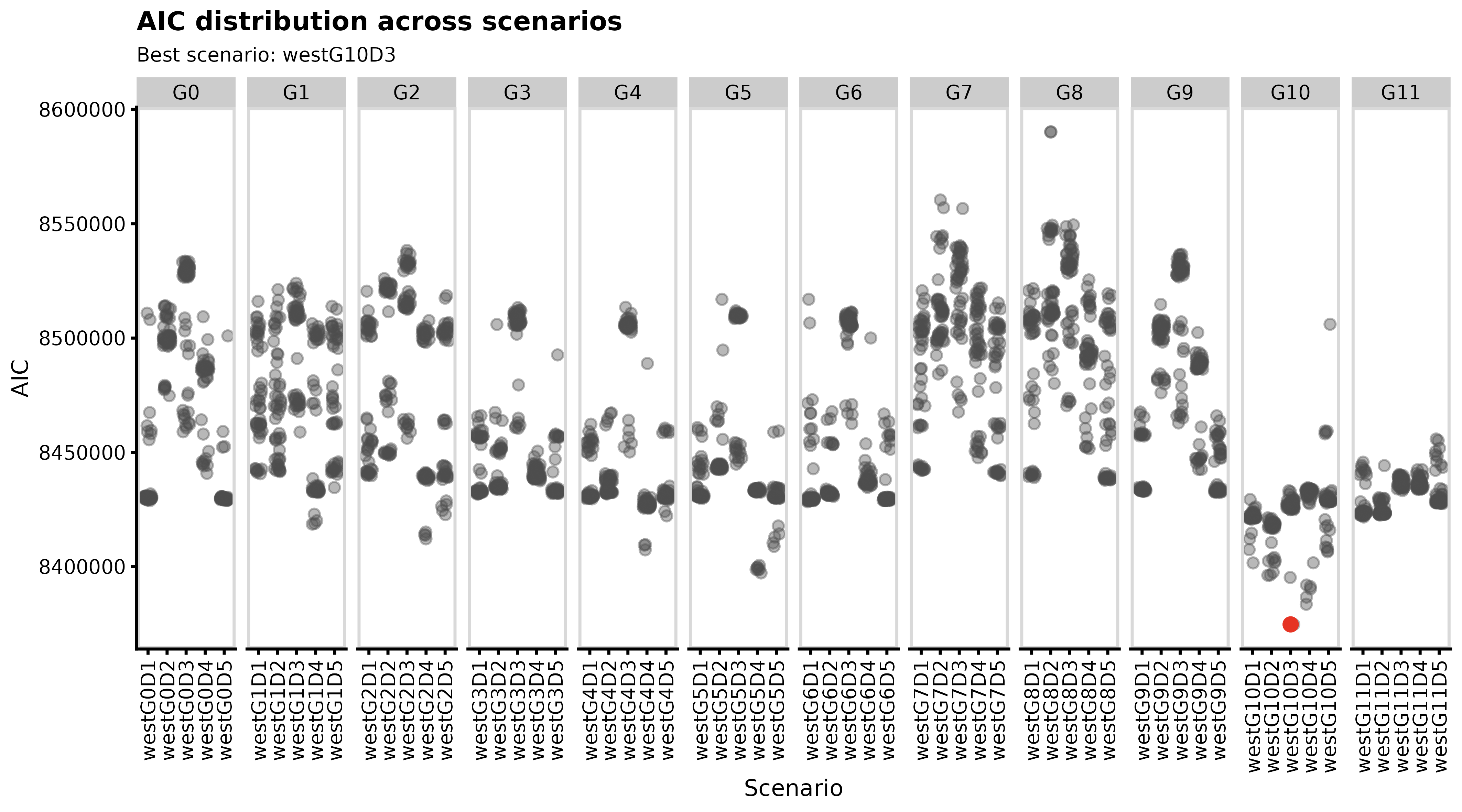

**Supplementary Figure 32 | Akaike information criterion (AIC) values for Occidental pear demographic models simulated with fastsimcoal2.**

Each point represents an independent run showing the best likelihood result obtained from 100,000 simulations per scenario, evaluated using the embedded MaxEstLhood function in fastsimcoal2. Each demographic scenario was replicated 50 times (50 × 100,000 simulations). The red point marks the best-fitting scenario with the lowest AIC value. The names of the scenarios correspond to the demographic models illustrated in Supplementary Figure 30.

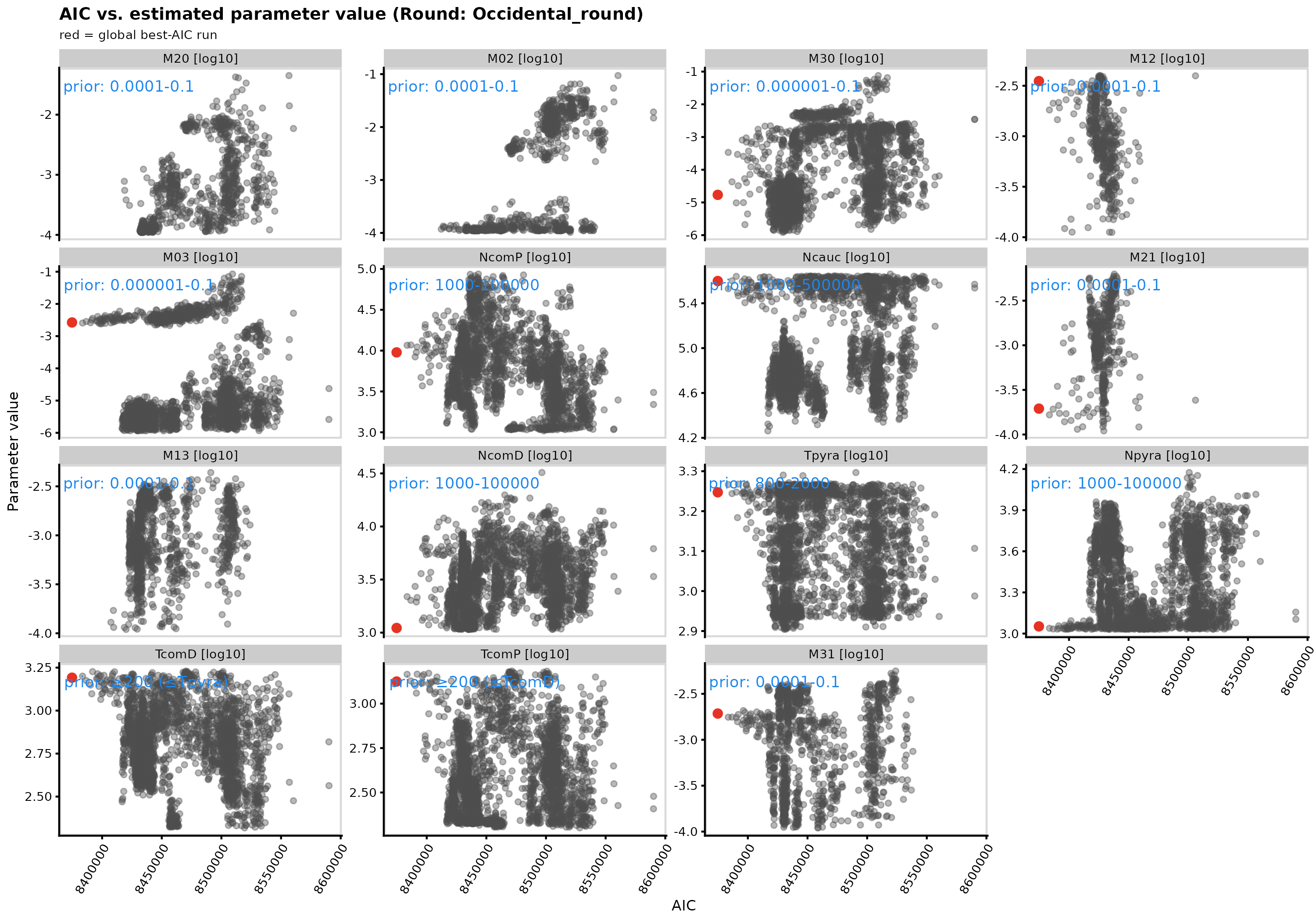

**Supplementary Figure 33 | Diagnostic relationship between Akaike information criterion (AIC) and estimated parameter values across fastsimcoal2 runs for the Occidental pear dataset.**

Each panel shows one demographic parameter (effective population size *N*, divergence time *T*, or migration rate *M*) plotted against the AIC of the run that produced it; values are log10-transformed for parameters spanning several orders of magnitude. Each point represents an independent run showing the best likelihood result obtained from 100,000 simulations, evaluated using the embedded MaxEstLhood function in fastsimcoal2. The blue text in each panel gives the prior range used for that parameter. The red point marks the global best-AIC run across all scenarios and replicates in this round.

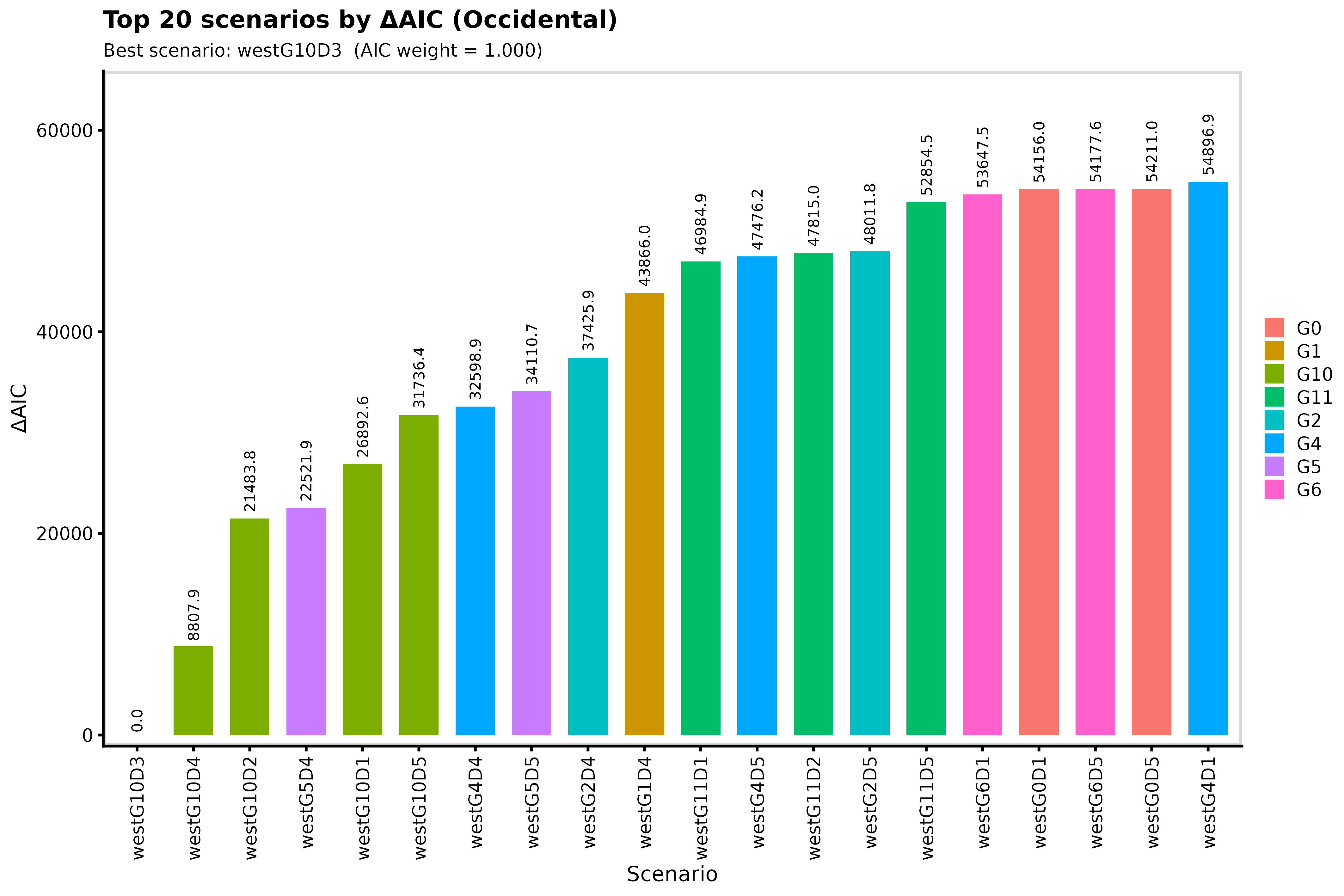

**Supplementary Figure 34 | Model selection support among the top 20 Occidental pear demographic scenarios based on Akaike information criterion.**

Scenarios are ranked by ΔAIC (each scenario's AIC minus the lowest AIC among all candidate scenarios tested for the Occidental lineage). Bars are colored by model group. The best-supported scenario has an Akaike weight of 1.000.

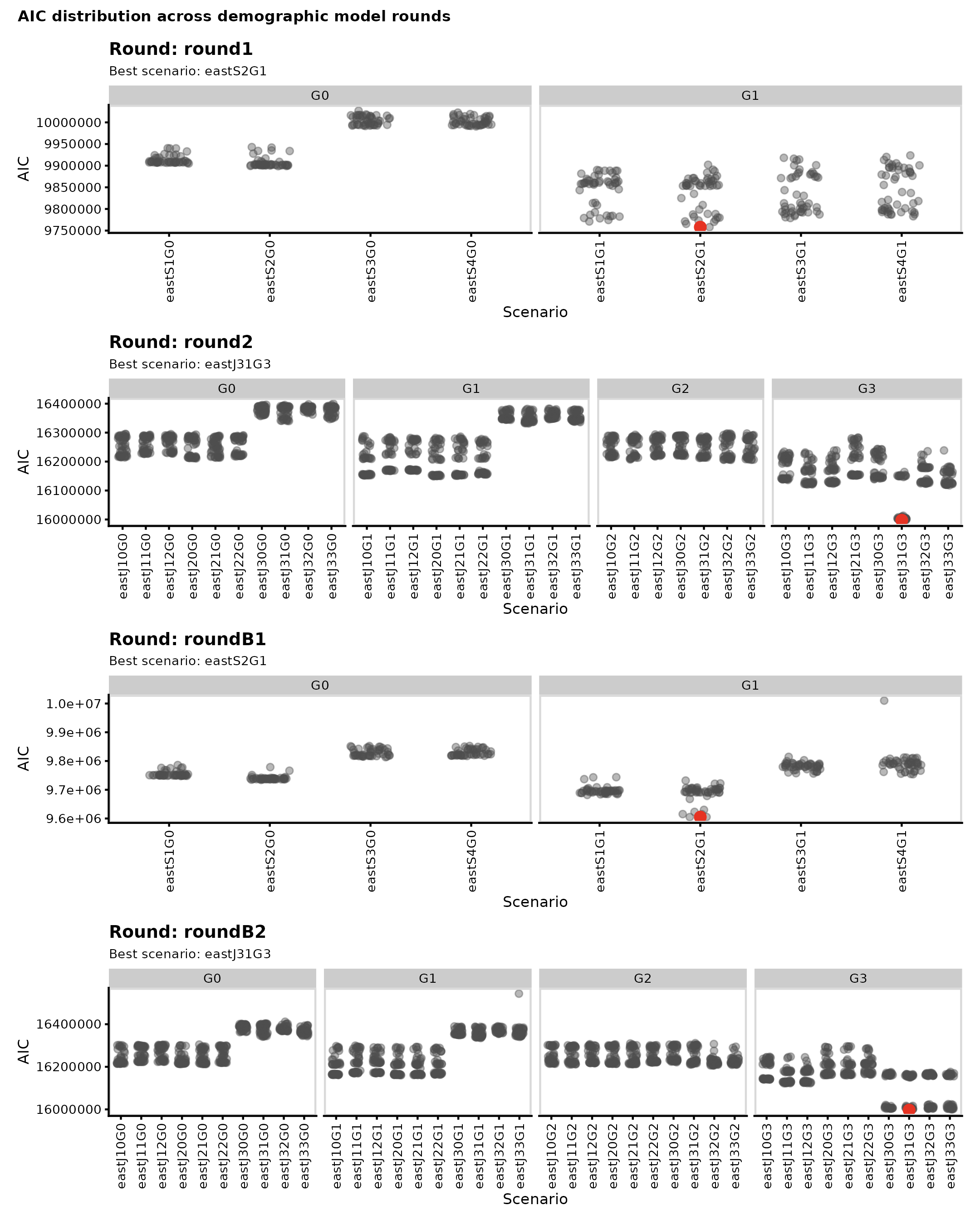

**Supplementary Figure 35 | AIC values for Oriental pear demographic models simulated with fastsimcoal2 (Rounds 1, 2, B1, and B2).**

Each point represents an independent run showing the best likelihood result obtained from 100,000 simulations per scenario, evaluated using the embedded MaxEstLhood function in fastsimcoal2. Each demographic scenario was replicated 50 times (50 × 100,000 simulations). The red point marks the best-fitting scenario with the lowest AIC value in each round. The names of the scenarios correspond to the demographic models illustrated in Supplementary Figure 31.

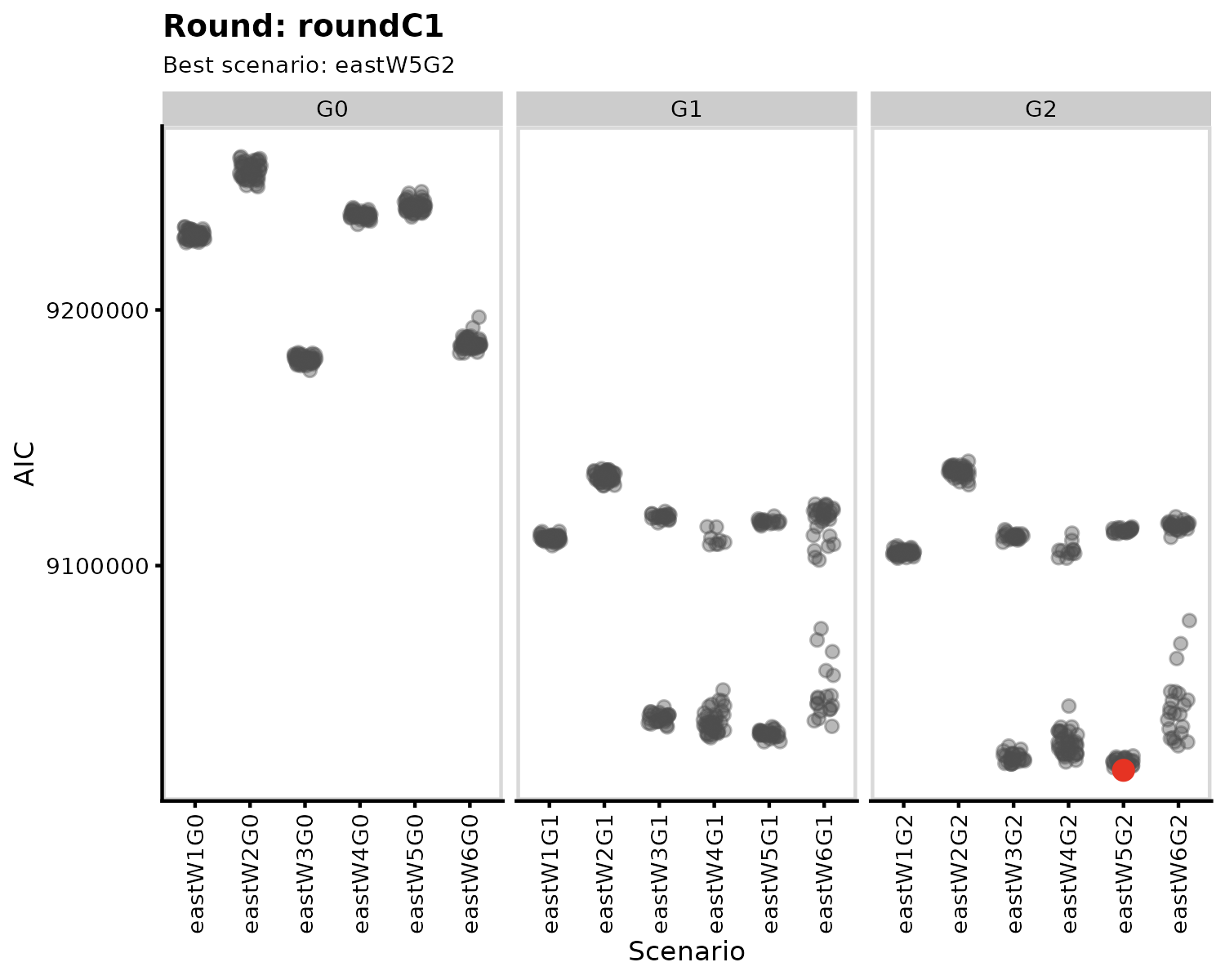

**Supplementary Figure 35 (continued) | AIC values for Oriental pear demographic models simulated with fastsimcoal2 (Round C1).**

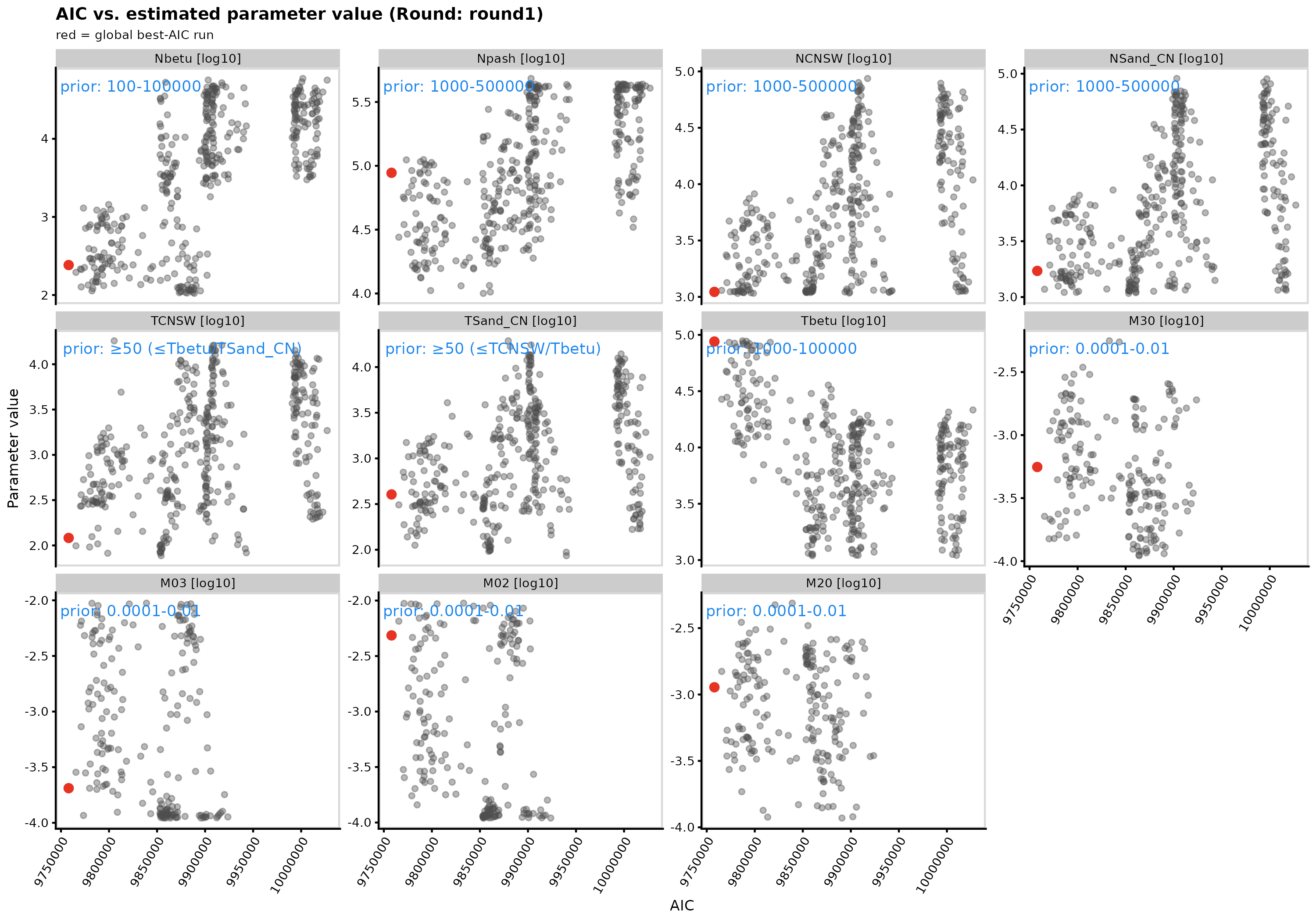

**Supplementary Figure 36 | Diagnostic relationship between the Akaike information criterion (AIC) and estimated parameter values across fastsimcoal2 runs for the Oriental pear dataset (Round 1).**

Each panel shows one demographic parameter (effective population size *N*, divergence time *T*, or migration rate *M*) plotted against the AIC of the run that produced it; values are log10-transformed for parameters spanning several orders of magnitude. Each point represents an independent run showing the best likelihood result obtained from 100,000 simulations, evaluated using the embedded MaxEstLhood function in fastsimcoal2. The blue text in each panel gives the prior range used for that parameter. Prior ranges are not shown for some divergence times (*T*), because their priors depend on the model topology and are therefore not fixed across scenarios. The red point marks the global best-AIC run across all scenarios and replicates in this round.

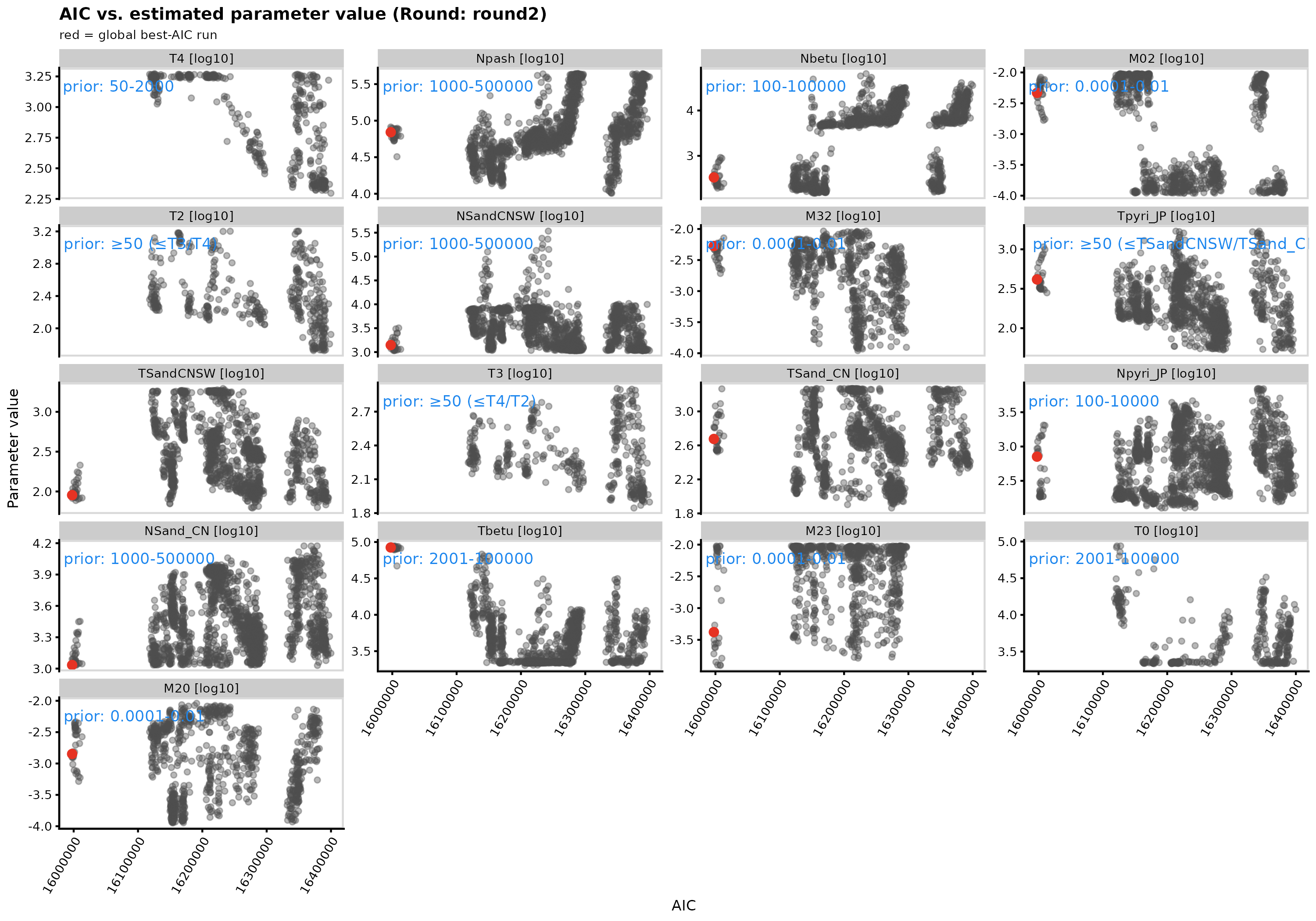

**Supplementary Figure 36 (continued) | Diagnostic relationship between the Akaike information criterion (AIC) and estimated parameter values across fastsimcoal2 runs for the Oriental pear dataset (round 2).**

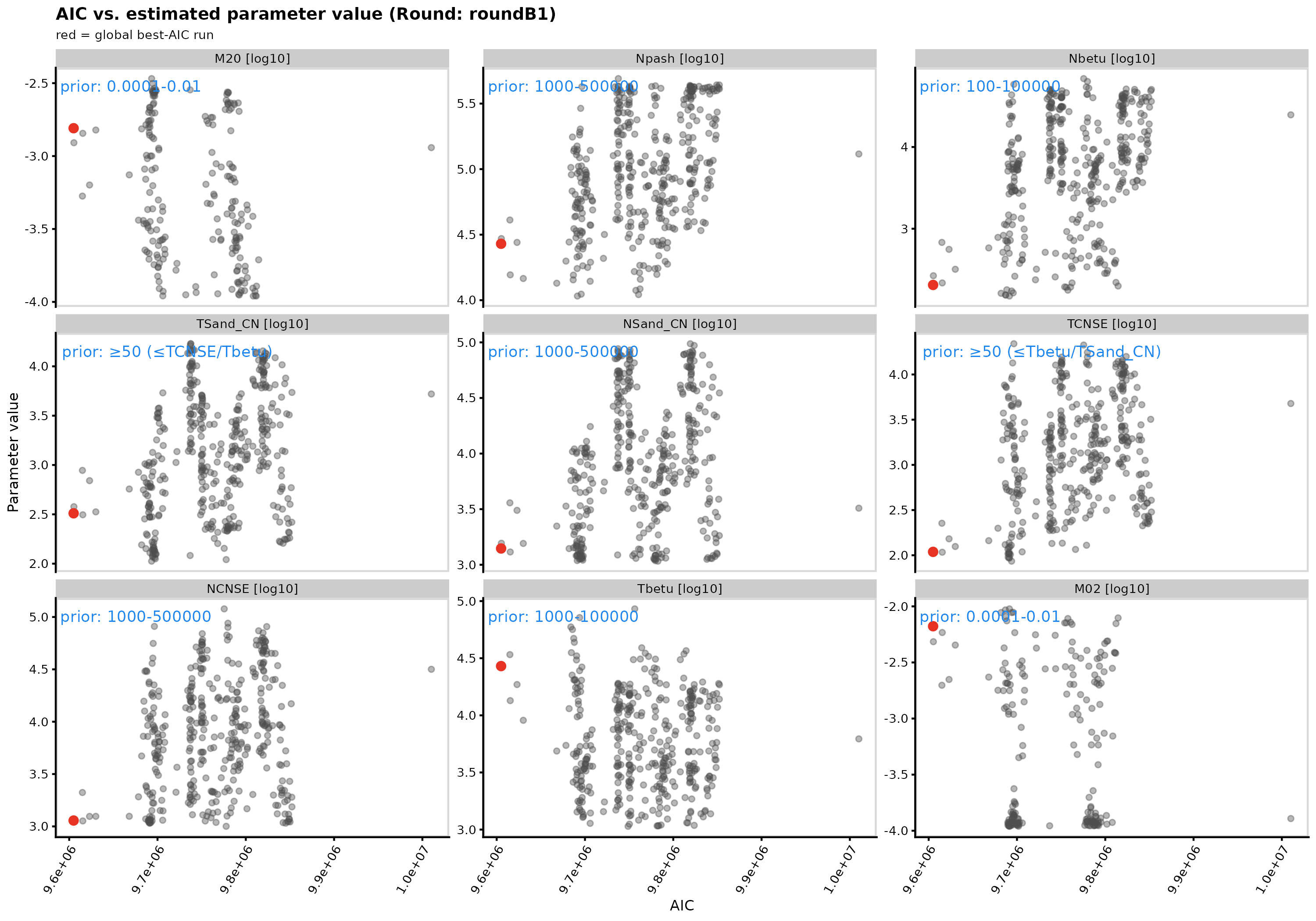

**Supplementary Figure 36 (continued) | Diagnostic relationship between the Akaike information criterion (AIC) and estimated parameter values across fastsimcoal2 runs for the Oriental pear dataset (round B1).**

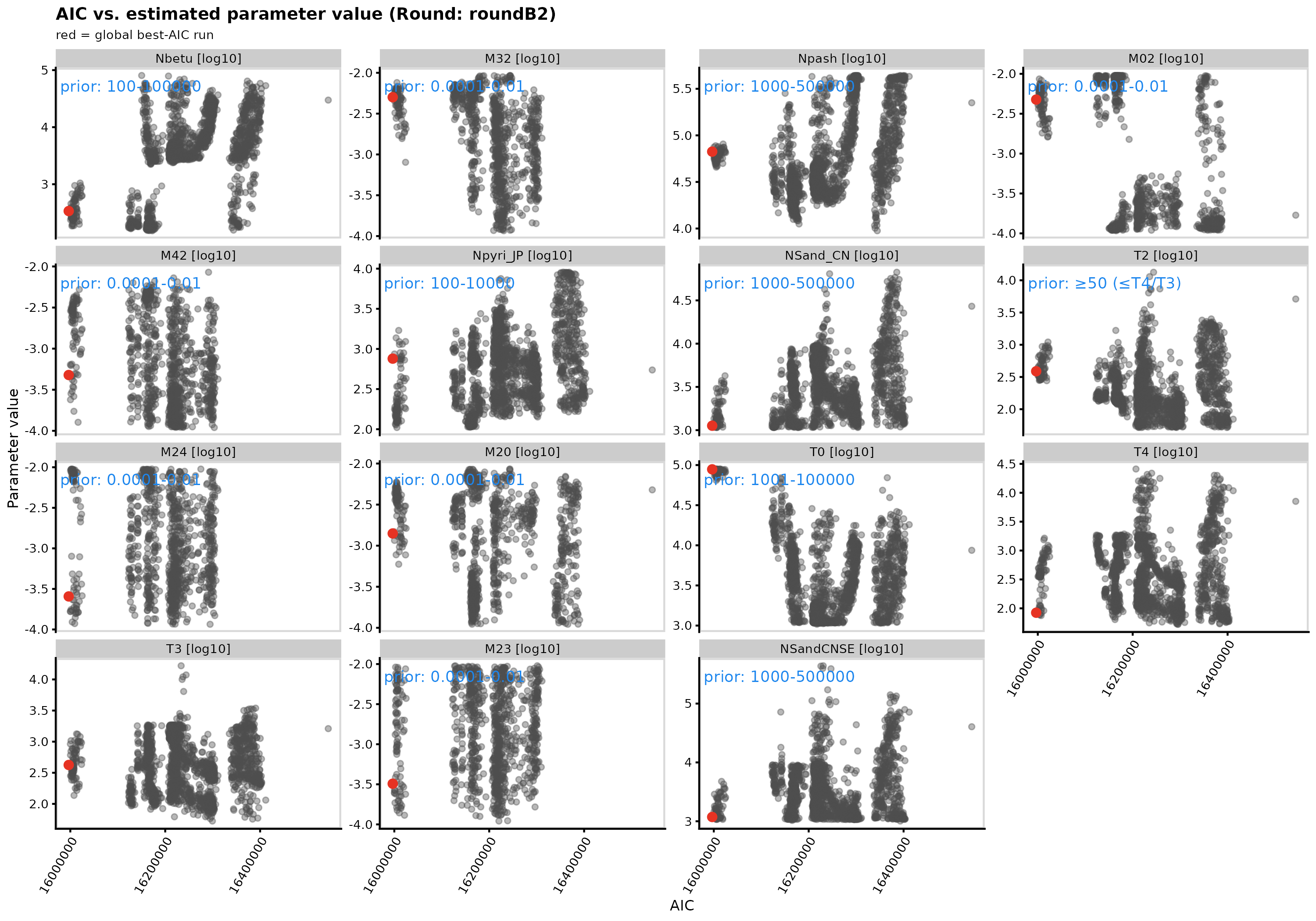

**Supplementary Figure 36 (continued) | Diagnostic relationship between the Akaike information criterion (AIC) and estimated parameter values across fastsimcoal2 runs for the Oriental pear dataset (round B2).**

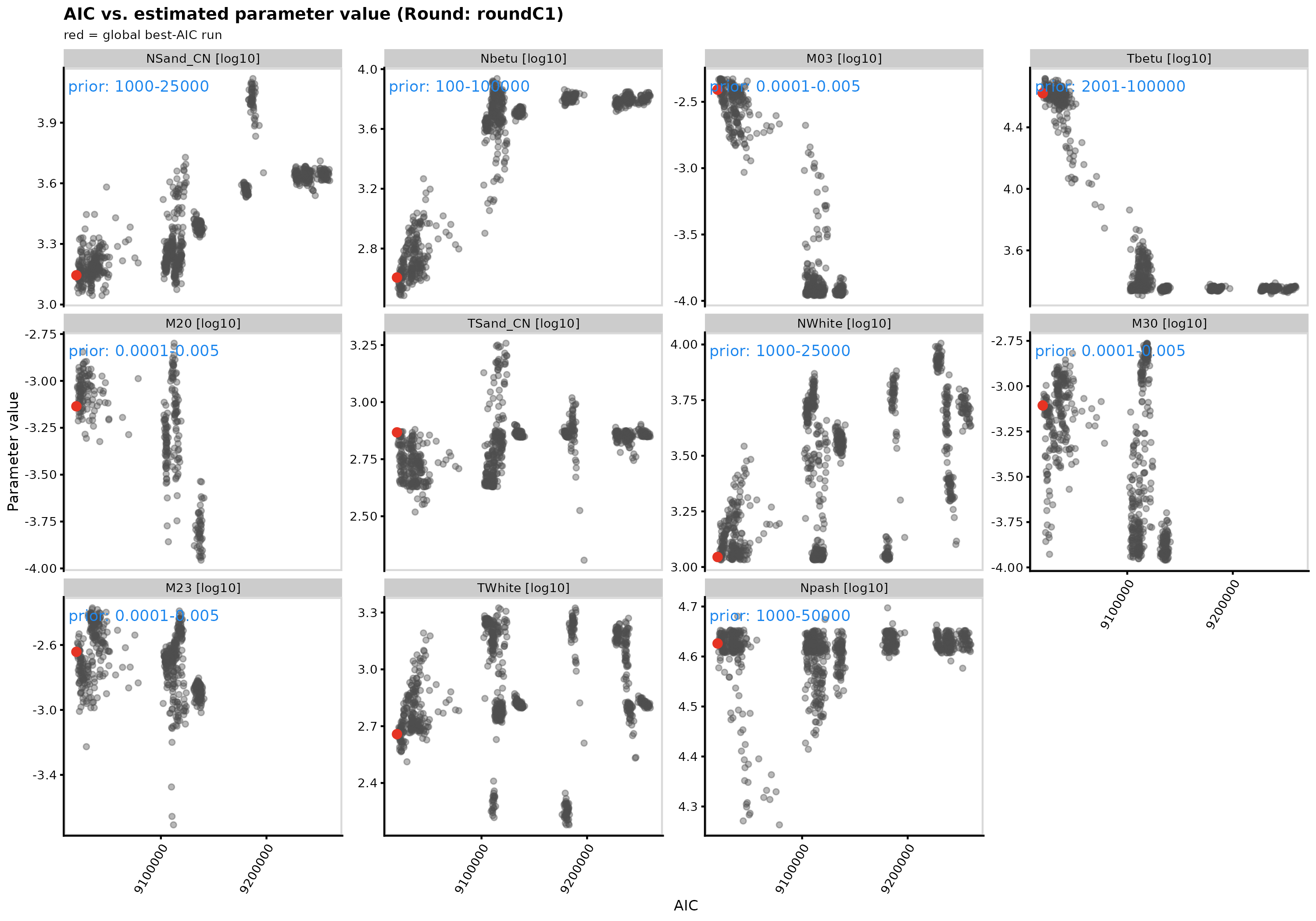

**Supplementary Figure 36 (continued) | Diagnostic relationship between the Akaike information criterion (AIC) and estimated parameter values across fastsimcoal2 runs for the Oriental pear dataset (round C1).**

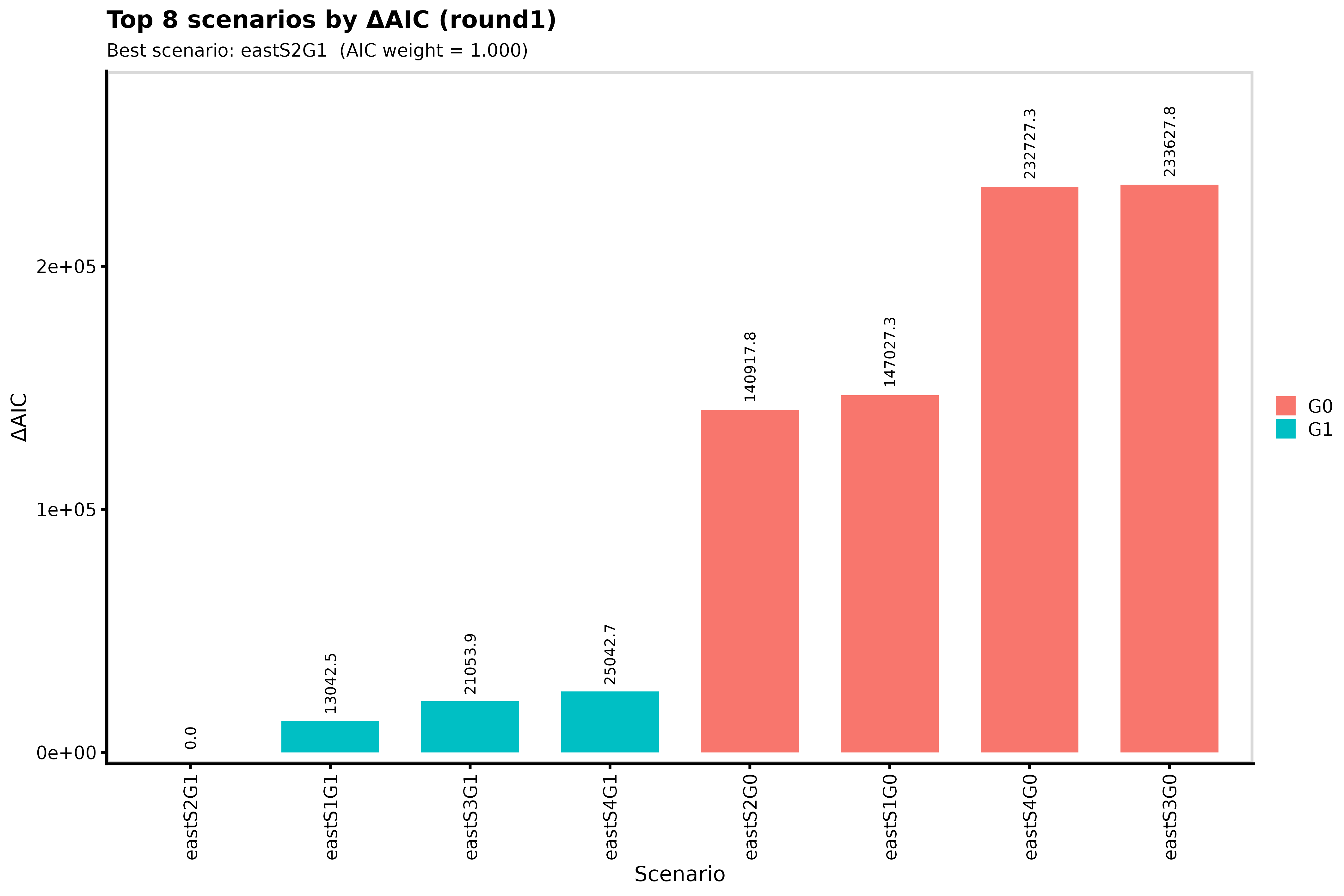

**Supplementary Figure 37 | Model selection support among the Oriental pear demographic scenarios based on Akaike information criterion (Round 1).**

Scenarios are ranked by ΔAIC (each scenario's AIC minus the lowest AIC among all candidate scenarios tested for the Oriental lineage). Bars are colored by model group. The best-supported scenario has an Akaike weight of 1.000.

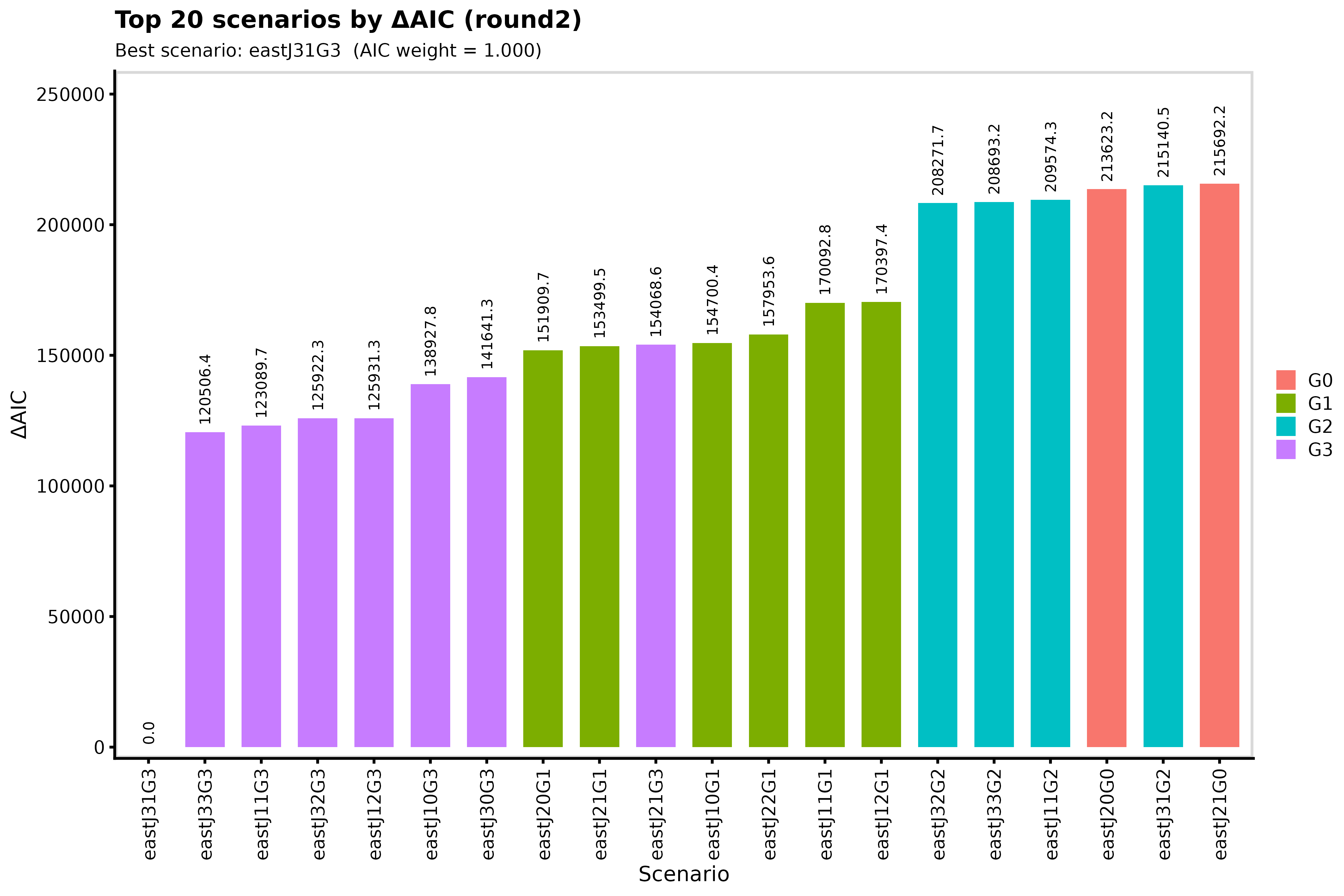

**Supplementary Figure 37 (continued) | Model selection support among the Oriental pear demographic scenarios based on Akaike information criterion (round 2).**

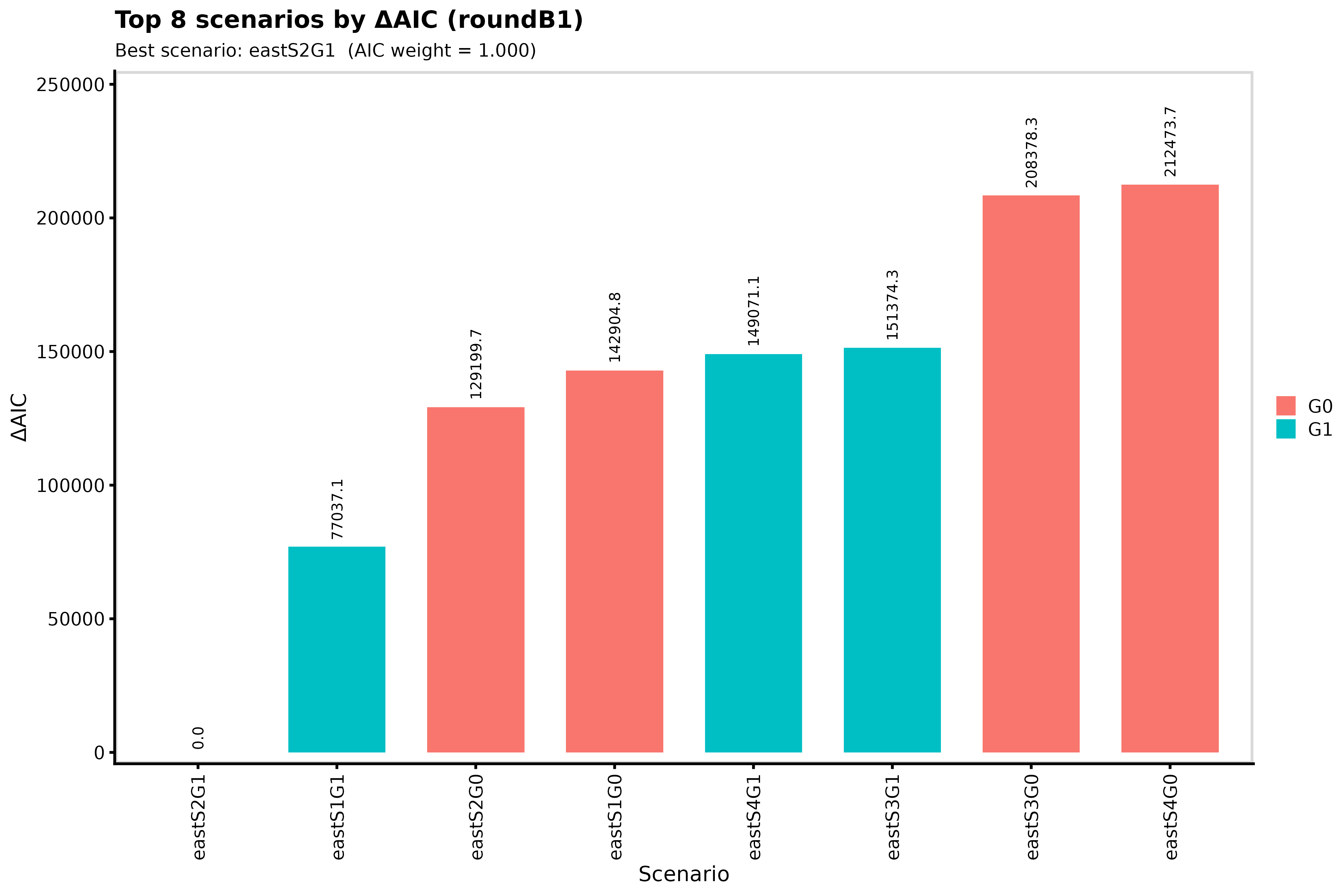

**Supplementary Figure 37 (continued) | Model selection support among the Oriental pear demographic scenarios based on Akaike information criterion (round B1).**

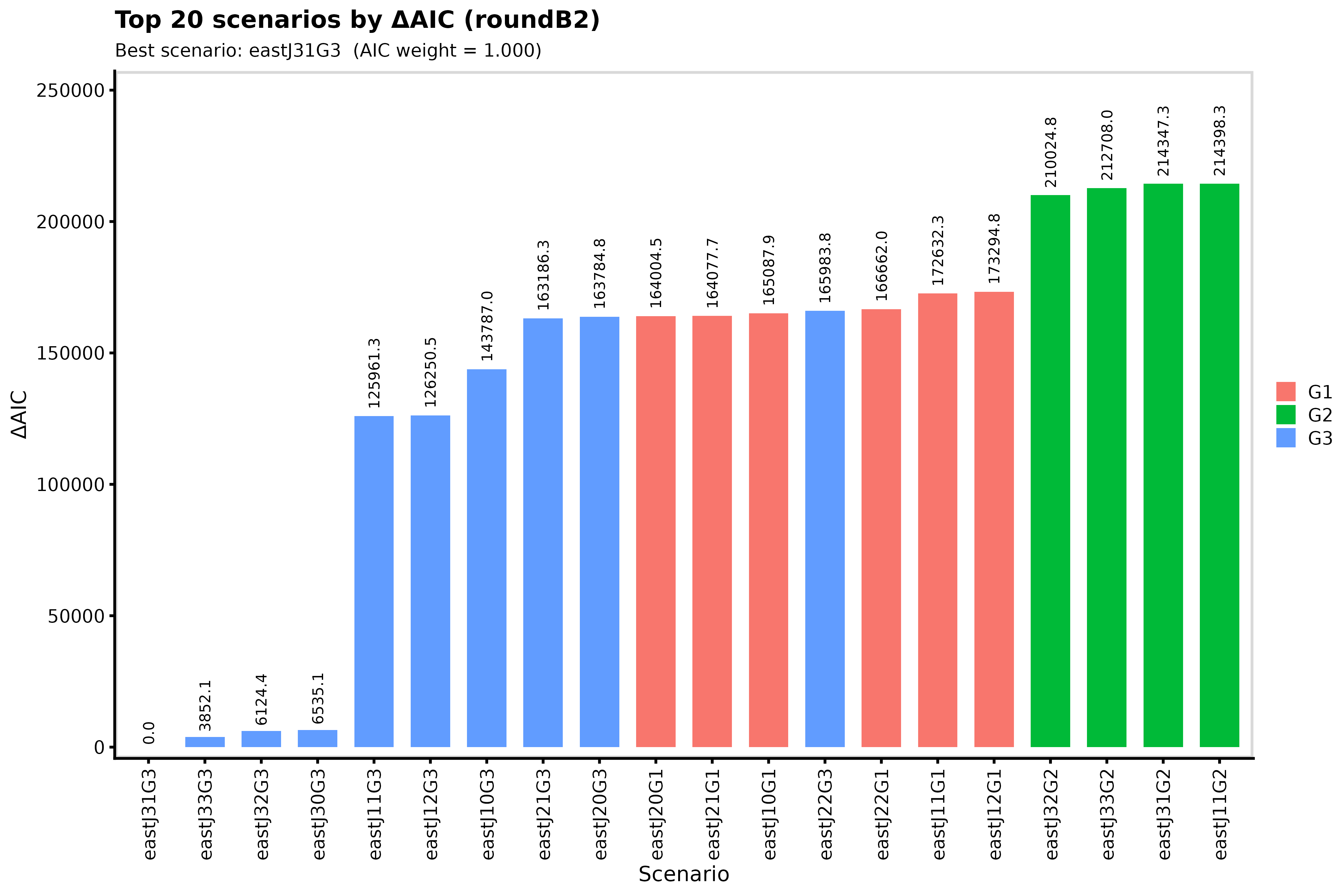

**Supplementary Figure 37 (continued) | Model selection support among the Oriental pear demographic scenarios based on Akaike information criterion (round B2).**

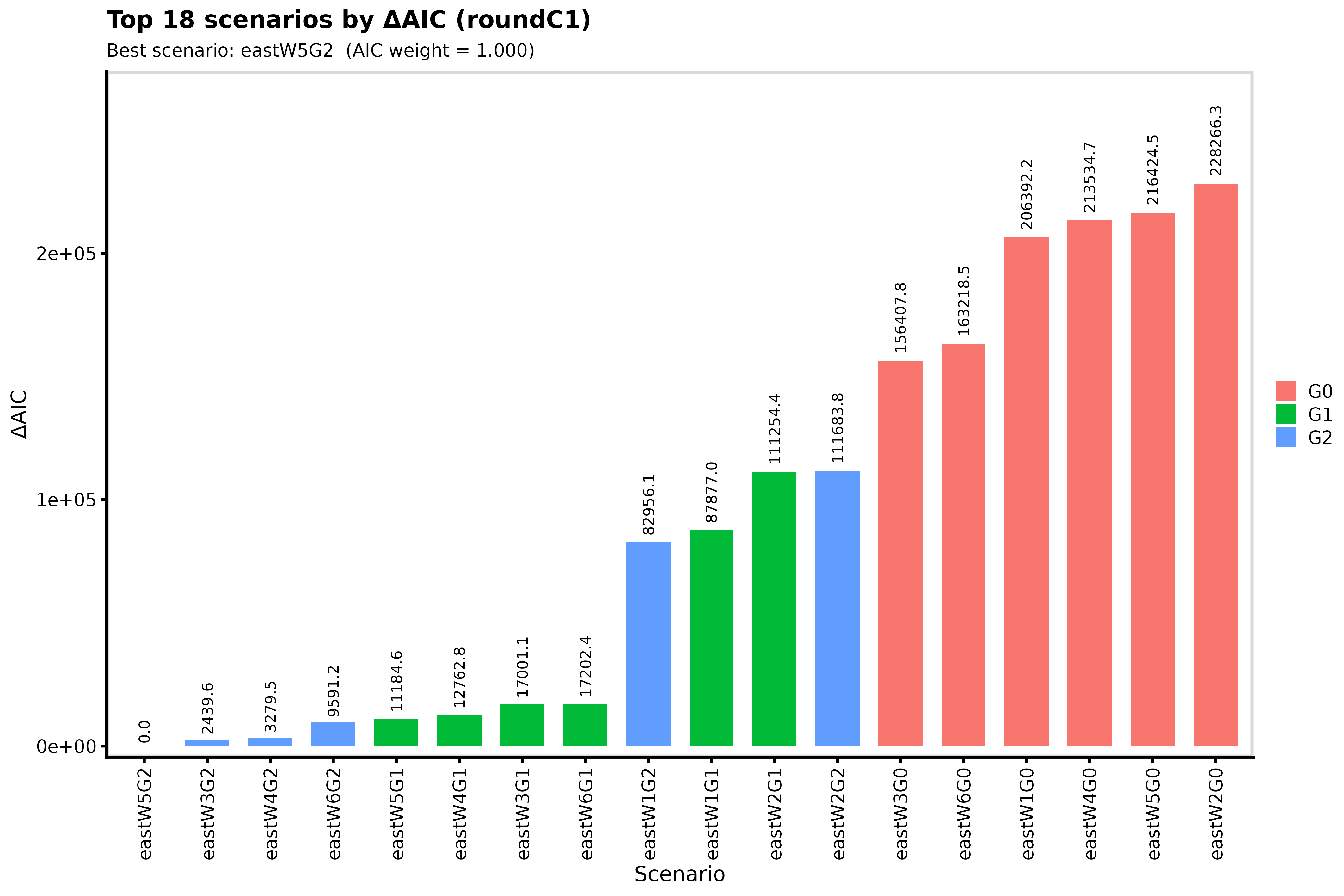

**Supplementary Figure 37 (continued) | Model selection support among the Oriental pear demographic scenarios based on Akaike information criterion (round C1).**

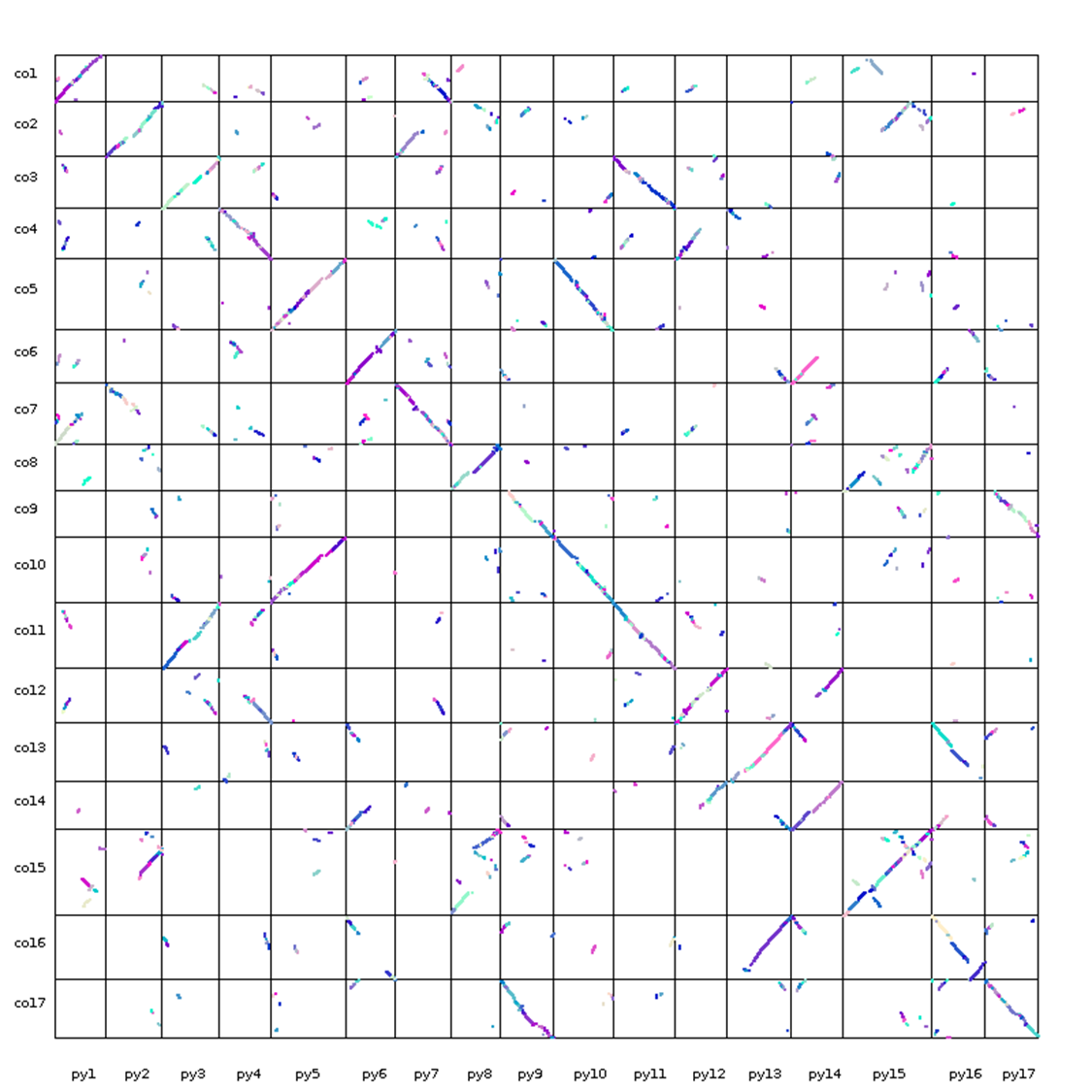

**Supplementary Figure 38 | Dot plot of syntenic genes between the two pear reference genomes.**

The analysis compared the *Pyrus communis* ‘Bartlett DH Genome v2.0’ (co) and *Pyrus pyrifolia* ‘Cuiguan’ (py) assemblies. Syntenic relationships were identified and visualized using MCScanX.

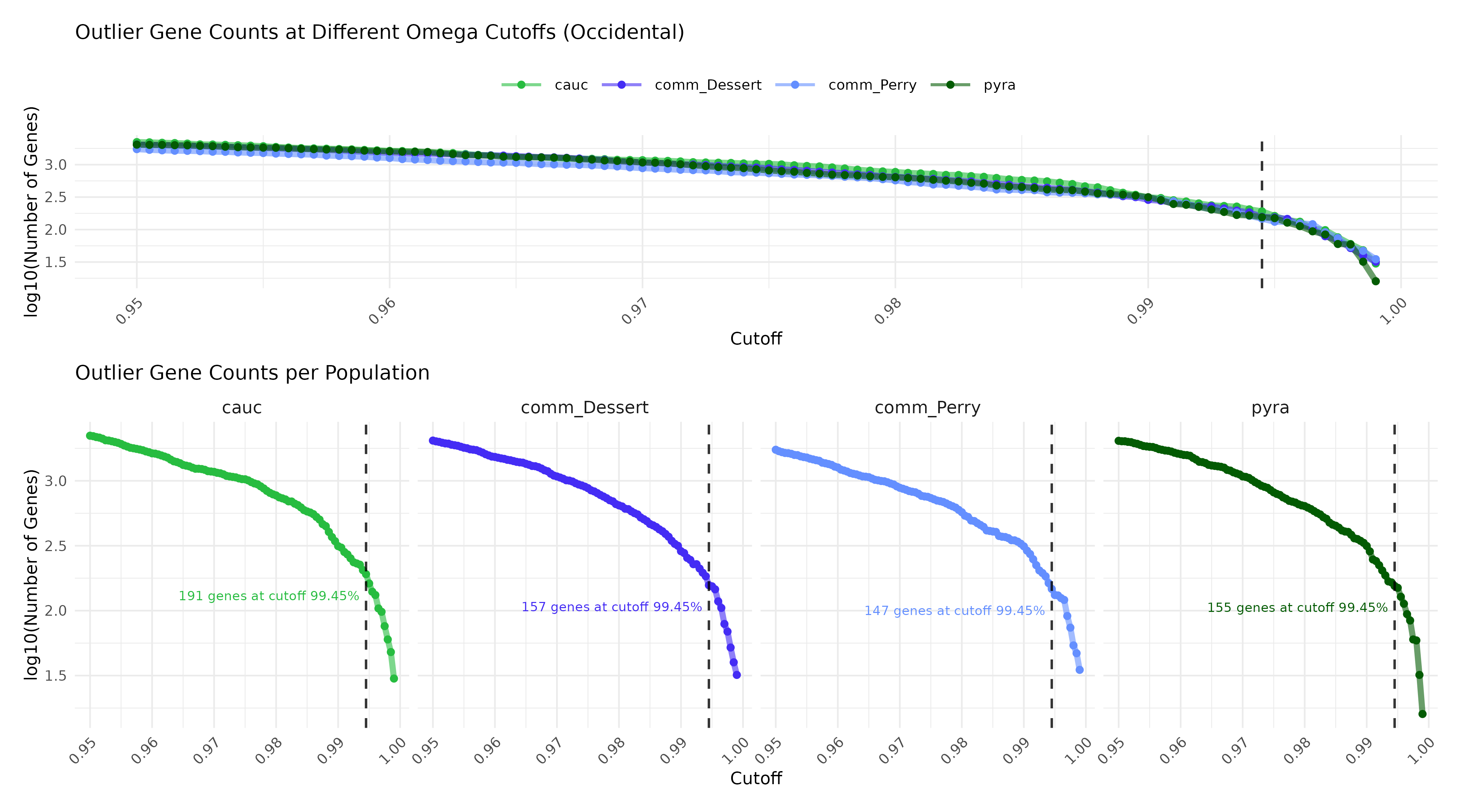

**Supplementary Figure 39 | Decay curves of candidate gene numbers captured at varying OmegaPlus score cutoffs in Occidental pear populations.**

The upper panel shows the log-transformed number of outlier genes detected across a range of percentile cutoffs (95–99.9%) for OmegaPlus scores (*ω*) in all Occidental populations using *P. communis* as the reference genome. The lower panels show population-specific decay curves, with the vertical dashed lines indicating the cutoff applied in the main analysis (99.45%). Labeled values represent the number of outlier genes identified at this threshold.

Population abbreviations:

cauc, *P. caucasica;*

comm Dessert, *P. communis* dessert;

comm Perry, *P. communis* perry;

pyra, *P. pyraster.*

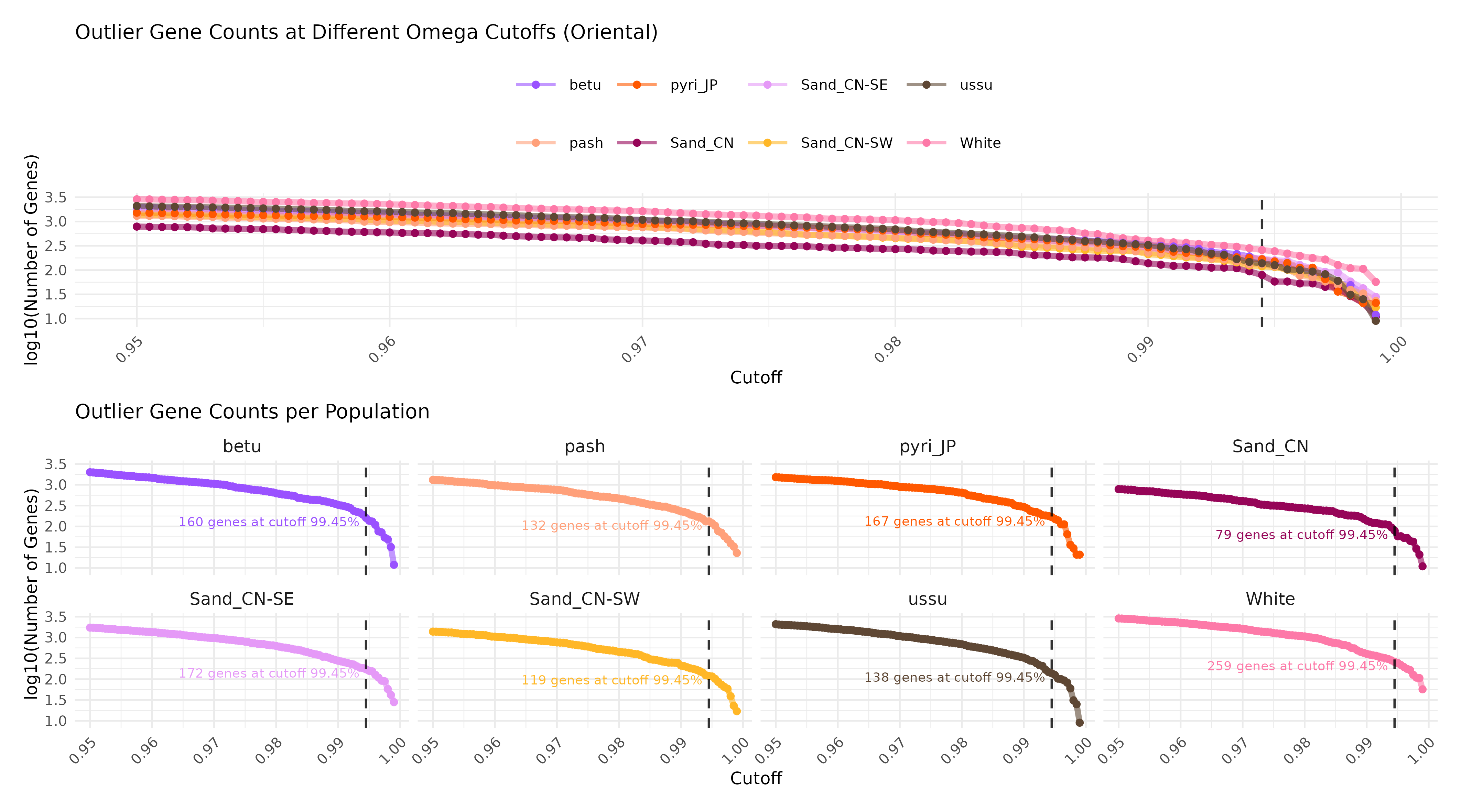

**Supplementary Figure 40 | Decay curves of candidate gene numbers captured at varying OmegaPlus score cutoffs in Oriental pear populations.**

The upper panel shows the log-transformed number of outlier genes detected across a range of percentile cutoffs (95–99.9%) for OmegaPlus scores (*ω*) in all Oriental populations, using *P. pyrifolia* as the reference genome. The lower panels show population-specific decay curves, with the vertical dashed line indicating the cutoff applied in the main analysis (99.45%). Labeled values represent the number of outlier genes identified at this threshold.

Population abbreviations:

betu, *Pyrus betulifolia*;

pash, *Pyrus pashia*;

pyri JP, *Pyrus pyrifolia* Nashi Japan;

Sand CN, *Pyrus pyrifolia* sand China-other;

Sand CN-SE, *Pyrus pyrifolia* sand China-Southeast;

Sand CN-SW, *Pyrus pyrifolia* sand China-Southwest;

ussu, *Pyrus ussuriensis*;

White, *Pyrus pyrifolia* white China.

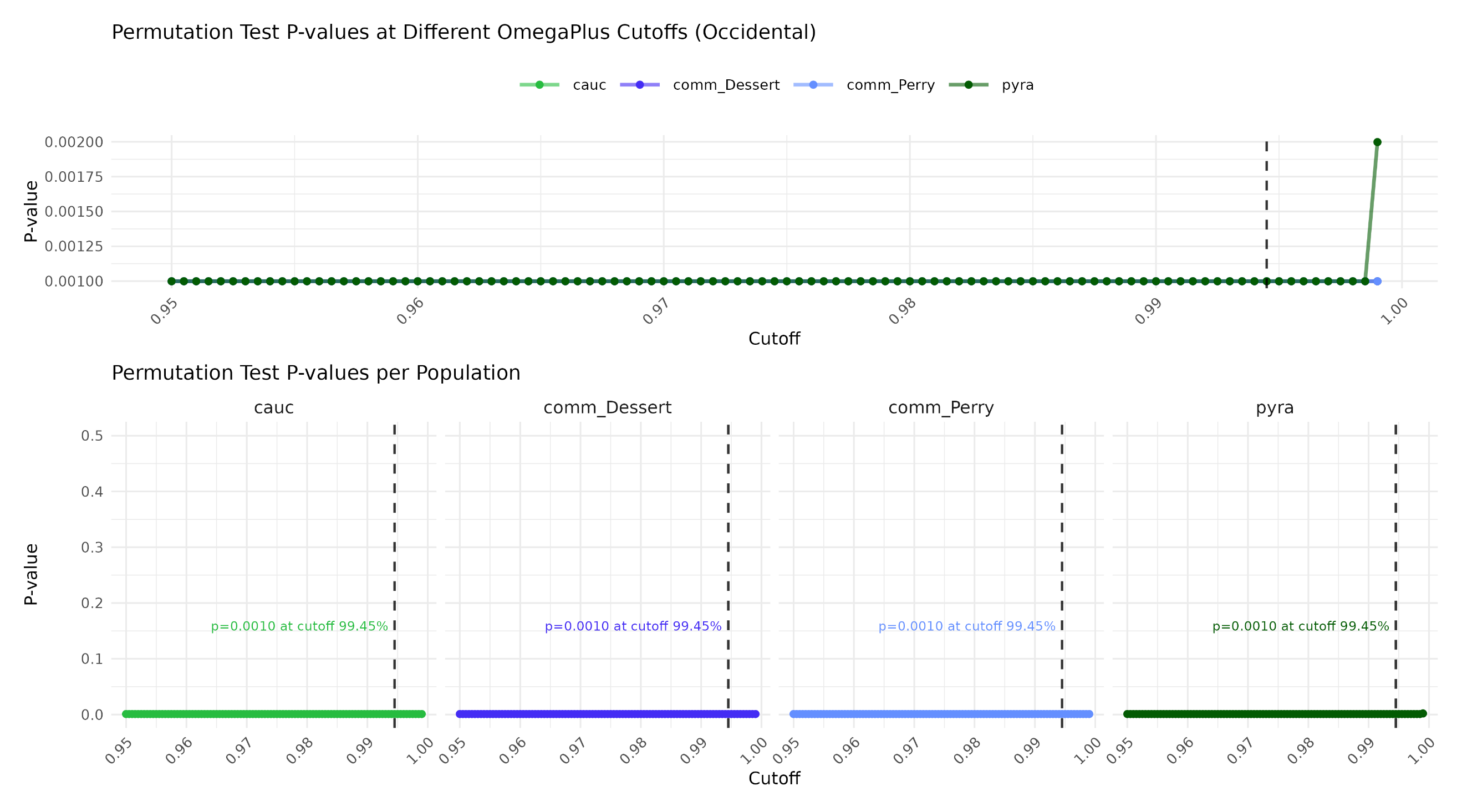

**Supplementary Figure 41 | Permutation** **test *p*-values for outlier gene sets identified at varying OmegaPlus cutoffs in Occidental pear populations.**

The upper panel shows permutation test *p-*values estimated across a range of OmegaPlus percentile cutoffs (95–99.9%) for Occidental populations, using *P. communis* as the reference genome. The lower panels show population-specific results, with the vertical dashed lines indicating the main analysis threshold (99.45%) and the corresponding *p*-value, *p* = 0.001. The *p*-values were calculated using the randomizeRegions method with the regioneR package to assess the statistical significance of the observed overlap between outlier gene regions and the expected value of a random region.

Population abbreviations:

cauc, *P. caucasica;*

comm Dessert, *P. communis* dessert;

comm Perry, *P. communis* perry;

pyra, *P. pyraster.*

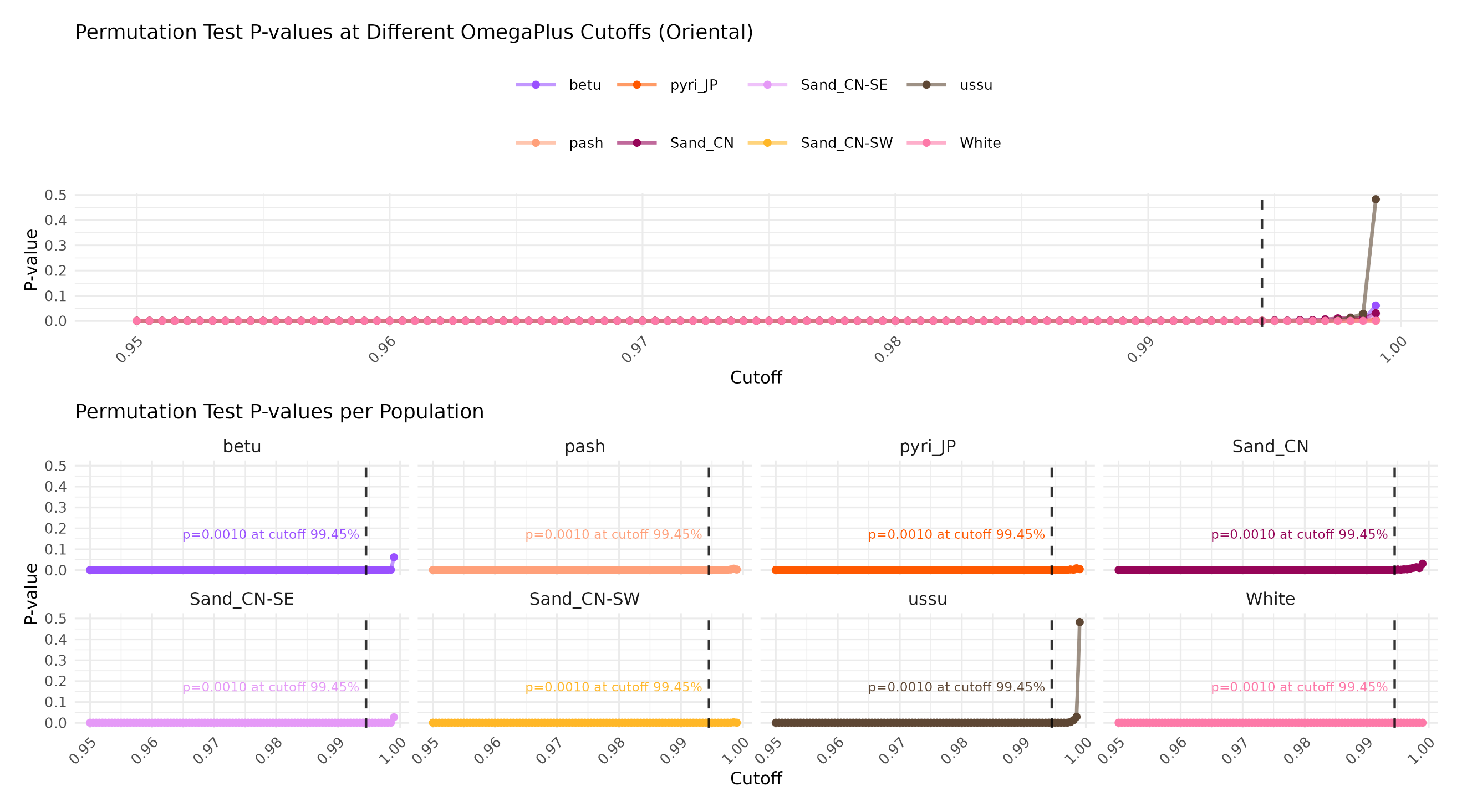

**Supplementary Figure 42 | Permutation test *p*-values for outlier gene sets identified at varying OmegaPlus cutoffs in Oriental pear populations.**

The upper panel shows permutation test *p*-values estimated across a range of OmegaPlus percentile cutoffs (95–99.9%) for Oriental populations, using *P. pyrifolia* as the reference genome. The lower panels show population-specific results, with the vertical dashed lines indicating the main analysis threshold (99.45%) and the corresponding *p*-value, *p* = 0.001. *P*-values were calculated using the randomizeRegions method implemented in the regioneR package to assess the statistical significance of the observed overlap between outlier gene regions and the expected value of a random region.

Population abbreviations:

betu, *Pyrus betulifolia*;

pash, *Pyrus pashia*;

pyri JP, *Pyrus pyrifolia* Nashi Japan;

Sand CN, *Pyrus pyrifolia* sand China-other;

Sand CN-SE, *Pyrus pyrifolia* sand China-Southeast;

Sand CN-SW, *Pyrus pyrifolia* sand China-Southwest;

ussu, *Pyrus ussuriensis*;

White, *Pyrus pyrifolia* white China.

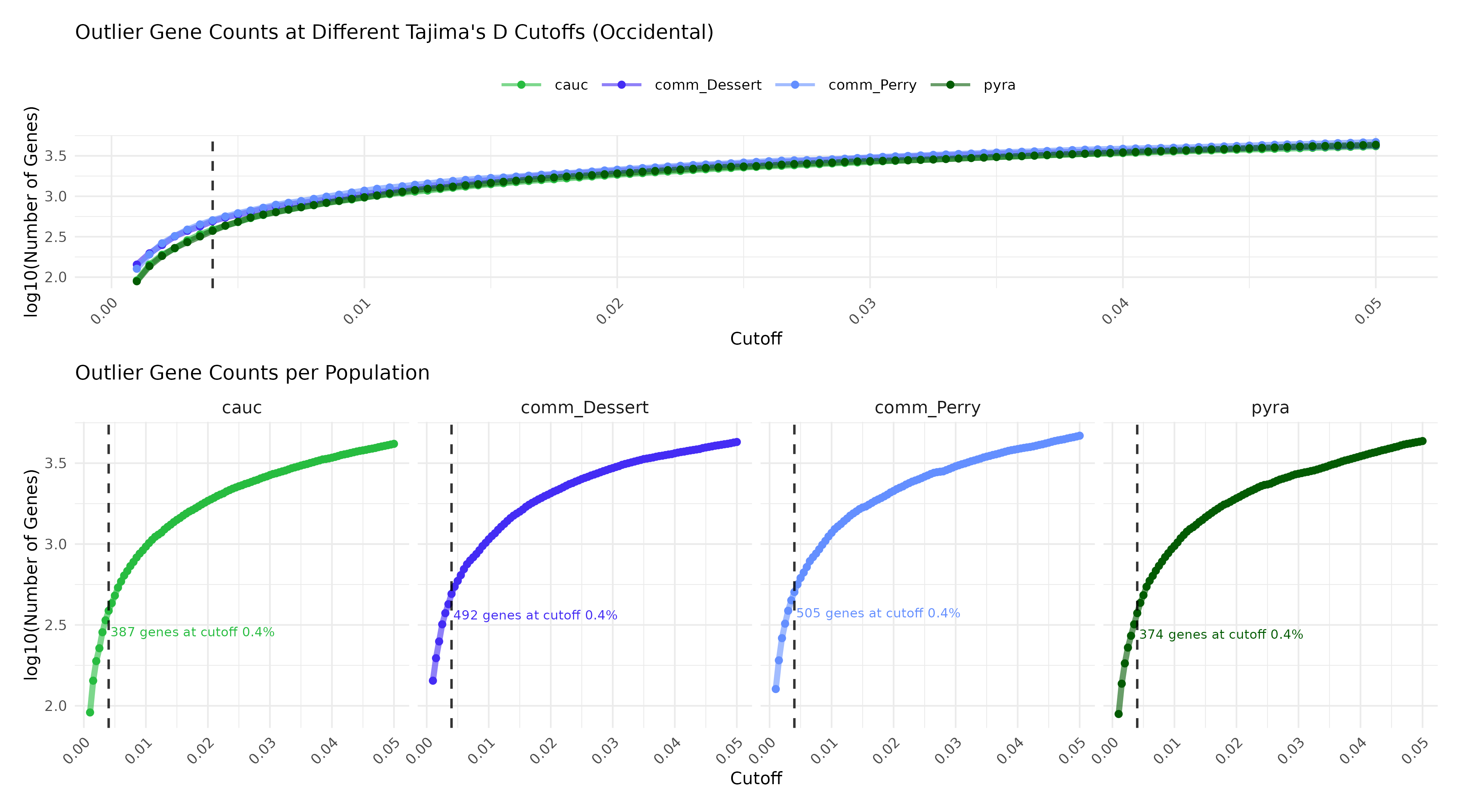

**Supplementary Figure 43 | Decay curves of candidate gene numbers captured at Tajima’s *D* score cutoffs in Occidental pear populations.**

The upper panel shows the log-transformed number of outlier genes detected across a range of percentile cutoffs (0.1–5%) for unbiased Tajima’s *D* scores (pixy) in all Occidental populations, using *P. communis* as the reference genome. The lower panels show population-specific decay curves, with the vertical dashed lines indicating the cutoff applied in the main analysis (0.4%). Labeled values represent the number of outlier genes identified at this threshold.

Population abbreviations:

cauc, *P. caucasica;*

comm Dessert, *P. communis* dessert;

comm Perry, *P. communis* perry;

pyra, *P. pyraster.*

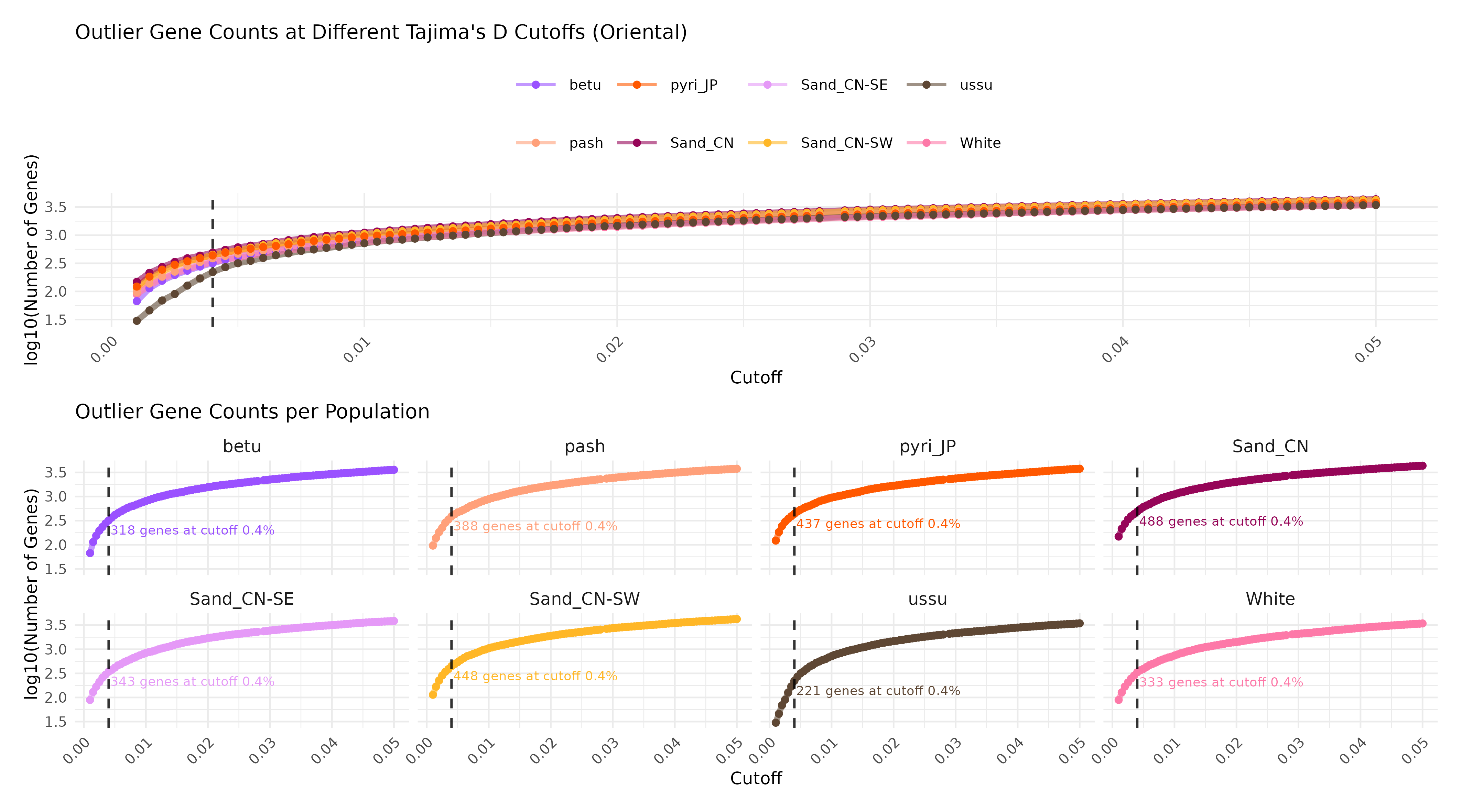

**Supplementary Figure 44 | Decay curves of candidate gene numbers captured at Tajima’s *D* score cutoffs in Oriental pear populations.**

The upper panel shows the log-transformed number of outlier genes detected across a range of percentile cutoffs (0.1–5%) for unbiased Tajima’s *D* scores (pixy) in all Oriental populations, using *P. pyrifolia* as the reference genome. The lower panels show population-specific decay curves, with the vertical dashed lines indicating the cutoff applied in the main analysis (0.4%). Labeled values represent the number of outlier genes identified at this threshold.

Population abbreviations:

betu, *Pyrus betulifolia*;

pash, *Pyrus pashia*;

pyri JP, *Pyrus pyrifolia* Nashi Japan;

Sand CN, *Pyrus pyrifolia* sand China-other;

Sand CN-SE, *Pyrus pyrifolia* sand China-Southeast;

Sand CN-SW, *Pyrus pyrifolia* sand China-Southwest;

ussu, *Pyrus ussuriensis*;

White, *Pyrus pyrifolia* white China.

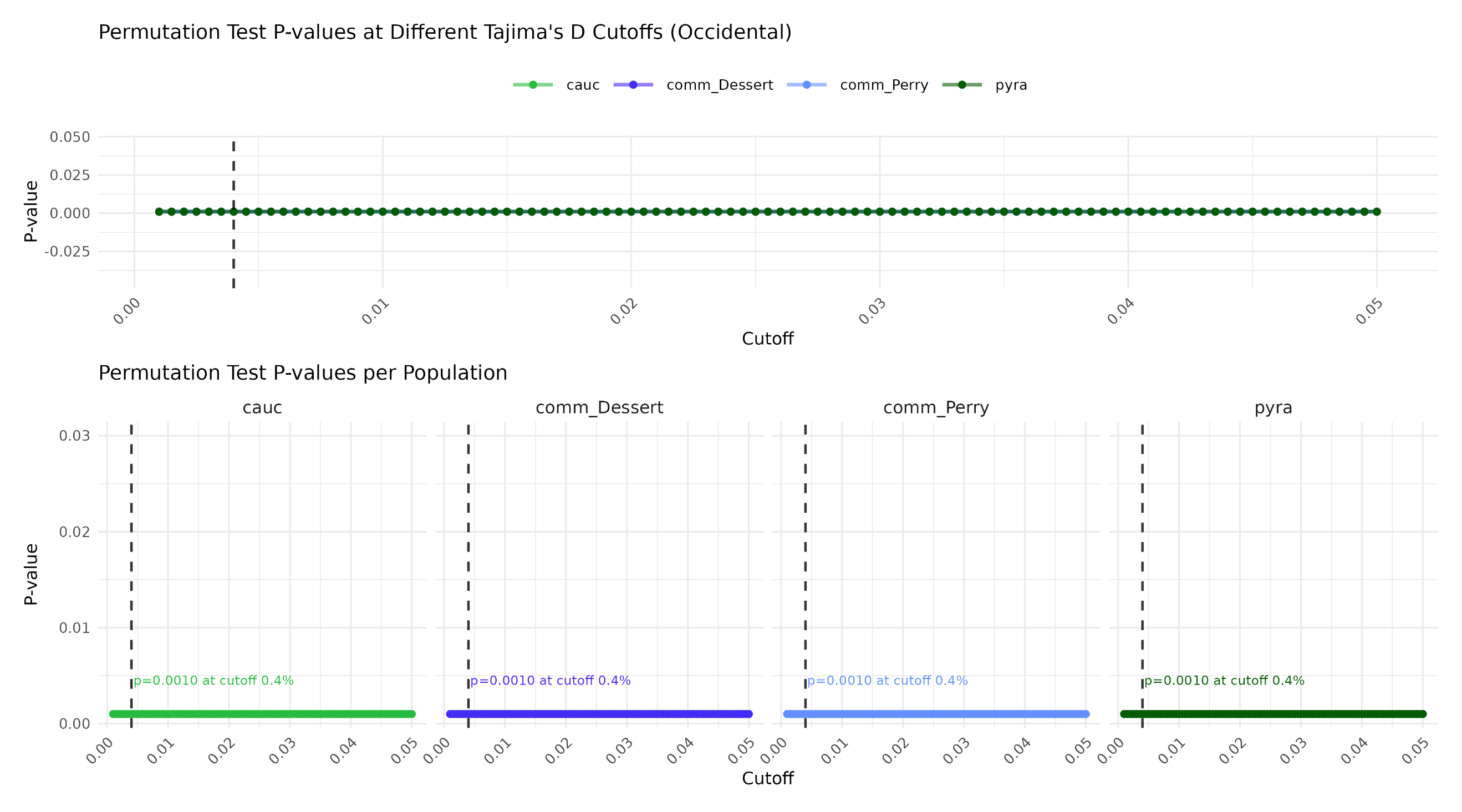

**Supplementary Figure 45 | Permutation test *p*-values for outlier gene sets identified at varying Tajima’s *D* cutoffs in Occidental pear populations.**

The upper panel shows permutation test *p*-values estimated across a range of Tajima’s *D* percentile cutoffs (0.1–5%) for all Occidental populations. The lower panels show population-specific results, with the vertical dashed lines indicating the main analysis threshold (0.4%) and the corresponding *p*-value, *p* = 0.001. The *p*-values were calculated using the randomizeRegions method with the regioneR package to assess the statistical significance of the observed overlap between outlier gene regions and the expected value of a random region.

Population abbreviations:

cauc, *P. caucasica;*

comm Dessert, *P. communis* dessert;

comm Perry, *P. communis* perry;

pyra, *P. pyraster.*

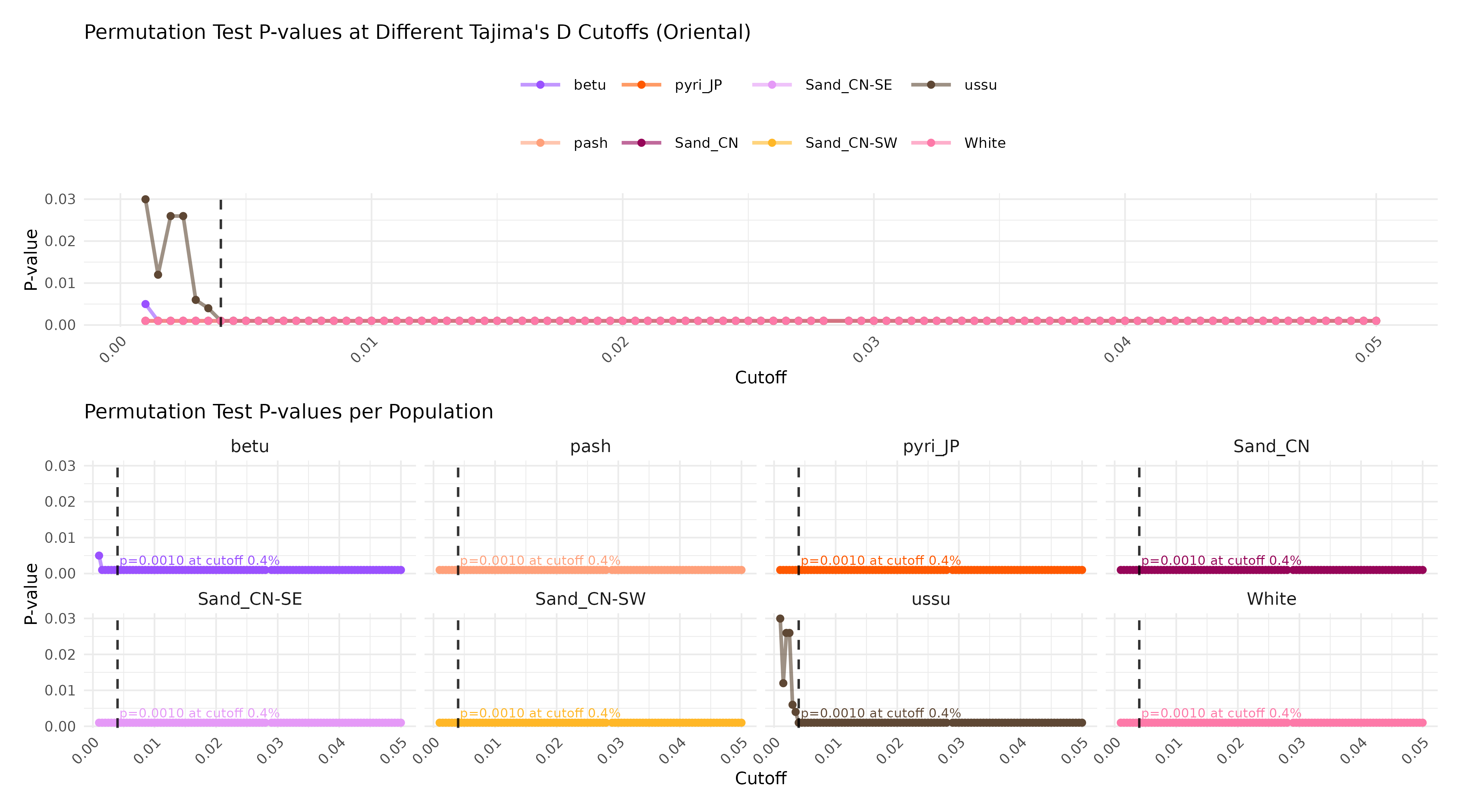

**Supplementary Figure 46 | Permutation test *p*-values for outlier gene sets identified at varying Tajima’s *D* cutoffs in Oriental pear populations.**

The upper panel shows permutation test *p*-values estimated across a range of Tajima’s *D* percentile cutoffs (0.1–5%) for all Oriental populations, using *P. pyrifolia* as the reference genome. The lower panels show population-specific results, with the vertical dashed lines indicating the main analysis threshold (0.4%) and the corresponding *p*-value, *p* = 0.001. The *p-*values were calculated using the ‘randomizeRegions’ method with the regioneR package to assess the statistical significance of the observed overlap between outlier gene regions and the expected value of a random region.

Population abbreviations:

betu, *Pyrus betulifolia*;

pash, *Pyrus pashia*;

pyri JP, *Pyrus pyrifolia* Nashi Japan;

Sand CN, *Pyrus pyrifolia* sand China-other;

Sand CN-SE, *Pyrus pyrifolia* sand China-Southeast;

Sand CN-SW, *Pyrus pyrifolia* sand China-Southwest;

ussu, *Pyrus ussuriensis*;

White, *Pyrus pyrifolia* white China.

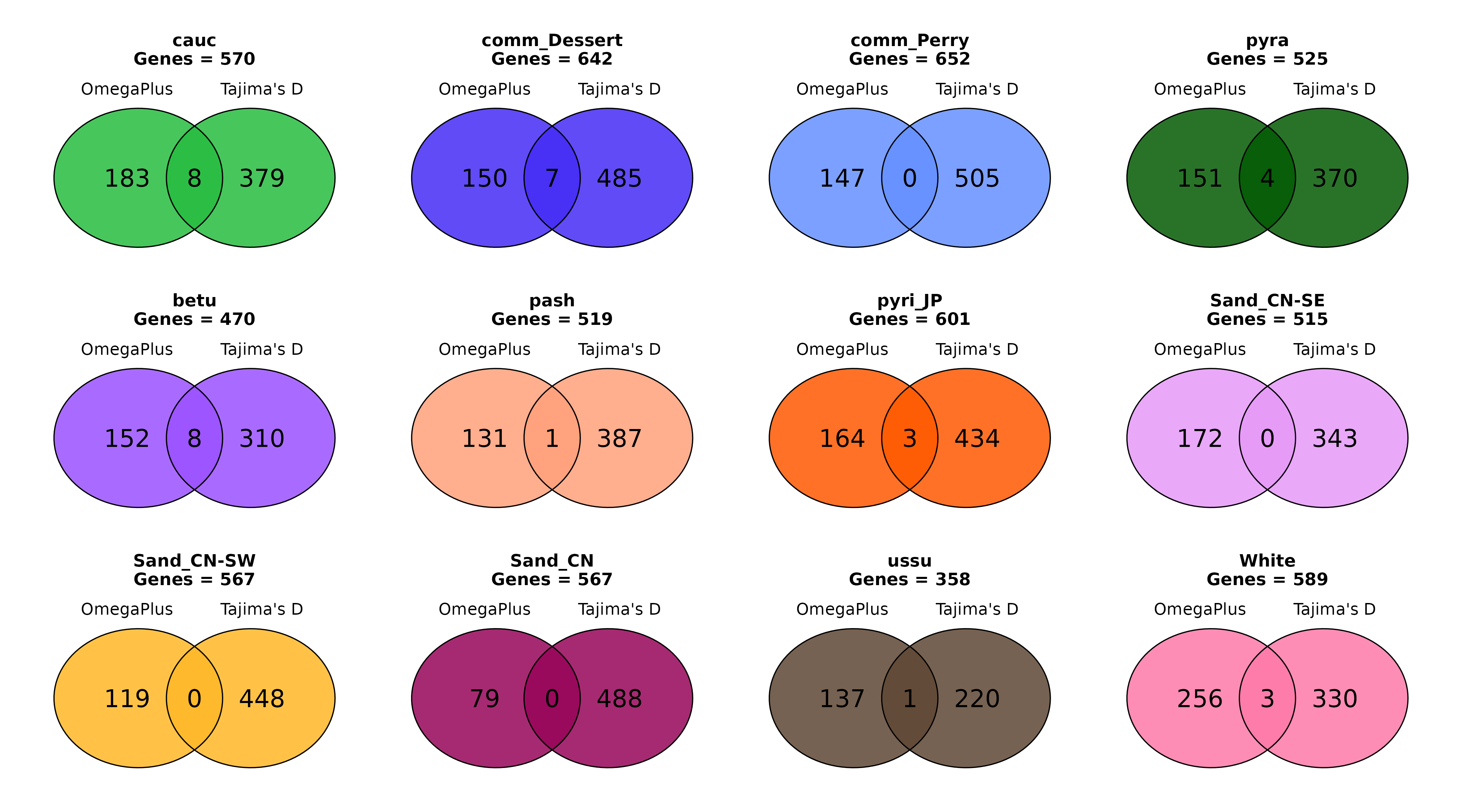

**Supplementary Figure 47 | Extent of overlap between candidate genes under selection in the 12 pear populations detected using two complementary methods.**

Venn diagrams show the number of candidate genes identified for selective sweeps in each population using OmegaPlus and unbiased Tajima’s *D* (calculated via pixy). The intersection indicates genes detected by both methods.

Population abbreviations:

cauc, *P. caucasica;*

comm Dessert, *P. communis* dessert;

comm Perry, *P. communis* perry;

pyra, *P. pyraster*;

betu, *Pyrus betulifolia*;

pash, *Pyrus pashia*;

pyri JP, *Pyrus pyrifolia* Nashi Japan;

Sand CN, *Pyrus pyrifolia* sand China-other;

Sand CN-SE, *Pyrus pyrifolia* sand China-Southeast;

Sand CN-SW, *Pyrus pyrifolia* sand China-Southwest;

ussu, *Pyrus ussuriensis*;

White, *Pyrus pyrifolia* white China.

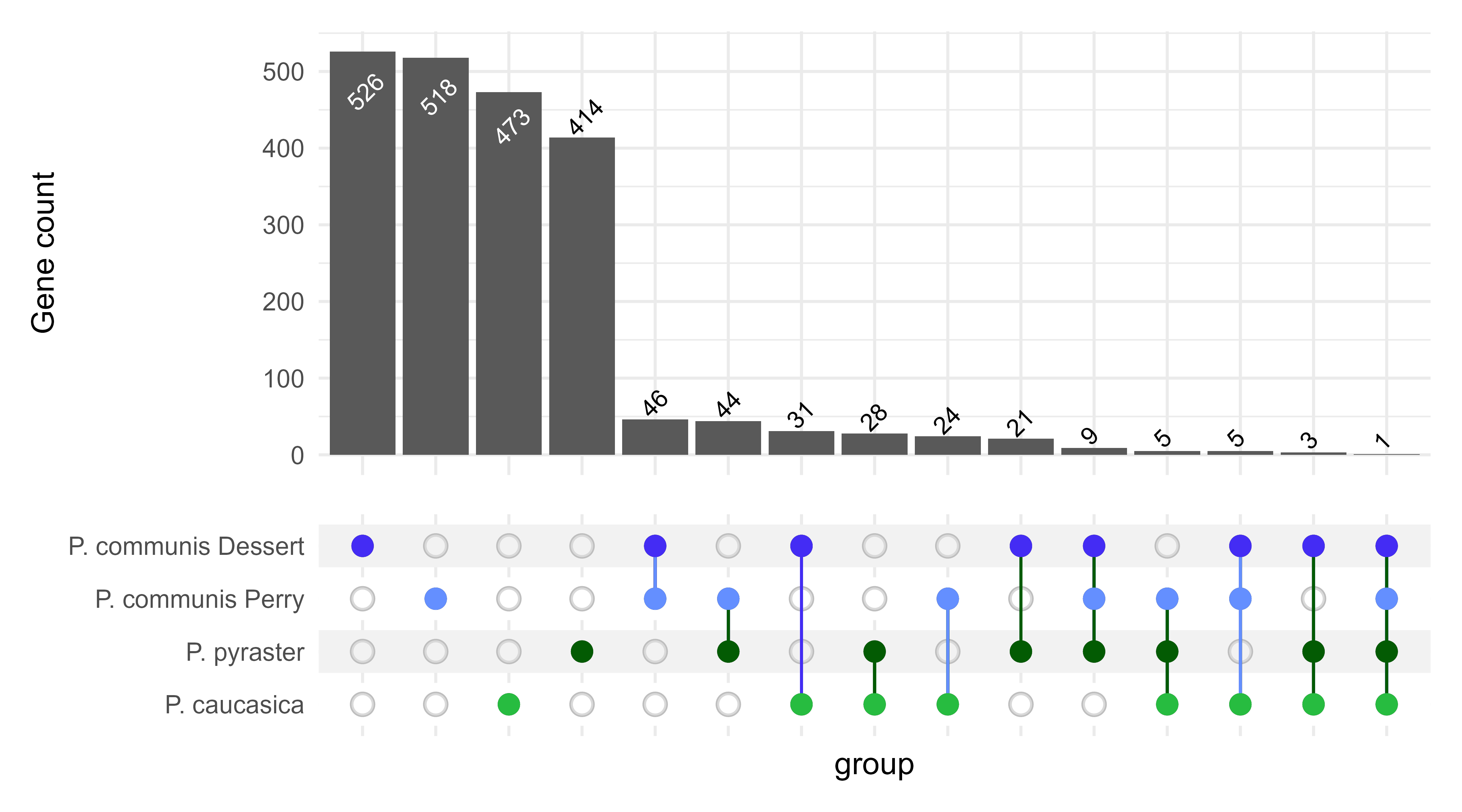

**Supplementary Figure 48 | Genes of interest in selective sweep regions in Occidental pear populations.**

The candidate genes were identified using Tajima’s *D* and *ω* statistics.

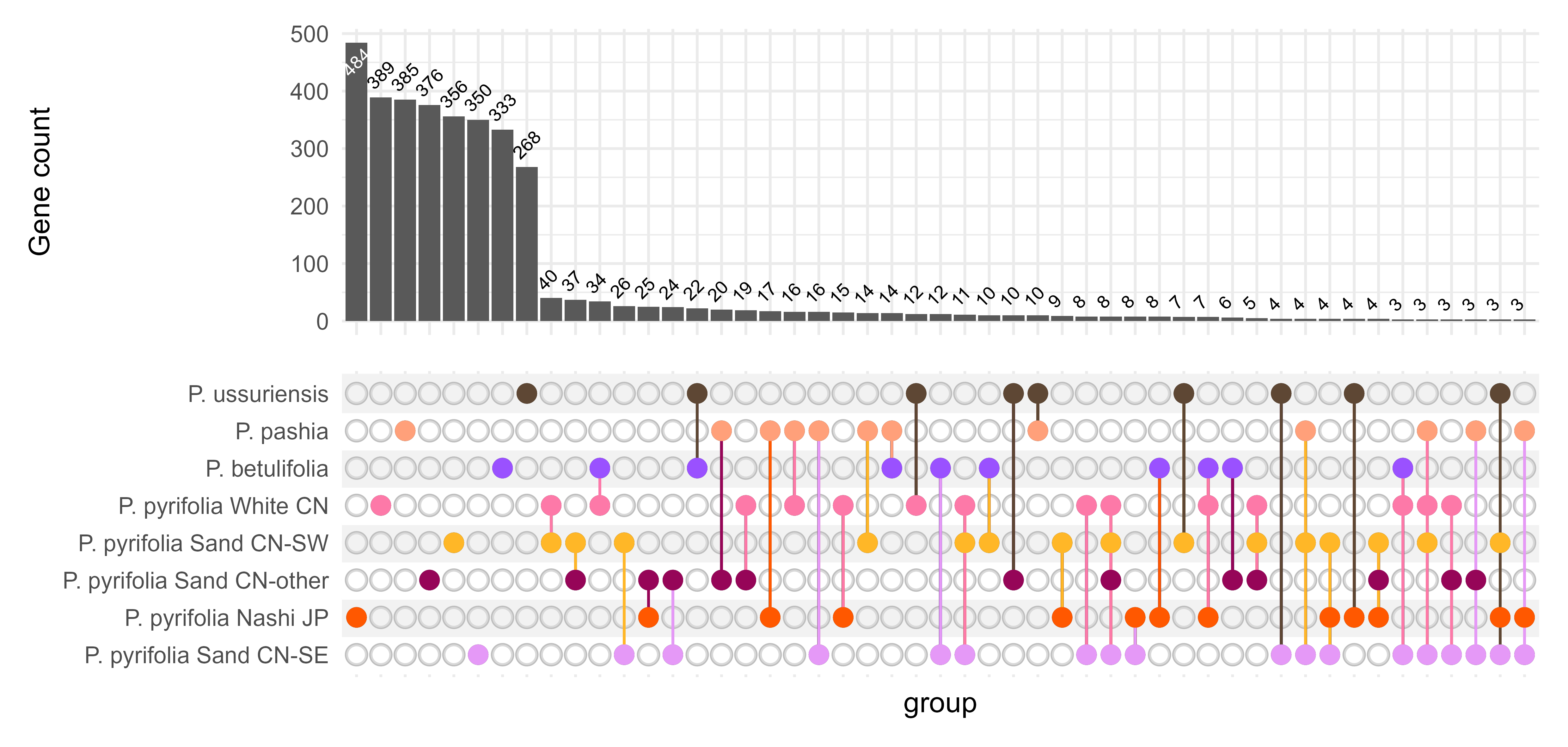

**Supplementary Figure 49 | Genes of interest in selective sweep regions in Oriental pear populations.**

The candidate genes were identified using Tajima’s *D* and *ω* statistics. Sets with fewer than three genes were not included in this plot.

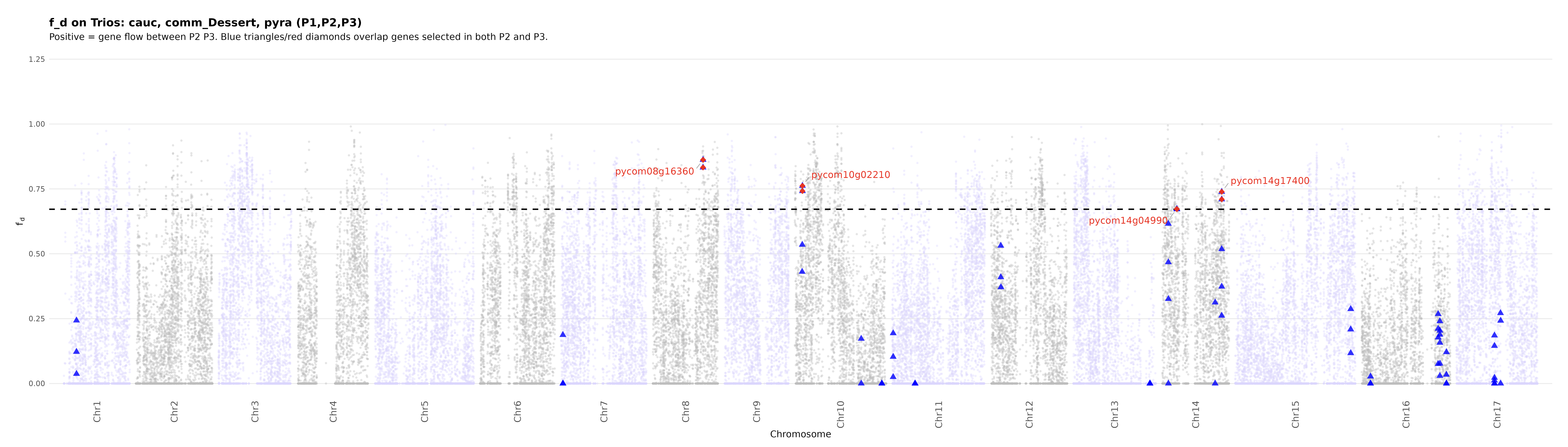

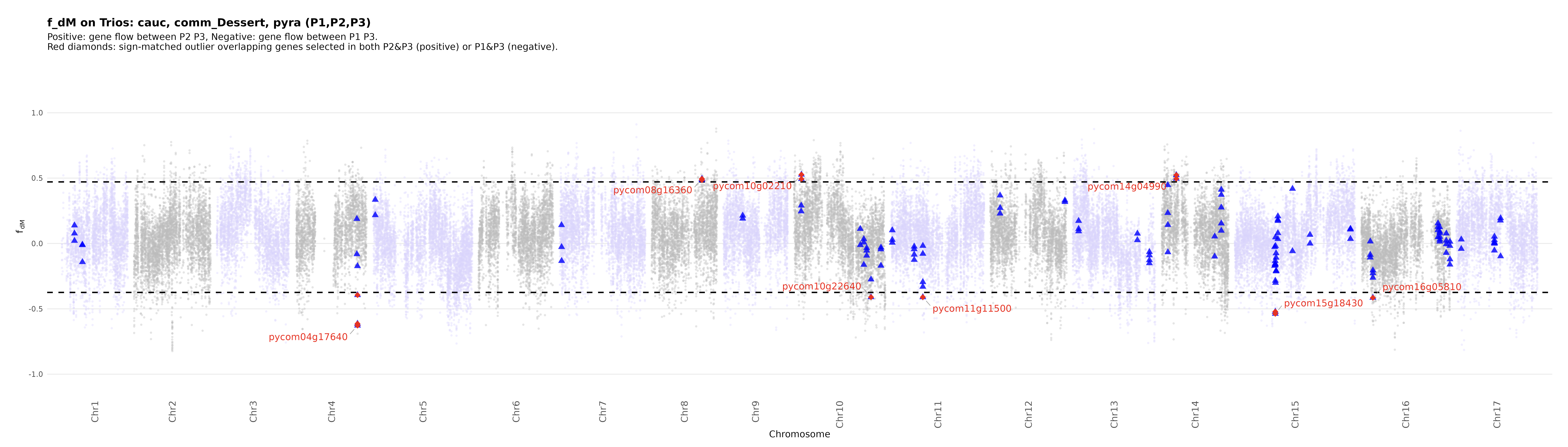

**Supplementary Figure 50 | Local introgression signals overlapping positive-selection candidate genes in Occidental pears.**

Genome-wide sliding-window estimates of the *f*_d_ (top panel) and *f*_dM_ (bottom panel) statistics, computed with Dsuite Dinvestigate, for the trio (((P1, P2), P3), O) with P1 = *Pyrus caucasica* (cauc), P2 = *P. communis* dessert (comm_Dessert), P3 = *P. pyraster* (pyra), and O = *P. ussuriensis* (ussu). Each gray point is one window, with alternating shading by chromosome. The *f*_d_ is one-tailed: positive values indicate excess allele sharing (gene flow) between P2 and P3, and the dashed line marks the top-5% outlier cutoff. The *f*_dM_ is two-tailed: positive values indicate gene flow between P2 and P3, and negative values indicate gene flow between P1 and P3, with dashed lines marking the top-2.5% and bottom-2.5% outlier cutoffs. Blue triangles denote windows overlapping genes under shared positive selection in the two relevant related populations (P2 and P3 for *f*_d_ and for positive *f*_dM_; P1 and P3 for negative *f*_dM_). Red diamonds denote sign-matched outlier windows that overlap these shared-selection genes (i.e., candidate regions of adaptive introgression) and are labeled with the overlapping gene identifiers. Positive-selection candidate genes were identified using *ω* and Tajima’s *D* statistic.

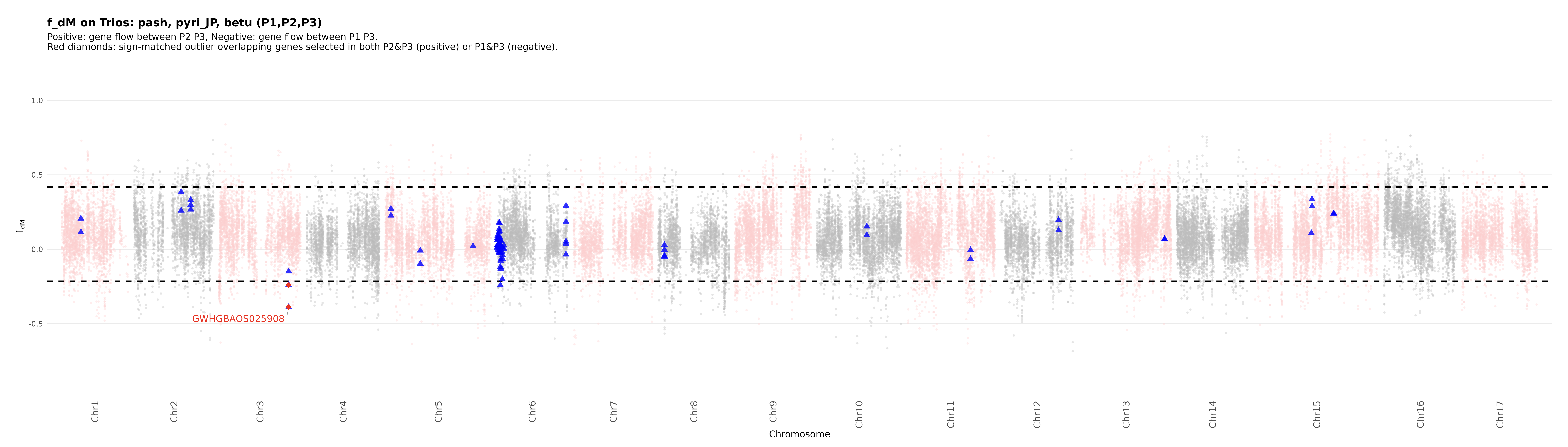

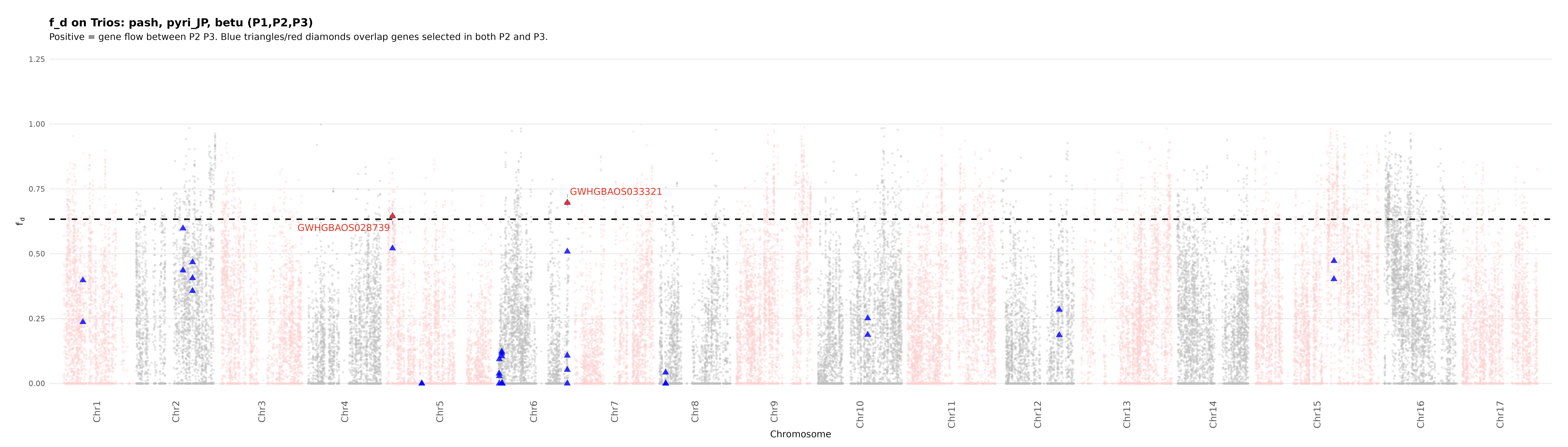

**Supplementary Figure 51 | Local introgression signals overlapping positive-selection candidate genes in Oriental pears.**

Genome-wide sliding-window estimates of the *f*_d_ (top panel) and *f*_dM_ (bottom panel) statistics, computed with Dsuite Dinvestigate, for the trio (((P1, P2), P3), O) with P1 = *Pyrus pashia* (pash), P2 = *P. pyrifolia* Nashi Japan (pyri_JP), P3 = *P. betulifolia* (betu), and O = *P. caucasica* (cauc). Each gray point is one window, with alternating shading by chromosome. The *f*_d_ is one-tailed: positive values indicate excess allele sharing (gene flow) between P2 and P3, and the dashed line marks the top-5% outlier cutoff. The *f*_dM_ is two-tailed: positive values indicate gene flow between P2 and P3, and negative values indicate gene flow between P1 and P3, with dashed lines marking the top-2.5% and bottom-2.5% outlier cutoffs. Blue triangles denote windows overlapping genes under shared positive selection in the two relevant related populations (P2 and P3 for *f*_d_ and for positive *f*_dM_; P1 and P3 for negative *f*_dM_). Red diamonds denote sign-matched outlier windows that overlap these shared-selection genes (i.e., candidate regions of adaptive introgression) and are labeled with the overlapping gene identifiers. Positive-selection candidate genes were identified using *ω* and Tajima’s *D* statistic.

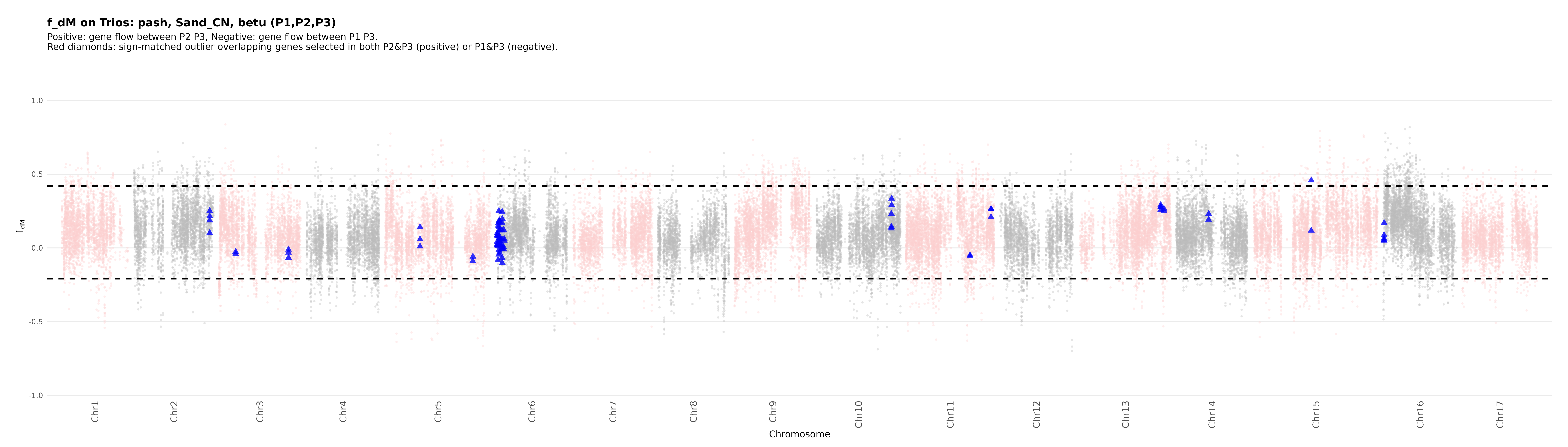

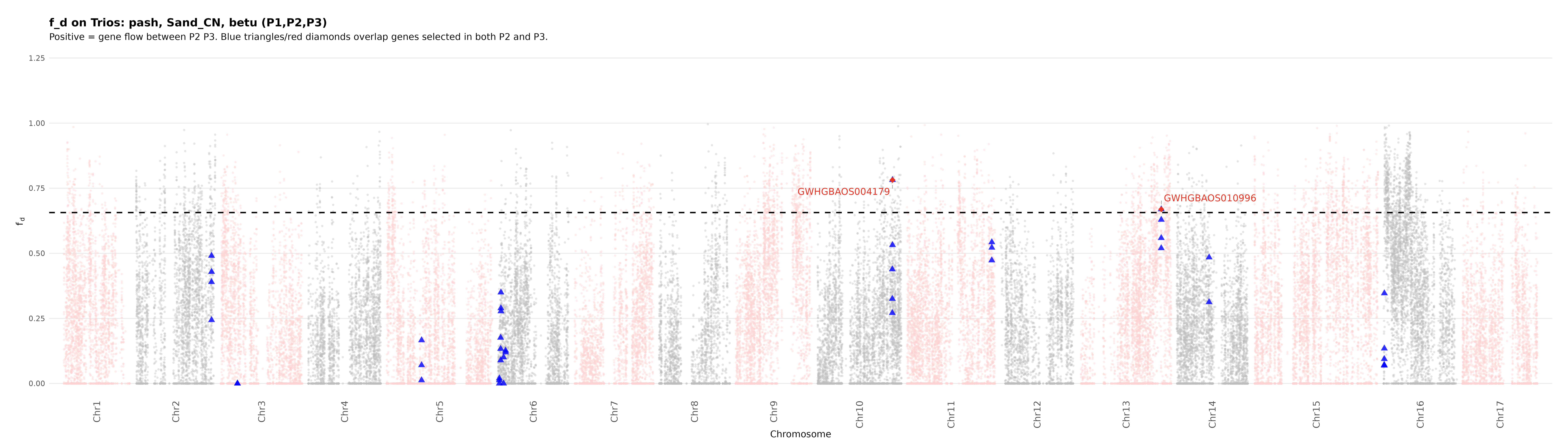

**Supplementary Figure 51 (continued) | Local introgression signals overlapping positive-selection candidate genes in Oriental pears.**

For the trio (((P1, P2), P3), O) with P1 = *Pyrus pashia* (pash), P2 = *P. pyrifolia* sand China-other (Sand_CN), P3 = *P. betulifolia* (betu), and O = *P. caucasica* (cauc).

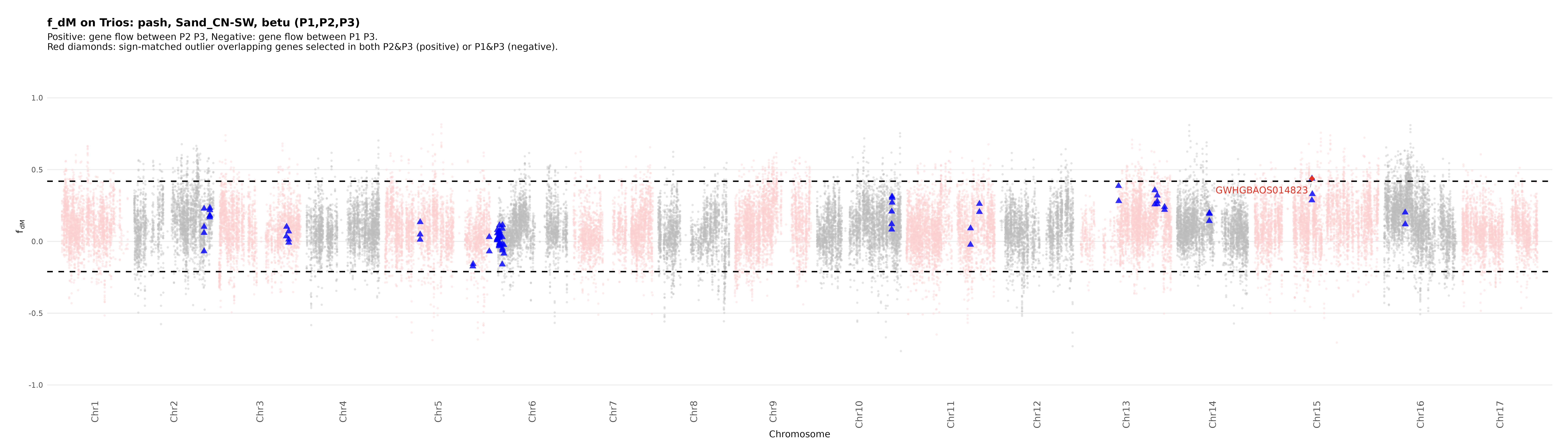

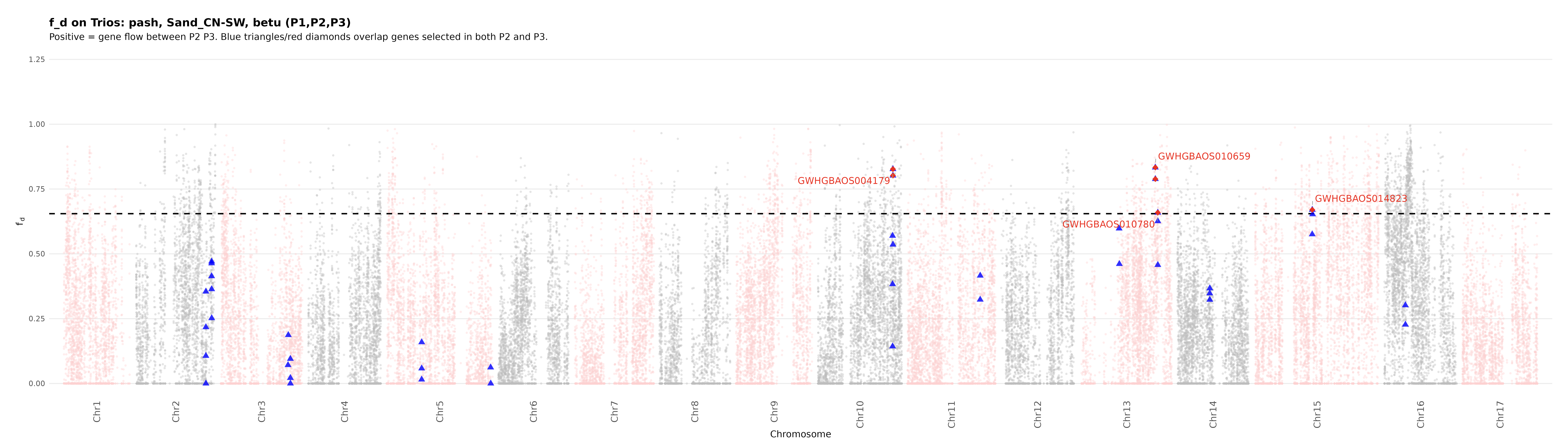

**Supplementary Figure 51 (continued) | Local introgression signals overlapping positive-selection candidate genes in Oriental pears.**

For the trio (((P1, P2), P3), O) with P1 = *Pyrus pashia* (pash), P2 = *P. pyrifolia* sand China-Southwest (Sand_CN-SW), P3 = *P. betulifolia* (betu), and O = *P. caucasica* (cauc).

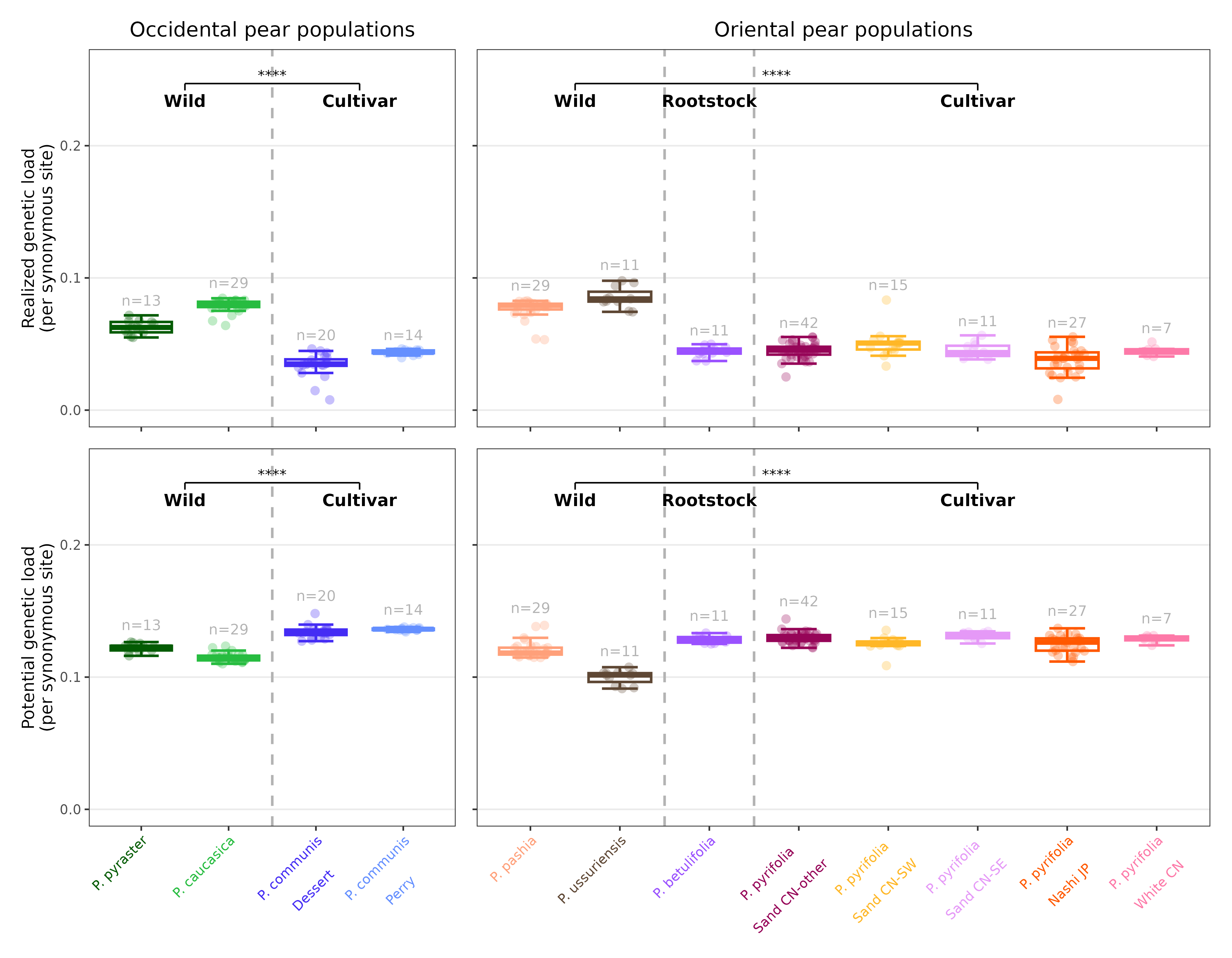

**Supplementary Figure 52 | Realized and potential genetic load in the homozygous and heterozygous states.**

Number of homozygous (top) and heterozygous (bottom) deleterious mutations (SIFT score < 0.05; SNPs) per synonymous site across all sites (selection and control regions combined), using *P. communis* and *P. pyrifolia* as reference genomes for Occidental and Oriental pear populations, respectively. Each point is one individual; boxes show the median and interquartile range. Wild vs. cultivar groups were compared with a two-sided Wilcoxon rank-sum test pooling individuals within each group. Source data are provided in Supplementary Table 16; exact *P* values are provided in Supplementary Table 17. Significance levels: ns, *P* > 0.05 (not significant); * *P* ≤ 0.05; ** *P* ≤ 0.01; *** *P* ≤ 0.001; **** *P* ≤ 0.0001.

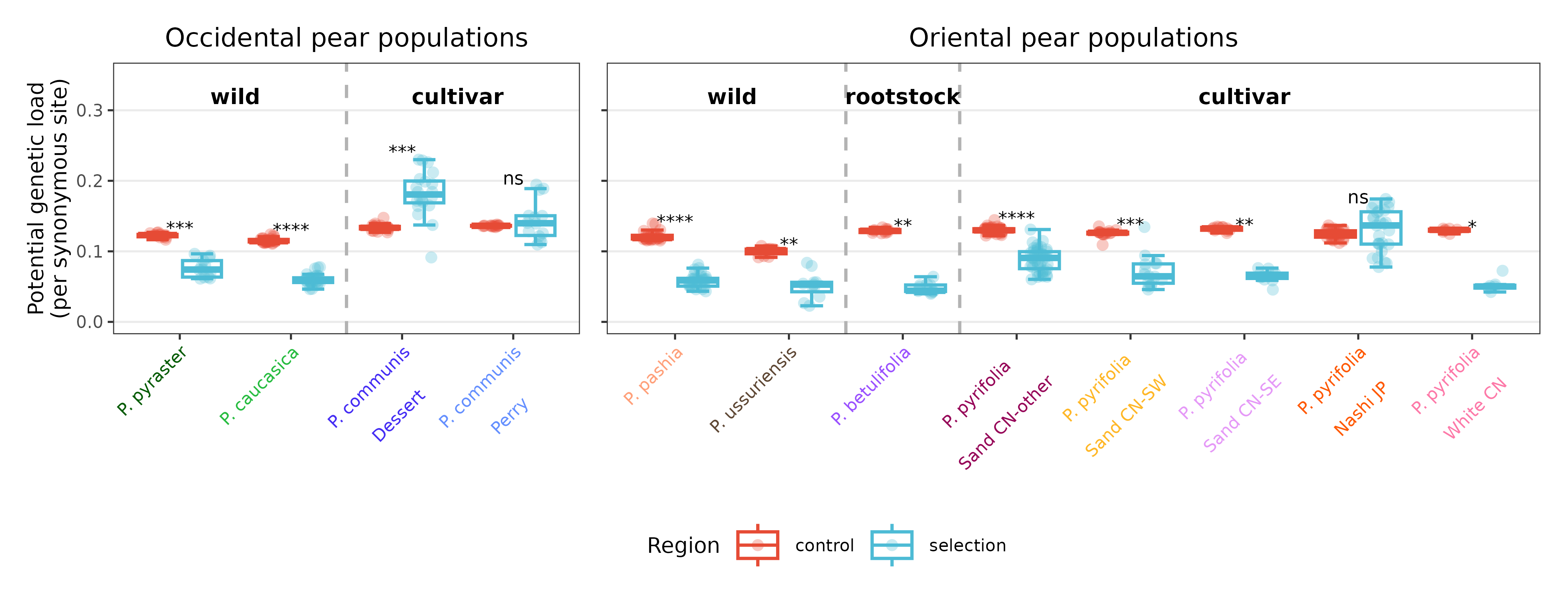

**Supplementary Figure 53 | Potential genetic load, selection vs. control regions.**Number of heterozygous deleterious mutations (SIFT score < 0.05; SNPs) per synonymous site in selection and control regions, using *P. communis* and *P. pyrifolia* as reference genomes for Occidental and Oriental pear populations, respectively. Each point is one individual; boxes show the median and interquartile range. Within each population, selection and control values come from the same individuals and were compared with a two-sided paired Wilcoxon signed-rank test; *P* values were Holm-adjusted within each lineage. Source data are provided in Supplementary Table 16. *P* values are provided in Supplementary Table 17. Significance levels: ns, *P* > 0.05 (not significant); * *P* ≤ 0.05; ** *P* ≤ 0.01; *** *P* ≤ 0.001; **** *P* ≤ 0.0001.

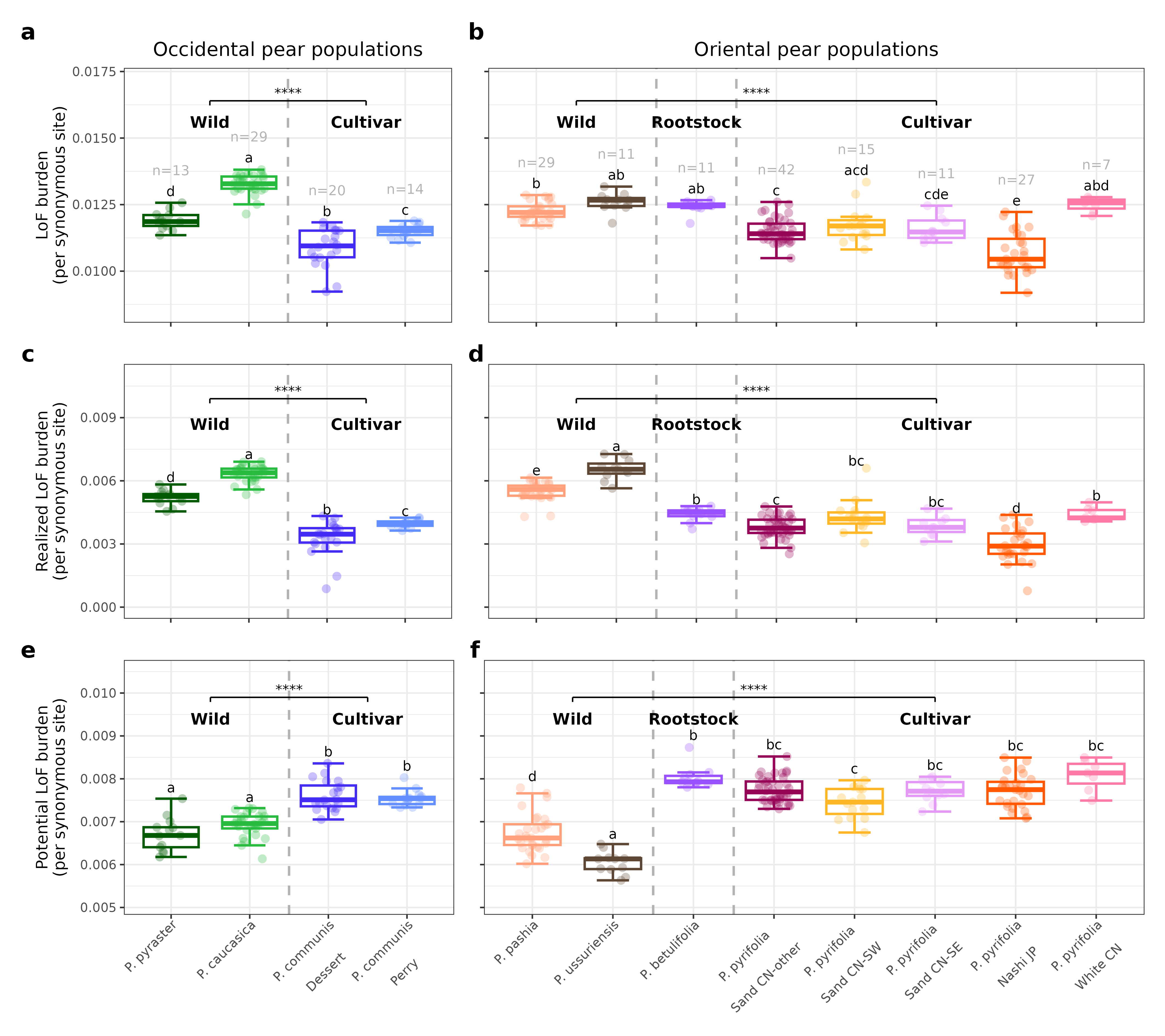

**Supplementary Figure 54 | Genome-wide loss-of-function burden.**

Number of high-confidence loss-of-function (LoF) alleles (stop-gained, splice-donor, and splice-acceptor SNPs predicted as LoF by both VEP and SnpEff) per synonymous site, using *P. communis* and *P. pyrifolia* as reference genomes for Occidental and Oriental pear populations, respectively. **a–b**, total LoF burden; **c–d**, LoF burden in the homozygous state (realized burden); **e–f**, LoF burden in the heterozygous state (potential burden). Each dot represents one individual; n, number of individuals per population. Letters indicate significant differences among populations within each lineage (pairwise Wilcoxon test, Holm-adjusted *P* < 0.01). Brackets compare wild and cultivated populations (rootstock excluded). Source data are provided in Supplementary Table 19. *P* values are provided in Supplementary Table 20. Significance levels (Wilcoxon test): ns, *P* > 0.05 (not significant); * *P* ≤ 0.05; ** *P* ≤ 0.01; *** *P* ≤ 0.001; **** *P* ≤ 0.0001.

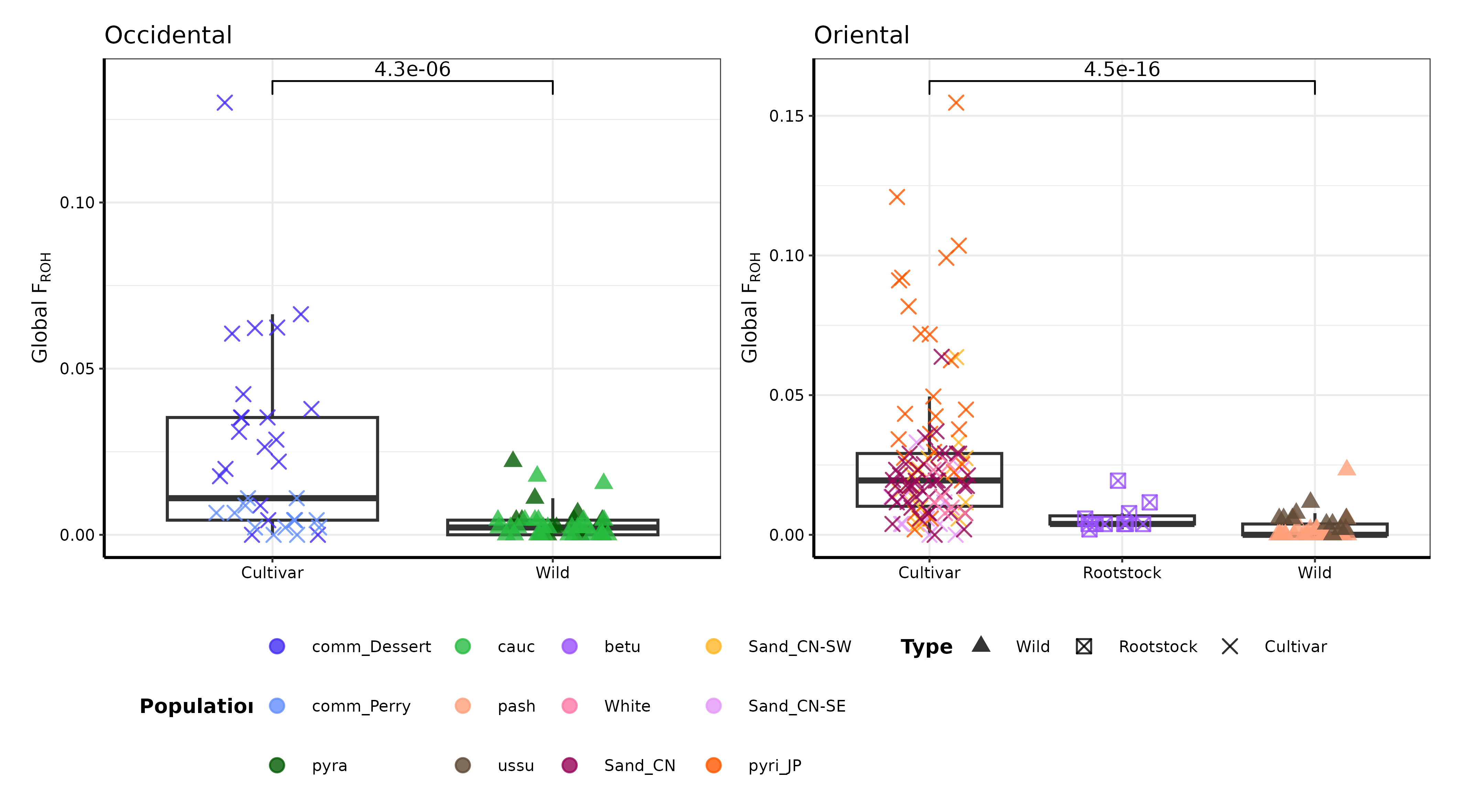

**Supplementary Figure 55 | Elevated *F*_ROH_ in cultivated relative to wild pear populations.**

Proportion of the genome in runs of homozygosity (*F*_ROH_, estimated with ROHan; Supplementary Note 2), shown by germplasm type for the Occidental (left) and Oriental (right) lineages. Boxes show the median and interquartile range; individual points are jittered horizontally, colored by population, and shaped by population type (Wild, Rootstock, Cultivar). *F*_ROH_ is significantly higher in Cultivar than in Wild individuals in both lineages (two-sided Wilcoxon rank-sum test).

Population abbreviations:

cauc, *P. caucasica;*

comm dessert, *P. communis* dessert;

comm perry, *P. communis* perry;

pyra, *P. pyraster*;

betu, *Pyrus betulifolia*;

pash, *Pyrus pashia*;

pyri JP, *Pyrus pyrifolia* Nashi Japan;

Sand CN, *Pyrus pyrifolia* sand China-other;

Sand CN-SE, *Pyrus pyrifolia* sand China-Southeast;

Sand CN-SW, *Pyrus pyrifolia* sand China-Southwest;

ussu, *Pyrus ussuriensis*;

White, *Pyrus pyrifolia* white China.

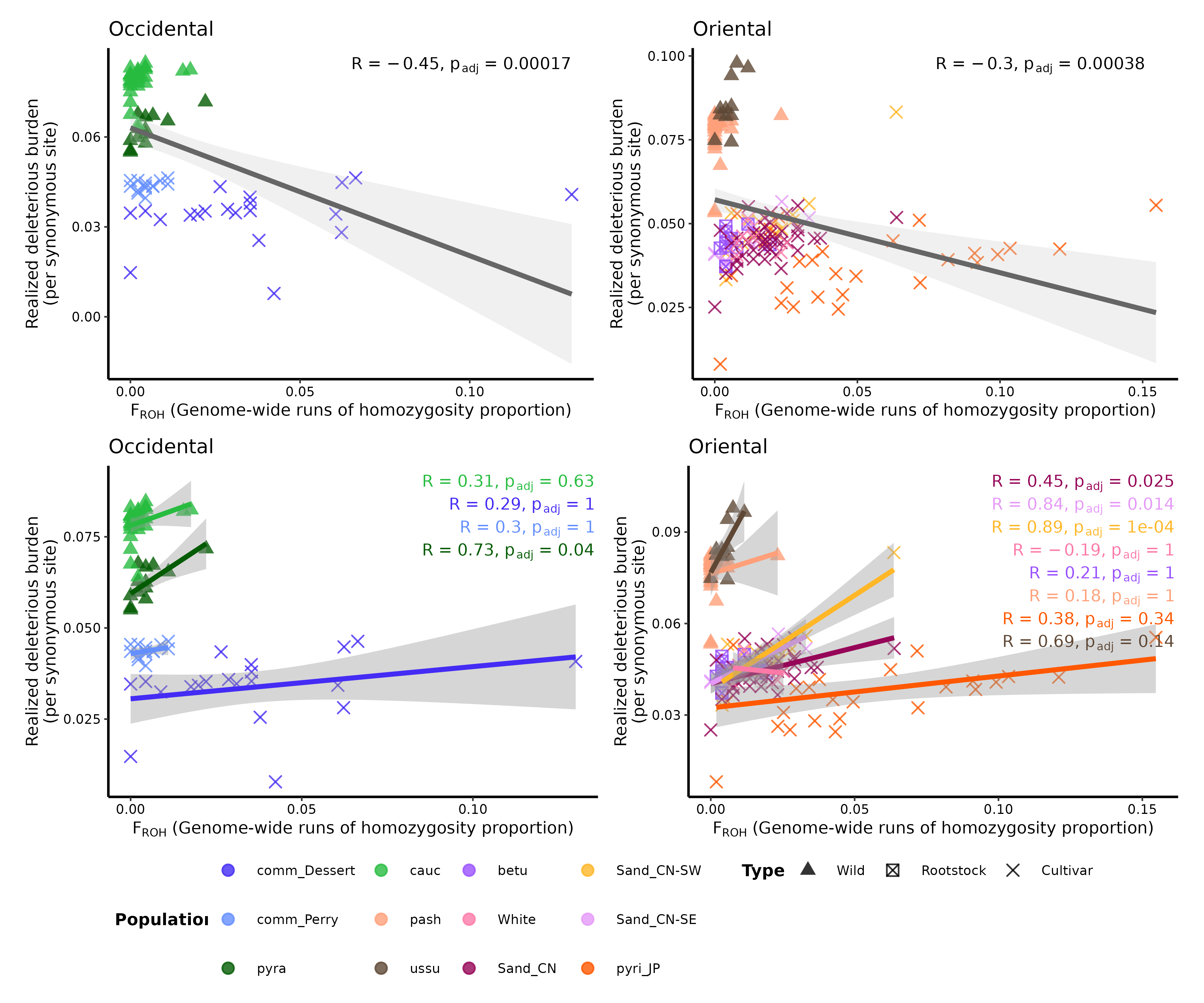

**Supplementary Figure 56 | Deleterious genetic burden scales with genome-wide *F*_ROH_ within each pear lineage (Realized deleterious burden).**

Relationship between genome-wide *F*_ROH_ and the deleterious genetic burden for the Occidental (left column) and Oriental (right column) lineages. Top row: ordinary least-squares linear regression (gray line; gray band, 95% confidence interval) fitted to all individuals pooled within each lineage. Bottom row: linear regressions fitted separately for each population (lines colored by population; gray bands, 95% confidence intervals). Pearson's correlation coefficient (R) and Holm-adjusted two-sided P-value (*P*_adj_) are shown in each panel. Each point is one individual, colored by population and shaped by type (Wild, Rootstock, or Cultivar).

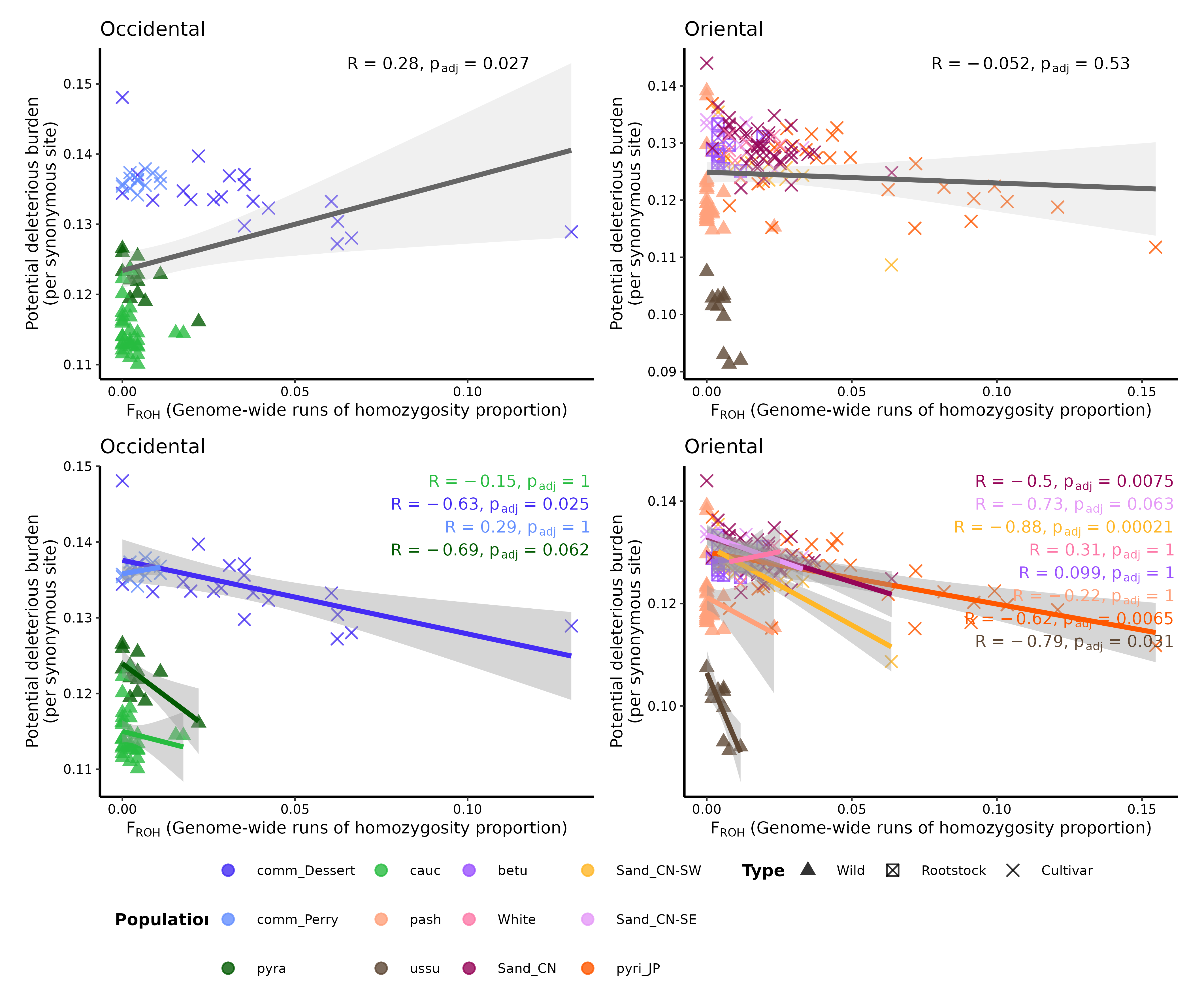

**Supplementary Figure 56 (continued) | Deleterious genetic burden scales with genome-wide *F*_ROH_ within each pear lineage (Potential deleterious burden).**

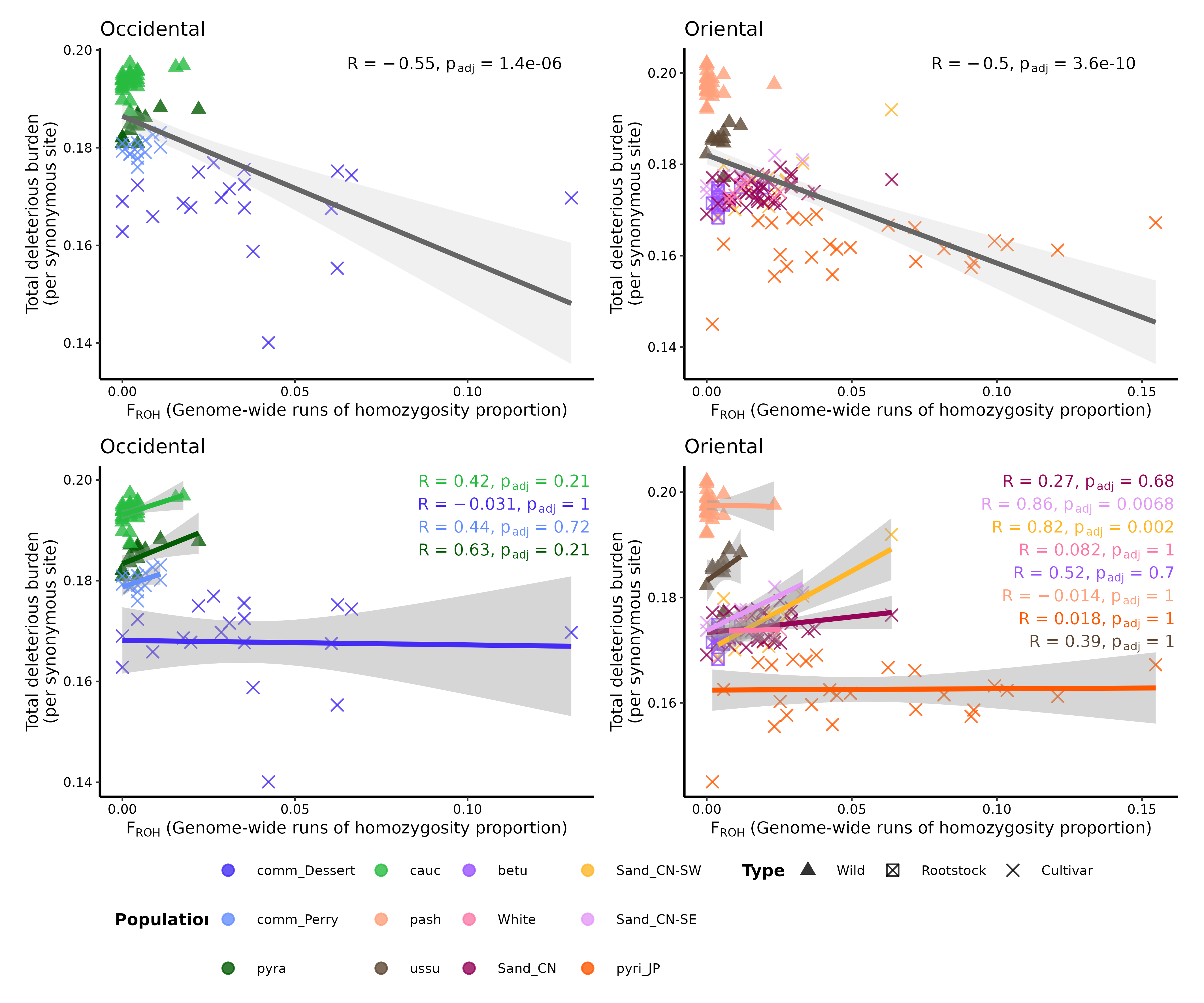

**Supplementary Figure 56 (continued) | Deleterious genetic burden scales with genome-wide *F*_ROH_ within each pear lineage (Total deleterious burden).**

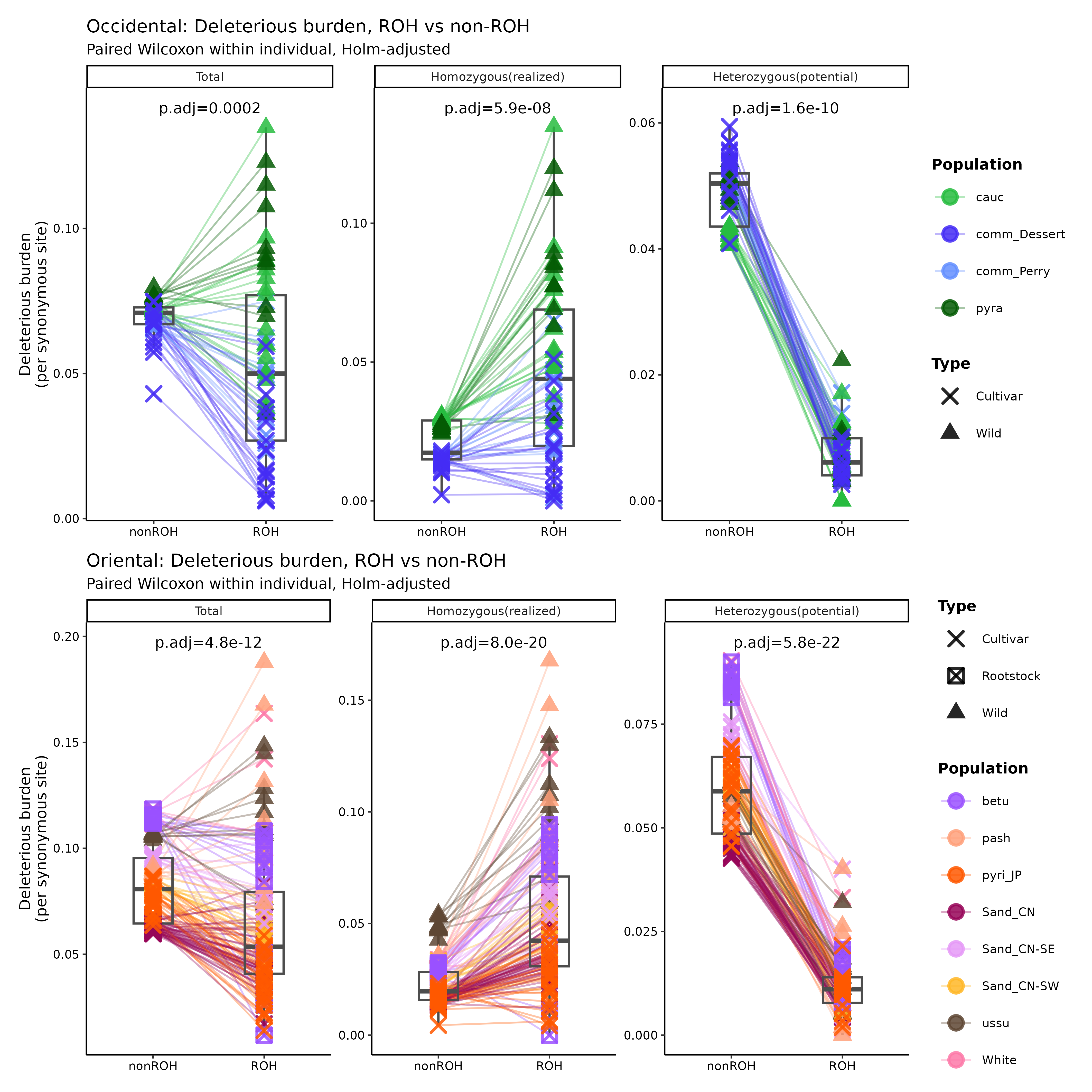

**Supplementary Figure 57 | Deleterious burden in ROH and non-ROH regions within individuals.**

Paired comparison of deleterious burden between ROH and non-ROH regions of the same individual, for the Occidental (top row; *n* = 57) and Oriental (bottom row; *n* = 128) lineages. Burden is shown for all deleterious alleles combined (Total) and separately for homozygous (realized) and heterozygous (potential) genotypes (left to right). Individuals without detected ROH were excluded. Each point represents one individual, colored by population and shaped by type (Wild, Rootstock, or Cultivar). Lines connect the ROH and non-ROH values of the same individual. Boxplots show the median and interquartile range, with whiskers extending to 1.5× the interquartile range. *P*-values are from two-sided paired Wilcoxon signed-rank tests (ROH vs. non-ROH within individuals), Holm-adjusted (*P*_adj_).

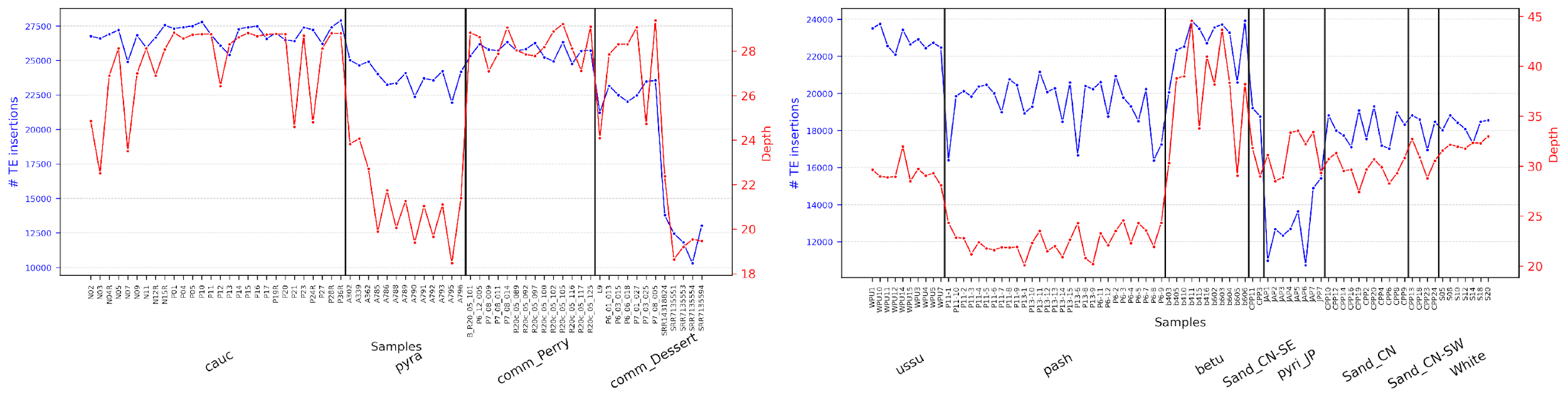

**Supplementary Figure 58 | Line plots showing the number of TE insertions and sequencing depth across samples from different populations.**

Blue lines represent the number of TE insertions, and red lines represent sequencing depth. Black vertical lines separate population groups. Occidental populations are shown on the left and Oriental populations on the right. To assess sequencing quality, TE insertions were scored as present or absent, with homozygous sites (1/1) counted once.

Population abbreviations:

cauc, *P. caucasica;*

comm dessert, *P. communis* dessert;

comm perry, *P. communis* perry;

pyra, *P. pyraster*;

betu, *Pyrus betulifolia*;

pash, *Pyrus pashia*;

pyri JP, *Pyrus pyrifolia* Nashi Japan;

Sand CN, *Pyrus pyrifolia* sand China-other;

Sand CN-SE, *Pyrus pyrifolia* sand China-Southeast;

Sand CN-SW, *Pyrus pyrifolia* sand China-Southwest;

ussu, *Pyrus ussuriensis*;

White, *Pyrus pyrifolia* white China.

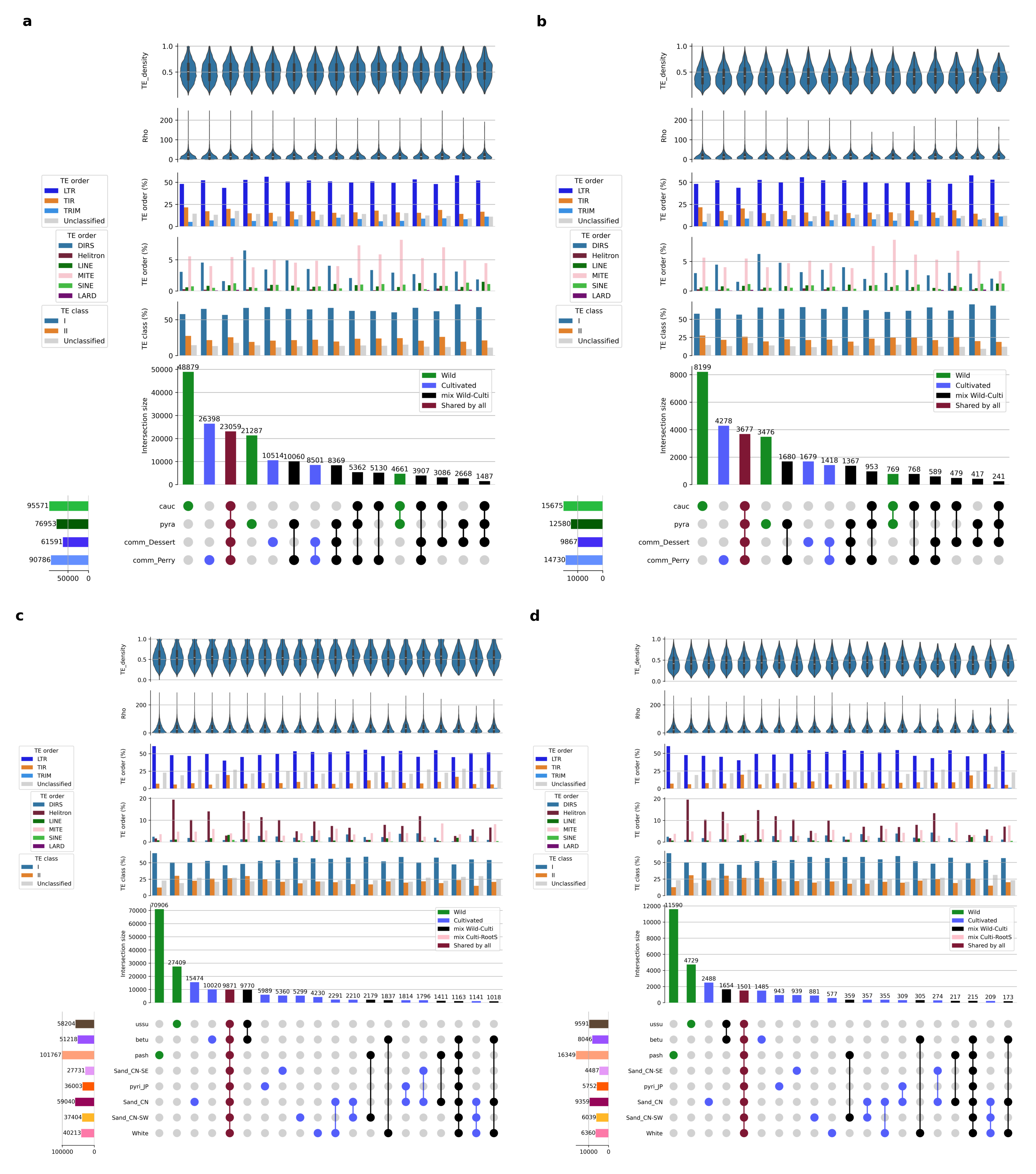

**Supplementary Figure 59 | Genome-wide and gene-proximal TE polymorphism landscape in Occidental and Oriental pear populations.**

Genome-wide **(a,c)** and gene-proximal **(b,d)** TE polymorphisms detected by MEGAnE, considering genes not under positive selection and their 2-kb upstream regions. Throughout the figure, panels (**a, b**) correspond to Occidental populations, while panels (**c, d**) correspond to Oriental populations. The TE density was calculated using a 50-kb window (number of base pairs annotated as TE per window). The distribution of Rho (ρ) represents an approximation of the recombination rate across the genome. To improve visualization clarity, the TE orders were divided into two separate panels. In the Oriental pear dataset, only the 20 largest intersections were displayed to enhance clarity.

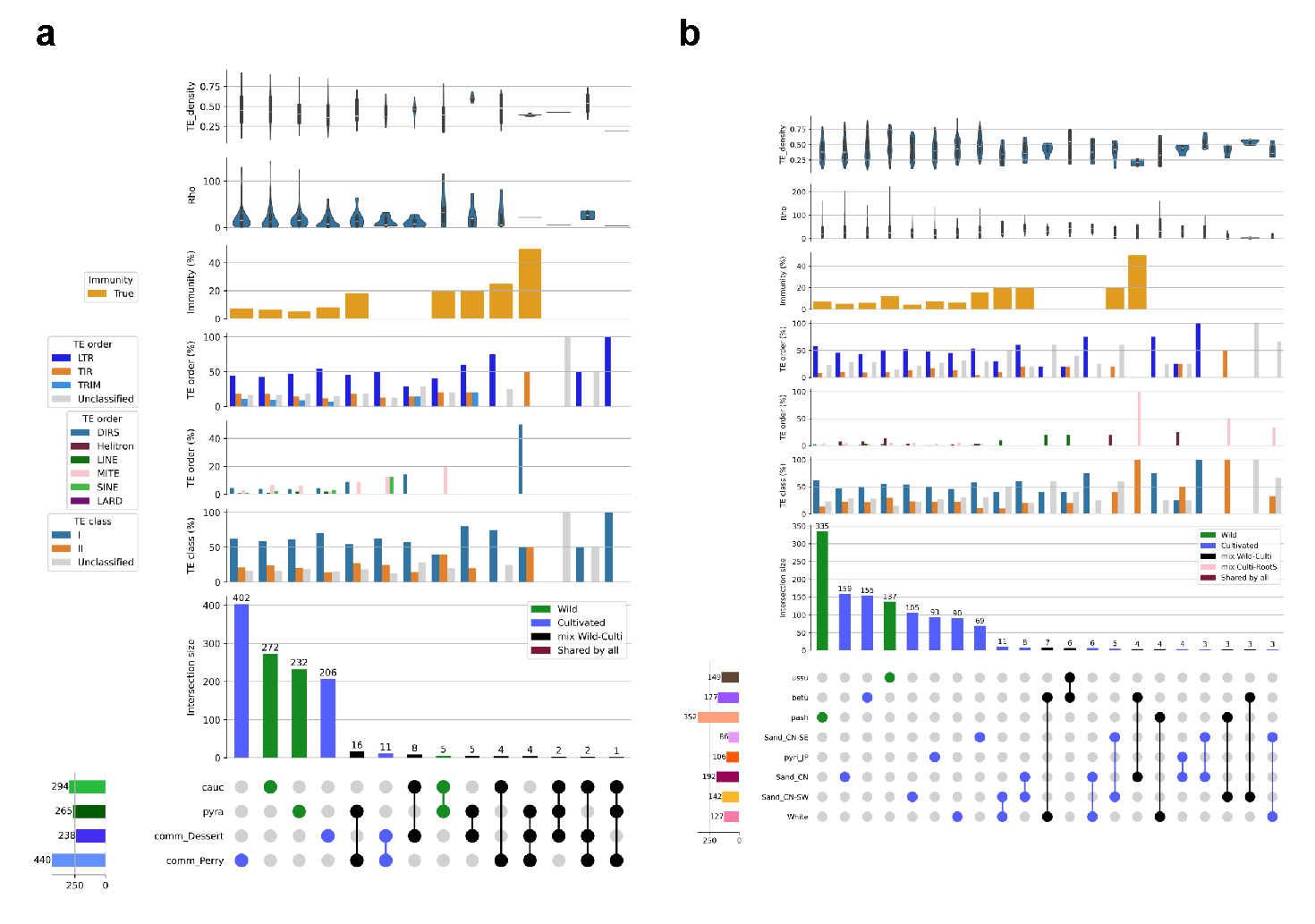

**Supplementary Figure 60 | Shared and fixed transposable element insertions in Occidental and Oriental pear populations.**

**a,b** UpSet plots showing TE polymorphisms located 2 kb upstream of genes under positive selection. The distribution of Rho (*ρ*) approximates the recombination rate across the genome. The TE density was calculated using a 50-kb window (number of base pairs annotated as TE per window). To improve visualization clarity, the TE orders were divided into two separate panels. In the Oriental pear dataset, only the 20 largest intersections were displayed to enhance clarity. DIRS, Dictyostelium intermediate repeat sequence; LARD, Large Retrotransposon Derivative; LINE, Long Interspersed Nuclear Element; LTR, Long Terminal Repeat; SINE, Short Interspersed Nuclear Element; TRIM, Terminal repeat Retrotransposon in Miniature; Helitron, Helitron transposon; MITE, Miniature Inverted repeat Transposable Element; TIR, Terminal Inverted Repeat.

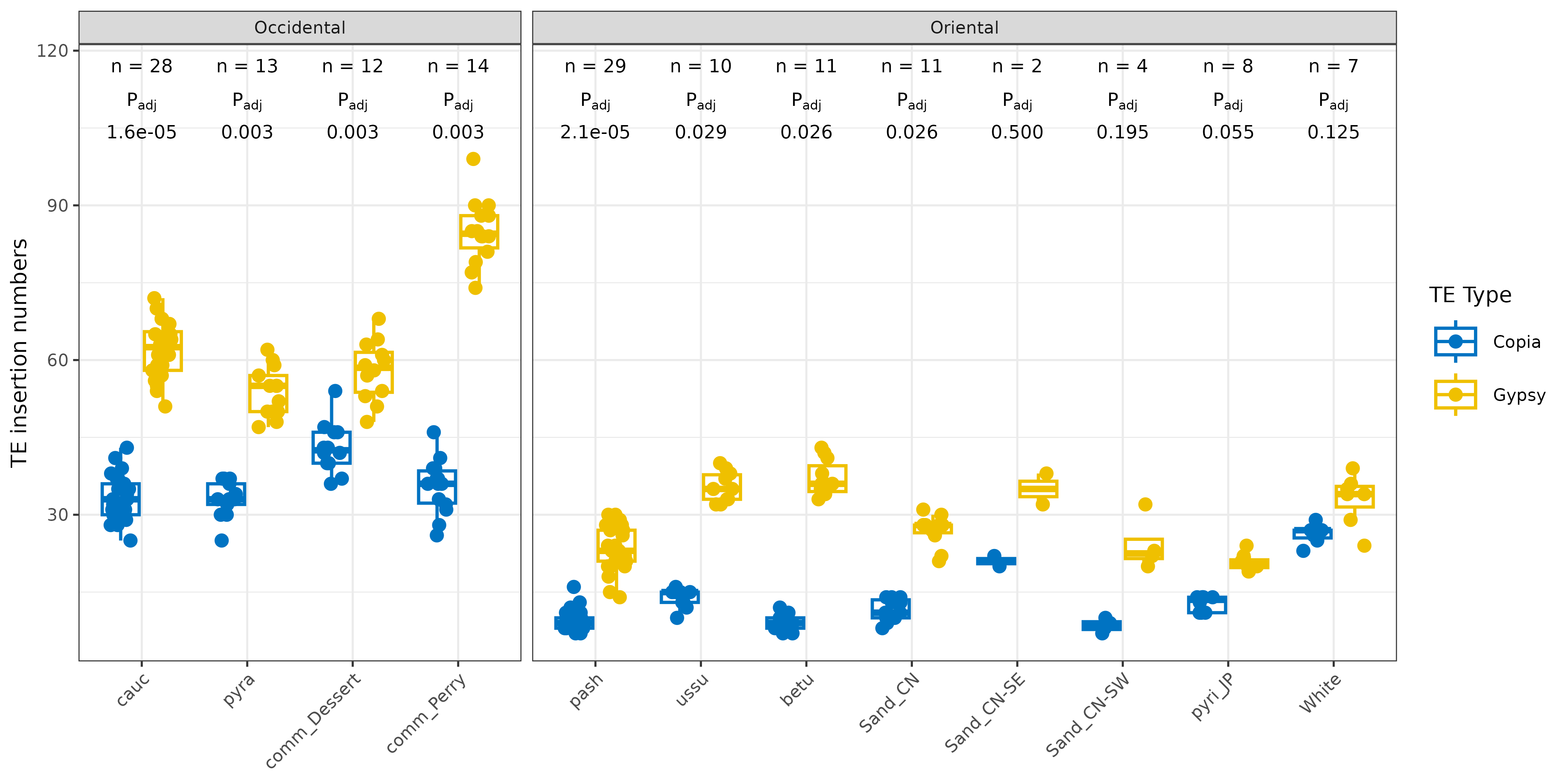

**Supplementary Figure 61 | Number of Copia and Gypsy TEs present per individual across pear populations.**

Each point represents one individual. Boxplots show the median and interquartile range of per-individual present TE insertion counts for the Copia (blue) and Gypsy (yellow) superfamilies, grouped by population within the Occidental and Oriental pear groups. Only TE polymorphisms overlapping the putative selective sweep regions of the individual's own population are counted. Above each population, n is the number of individuals, and *P*_adj_ is from a two-sided paired Wilcoxon signed-rank test (Gypsy vs Copia), Holm-corrected within the Occidental and Oriental lineages.

Population abbreviations:

cauc, *P. caucasica;*

comm dessert, *P. communis* dessert;

comm perry, *P. communis* perry;

pyra, *P. pyraster*;

betu, *Pyrus betulifolia*;

pash, *Pyrus pashia*;

pyri JP, *Pyrus pyrifolia* Nashi Japan;

Sand CN, *Pyrus pyrifolia* sand China-other;

Sand CN-SE, *Pyrus pyrifolia* sand China-Southeast;

Sand CN-SW, *Pyrus pyrifolia* sand China-Southwest;

ussu, *Pyrus ussuriensis*;

White, *Pyrus pyrifolia* white China.

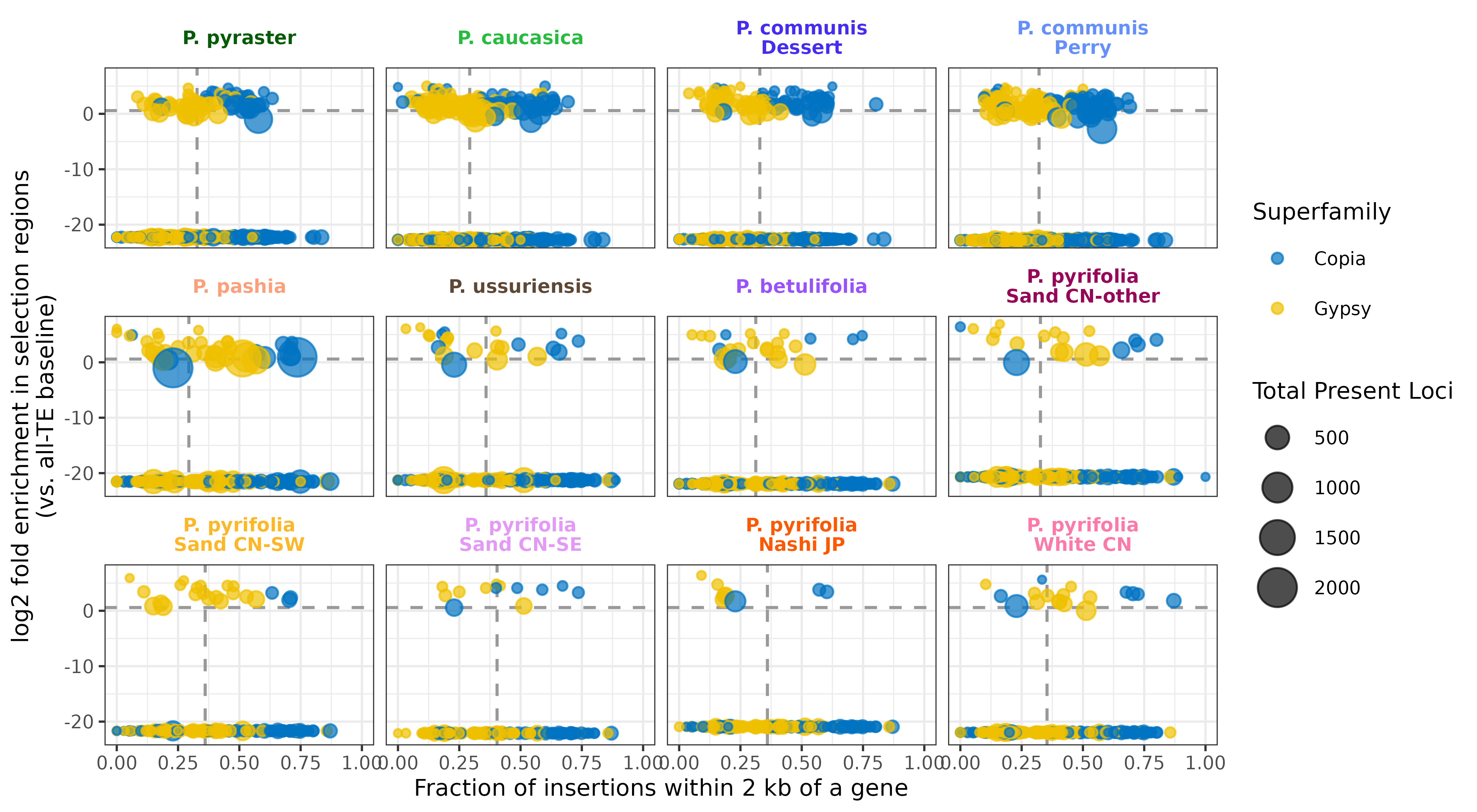

**Supplementary Figure 62 | Consensus-level enrichment of Copia and Gypsy retrotransposons in selection regions relative to their proximity to genes.**

For each population, individual points represent single TE consensus sequences, restricted to the Copia and Gypsy LTR-retrotransposon superfamilies and to consensi with ≥ 5 insertion loci in that population. The x-axis shows the fraction of a consensus's insertion loci located within 2 kb of an annotated gene. The y-axis shows its log2 fold enrichment in selection regions relative to the all-TE baseline of that population, *i.e*., log2[(fraction of the consensus's loci in selection regions) / (fraction of all TE loci in selection regions)]. Consensi with no loci in selection regions have an undefined log2 enrichment and are plotted at the bottom of each panel. Point color denotes superfamily (Copia, blue; Gypsy, gold); point size denotes the total number of present loci for that consensus. The horizontal dashed line marks 1.5-fold enrichment. It is an effect-size reference only, not a significance threshold. None of the consensi shown was significantly enriched or depleted (two-sided Fisher's exact test, Benjamini–Hochberg-adjusted q ≥ 0.05 in every population). The vertical dashed line marks the median fraction of insertions within 2 kb of a gene among the consensi shown in that population. Points in the upper-right region of a panel would indicate consensi that are both overrepresented in selection regions and preferentially located near genes.

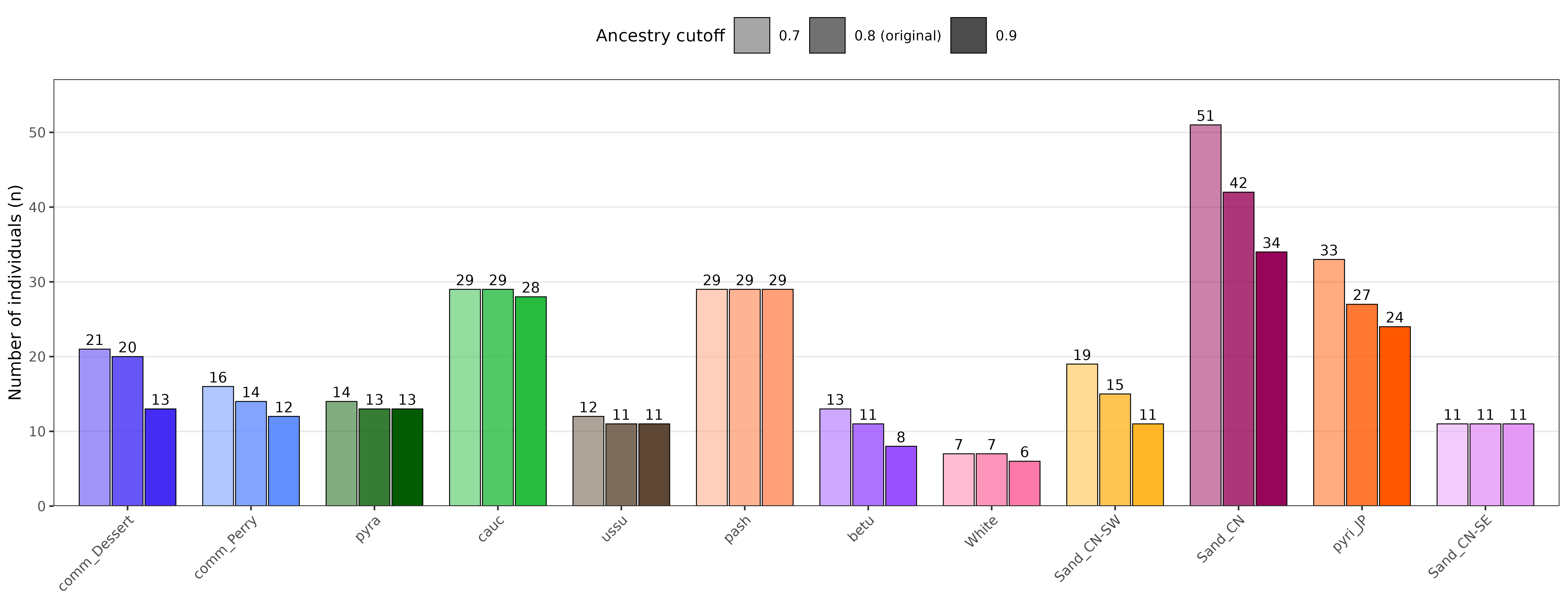

**Supplementary Figure 63 | Number of individuals in each population at three ancestry thresholds.**

Bars show the number of individuals assigned to each population when the ancestry threshold was 0.7, 0.8, or 0.9; darker shading indicates a higher threshold. The 0.8 threshold was used in the main analyses. Ancestry threshold, minimum ancestry proportion estimated by fastStructure. Numbers above bars are sample sizes.

Population abbreviations:

comm dessert, *P. communis* dessert;

comm perry, *P. communis* perry;

pyra, *P. pyraster*;

cauc, *P. caucasica;*

ussu, *Pyrus ussuriensis*;

pash, *Pyrus pashia*;

betu, *Pyrus betulifolia*;

White, *Pyrus pyrifolia* white China;

Sand CN-SW, *Pyrus pyrifolia* sand China-Southwest;

Sand CN, *Pyrus pyrifolia* sand China-other;

pyri JP, *Pyrus pyrifolia* Nashi Japan;

Sand CN-SE, *Pyrus pyrifolia* sand China-Southeast.

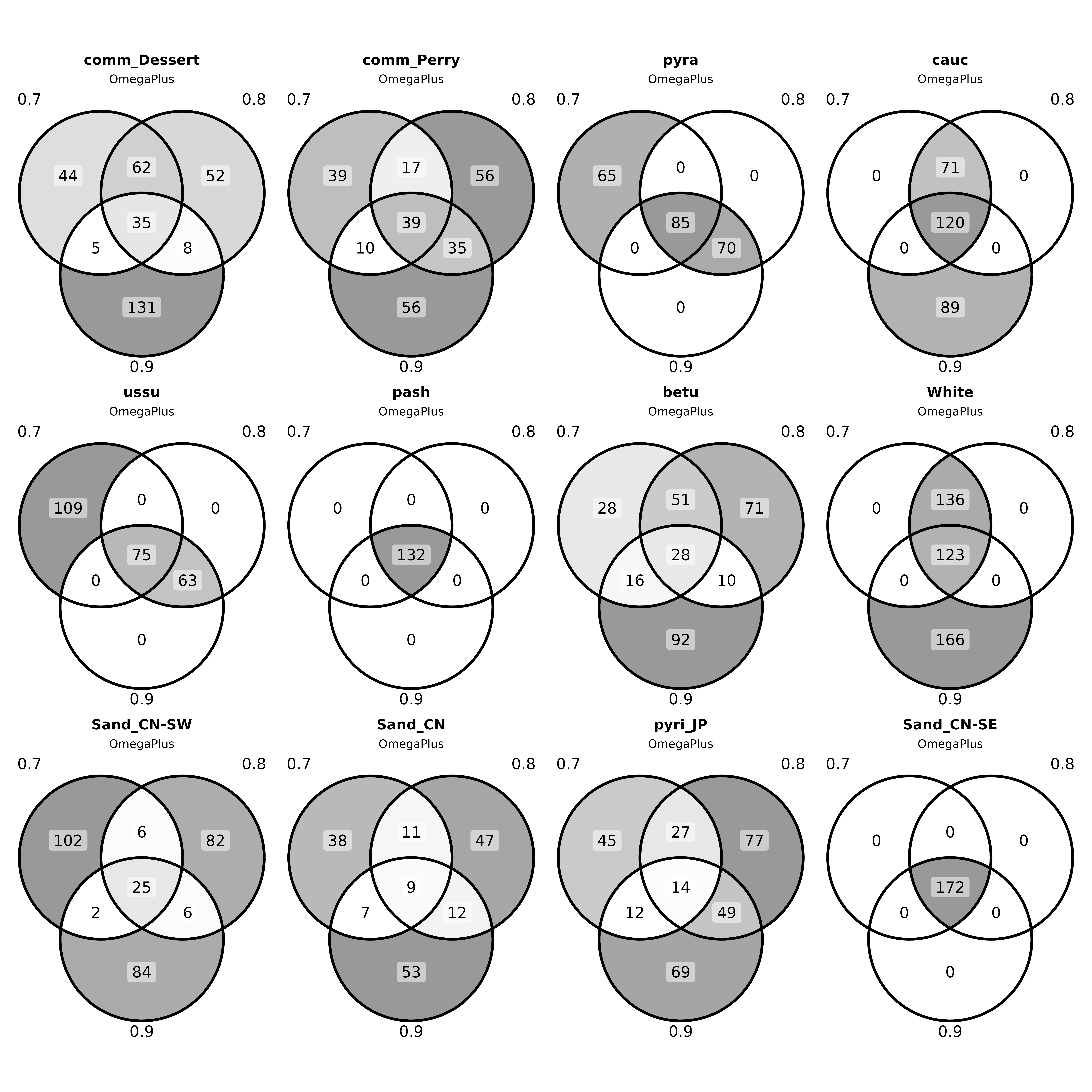

**Supplementary Figure 64 | Overlap of candidate genes under positive selection across three ancestry thresholds (OmegaPlus).**

For each population, Venn diagrams show candidate genes detected with populations defined at ancestry thresholds of 0.7, 0.8, and 0.9. Numbers are candidate gene counts identified for selective sweeps. The center shows genes detected in all three thresholds. Darker shading indicates more genes.

Population abbreviations:

comm dessert, *P. communis* dessert;

comm perry, *P. communis* perry;

pyra, *P. pyraster*;

cauc, *P. caucasica;*

ussu, *Pyrus ussuriensis*;

pash, *Pyrus pashia*;

betu, *Pyrus betulifolia*;

White, *Pyrus pyrifolia* white China;

Sand CN-SW, *Pyrus pyrifolia* sand China-Southwest;

Sand CN, *Pyrus pyrifolia* sand China-other;

pyri JP, *Pyrus pyrifolia* Nashi Japan;

Sand CN-SE, *Pyrus pyrifolia* sand China-Southeast.

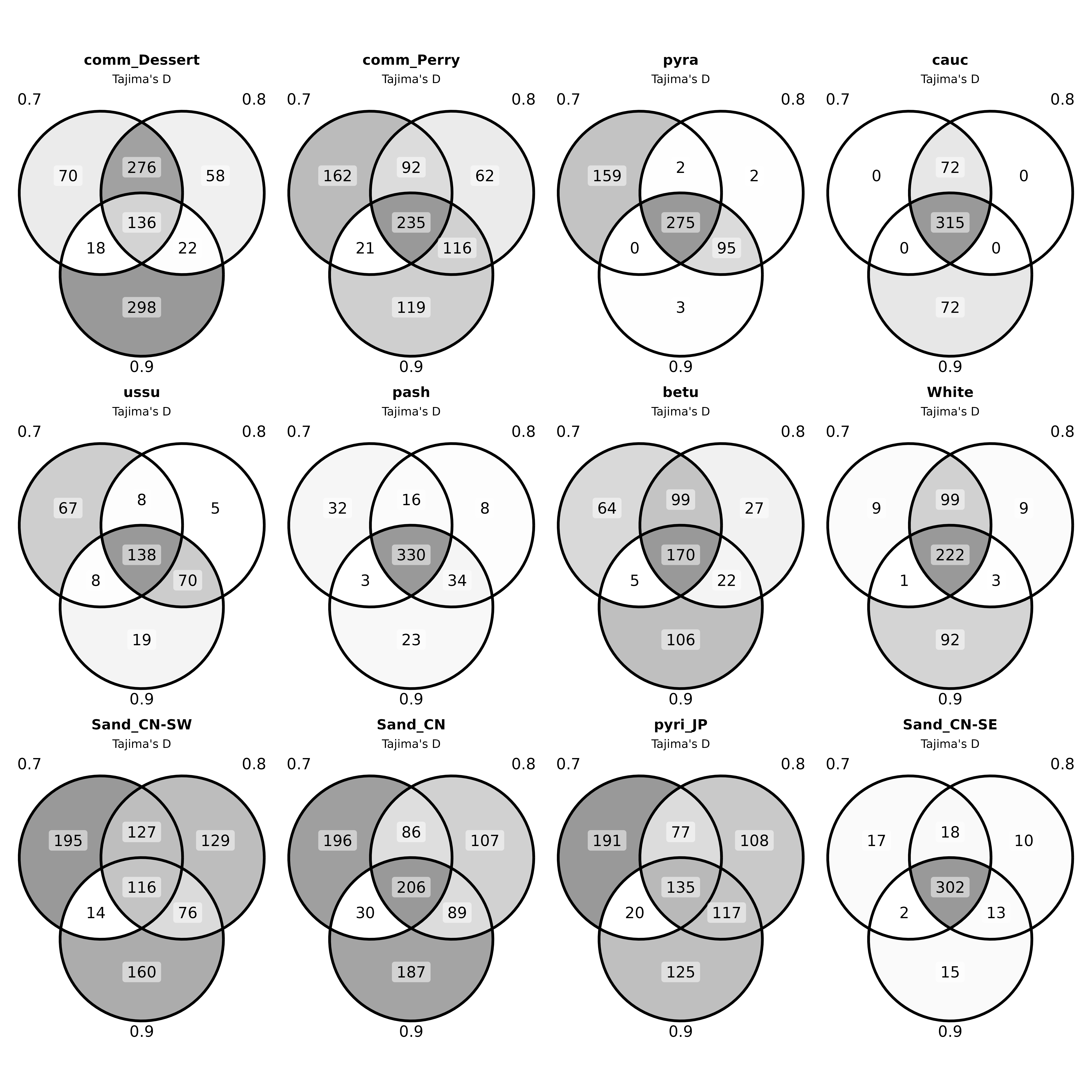

**Supplementary Figure 64 (continued) | Overlap of candidate genes under positive selection across three ancestry thresholds (Tajima’s *D*).**

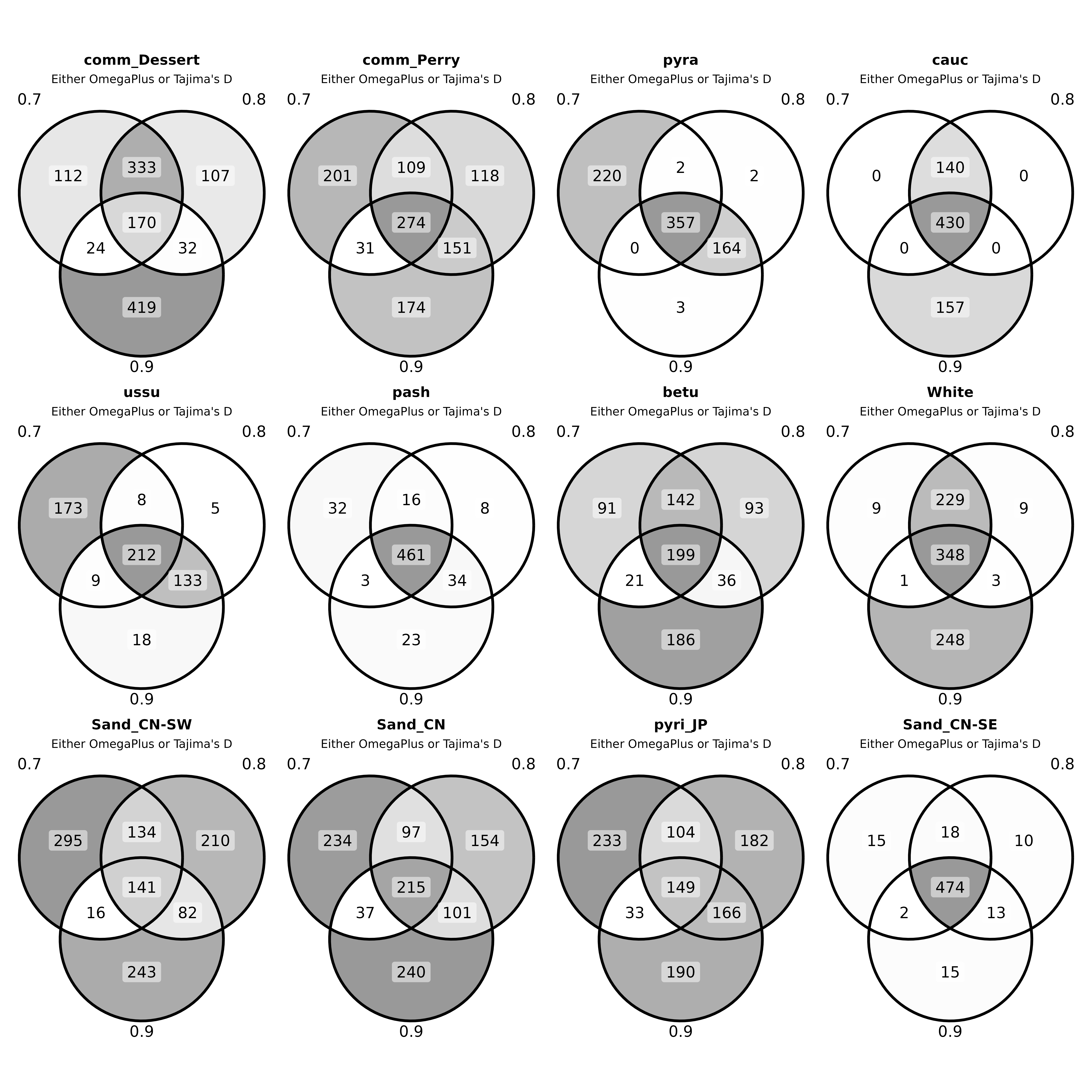

**Supplementary Figure 64 (continued) | Overlap of candidate genes under positive selection across three ancestry thresholds (either OmegaPlus or Tajima’s *D*).**

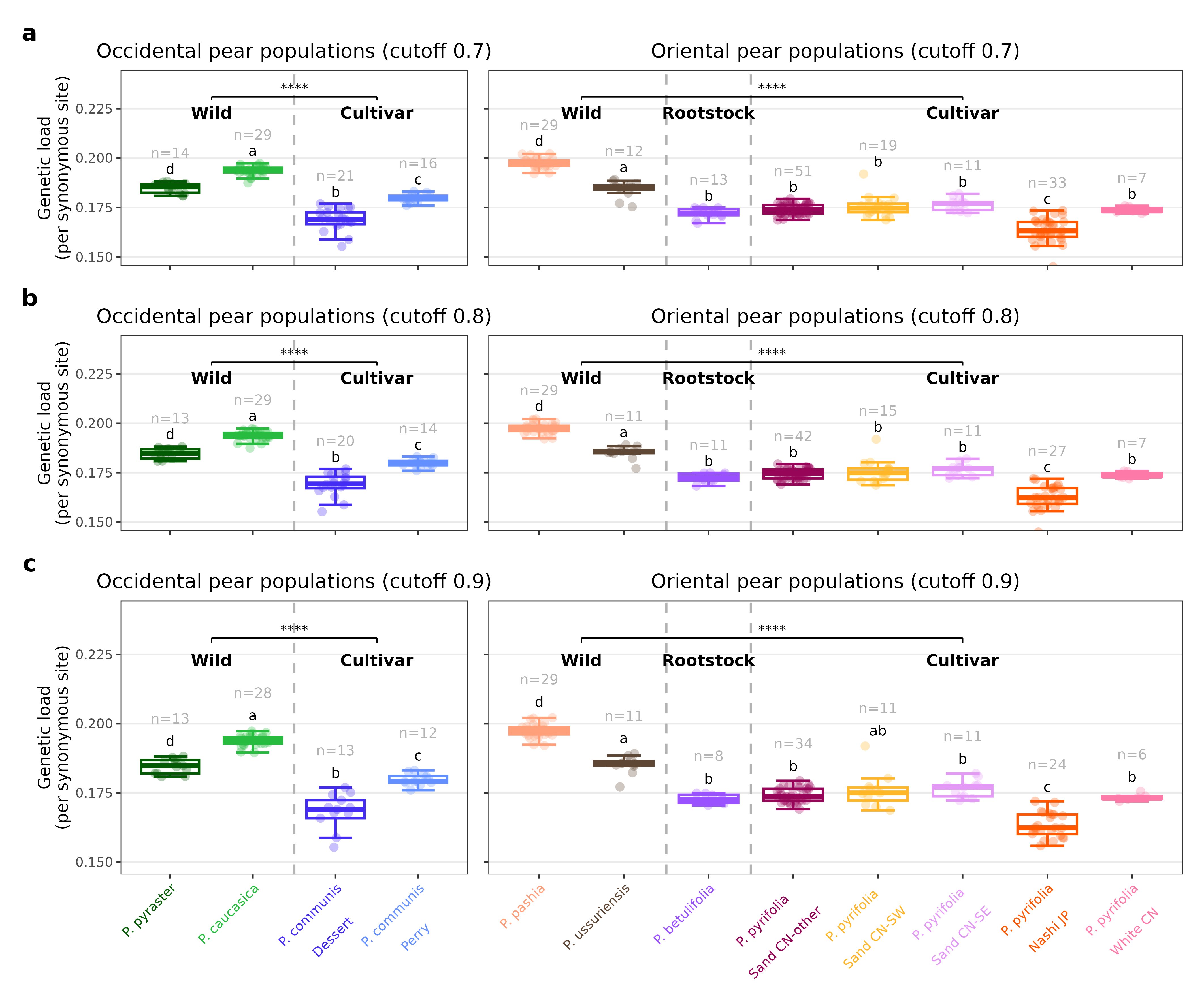

**Supplementary Figure 65 | Genetic load of pear populations defined with different ancestry cutoffs.**

**a–c**, Number of deleterious mutations (SIFT score < 0.05; SNPs or fixed sites) per carried synonymous site in each population. Populations were defined with ancestry cutoffs of 0.7 (a), 0.8 (b), and 0.9 (c). The 0.8 cutoff is the population set used in the main analyses (Fig. 4a). *P. communis* and *P. pyrifolia* were used as reference genomes for the Occidental and Oriental pear populations, respectively. Deleterious mutations were counted across all masked coding regions (selective sweep regions plus control regions), including both homozygous and heterozygous states. Stars above the brackets indicate significant differences between wild and cultivar populations (two-sided Wilcoxon rank-sum test). Different letters above the boxplots indicate significant differences in the number of deleterious alleles among populations (pairwise two-sided Wilcoxon rank-sum test, Holm-adjusted, *P* < 0.01). Numbers above the letters give the number of individuals in each population. Comparisons were performed separately for the Occidental and Oriental populations. Boxplot elements: center line, median; box limits, 25th and 75th percentiles; whiskers, extending to the most extreme data point within 1.5 × the interquartile range from the box; points, individual data. Significance levels: ns, *P* > 0.05 (not significant); * *P* ≤ 0.05; ** *P* ≤ 0.01; *** *P* ≤ 0.001; **** *P* ≤ 0.0001.

### Supplementary Table Legends

**Supplementary Table 1 | Quality metrics and passport information for the sequencing data used in this study.**

Summary of all *Pyrus* accessions analyzed, including sequencing quality metrics, metadata, and standardized identifiers used throughout this study.

**Headers:**

- **ShortNum,** Unique three-digit identifier assigned to each sample.
- **ID_Uniform,** Uniform identifier used across analyses and plots, coded as “AAAA_BB_CC000,” integrating species abbreviation (AAAA), sampling country code (BB; ISO 3166-1 alpha-2, “xx” = unknown), and sample-type code plus serial number (CC000). Sample-type codes: CW, cultivated Occidental; CE, cultivated Oriental; RE, rootstock Oriental; WW, wild Occidental; WE, wild Oriental.
- **ID_vcf,** Sample ID used in variant-calling (VCF) files.
- **ID_Run_Accession,** Accession number of public sequencing data.
- **ID_Original,** Sample ID reported in previous studies or by the original collector.
- **If_Kept,** Indicates whether the individual was retained for downstream analysis or removed (reason provided).
- **If_call_TIP,** Indicates whether the individual was retained for TIP calling or removed (reason provided).
- **Clean_Bases,** Total number of bases retained after quality control.
- **RawDepth,** Estimated sequencing depth, calculated as total bases divided by reference genome size (540 Mb).
- **MappedDepth (X),** Average depth of reads mapped to the reference genome (one column per reference).
- **MappedCoverage (%),** Percentage of the reference genome covered by mapped reads (one column per reference).
- **BioProject,** BioProject accession number for the sequencing dataset.
- **Resource,** Source of the sequencing data (publication, project, or sampler).
- **Accession,** ID of the plant material provided by the sampler.
- **CultivarOrCommonName,** Cultivar or common name of the individual.
- **CommonNameInChinese,** Common name in Chinese.
- **Usage,** Intended use (dessert, perry, or NA for unassigned).
- **SpeciesName,** Scientific name of the accession.
- **SpeciesName_TengSystem,** Species name following Teng’s taxonomic system (e.g., *Pyrus bretschneideri* → *Pyrus pyrifolia* White Pear Group).
- **Crop_or_Wild,** Indicates whether the individual is cultivated or wild.
- **Occidental_or_Oriental,** Classification into the Occidental or Oriental gene pool.
- **Country,** Country of sampling.
- **Country_Code_ISO_3166-1_alpha-2,** ISO 3166-1 alpha-2 code of the sampling country (“xx” = unknown).
- **City,** City or locality of sampling.
- **LATITUDE / LONGITUDE,** Geographical coordinates of the sampling location.

**Supplementary Table 2 | Population ancestry coefficients (Q values) inferred from population-structure analyses.**

Q values estimated from fastSTRUCTURE analyses using the genomes of all 396 *Pyrus* individuals mapped onto the *P. pyrifolia* ‘Cuiguan’ reference genome. Each individual is identified by standardized sample IDs and its proportional cluster membership in the full (Occidental and Oriental) datasets.

**Headers:**

**ID_vcf**, Sample identifier used in the VCF files.

**ID_uniform**, Standardized identifier used across analyses (see Supplementary Table 1).

**full_dataset_Cluster1 – full_dataset_Cluster10**, Ancestry coefficients (Q values) for ten clusters inferred from the full dataset.

**Assigned_cluster_FULL_DATASET**, Cluster with the highest Q value in the full dataset, both Occidental and Oriental.

**Occidental_dataset_Cluster1 – Occidental_dataset_Cluster9**, Ancestry coefficients for clusters within the Occidental subset.

**Assigned_cluster_Occidental_dataset**, Most likely cluster assignment for each Occidental individual.

**Oriental_dataset_Cluster1 – Oriental_dataset_Cluster10**, Ancestry coefficients for clusters within the Oriental subset.

**Assigned_cluster_Oriental_dataset_cutoff0.7,** Most likely cluster assignment with ancestry cutoff 0.7 for each Oriental individual.

**Assigned_cluster_Oriental_dataset_cutoff0.9,** Most likely cluster assignment with ancestry cutoff 0.9 for each Oriental individual.

**Assigned_cluster_Oriental_dataset**, Most likely cluster assignment for each Oriental individual.

**Pure_or_Admixed_in_separate_dataset**, Classification of each individual as pure (dominant cluster membership ≥ 0.8) or admixed in its corresponding regional dataset.

**Supplementary Table 3 | Population ancestry coefficients (Q values) of Occidental pears mapped onto the *Pyrus communis* reference genome.**

Q values estimated from fastSTRUCTURE analysis of the Occidental pear subset, using the *P. communis* ‘Bartlett DH Genome v2.0’ reference assembly. Each individual is identified by standardized sample IDs and its proportional cluster membership across the inferred genetic clusters.

**Headers:**

**ID_vcf**, Sample identifier used in the VCF files.

**ID_uni**, Standardized identifier used across analyses (see Supplementary Table 1).

**Occidental_dataset_on_P.communis_Cluster1 – Occidental_dataset_on_P.communis_Cluster15**, Ancestry coefficients (*Q* values) for 15 clusters inferred from the Occidental dataset mapped to *P. communis*.

**Assigned_cluster_Occidental_dataset_cutoff0.7,** Cluster showing the highest *Q* value above cutoff 0.7 for each individual when mapped to *P. communis*.

**Assigned_cluster_Occidental_dataset_cutoff0.9,** Cluster showing the highest *Q* value above cutoff 0.9 for each individual when mapped to *P. communis*.

**Assigned_cluster_mapped_onto_P.communis**, Cluster showing the highest *Q* value for each individual when mapped to *P. communis*.

**pure_or_admixed_when_mapped_onto_P.communis**, Classification of each individual as pure (dominant cluster membership ≥ 0.8) or admixed in the *P. communis*-based analysis.

**Assigned_cluster_mapped_onto_P.pyrifolia**, Most likely cluster assignment for the same individuals when mapped to the *P. pyrifolia* reference genome.

**pure_or_admix_when_mapped_onto_P.pyrifolia**, Classification of each individual as pure or admixed in the *P. pyrifolia*-based analysis.

**Supplementary Table 4 | Mapping of sample identifiers to population assignments.**

Correspondence between sample IDs and population designations used throughout this study, linking individuals to their population codes, names, and color schemes.

**Headers:**

**ID_vcf**, Sample identifier used in the VCF files.

**ID_uni**, Standardized identifier used across analyses (see Supplementary Table 1).

**Population_ID**, Abbreviated population code used in analysis scripts and figures.

**Population_name,** Full name of the population.

**Population_type**, Population category (e.g., cultivar, rootstock, or wild).

**Assigned_population_color_code**, Default color code used for consistent visualization of populations across figures.

**Related_fastStructure_cluster**, Cluster assignment corresponding to the results of fastSTRUCTURE analysis.

**Population_ID_when_ancestry_cutoff_0.7,** Population code used in analysis scripts and figures, assigned to samples with an ancestry cutoff of 0.7. The suffix “-C7” denotes this threshold.

**Population_ID_when_ancestry_cutoff_0.9,** As above, with an ancestry coefficient threshold of 0.9 (suffix “-C9”).

**Supplementary Table 5 | Results of the *D*-statistic and *f*_4_-ratio tests for introgression among Occidental pear populations using Oriental pears as an outgroup.**

Results of ABBA–BABA (*D*-statistic) and *f_4_*-ratio analyses used to test for gene flow between Occidental pear populations, with Oriental populations serving as the outgroup. The significance of the *D*-statistic was evaluated using a standard block-jackknife procedure.

**Supplementary Table 6 | Results of the *D*-statistic and *f*_4_-ratio tests for introgression among Oriental pear populations using Occidental pears as an outgroup.**

Results of ABBA–BABA (*D*-statistic) and *f*_4_-ratio analyses used to test for gene flow between Oriental pear populations, with Occidental populations serving as the outgroup. The significance of the *D*-statistic was evaluated using a standard block-jackknife procedure.

**Supplementary Table 7 | Demographic simulation parameters inferred from fastsimcoal2 analyses.**

Best-fit demographic parameters estimated for both Oriental and Occidental datasets.

Each entry corresponds to the best run (lowest AIC) selected from 50 independent fastsimcoal2 runs (100,000 simulations per likelihood estimate) within a given round, as defined by the models described in Supplementary Figures 30 and 31.

Parameters include effective population sizes, divergence times, and migration rates inferred under the selected demographic scenarios.

**Headers:**

**Lineage,** Oriental or Occidental.

**Round_ID,** Identifier of the simulation round; each round represents a distinct set of demographic models differing in population composition, topology, and migration events (gene flow).

**N_Scenarios,** Number of scenarios in this round.

**Best_Scenario,** Name of the best-fitting demographic model based on AIC. Model structures are illustrated in Supplementary Figures 30 and 31.

**MaxEstLhood,** Maximum estimated log_10_ likelihood of the observed site-frequency spectrum under the best-fit model.

**MaxObsLhood,** Maximum possible log_10_ likelihood reached if the expected site-frequency spectrum matched the observed spectrum.

**AIC,** Akaike Information Criterion, calculated as

$$AIC=2k-2\left( \frac{\text{MaxEstLhood}}{\log_{10} e} \right)$$

where k is the number of free parameters; a lower AIC indicates a better model fit.

**Simulation_Parameter**, Model parameter name. Parameters starting with N denote effective population sizes (*N*_e_), those with T represent divergence times to the present (in generations), and those with M indicate migration rates between the corresponding populations.

**Simulation_Value,** Estimated value of each parameter in the best-fitting run (model units).

**Interpretation_of_Simulation_Parameter**, Definition of each parameter.

**Dominant_Gene_Flow_Direction,** For each pair of reciprocal migration rates, the direction with the higher point estimate.

**CI_Lower_95pct**, Lower bound of the 95% confidence interval (2.5th percentile of the bootstrap estimates).

**CI_Upper_95pct,** Upper bound of the 95% confidence interval (97.5th percentile of the same bootstrap estimates).

**Supplementary Table 8 | List of candidate genes under positive selection identified in Occidental pear populations.**

List of genes located within candidate genomic regions under positive selection in Occidental pear populations. Each gene entry includes its genomic position, detection methods, annotation details, and predicted functions.

**Supplementary Table 9 | List of candidate genes under positive selection identified in Oriental pear populations.**

List of genes located within candidate genomic regions under positive selection in Oriental pear populations. The table summarizes gene positions, detection methods, and predicted functions.

**Supplementary Table 10 | GO enrichment analysis of candidate genes in selective-sweep regions shared among or specific to Occidental pear populations.**

Results of Gene Ontology (GO) term enrichment analysis for candidate genes in selective sweep regions in Occidental pear populations, showing shared and population-specific candidate genes.

**Supplementary Table 11 | GO enrichment analysis of candidate genes in selective-sweep regions shared among or specific to Oriental pear populations.**

Results of GO term enrichment analysis for candidate genes in selective-sweep regions in Oriental pear populations, showing shared and population-specific candidate genes.

**Supplementary Table 12 | KEGG enrichment analysis of Occidental and Oriental populations.**

Results of Kyoto Encyclopedia of Genes and Genomes (KEGG) pathway enrichment analysis for candidate genes in selective-sweep regions in both Occidental and Oriental pear populations.

**Supplementary Table 13 | Local gene-flow statistic outliers overlapping positive-selection candidate genes in Occidental pears.**

Window-based *D*-statistics from Dsuite Dinvestigate (50-SNP windows, 25-SNP step) for the trio (((P1, P2), P3), O), tested for local introgression overlapping genes under shared positive selection. Positive-selection candidates were defined from *ω* (OmegaPlus) and Tajima’s *D* (pixy); “shared” means selected in both named populations.

**Header definitions:**

**Trio / P1 / P2 / P3,** the three ingroup populations of the tested topology (((P1, P2), P3), O). Introgression is evaluated between P3 and P2 (positive statistic) or between P3 and P1 (negative *f*_dM_).

**chr / windowStart / windowEnd**, chromosome and start/end coordinates (bp, reference numbering) of the sliding window.

***D***, Patterson's *D* for the window.

**f_d**, *f*_d_ admixture proportion; one-tailed, higher positive values indicate gene flow between P2 and P3.

**f_dM**, *f*_dM_ statistic; two-tailed, positive indicates gene flow between P2 and P3, negative between P1 and P3.

**If_top_5_percent_f_d_outlier,** logical; TRUE if the window falls in the top 5% of the genome-wide *f*_d_ distribution.

**f_d_outlier_overlap_selection_gene_ID**, for *f*_d_ outlier windows, comma-separated IDs of shared-selection genes (P2 and P3) overlapping the window; NA if the window is not an *f*_d_ outlier or overlaps no such gene.

**If_top_2.5_percent_f_dM_outlier**, logical; TRUE if the window is in the top 2.5% of *f*_dM_.

**f_dM_top_outlier_overlap_selection_gene_ID**, for top *f*_dM_ outliers, IDs of shared-selection genes (P2 and P3) overlapping the window; NA otherwise.

**If_bottom_2.5_percent_f_dM_outlier**, logical; TRUE if the window is in the bottom 2.5% of *f*_dM_.

**f_dM_bottom_outlier_overlap_selection_gene_ID**, for bottom *f*_dM_ outliers, IDs of shared-selection genes (P1 and P3) overlapping the window; NA otherwise.

**Supplementary Table 14 | Local gene-flow statistic outliers overlapping positive-selection candidate genes in Oriental pears.**

Window-based *D*-statistics from Dsuite Dinvestigate (50-SNP windows, 25-SNP step) for the trio (((P1, P2), P3), O), tested for local introgression overlapping genes under shared positive selection. Positive-selection candidates were defined from *ω* (OmegaPlus) and Tajima's *D* (pixy); "shared" means selected in both named populations.

**Supplementary Table 15 | Enrichment of local gene-flow (*f*_d_/*f*_dM_) outliers within selective-sweep regions.**

Tests whether windows with outlier *f*_d_ or *f*_dM_ values are over-represented in selective-sweep regions shared by the two populations, whose allele sharing the outlier tail measures.

**Header descriptions:**

**lineage,** Occidental (*P. communis* reference) or Oriental (*P. pyrifolia* reference).

trio, Dinvestigate trio P1_P2_P3.

**P1,** population P1.

**P2,** population P2.

**P3,** population P3.

**statistic,** introgression statistic: *f*_d_ or *f*_dM_ (Dsuite Dinvestigate, 50-SNP windows, step 25).

**tail,** outlier tail: f_d top 5% (one-tailed); f_dM top 2.5% (P2-P3 sharing) or bottom 2.5% (P1-P3 sharing).

**gene_flow_tested,** pair whose allele sharing the tail measures.

**sweep_regions,** sweep regions tested: intersection of the two populations' sweep BEDs.

**n_windows_total,** windows in the test universe (≥ 80% of the window within callable regions, i.e., GenMap PASS with CentIER-inferred centromeres removed; valid statistic).

**n_outlier_windows,** outlier windows in the universe.

**n_windows_in_sweep,** windows overlapping the shared sweep regions.

**n_permutations,** circular permutations (rotations of sweep intervals) used by every permutation test; minimum p = 1/(n+1).

**uncontrolled_outliers_in_sweep_observed, outlier** windows inside shared sweeps.

**uncontrolled_outliers_in_sweep_expected,** mean outlier count inside sweeps over permutations (no covariates).

**uncontrolled_fold_enrichment,** observed/expected.

**uncontrolled_z,** (observed - expected) / SD of the permutation null.

**uncontrolled_perm_p,** one-sided circular-permutation p, outlier count.

**uncontrolled_perm_p_BH,** BH-adjusted within lineage.

**uncontrolled_meanstat_in_minus_out_observed,** mean statistic inside minus outside sweeps (*f*_dM_ sign flipped for the bottom tail).

**uncontrolled_meanstat_in_minus_out_expected,** same difference, mean over permutations.

**uncontrolled_meanstat_z,** z of the mean difference against the permutation null.

**uncontrolled_meanstat_perm_p,** one-sided circular-permutation p, mean difference.

**uncontrolled_meanstat_perm_p_BH,** BH-adjusted within lineage.

**uncontrolled_fisher_n_windows_thinned,** non-overlapping windows used for Fisher.

**uncontrolled_fisher_outlier_in_sweep,** 2x2 cell: outlier, inside sweep.

**uncontrolled_fisher_outlier_outside_sweep,** 2x2 cell: outlier, outside sweep.

**uncontrolled_fisher_nonoutlier_in_sweep,** 2x2 cell: non-outlier, inside sweep.

**uncontrolled_fisher_nonoutlier_outside_sweep,** 2x2 cell: non-outlier, outside sweep.

**uncontrolled_fisher_OR,** Fisher odds ratio, thinned windows.

**uncontrolled_fisher_p,** one-sided Fisher p.

**uncontrolled_fisher_p_BH,** BH-adjusted within lineage.

**pi_adjusted_outliers_in_sweep_expected,** outliers expected inside sweeps from the covariate-only logistic model (sum of fitted probabilities).

**pi_adjusted_obs_over_exp,** observed / model-expected outliers inside sweeps.

**pi_adjusted_excess_obs_minus_exp,** observed - model-expected outliers inside sweeps (test statistic U).

**pi_adjusted_perm_p,** one-sided circular-permutation p of U (main pi-controlled test).

**pi_adjusted_perm_p_BH,** BH-adjusted within lineage.

**pi_adjusted_perm_p_two_sided,** two-sided circular-permutation p of U.

**pi_adjusted_perm_null_excess_median,** median of U over permutations.

**pi_adjusted_perm_null_excess_mean,** mean of U over permutations.

**pi_adjusted_perm_null_excess_sd,** SD of U over permutations.

**pi_adjusted_perm_n_rotations_without_sweep_window,** rotations that hit no window (U = 0, kept).

**pi_adjusted_logit_complete_separation,** TRUE if no outlier inside sweeps: in-sweep coefficient and Wald p not interpretable.

**pi_adjusted_logit_converged,** glm converged.

**pi_adjusted_logit_coef_in_sweep,** log-odds coefficient of in_sweep in the full model.

**pi_adjusted_logit_se_in_sweep,** SE of the in_sweep coefficient.

**pi_adjusted_logit_OR_in_sweep,** exp(coefficient of in_sweep).

**pi_adjusted_logit_wald_p_in_sweep,** Wald p of in_sweep; assumes independent windows (overlapping windows: anticonservative), use the permutation p.

**pi_adjusted_logit_coef_log_pi_P1,** coefficient of log pi in P1 (pixy on the same windows).

**pi_adjusted_logit_coef_log_pi_P2,** coefficient of log pi in P2.

**pi_adjusted_logit_coef_log_pi_P3,** coefficient of log pi in P3.

**pi_adjusted_logit_coef_log_window_length,** coefficient of log window length (SNP density = 50/length, collinear, not modeled).

**pi_adjusted_logit_coef_gene_density,** coefficient of gene density.

**pi_matched_n_windows_thinned,** non-overlapping (thinned) windows available for matching.

**pi_matched_n_sweep_windows,** thinned windows inside sweeps.

**pi_matched_n_sweep_matched_full,** sweep windows with 5 controls within the caliper (0.10 on log scale).

**pi_matched_n_sweep_matched_partial,** sweep windows with 1-4 controls.

**pi_matched_n_sweep_unmatched,** sweep windows without any control (excluded from the test).

**pi_matched_n_controls,** matched control windows (no replacement).

**pi_matched_sweep_outlier,** 2x2 cell: matched sweep windows that are outliers.

**pi_matched_sweep_nonoutlier,** 2x2 cell: matched sweep windows that are not outliers.

**pi_matched_control_outlier,** 2x2 cell: control windows that are outliers.

**pi_matched_control_nonoutlier,** 2x2 cell: control windows that are not outliers.

**pi_matched_outlier_rate_sweep,** outlier rate, matched sweep windows.

**pi_matched_outlier_rate_control,** outlier rate, controls (pooled).

**pi_matched_outlier_rate_control_weighted,** outlier rate, controls weighted so each sweep window counts once.

**pi_matched_outlier_rate_all_sweep_windows,** outlier rate, all thinned sweep windows incl. unmatched.

**pi_matched_outlier_rate_nonsweep_windows,** outlier rate, all thinned non-sweep windows.

**pi_matched_OR, Fisher odds ratio,** matched sweep vs pooled controls (not stratified by match set).

**pi_matched_OR_ci95_low,** 95% CI lower bound.

**pi_matched_OR_ci95_high,** 95% CI upper bound.

**pi_matched_fisher_p,** one-sided Fisher *P*.

**pi_matched_fisher_p_BH,** BH-adjusted within lineage.

**pi_matched_fisher_p_two_sided,** two-sided Fisher *P*.

**Supplementary Table 16 | Per-individual deleterious mutation burden.**

Summary of deleterious allele counts and zygosity information for each individual across populations. These values underlie Fig. 4.

**Header descriptions:**

**Sample_ID,** Sample identifier.

**Population,** Population code, as in Supplementary Table 4.

**Allelic_state,** Zygosity class. Hom, homozygous for the non-reference allele; Het, heterozygous; Total, Hom + Het.

**Deleterious_allele_count,** Number of non-reference alleles at nonsynonymous sites predicted deleterious by SIFT4G (score < 0.05; low-confidence predictions excluded). A homozygous genotype contributes two alleles, and a heterozygous genotype one.

**Deleterious_per_synonymous_SNP**, Deleterious_allele_count divided by Synonymous_SNP_count.

**Region_type**, Set of coding sequences analyzed. all_masked, CDS retained after masking; selection, CDS overlapping the population’s selective-sweep regions; control, CDS outside these regions.

**Synonymous_SNP_count**, Number of synonymous sites at which the individual carries at least one non-reference allele, within the same region.

**Supplementary Table 17 | Results of statistical tests for population- and group-level comparisons of genetic load underlying Fig. 4.**

Tests underlying Fig. 4 and Supplementary Figs. 52 and 53; burden values are provided in Supplementary Table 16. Populations within each lineage were compared pairwise using two-sided Wilcoxon rank-sum tests, with Holm adjustment across all pairs within each lineage (compact letter display at adjusted *P* < 0.01). Wild and cultivated groups were compared by pooling individuals from the wild and cultivated populations of each lineage (excluding *P. betulifolia* rootstock), without adjustment. Selection versus control regions and homozygous versus heterozygous burden were compared within each population using two-sided paired Wilcoxon signed-rank tests, with Holm adjustment across populations within each lineage.

**Column descriptions:**

**Figure_panel,** Figure panel to which the test applies.

**Lineage,** Occidental or Oriental.

**Population,** Populations compared (pairwise and pooled tests) or population tested (paired tests).

**Comparison,** Type of comparison.

**Group1, Group2,** Groups compared.

**Test,** Statistical test.

**P_value,** Unadjusted *P* value.

**P_adj,** Holm-adjusted *P* value.

**Adj_method,** Adjustment method and the set of tests within which it was applied.

**Supplementary Table 18 | Quality-control analyses of the deleterious mutation burden: sequencing depth, genotype missingness, SIFT coverage, callable CDS length, and realized versus potential burden.**

Burden is the number of deleterious alleles per carried synonymous site (Supplementary Table 16). The 11 rootstock individuals (*P. betulifolia*) are excluded throughout. Methods for each panel are given in the notes below it.

**Supplementary Table 19 | Per-individual loss-of-function (LoF) burden.**

Summary of LoF allele counts and zygosity information for each individual across populations. LoF alleles were predicted as high-confidence LoF by both VEP and SnpEff. These values underlie Supplementary Fig. 54.

**Supplementary Table 20 | Results of statistical tests for population- and group-level comparisons of LoF load.**

Populations within each lineage were compared pairwise using two-sided Wilcoxon rank-sum tests, with Holm adjustment across all pairs within each lineage (compact letter display at adjusted *P* < 0.01). Wild and cultivated groups were compared by pooling individuals from the wild and cultivated populations of each lineage (excluding *P. betulifolia* rootstock), without adjustment. Selection versus control regions and homozygous versus heterozygous burden were compared within each population using two-sided paired Wilcoxon signed-rank tests, with Holm adjustment across populations within each lineage.

**Supplementary Table 21 | Overlap between TEs and genes under positive selection identified in Occidental pear populations.**

Overlap analysis of transposable element (TE) insertions and genes located within genomic regions under positive selection in Occidental pear populations. The table summarizes TE annotations, overlap statistics, and corresponding gene information.

Column descriptions are provided on the next page.

**Supplementary Table 22 | Overlap between TEs and genes under positive selection identified in Oriental pear populations.**

Overlap analysis of TE insertions and genes located within genomic regions under positive selection in Oriental pear populations. The table summarizes TE annotations, overlap statistics, and corresponding gene information.

**Supplementary Tables 21 and 22 column descriptions:**

Each row corresponds to one TE insertion polymorphism. When several values are possible (e.g., several genes or GO terms), they are separated by commas. “NA” means no information.

**TE presence by population**

**Population name,** TRUE/FALSE, TE is present in the population in at least one sample

**Genomic coordinates and variant information**

**CHROM**, chromosome or scaffold

**POS**, genomic position of the TE insertion polymorphism

**END**, end coordinate of the TE insertion polymorphism

**SVLEN**, length of the TE insertion polymorphism

**ID**, unique identifier of the TE insertion polymorphism

**Samples and populations**

**samples**, list of samples carrying the TE (comma-separated)

**populations**, list of populations corresponding to the samples

**nb_TEs**, number of TE consensus sequences associated with this insertion (data from MEGAnE output)

**consensus**, TE consensus ID(s)

**TE classification**

**class**, TE class (e.g., I, II), possibly multiple values

**order**, TE order (e.g., LTR, TIR), possibly multiple values

**family**, TE family, possibly multiple values

**Gene annotation**

**gene_ID**, gene identifier(s) whose 2 kb upstream region contains the TE insertion

**gene_Start**, gene start coordinate(s)

**gene_End**, gene end coordinate(s)

**gene_Strand**, gene strand(s)

**gene_Symbol**, gene symbol(s), if available

**positive_selection**, TRUE/FALSE indicating whether the gene is under positive selection

**Functional annotation**

**GO_ID**, Gene Ontology term(s) associated with the gene(s)

**Ath_description**, functional description based on Arabidopsis annotation

**Curated TE classification**

**class_curated**, dominant TE class

**order_curated**, dominant TE order

**family_curated**, dominant TE family

**Recombination context**

**Start_Rho**, start coordinate of the recombination window

**End_Rho**, end coordinate of the recombination window

**Rho**, recombination rate value

**TE density context**

**Start_TEcov**, start of the genomic window used for TE density

**End_TEcov**, end of the genomic window used for TE density

**NB_TE_OVERLAP**, number of TEs overlapping the window

**BASES_COVERED**, number of bases covered by TEs in the window

**WINDOW_SIZE**, size of the window

**TE_density**, proportion of the window covered by TEs

GO summary and categorization

**GO_Level0**, main GO domains associated with the gene(s)

**GO_Level2**, GO level-2 terms associated with the gene(s)

**GO_Level2_Category**, broad functional category inferred from GO level-2 terms

**Immunity-related annotation**

**Immunity**, TRUE/FALSE, indicates whether the gene(s) are associated with immunity-related functions

**Intersection key and summary metrics**

**_key**, binary code representing the combination of populations in which the TE is present

**TE class (%)**, within each population-combination (_key), percentage of gene-TE association rows assigned to each curated TE class

**TE order (%)**, within each population-combination (_key), percentage of gene-TE association rows assigned to each curated TE order

**GO Level 2 (%)**, within each population-combination (_key), percentage of gene-TE association rows assigned to each GO level-2 functional category

**Immunity (%)**, within each population-combination (_key), percentage of gene-TE association rows associated with immunity-related genes

**Supplementary Table 23 | Summary of TIPs associated with selected genes.**

This table provides, for each population (both Occidental and Oriental), gene-level selection statistics derived from SNP-based scans and the associated TIP counts, including the proportion of TIPs overlapping genes under positive selection.

**Supplementary Table 24 | Pairwise comparisons of TIP occurrence upstream of positively selected genes across pear populations.**

Pairwise population comparisons were performed using generalized estimating equations (GEE) adjusted for local recombination rate, TE density, gene density, and log-transformed gene length. *Lineage* indicates Occidental (PCOM) or Oriental (PPY) pears; *Gene_category* indicates all positively selected genes or the immunity-associated subset; *Population1* and *Population2* identify the compared populations, and *Group1* and *Group2* indicate their cultivated or wild status. *N_selected_genes_pop1* and *N_selected_genes_pop2* indicate the numbers of positively selected genes in each population, while *N_shared_selected_genes* indicates the number shared between them. *OR_pop1_vs_pop2* is the adjusted odds ratio for TIP presence in Population1 relative to Population2, with corresponding 95% confidence limits (*CI95_lower*, *CI95_upper*). *Log_OR* is the natural logarithm of the odds ratio, and *SE_robust* is its robust standard error. *p_value* is the two-sided Wald test p-value, and *FDR_BH* is the Benjamini–Hochberg-adjusted p-value. *Higher_TIP_odds_population* identifies the population with higher adjusted TIP odds, and *Significant_FDR05* indicates whether FDR < 0.05.

**Supplementary Table 25 | Association between transposable element insertion polymorphisms (TIPs) and positive selection across pear populations and genomic compartments.**

Associations between TIP occurrence and positive selection were assessed using Cochran–Mantel–Haenszel (CMH) tests stratified by PLM gene cluster, generalized estimating equations (GEE) adjusted for genomic covariates, and matched-control analyses. *Lineage* indicates Occidental (PCOM) or Oriental (PPY) pears; *population* identifies the population analyzed; and *compartment* indicates the genomic region examined (CDS, intron, 2-kb upstream or downstream, and 5′ or 3′ UTR in PPY only). *n_selected* and *n_other* indicate the numbers of positively selected and non-selected genes, respectively, while *n_selected_with_tip* and *n_other_with_tip* indicate the corresponding numbers containing at least one TIP. *pct_sel_with_tip* and *pct_other_with_tip* represent the percentages of genes containing TIPs in each category. *n_strata* indicates the number of PLM gene clusters included in the CMH analysis. *cmh_or*, *cmh_lo*, *cmh_hi* and *cmh_p* report the CMH odds ratio, its 95% confidence limits and associated P value. *adj_or*, *adj_lo*, *adj_hi* and *adj_p* report the GEE-adjusted odds ratio, its 95% confidence limits and P value, accounting for gene length, local TE density, gene density and recombination rate. Odds ratios compare positively selected genes with non-selected genes, with values below one indicating TIP depletion among positively selected genes. *ctrl_mean_pct*, *ctrl_lo_pct* and *ctrl_hi_pct* indicate the mean TIP frequency and its 95% empirical interval across 1,000 matched sets of non-selected genes. *perm_p* indicates the empirical P value from the matched-control analysis. *cmh_fdr*, *adj_fdr* and *perm_fdr* are the Benjamini–Hochberg-adjusted P values for the CMH, GEE and matched-control analyses, respectively, calculated separately for PCOM and PPY across population-by-compartment comparisons.

**Supplementary Table 26 | SNP datasets used throughout this study.**

For each dataset: number of SNPs and individuals, reference genome, filtering criteria, rationale for the subset and the analyses that use it.

**Supplementary Table 27 | Ancestry-cutoff sensitivity analysis of fastsimcoal2 demographic inference.**

Best-fitting models (Oriental: eastS2G1; Occidental: westG10D3) were re-estimated using samples selected with ancestry-proportion cutoffs of 0.7 (C7) and 0.9 (C9). Point estimates are compared with the 95% confidence intervals of the main analysis, obtained from 50 bootstrap replicates. Overall, 35 of 46 re-estimated parameters (76%) fell within the main-analysis CIs (Oriental 16/22; Occidental 19/24).

**Supplementary Table 28 | Callable regions retained after mappability and centromere filtering in the *Pyrus pyrifolia* and *Pyrus communis* reference genomes.**

Genomic intervals retained for all window-based analyses. Mappability profiles were generated with GenMap (*k* = 140, *e* = 0), mean mappability was computed in 100-kb windows with a 50-kb step, and windows with mean mappability > 0.9 were retained; centromeric regions inferred with CentIER were then subtracted. The resulting set comprises 302.3 Mb (59.4%) of the 17 *P. pyrifolia* chromosomes and 342.3 Mb (76.7%) of the 17 *P. communis* chromosomes; unplaced scaffolds were excluded. Coordinates follow the BED convention (0-based, half-open: the start position is included and the end position is excluded), and are given separately for *P. pyrifolia* Cuiguan and *P. communis* Bartlett DH Genome v2.0.

**Supplementary Table 29 | Fastsimcoal2 model comparison in the main demographic analysis of the Oriental and Occidental pear lineages.**

Each row is one demographic scenario tested in the main analysis (five rounds for the Oriental lineage and one for the Occidental lineage; 169 scenarios in total).

**Column descriptions:**

**lineage,** Oriental or Occidental.

**round,** Simulation round, as in Supplementary Table 7.

**scenario_id,** Scenario name.

**topology,** Population tree of the scenario in Newick format.

**topology_description,** Order of divergence events.

**wild_root_proxy,** Wild population used as the ancestral proxy (*P. pashia* for Oriental and *P. caucasica* for Occidental pears).

**gene_flow,** Migration edges included in the scenario (source -> recipient, forward in time); none, no gene flow.

**dominant_gene_flow,** For each pair of reciprocal migration rates, the direction with the higher point estimate.

**MaxEstLhood, AIC,** As in Supplementary Table 7.

**ΔAIC,** AIC difference from the best scenario of the round.

**best_scenario,** “yes” for the lowest-AIC scenario of the round.

**Supplementary Table 30 | Sensitivity of Oriental fastsimcoal2 model selection to the choice of ancestral proxy (*P. pashia* vs *P. ussuriensis*).**

Rounds 2, B2, and C1 of the main analysis (Supplementary Tables 7 and 29) were repeated with *P. ussuriensis* replacing *P. pashia* as the wild ancestral proxy (Round_ID suffix “ussu”). AIC values are comparable only within a round, because the two proxies yield different site-frequency spectra.
